# Experimental trophic flexibility guides a compartmentalized genome-scale model of red microalgae *Galdieria sulphuraria*

**DOI:** 10.64898/2026.10.15.759342

**Authors:** Agris Pentjuss, Sergejs Kolesovs, Mara Grube, Rita Scerbaka, Karlis Svirksts, Egils Stalidzans, Armands Vigants, Sergiy Smetana, Daniel Pleissner

## Abstract

The thermoacidophilic red microalga *Galdieria sulphuraria* combines broad organic-carbon utilization with biotechnological robustness. We integrated cultivation data for strain SAG 107.79 with an organelle-resolved genome-scale metabolic model describing autotrophic, mixotrophic and heterotrophic metabolism. Heterotrophic cultures produced 2.29, 2.03 and 2.14 g L⁻¹ biomass on glucose, xylose and their mixture, respectively, while mixed-sugar cultures consumed glucose before xylose. Biomass composition was similar under mixotrophy and heterotrophy, and stationary-phase glucose consumption supported a non-growth-associated maintenance demand of 1.7 mmol ATP gDW⁻¹ h⁻¹. These measurements informed trophic-state-specific biomass objective functions. The final model contains 2,487 reactions, 2,270 compartment-specific metabolites and 1,050 genes. The model reproduced the experimentally scored growth phenotypes, highlighted polyol-associated product capacity under heterotrophy and scored 90% in the archived MEMOTE report. To our knowledge, this is the first experimentally based, organelle-resolved genome-scale metabolic model for *G. sulphuraria*, providing a quantitative framework linking trophic state and substrate identity with carbon use, flux flexibility and product capacity. A user-friendly supplementary package provides the executable SBML model, modular biomass resources, reproducible scripts, quality reports and underlying experimental data.

## Introduction

Microalgae are increasingly considered as promising sources of nutrients, pigments, lipids and other high-value compounds, and several species are being explored as novel food and biotechnology platforms. Their industrial application depends on understanding how cultivation conditions, nutrient availability and trophic state influence growth, biomass composition and product formation. Successful *in silico* modelling of such biotechnological processes, including cultivation, fermentation and physiological adaptation, therefore requires accurate representations of cellular metabolism. Genome-scale metabolic models provide a framework for connecting this cellular metabolism with environmental and process conditions.

Systems biology and genome-scale metabolic modelling link genotype, metabolic capacity and environmental conditions in microalgal biotechnology [1]. A genome-scale model (GEM) combines reaction stoichiometry, gene-protein-reaction associations and physicochemical constraints, while expression data can support condition-specific interpretation [2, 3]. In microalgae, applications include biomass and lipid partitioning in *Phaeodactylum tricornutum* [4], TAG-engineering strategies in *Nannochloropsis salina* [5], and EPA and fatty-acid production for scalable cultivation [6]. Metabolic engineering has also enhanced omega-3 EPA in *Nannochloropsis* [7] and ketocarotenoids in *Chlamydomonas reinhardtii* [8]. Beyond microalgae, model-based approaches have addressed acetaldehyde and ethanol production [9, 10], amino-acid and other industrial bioproduct pathways [11, 12], and biotechnological potential in *Zymomonas mobilis* [13, 14], *Kluyveromyces marxianus* [15], *Saccharomyces cerevisiae* [16] and *Pichia pastoris* [17].

For microalgae, realistic compartmentation is important because carbon, cofactors, reducing equivalents and biosynthetic precursors are distributed among organelles. In *C. reinhardtii*, iRC1080 resolved chloroplast and thylakoid light metabolism, while iCre1355 expanded gene-reaction coverage across autotrophic (AUTO), mixotrophic (MIXO) and heterotrophic (HETERO) conditions [18, 19]. The six-compartment *Chlorella vulgaris* model iCZ843 predicted phenotypes across trophic modes, and iAJ526 extended compartmentalized modelling to *Chlorella variabilis* biofuel potential [20, 21]. Organelle-resolved models also improved analysis of lipid, energy and redox metabolism: iLB1027_lipid for *P. tricornutum* contained 4,456 reactions across six compartments and integrated localization prediction with measured biomass composition [4]. *Nannochloropsis* models iRJ1321 and iNS934 addressed trophic flexibility, nitrogen metabolism and lipid/TAG production [5, 22, 23], while iTps1432 for *Thalassiosira pseudonana* represented organelle organization for photosynthetic and redox analysis [24]. More recently, iMgadit23 refined ER and plastid glycerolipid metabolism in *Microchloropsis gaditana* and improved lipid-mutant predictions [25].

Genome-scale resources for red microalgae remain limited. CcrGEM reconstructed metabolism of the red macroalga *Chondrus crispus* [26], while EctoGEM for the brown macroalga *Ectocarpus siliculosus* provides a comparative algal resource [27]. Neither represents the distinctive thermoacidophilic and trophically flexible physiology of unicellular *Galdieria sulphuraria*. In *G. sulphuraria*, photosynthetic and respiratory metabolism operate across AUTO, MIXO and HETERO states, together with broad organic-carbon utilization and trophic-state-dependent storage and lipid metabolism [28, 29, 30].

*G. sulphuraria, a unicellular* red microalga from the class Cyanidiophyceae, stands out as an extremophile with exceptional promise for these purposes. It is well known for thriving under extreme conditions, including highly acidic environments (pH 1–5) and temperatures reaching 56 °C [31]. Because such conditions suppress the growth of most contaminating organisms, this species is especially well-suited for large-scale, non-sterile industrial processes that rely on complex organic waste streams ([32]; [33]; [34]; [35]; [36]). Furthermore, its remarkable metabolic versatility allows it to grow on a wide range of organic substrates, including those originating from aquaculture waste [37].

The biochemical makeup of *G. sulphuraria* biomass further underscores its potential value. The alga can accumulate over 30% (w/w) protein with a well-balanced amino acid composition, making it highly suitable for use in animal feed and human nutrition [36, 38, 39]. It also synthesizes considerable amounts of highly branched glycogen along with the blue pigment phycocyanin – a compound already commercially recognized as a natural colorant and increasingly utilized in the food and nutraceutical sectors [40, 41]. Collectively, these attributes make *G. sulphuraria* an outstanding candidate for the bio-based transformation of aquaculture residues into high-value products (HVP), supporting environmental remediation, economic valorization, and the advancement of circular resource systems.

Our cultivation experiments provided the biological basis for genome-scale reconstruction. Under HETERO conditions, glucose supported greater biomass and biomass yield than xylose and was consumed first in mixed-sugar cultures. MIXO and HETERO biomass composition was broadly similar, whereas stationary-phase glucose consumption under HETERO conditions provided an organism-specific maintenance constraint. These observations raised the question whether glucose-first utilization reflected metabolic efficiency and motivated an organelle-resolved *G. sulphuraria* model with measured MIXO/HETERO composition incorporated into trophic-state-specific biomass objectives. Standardized growth-capacity and fixed-growth comparisons confirmed a modest glucose advantage, while HETERO minimum-carbon optimization selected glucose under the tested mixed-sugar condition. Together, these analyses distinguish substrate feasibility from preference and quantify how trophic state changes carbon use, rather than treating all growth-supporting substrates as metabolically equivalent. The model was then extended to broader substrate utilization, trophic-state-dependent carbon allocation and HVP potential using metabolic modeling methods like Flux Balance Analysis (FBA) and Flux Variability Analysis (FVA).

## Results

### Growth on glucose, xylose and mixed sugars

Initial experiment tested whether *G. sulphuraria* uses a hexose and a pentose equally well. HETERO screening compared 5 g L⁻¹ glucose, 5 g L⁻¹ xylose and a 2.5 + 2.5 g L⁻¹ glucose– xylose mixture. Final biomass concentrations were 2.29 ± 0.04, 2.03 ± 0.04 and 2.14 ± 0.04 g L⁻¹ (dry weight), and the corresponding biomass yields on consumed substrate were 0.442 ± 0.006, 0.397 ± 0.008 and 0.417 ± 0.008 g g⁻¹, respectively. This confirmed that *G. sulphuraria* used both hexose and pentose carbon under organic-carbon-supported HETERO growth, with glucose resulting in the highest biomass and yield (p<0.001), while xylose alone and the mixed feed both supported substantial biomass formation. These results established glucose and xylose uptake as experimentally supported model phenotypes rather than annotation-derived assumptions. Individual measurements, including replicate counts and summary statistics are provided in Supplementary Data 7.

### Mixed-sugar kinetics reveal ordered carbon-source utilization

(11E) The mixed-sugar kinetic experiment independently showed rapid growth but resolved the order of substrate depletion. By day 4, mean glucose had reduced from 2.51 to 0.28 g L⁻¹ in MIXO cultures and to 0.80 g L⁻¹ in HETERO cultures, whereas xylose remained near 2.35–2.37 g L⁻¹. By day 5, both sugars were below the recorded detection level and biomass reached 2.12 g L⁻¹ under MIXO and 1.83 g L⁻¹ under HETERO conditions. Log-linear fits over days 1–5 gave apparent rates of 1.23 d⁻¹ (R² = 0.94) and 1.44 d⁻¹ (R² = 0.96), respectively. The data therefore support preferential glucose use followed by rapid xylose utilization, rather than simultaneous depletion at equal rates. The underlying biomass and sugar-concentration measurements are provided in Supplementary Data 7.

An additional glucose-supported kinetic test provided orthogonal optical-density and cell-count measurements. Over days 3–5, OD750-derived apparent growth rates were 1.48 d⁻¹ for MIXO and 1.65 d⁻¹ for HETERO cultures, while cell-count-derived values were 1.54 and 1.65 d⁻¹, respectively. The agreement between the two measurement types supports the occurrence of a rapid exponential phase, while the sparse sampling limits precision. Underlying OD750 and cell-count measurements, including the HET cell-count rate, are provided in Supplementary Data 7.

### Biomass composition is dominated by protein

When *G. sulphuraria* biomass was sampled on day 5, the measured composition was similar under MIXO and HETERO conditions. Protein represented 41.47 ± 1.54% and 40.16 ± 0.62% of dry biomass, carbohydrate 33.77 ± 0.37% and 36.09 ± 1.36%, nucleic acids 7.75 ± 0.51% and 7.65 ± 0.47%, and lipids 7.01 ± 0.65% and 6.10 ± 0.28%, respectively. The measured categories summed to 90% in each condition, leaving a 10% residual pool that may include ash and unmeasured material. These measurements define a common experimental core for the two organic-carbon-supported biomass functions, but they do not by themselves justify coefficients for the unmeasured pool. Biological-replicate values and summary statistics are provided in Supplementary Data 7.

### Estimation of non-growth associated maintenance from stationary-phase glucose consumption

In half-strength ACM containing approximately 18 g L⁻¹ glucose, biomass increased to 5.12 ± 0.05 g L⁻¹ by day 7, while glucose declined to 8.68 ± 0.02 g L⁻¹. Between days 7 and 8, glucose decreased by 1.25 g L⁻¹, whereas biomass changed from 5.12 ± 0.05 to 4.96 ± 0.12 g L⁻¹ (Fig. 4). Because no positive biomass formation was observed, this interval was treated as an apparent stationary-phase maintenance period.

**Figure 1.**
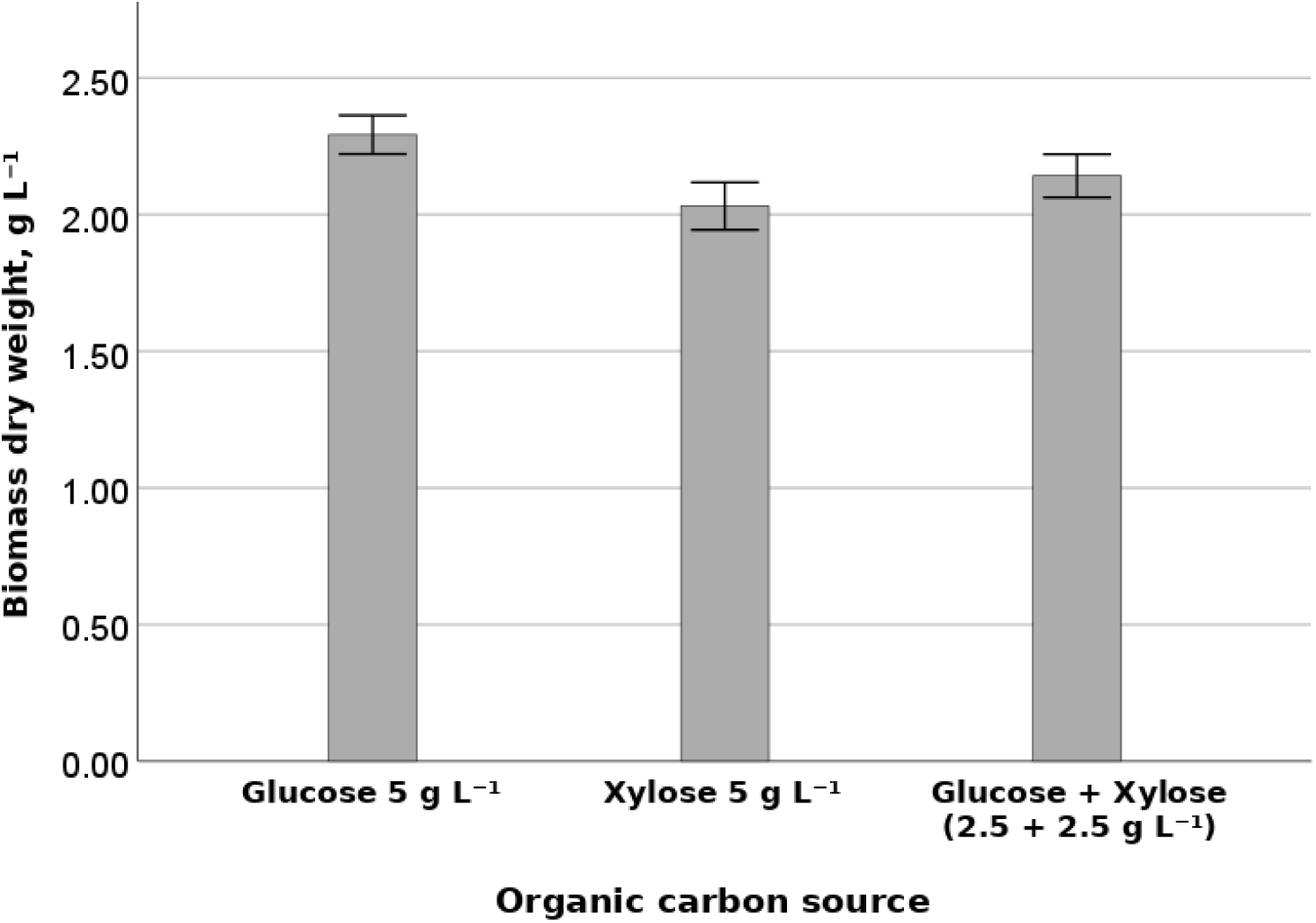
G. sulphuraria biomass dry weight after 7 days of HETERO cultivation in ACM containing 5 g L⁻¹ glucose, 5 g L⁻ ¹ xylose, or 2.5 + 2.5 g L⁻ ¹ glucose and xylose (n = 4 for glucose; n = 5 for xylose and the mixture). Source data are provided in Supplementary Data 7.

**Figure 2.**
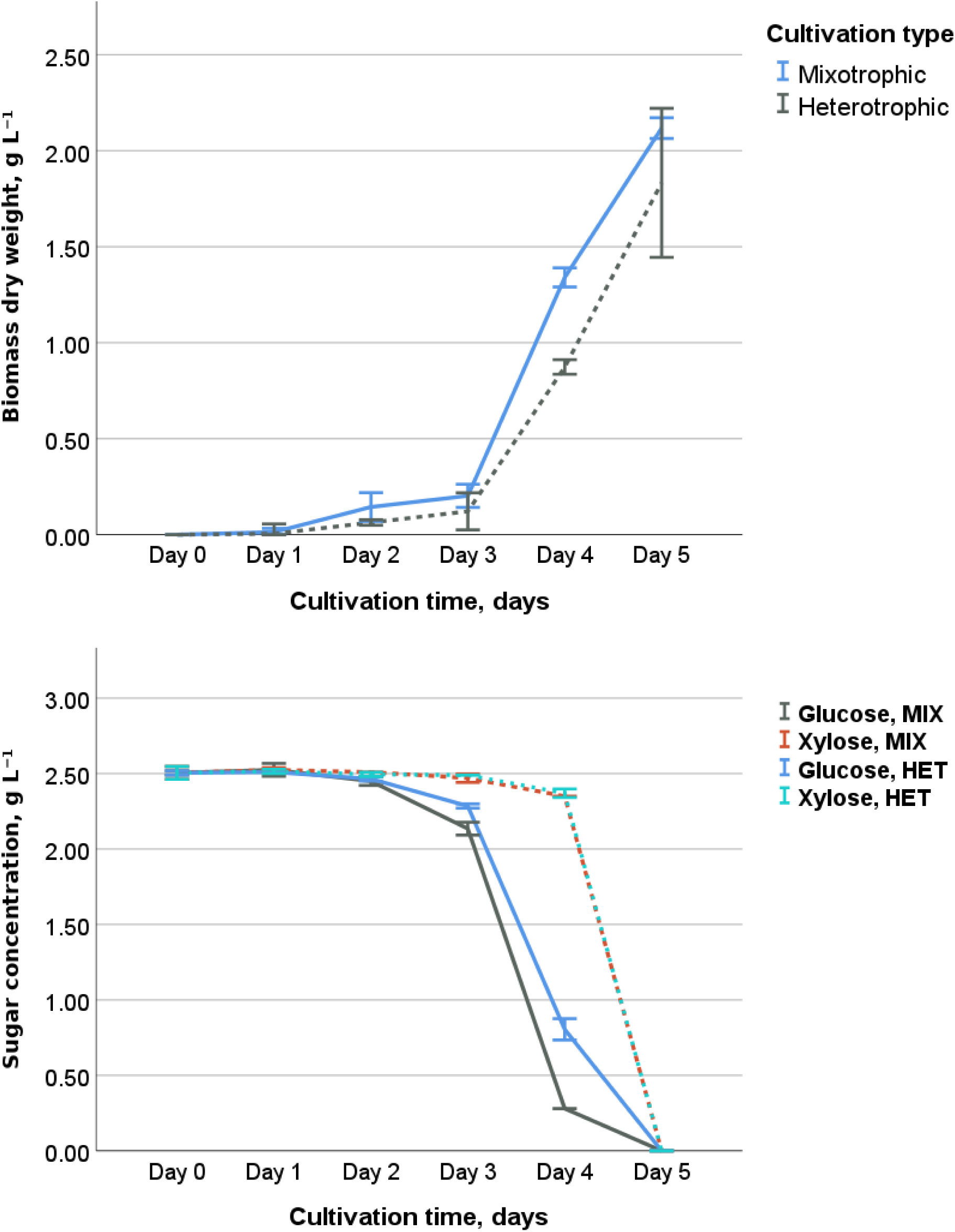
a) G. sulphuraria biomass synthesis over 5 days of MIXO or HETERO cultivation in ACM containing 2.5 + 2.5 g L⁻¹ glucose and xylose (n = 3); b) Changes in sugar concentrations (xylose and glucose) over 5 days of mixotrophic (MIX) or heterotrophic (HET) cultivation in ACM containing 2.5 + 2.5 g L⁻¹ glucose and xylose (n = 2). Source data are provided in Supplementary Data 7.

**Figure 3.**
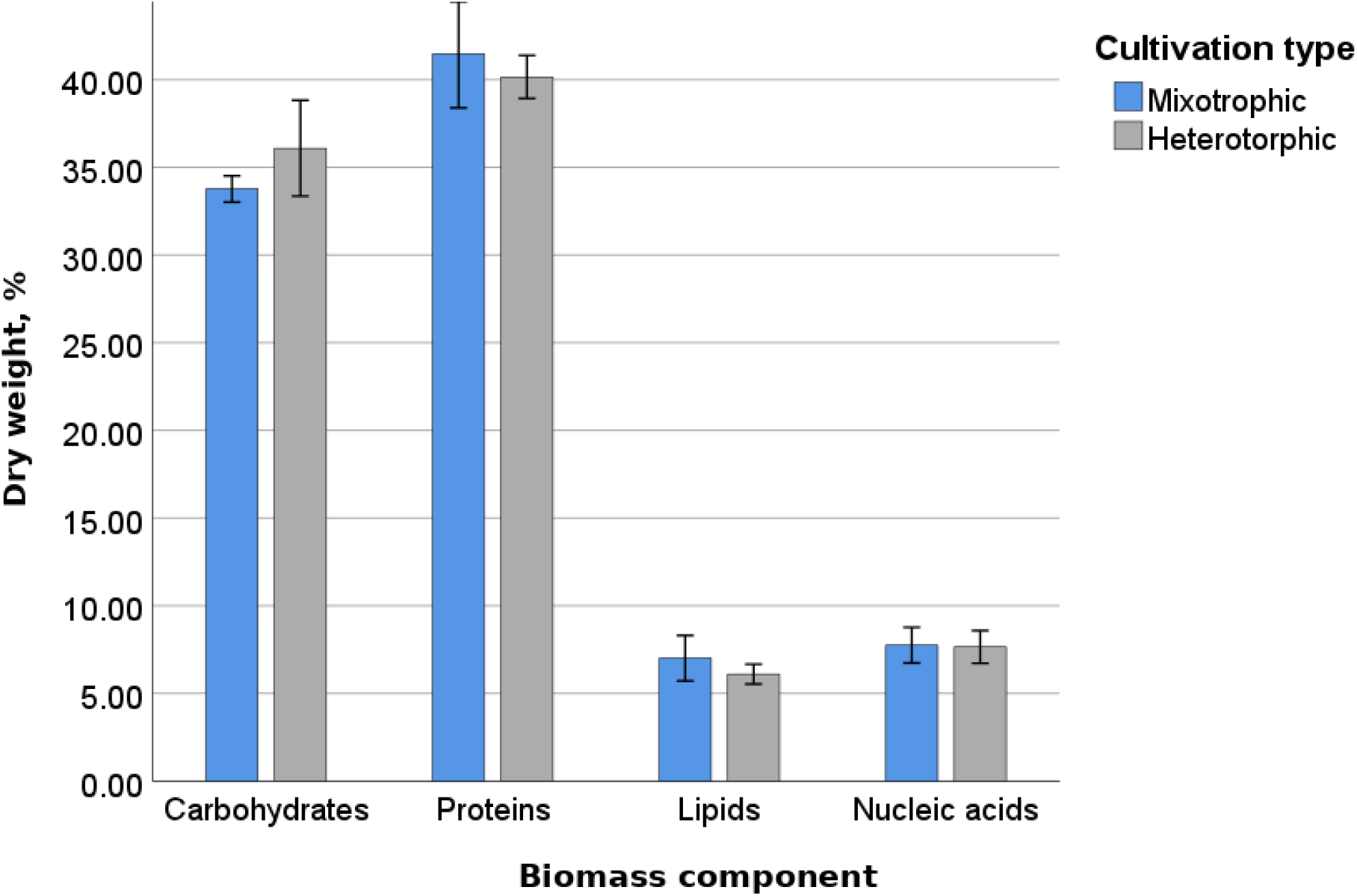
G. sulphuraria biomass composition on day 5 under MIXO and HETERO conditions (n = 2 biological replicates). Values are percentages of biomass dry weight. Source data are provided in Supplementary Data 7.

**Figure 4.**
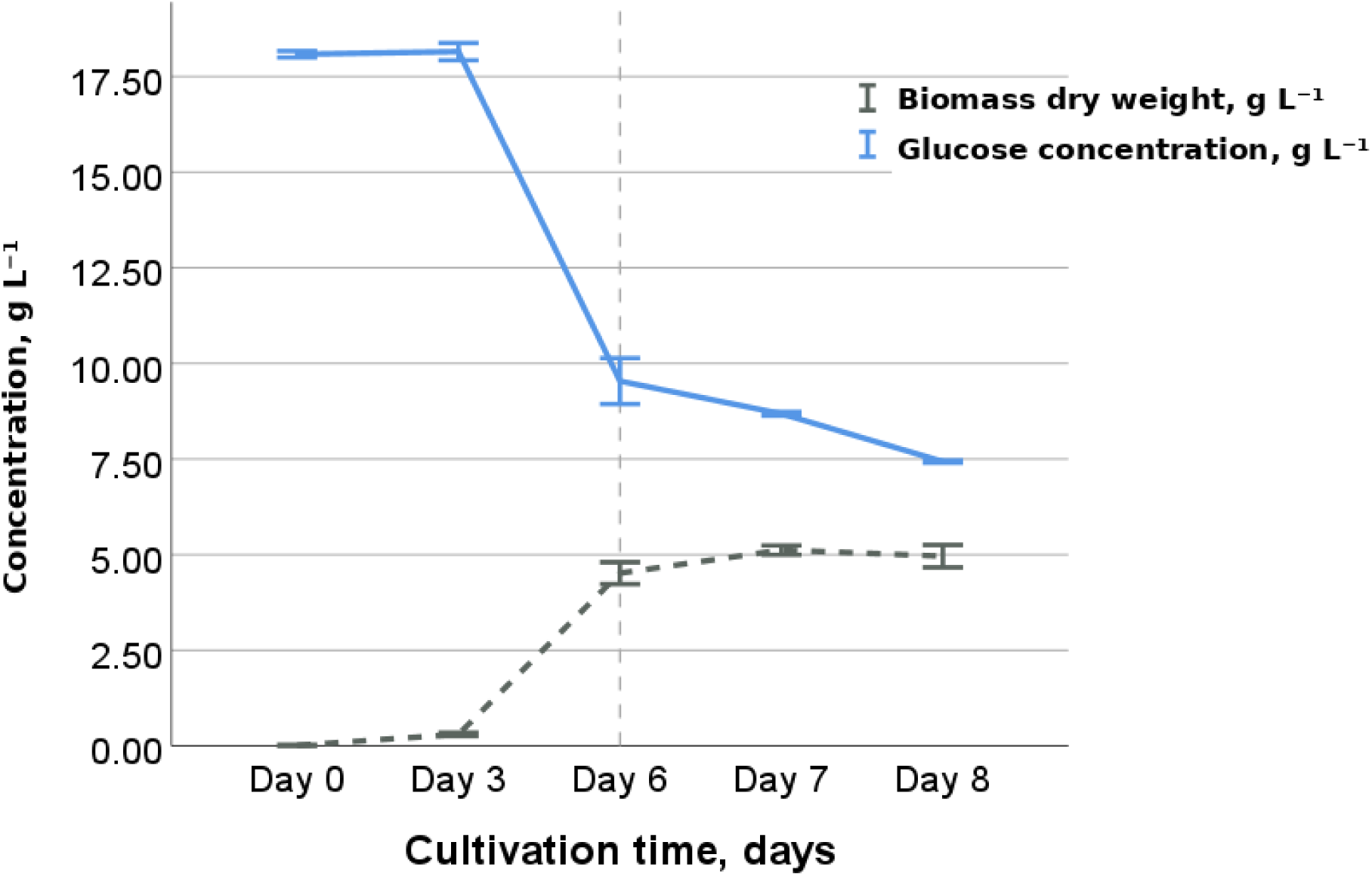
Changes in biomass synthesis (g L⁻¹) and glucose concentrations (g L⁻¹) during 8 days of HETERO cultivation on twice-diluted ACM with 18 g L⁻¹ glucose (n≥3). Source data are provided in Supplementary Data 7

The day-7 to day-8 glucose decrease was 1.2503 g L⁻¹ d⁻¹ and mean biomass was 5.04 gDW L⁻¹. These values give an unrounded maintenance-associated glucose demand of 0.2481 g gDW⁻¹ d⁻¹ (0.01034 g gDW⁻¹ h⁻¹; 0.05738 mmol gDW⁻¹ h⁻¹) (Formula 1). For the reported ATP calibration in Supplementary Data 7, the daily rate was rounded to 0.25 g gDW⁻¹ d⁻¹, equivalent to 0.05782 mmol glucose gDW⁻¹ h⁻¹. An aerobic yield of 26–30 mmol ATP per mmol glucose was assumed.

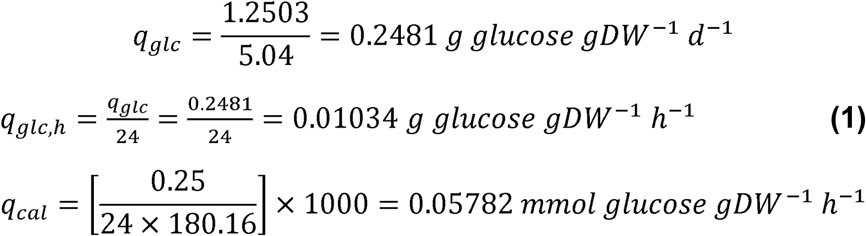

The ATP-based non-growth-associated maintenance (NGAM) estimate uses the rounded calibration rate and the assumed aerobic ATP yield (Formula 2):

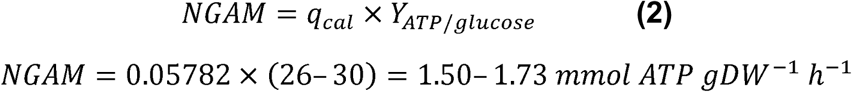

The assumed ATP yield from glucose (26-30 mmol ATP per mmol glucose) gives an NGAM, i.e. the maintenance energy requirement, of ≈1.5-1.7 mmol ATP gDW⁻¹ h⁻¹. This is a model-dependent estimate rather than a direct ATP measurement because the conversion depends on the respiratory ATP yield represented by the genome-scale model. GAM cannot be calculated from this stationary-phase interval because no positive biomass formation occurred. The underlying measurements and formula-based calculation chain are provided in Supplementary Data 7.

### Genome-scale reconstruction and compartmentation

The reconstructed *G. sulphuraria* GEM contains 2,487 reactions, 2,270 compartment-specific metabolites and 1,050 genes distributed across cytosolic, plastid, thylakoid, mitochondrial and peroxisomal metabolism, together with explicit transport and exchange reactions. Compartment assignments represent plastid/thylakoid photosynthetic and redox processes and mitochondrial energy metabolism [28], cytosolic storage-carbon and organic-substrate metabolism [29, 42], plastid carbon transport and fatty-acid precursor supply [43], photorespiration [28] and lipid metabolism [29]. Experimentally supported pathways that were incomplete or absent from the initial genome-derived set were restored by evidence-based gap filling. Reaction-level compartmentation, pathway assignment and supporting evidence are provided in Supplementary Data 5 together with the final executable SBML model.

### Model quality assessment with MEMOTE and comparison with published algal GEMs

The final reconstruction contains 2,487 reactions, 2,270 compartment-specific metabolites and 1,050 genes. Its MEMOTE report scored 90.182% overall and 99.500% for consistency; 49 reactions were mass-unbalanced and 28 were charge-unbalanced (Supplementary Data 5–6). To conserve symbolic photon stoichiometry, the final model uses Photon_USED_Plas and Photon_USED_Thyl pseudo-metabolites. Light-dependent reactions produce the compartment-matched carrier, which is removed through a demand reaction. In PSII, four Photon_Thyl units produce four Photon_USED_Thyl units without changing the biochemical chemistry; the same principle was applied to photosystem I and plastid protochlorophyllide oxidoreductase. For comparison, reported overall MEMOTE scores for selected photosynthetic algal GEMs include iNS934 (18%), iRJ1321 (25.3%), the AuCoMe *Microchloropsis gaditana* reconstruction (17.7%) and iMgadit23 (67%) (Supplementary Data 6). The report and comparator workbook describe the final SBML model.

### Condition-specific biomass objective functions

Separate biomass objective functions were implemented for AUTO, MIXO and HETERO growth. Each BOF contains protein, carbohydrate, lipid, RNA, DNA and pigment fractions, with trophic-state-specific coefficients retained where supporting measurements were available. Supplementary Data 2 retains the literature-informed reference reactions and relative molecular compositions; the reported MIXO and HETERO experimental BOFs are assembled from these proportions by the scripts in Supplementary Data 3.

Protein was treated as a condition-dependent biomass component. Literature-supported BOF ranges were 0.61–0.62, 0.56–0.59 and 0.25–0.42 g gDW⁻¹ for AUTO, MIXO and HETERO growth, respectively. The substantially broader HETERO interval reflects substrate- and nutrient-dependent variation reported for *G. sulphuraria* [36, 37, 38, 44, 45].

Carbohydrate fractions were likewise trophic-state dependent, with BOF ranges of 0.16–0.23, 0.20–0.25 and 0.10–0.36 g gDW⁻¹ for AUTO, MIXO and HETERO states, respectively. The broad HETERO interval reflects direct observations ranging from carbohydrate-poor to strongly storage-rich biomass depending on substrate and cultivation regime [36, 37, 45].

The pigment BOFs contained chlorophyll a, zeaxanthin, β-carotene and phycocyanobilin (PCB), with pigment investment decreasing from AUTO to MIXO and HETERO states [38, 46]. Only the PCB chromophore was represented from phycocyanin because its apoprotein contribution was already included within the protein fraction, avoiding double counting.

Lipids were represented using trophic-state-specific total lipid contents together with detailed fatty-acid and membrane-lipid composition rather than a single generic lipid pool. Stable lipid was partitioned between membrane lipids and TAG using AUTO/MIXO/HETERO allocations of 0.75:0.25, 0.65:0.35 and 0.50:0.50; these exact ratios are modelling assumptions, whereas the experimentally observed direction of trophic membrane/storage remodeling is supported by [29] and [47]. The membrane fraction was further resolved into MGDG, DGDG, SQDG, PG, PE, PC, PI, PS and DGTS, with condition-specific fatty-acid profiles used to obtain molecular-species BOF coefficients.

RNA and DNA were represented separately using genome-derived nucleotide composition, while common RNA and DNA mass fractions were applied across trophic states because condition-specific measurements were unavailable. Importantly, maintenance demand was additionally informed by the stationary-phase experiment performed in this study. The observed glucose consumption corresponded to an apparent ATP-maintenance demand of approximately 1.7 mmol ATP gDW⁻¹ h⁻¹, and NGAM = 1.7 mmol ATP gDW⁻¹ h⁻¹ was therefore implemented as a separate ATP-maintenance constraint rather than incorporated into biomass stoichiometry.

Reference biomass reactions and the integration guide are provided in Supplementary Data 2; the scripts that apply measured MIXO and HETERO fractions are provided in Supplementary Data 3.

### Literature growth-rate evidence defines the validation envelope

The curated *G. sulphuraria* specific-growth-rate dataset comprised 135 condition-specific literature records: 45 AUTO records spanning μ = 0.128–0.500 d⁻¹ (0.0053–0.0208 h⁻¹), 10 MIXO records spanning 0.228–1.490 d⁻¹ (0.0095–0.0621 h⁻¹), and 80 HETERO records spanning 0.020–1.530 d⁻¹ (0.0008–0.0638 h⁻¹). Because studies differed in strain, substrate, pH, temperature and cultivation system, these ranges were used as condition-specific validation envelopes rather than pooled estimates of a species-wide μmax.

Records included primary measurements, doubling-time-derived values, operational steady-state estimates and secondary extractions. Row-level values, uncertainty provenance and source mapping are provided in Supplementary Data 4.

### Model evaluation against cultivation experiments

All seven experimental substrate-condition tests and eight quantitative growth measurements were feasible *in silico*. Experimental growth rates were imposed as biomass fluxes, and the minimum organic-carbon requirement needed to sustain each rate was calculated. At the HETERO KIN2 reference growth rate (μ = 0.060 h⁻¹), the minimum carbon demand was 3.223 C-mmol gDW⁻¹ h⁻¹. Detailed results are provided in Supplementary Data 3.

### Sole substrate validation

The final model was evaluated across 29 experimentally scored substrate entries, representing 28 unique model exchange conditions (Supplementary Data 3). All scored growth phenotypes were reproduced under both HETERO and MIXO conditions. Starch was treated as an intracellular storage compound and excluded from qualitative sole-substrate phenotype scoring. The CO₂-only AUTO reference supported a maximum growth rate of 0.041 h⁻¹ at a photon uptake limit of 20 mmol gDW⁻¹ h⁻¹.

At 6 C-mmol gDW⁻¹ h⁻¹ organic-carbon supply, MIXO maximum growth exceeded HETERO capacity for the representative feasible substrates. The absolute growth-capacity predictions and differences from HETERO are shown in Table 1; these are constrained model capacities.

**Table 1.** Representative substrate comparisons under standardized model constraints and literature-supported experimental evidence.

| Substrate | Experimental evidence | Reference | HETERO theoretical $\mu_{\max}$ (h<sup>-1</sup>) | HETERO C demand at $\mu=0.060$ (C-mmol gDW<sup>-1</sup> h<sup>-1</sup>) | MIXO theoretical $\mu_{\max}$ (h<sup>-1</sup>) | CO<sub>2</sub> contribution in MIXO (%) |
| --- | --- | --- | --- | --- | --- | --- |
| D-Glucose | $\mu = 1.10 \pm 0.03$ d<sup>-1</sup>; $t_d = 50$ h | [46, 49] | 0.12 (reference) | 3.2 (reference) | 0.25 (+112.6%) | 35.9 |
| D-Xylose | $t_d = 282$ h; glucose specific growth rate reported 51% higher than xylose | [48, 49] | 0.11 (-5.8%) | 3.4 (+5.4%) | 0.25 (+123.6%) | 35.3 |
| meso-Erythritol | Growth-positive in the 27-carbon-source screen | [42] | 0.13 (+10.6%) | 2.9 (-9.5%) | 0.26 (+99.2%) | 38.1 |
| Glycerol | $\mu = 1.08 \pm 0.03$ d<sup>-1</sup>; doubling time = $0.64 \pm 0.02$ d | [46] | 0.13 (+8.9%) | 2.96 (-8.3%) | 0.26 (+99.9%) | 37.4 |
| Acetate | Controlled acetic-acid feeding: biomass productivity = $1.18 \pm 0.05$ g L<sup>-1</sup> d<sup>-1</sup>; | [48, 50] | 0.08 (-31.0%) | 4.5 (+39.8%) | 0.21 (+161.1%) | 24.4 |
| Pyruvate | pathway/regulatory evidence only | [51] | 0.08 (-29.9%) | 4.5 (+38.6%) | 0.22 (+170.3%) | 28.0 |

At μ = 0.060 h⁻¹, xylose required 5.4% more carbon than glucose, whereas meso-erythritol and glycerol required 9.5% and 8.3% less. Acetate and pyruvate required approximately 39% more carbon, showing that growth-positive substrates were not equivalent in carbon demand.

Glycerol supported HETERO growth with a predicted maximum growth rate of 0.129 h⁻¹. At μ = 0.060 h⁻¹, its feasible uptake range was 0.985–1.034 mmol gDW⁻¹ h⁻¹. Under MIXO, glycerol supported a maximum growth rate of 0.258 h⁻¹, with CO₂ accounting for the substrate-specific fraction of total external carbon reported in Table 1. Experimental HETERO growth on glycerol has been reported at μ = 1.08 ± 0.03 d⁻¹, with a doubling time of 0.64 ± 0.02 d [46].

The predicted HETERO growth capacity on D-xylose was 5.836% lower than on D-glucose (0.112 versus 0.119 h⁻¹). The direction agrees with experimental observations for *G. sulphuraria* UTEX 2919, where specific growth on glucose was reported to be 51% higher than on xylose [48]. Earlier substrate-specific doubling times are reported in [49].

Acetate utilization is supported experimentally under controlled cultivation conditions, including a biomass productivity of 1.18 ± 0.05 g L⁻¹ d⁻¹ during acetic-acid feeding [50], while its effect depends strongly on cultivation regime and concentration [48]. Direct quantitative sole-pyruvate growth data for *G. sulphuraria* remain limited; [51] provides metabolic and regulatory context rather than a direct growth-rate measurement.

### Targeted flux variability analysis

Targeted FVA tested six representative carbon sources under HETERO conditions at μ = 0.060 h⁻¹. After minimum substrate demand was determined, uptake was limited to 105% of that value and 5–8 pathway, respiratory and biomass reactions were analysed. Intervals excluding zero identified obligatory reactions [52, 53].

All six substrates were obligatorily consumed, and their minimum carbon requirements matched the fixed-growth substrate screen. The panel covered hexose, pentose, polyol and organic-acid entry routes; numerical uptake and carbon-demand values are reported in Table 2 [42].

**Table 2.** Targeted flux variability analysis of representative carbon sources under heterotrophic growth. Biomass was fixed at μ = 0.060 h⁻¹. The 6 C-mmol gDW⁻¹ h⁻¹ organic-carbon value was used only as a maximum availability. Substrate-uptake intervals are shown as positive uptake magnitudes after constraining uptake to ≤105% of the minimum requirement. Reaction fluxes are mmol gDW⁻¹ h⁻¹. All numerical results are reported to two decimal places.

| Carbon source | Minimum substrate uptake | FVA uptake range ($\leq 105\%$ minimum) | Key obligatorily active pathway reactions | Respiratory Complex III FVA range | Complex V FVA range |
| --- | --- | --- | --- | --- | --- |
| D-Glucose | 0.54 | 0.54–0.56 | Glucokinase; enolase | 1.51–2.09 | 3.22–4.54 |
| D-Xylose | 0.68 | 0.68–0.71 | Xylose conversion; D-xylose reductase; pentose-phosphate entry | 1.82–2.46 | 3.89–5.11 |
| meso-Erythritol | 0.73 | 0.73–0.77 | D-erythrulose kinase; erythritol dehydrogenase | 1.58–2.22 | 3.53–4.62 |
| Glycerol | 0.99 | 0.99–1.03 | Glycerol kinase; glycerol-3-phosphate oxidation; enolase | 1.97–2.57 | 3.41–4.50 |
| Acetate | 2.25 | 2.25–2.37 | Acetate-CoA ligase; malate synthase; PEP carboxykinase | 4.12–4.79 | 7.76–9.07 |
| Pyruvate | 1.49 | 1.49–1.56 | Mitochondrial citrate synthase | 2.51–3.14 | 4.87–6.47 |

Fixed-growth FVA retained the main substrate-screen pattern: meso-erythritol and glycerol required less carbon than glucose [46], and constraints were concentrated at substrate entry. Entry fluxes for sugars, polyols and acetate remained obligatory, consistent with reported pentose, glycerol and acetate utilization [48, 54, 55]. Downstream central metabolism was more flexible and substrate dependent, with especially redundant pyruvate entry. Respiratory Complex III and Complex V were obligatory across all six substrates, indicating conserved respiratory and ATP-generation requirements [56]. Detailed reaction intervals are reported in Table 2 and Supplementary Data 3.

### High-value product production potential across trophic states and substrates

Nine HVP targets covered storage-carbohydrate, lipid, pigment, compatible-solute and isoprenoid metabolism: glycogen/storage α-glucan, triacylglycerol (TAG), linoleic acid (C18:2), β-carotene, phycocyanobilin (PCB), floridoside, squalene, ergosterol and phytol. State-specific analyses prioritized HETERO glycogen, TAG, floridoside and ergosterol; MIXO glycogen, TAG, C18:2, PCB and ergosterol; and AUTO glycogen, C18:2, β-carotene, PCB and phytol. Squalene remained a supplementary model-supported target. The products were selected because they are experimentally observed or biochemically supported in *G. sulphuraria* and span distinct biotechnology applications. Definitions and priorities are given in Supplementary Table S2.

The analyses separated theoretical production capacity from standardized carbon efficiency. Capacity was optimized while maintaining ≥50% of each substrate-specific μmax; carbon efficiency was evaluated at μ = 0.060 h⁻¹. Organic-carbon availability was capped at 6 C-mmol gDW⁻¹ h⁻¹ in HETERO and MIXO. Storage α-glucan was treated as an intracellular reserve rather than extracellular starch uptake. Keeping the designs separate prevents theoretical capacity from being conflated with carbon efficiency at one growth state.

### Product-specific significance and trophic-state interpretation

Glycogen/storage α-glucan. *G. sulphuraria* Glycogen has 18% α-(1→6) linkages, short chains and small particles [57], with trophic-state-dependent accumulation [29]; it has been proposed as a starch alternative for slowly digestible and resistant glucose polymers [58]

#### Triacylglycerol (TAG)

Neutral-lipid accumulation changes with trophic state in *G. sulphuraria* [29], while microalgal TAGs are renewable feedstocks for biodiesel and related fuels [59].

Linoleic acid (C18:2). C18:2 comprised 45.2 ± 2.3% of fatty acids under AUTO and 19.5 ± 2.6% under HETERO [60]; linoleic-acid-rich oils serve food, pharmaceutical and cosmetic applications [61, 62].

#### β-Carotene

In *G. sulphuraria*, β-carotene was lower under HETERO than AUTO and more strongly inhibited by glucose than glycerol [46]. Uses include natural colorants, provitamin-A supplements and nutraceuticals [63].

#### Phycocyanobilin (PCB)

*G. sulphuraria* converts phycoerythrobilin to PCB through PEB:PCB isomerase [64], and phycocyanin abundance depends on carbon source and trophic state [46]. As the phycocyanin chromophore, PCB supports natural blue-colorant and fluorescent biomolecule applications [65].

#### Floridoside

Its pathway was characterized in *G. sulphuraria* [66], and osmotic stress induced up to 56.8 mg g⁻¹ dry biomass in glycerol-grown cells [67]. Potential applications include nutraceutical, cosmetic and pharmaceutical bioactive fractions [68].

#### Squalene

No validated study directly quantified its accumulation in *G. sulphuraria*, so it remains a supplementary model-supported sterol-pathway target; established uses include cosmetic emollients and lipid-based topical formulations [69].

#### Ergosterol

Levels were higher under HETERO than AUTO [70] and increased after glucose addition and under high light conditions [71]. Fermentation-derived ergosterol is an industrial vitamin-D₂ precursor [72].

#### Phytol

Abundance was higher under AUTO than HETERO [70] and lower under MIXO than AUTO [71]. Uses include fragrances and other value-added chemicals [73].

### Primary analysis: substrate-dependent theoretical HVP production capacity

HETERO production capacity was substrate dependent (Fig. 5). Dulcitol was the best substrate for glycogen, floridoside and ergosterol, while meso-erythritol was optimal for TAG, with improvements of 8.65–12.62% relative to D-glucose. These are theoretical product-flux improvements at ≥50% substrate-specific maximum growth and not measured productivities or carbon yields [42, 54].

**Figure 5.**
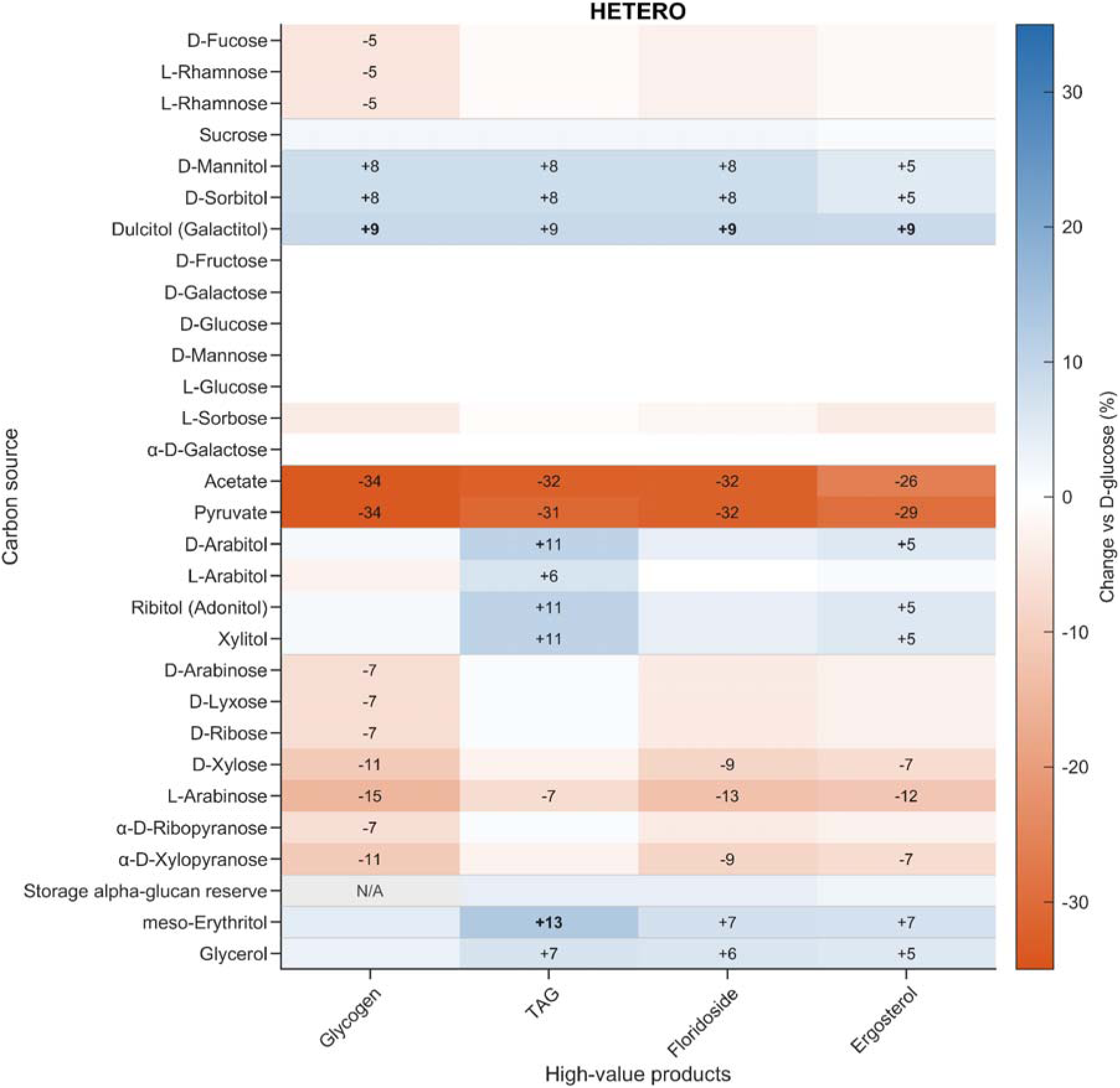
Substrate-dependent changes in theoretical HVP production capacity under HETERO conditions. Values show percentage change in optimized product flux relative to D-glucose while maintaining ≥50% of the substrate-specific maximum growth rate. Organic-carbon availability was normalized to a maximum of 6 C-mmol gDW⁻ ¹ h⁻ ¹. Numerical labels are shown for changes of ≥5% in absolute value relative to D-glucose.

MIXO responses were narrower, ranging from 1.74% to 5.18% among the five prioritized products (Table 3; Supplementary Fig. S1). In the updated simulations, meso-erythritol was optimal for PCB instead of dulcitol.

**Table 3.** Best theoretical HVP production-capacity responses relative to D-glucose. Values are product- flux capacity changes at ≥50% of each substrate-specific μmax and are not carbon yields.

| State | HVP | Best substrate | Change vs D-glucose (%) |
| --- | --- | --- | --- |
| HETERO | Glycogen | Dulcitol | 8.94 |
| HETERO | TAG | meso-Erythritol | 12.62 |
| HETERO | Floridoside | Dulcitol | 8.95 |
| HETERO | Ergosterol | Dulcitol | 8.65 |
| MIXO | Glycogen | Dulcitol | 1.74 |
| MIXO | TAG | meso-Erythritol | 3.19 |
| MIXO | Linoleic acid | meso-Erythritol | 3.30 |
| MIXO | PCB | meso-Erythritol | 2.58 |
| MIXO | Ergosterol | meso-Erythritol | 5.18 |

### Secondary analysis: standardized fixed-growth carbon efficiency

At fixed μ = 0.060 h⁻¹, polyols again gave the highest HETERO carbon yields (Supplementary Fig. S2; Supplementary Table S3). Meso-erythritol and dulcitol increased yields by 17.03– 23.11% across the four prioritized products, confirming that the advantage was not driven only by growth potential.

### MIXO performance is distinguished by carbon-source robustness and additional CO₂ assimilation

At μ = 0.060 h⁻¹, fixed-growth yields for the five prioritized MIXO products varied little across most substrates, whereas the CO₂-derived share of total assimilated carbon ranged from 19.8– 29.8%. Thus, MIXO combined robust organic-carbon conversion with substrate-dependent inorganic-carbon assimilation. Substrate identity changed how strongly inorganic carbon complemented organic carbon rather than eliminating product feasibility (Supplementary Table S4).

Relative to D-glucose, the best MIXO total-carbon yields increased only 0.639–1.832% across the five prioritized products. Both organic-C-normalized and total-assimilated-carbon yields are reported; the latter is preferred because product carbon can originate from CO₂. The complete profile is shown in Supplementary Fig. S3 and agrees with simultaneous respiratory and photosynthetic carbon metabolism in *G. sulphuraria* [28].

### AUTO production-capacity context

AUTO was evaluated separately as a CO₂-only condition and excluded from organic-substrate comparisons. Its targets were glycogen, C18:2, β-carotene, PCB and phytol. At a photon uptake limit of 20 mmol gDW⁻¹ h⁻¹, AUTO maximum growth capacity was 0.041 h⁻¹, corresponding to a ≥50% growth floor of approximately 0.020 h⁻¹. All five selected product outputs remained feasible under this light constraint

### Overall interpretation

Together, fixed-growth FVA separated obligatory substrate-entry and respiratory functions from flexible downstream metabolism, while HVP analyses distinguished trophic-state-specific production patterns. HETERO was most sensitive to substrate identity, MIXO combined more stable product yields with substrate-dependent CO₂ assimilation, and AUTO provided the CO₂-only reference for comparing production capacity and carbon efficiency across trophic contexts.

## Discussion

To our knowledge, this is the first experimentally anchored, organelle-resolved GEM for the unicellular thermoacidophilic red alga *G. sulphuraria*. The final reconstruction contains 2,487 reactions, 2,270 compartment-specific metabolites and 1,050 products across five compartments and is the model evaluated by the MEMOTE report. Explicit provenance, compartmentalization and symbolic photon bookkeeping make model assumptions auditable, while cultivation data constrain the biological interpretation (Supplementary Data 5–6).

The model was developed from cultivation phenotypes rather than genome annotation alone. Glucose supported greater biomass formation than xylose and was consumed first in mixed cultures, indicating unequal use of co-occurring substrates rather than absent xylose metabolism. This distinction matters for mixed-carbon side streams, on which *G. sulphuraria* has already been cultivated [36, 48, 74]. It is also consistent with reported glucose-mediated repression of xylose use [48, 51]. By separating feasibility from preferred carbon use, the model translates that cultivation phenotype into testable differences in carbon demand, pathway flexibility and product potential across trophic states. This provides a more useful basis for evaluating mixed feedstocks than a binary growth/no-growth classification (Supplementary Data 3).

Measured MIXO and HETERO biomass composition provided a second constraint. Because trophic state alters biomass allocation, separate AUTO, MIXO and HETERO objectives avoid biases introduced by one invariant biomass reaction [75, 76]. Condition-specific biomass functions are also used in other algal GEMs [4, 20, 77]. The reference compositions and molecular distributions in Supplementary Data 2 make these assumptions inspectable. The analysis scripts in Supplementary Data 3 apply the measured MIXO and HETERO fractions to those distributions without normalizing the measured totals to 100%.

The stationary-phase experiment supplied an organism-specific energetic anchor. Continued glucose consumption without positive biomass formation yielded a model-dependent non-growth-associated maintenance (NGAM) estimate of 1.7 mmol ATP gDW⁻¹ h⁻¹. Although this estimate depends on the assumed aerobic ATP yield and is not a direct intracellular ATP measurement, it is preferable to transferring a maintenance value from an unrelated organism. Separating NGAM from growth-associated maintenance also preserves the conceptual distinction between ATP used to sustain cells and ATP coupled to biomass formation. The experimental measurements and calculation are provided in Supplementary Data 7, while BOF and maintenance-reaction implementation remains documented in Supplementary Data 1–2.

The complementary analyses show that substrate feasibility, preference, carbon demand and product capacity are distinct properties. All experimentally scored substrate conditions in the archived analysis were feasible. The HETERO mixed-sugar minimum-carbon solution selected glucose, while the separate sole-substrate screen found both sugars feasible. MIXO increased growth capacity across the HETERO feasible substrate panel. At fixed growth, carbon demand still differed systematically among sugars, polyols and organic acids, so growth-positive feedstocks were not metabolically equivalent. Capacity and fixed-growth screens therefore provide complementary criteria for feedstock selection (Supplementary Data 3–4).

Targeted FVA identified the essential substrate-entry and respiratory functions and the flexibility of downstream metabolism. Entry constraints depended on substrate, whereas central pathways retained alternative feasible states; respiratory Complex III and Complex V remained required throughout. Agreement between optimization and FVA supports the inferred phenotypes without treating one optimum as unique [52, 53]. The flexible regions identify where uptake, secretion or isotope-labelling measurements would most improve model resolution (Supplementary Data 3).

High value products (HVP) predictions were more substrate sensitive in HETERO, where polyols generally favored glycogen, TAG, floridoside and ergosterol. MIXO responses were more robust and included inorganic-carbon assimilation, consistent with light-driven metabolism buffering organic-substrate effects. Agreement between capacity and fixed-growth analyses supports these qualitative differences without relying on maximum growth alone. These interpretations agree with experimental evidence for trophic-state-dependent storage, lipid, compatible-solute and sterol metabolism in *G. sulphuraria* [29, 66, 67, 70, 71]. The predictions identify candidate substrate–product pairs without substituting for measured titres or productivities (Supplementary Data 3).

Integration of quantitative omics was outside this study’s scope. Relevant *G. sulphuraria* transcriptomic, metabolomic and isotope-labelling datasets exist [28, 46, 71] and could support condition-specific refinement. Quantitative proteomics, uptake/secretion measurements and dedicated 13C-flux experiments would provide stronger constraints. Improved annotation, orthology, transporter assignment and localization would strengthen GPR coverage; expression-constrained, thermodynamic, dynamic and enzyme-constrained modelling could improve physiological realism ([2]; [3]; [12]; Supplementary Data 5).

*G. sulphuraria* combines acidothermophilic growth, broad substrate use and trophic flexibility with demonstrated applications in wastewater and residue valorization, protein-rich biomass, glycogen, lipids, phycocyanin and floridoside production [29, 36, 39, 44, 48, 67, 74, 78, 79]. The principal contribution is a compartmentalized GEM integrating trophic-state-specific biomass functions, maintenance demand, physiological validation, substrate-resolved growth, FVA and HVP optimization. The model, provenance, validation and optimization resources are distributed in Supplementary Data 1–7. These materials make the reconstruction reusable beyond the supplied SBML model and allow future experimental constraints or alternative objectives to be introduced transparently. By linking trophic state and organic-substrate identity to carbon efficiency, flux flexibility and product capacity, the model provides a common framework for hypothesis generation in algal systems biology, computational modelling, bioprocess engineering and circular-bioeconomy research.

Together, the experimental data showed that glucose supported higher HETERO biomass formation and yield than xylose (2.29 vs 2.03 g L⁻¹; 0.442 vs 0.397 g g⁻¹), while mixed-sugar cultures preferentially consumed glucose first. In the proposed model, HETERO polyols increased theoretical HVP capacity by 8.645–12.619% relative to glucose, whereas the corresponding MIXO gains were smaller (1.738–5.182%). Overall, the *G. sulphuraria* model links trophic state and substrate identity to carbon efficiency, flux flexibility, growth, and high-value product production potential. It provides a transparent and reusable framework for evaluating mixed-carbon side streams, identifying promising substrate–product combinations, guiding targeted experiments, and supporting future applications in algal biotechnology, bioprocess engineering, and circular bioeconomy research.

## Methods

### Strain and cultivation media

*G. sulphuraria* strain SAG 107.79 (SAG Culture Collection of Algae, Göttingen University, Germany) was maintained under axenic conditions. Allen’s Cyanidium Medium ACM [80] was prepared using reagents provided by Sigma-Aldrich (St. Louis, MO, USA). The medium pH was lowered to 4 prior to any cultivation, using 1 M HCl. MIXO and HETERO cultures used ACM supplemented with 5 g L⁻¹ glucose, xylose 5 g L⁻¹, and 2.5 + 2.5 g L⁻¹ mixture of glucose and xylose as the organic carbon source. For maintenance energy cost determination, cultivation was conducted HETERO in twofold-diluted ACM with 18 g L⁻¹ glucose in order to accelerate the transition to stationary phase.

### Experimental design

The trials were carried out in 50 mL Erlenmeyer flasks containing 25 mL medium and 10% inoculum (approximately 5 × 10 cells mL⁻¹ at day 0) in a Biosan orbital shaker-incubator ES-20 at (Biosan Ltd., Riga, Latvia) at 55 ± 2 μmol photons m⁻² s⁻¹ and 42°C, with 5, 7, or 8 cultivation days based on the trial. Prior to that, the inoculum was prepared photo AUTO for 10 days, under the same cultivation conditions. For HETERO cultivation, flasks were wrapped in aluminum foil to exclude light and incubated alongside the MIXO cultivated flasks and photoautotrophic controls.

### Biomass dry weight, productivity, and yield calculations

Dry weight determination was performed as follows: The culture volume was adjusted for evaporation using deionized water, after which a 20 mL sample of *G. sulphuraria* culture was centrifuged at 5000 rpm for 2 min, and the supernatant was discarded. The pellet was then rinsed with deionized water to remove residual medium components. The centrifugation and rinsing steps were repeated twice. Subsequently the concentrated samples were then transferred to pre-weighed weighing bottles, dried at 105°C for 24 h, and used for gravimetric determination of dry weight using an analytical balance. Biomass dry weight was determined by subtracting the mass of the empty (pre-weighed) weighing bottle from the mass of the weighing bottle containing the dried biomass. The resulting value was converted from g per 20 mL to g L⁻¹. Additionally, following volume adjustments for evaporation, assessment of cell count was conducted using a haemocytometer and a light microscope at ×400 magnification for additional control of *G. sulphuraria* growth.

Where sugar consumption data were available (HETERO cultivation), the biomass yield (Formula 3) from the consumed substrate (Y, g g⁻¹) was calculated using the following equation:

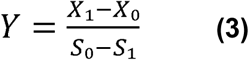

where X₀ and X₁ represent the biomass dry weight (g L⁻¹) at the beginning and end of the experiment, respectively, and S₀ and S₁ represent the sugar concentration (g L⁻¹) at the beginning and end of the experiment.

### Defined-sugar batch screens

*G. sulphurari*a was cultivated under MIXO and HETERO conditions for 7 d. One screen compared glucose and xylose as separate carbon sources. A second HET screen compared 5 g L⁻¹ glucose, 5 g L⁻¹ xylose and 2.5 g L⁻¹ glucose plus 2.5 g L⁻¹ xylose.

### Growth-kinetics experiments

Two mixed-sugar kinetic experiments used 2.5 g L⁻¹ glucose plus 2.5 g L⁻¹ xylose under MIXO and HET conditions. Biomass was sampled from day 0 to day 8 in the first series and from day 0 to day 5 in the second. Glucose and xylose were recorded separately in the second series. A complementary glucose-supported experiment measured dry biomass, OD750 and cell abundance during days 3–5 under MIXO and HETERO conditions, with a photo AUTO day-5 reference. Apparent specific growth rates were calculated by linear regression of ln(X) against time over the stated interval, where X was dry biomass, OD750 or cell count. Two-point rates are explicitly identified as interval-specific estimates.

### Biomass-composition and Sugar uptake measurements

For FTIR spectroscopy analyses *G. sulphuraria* biomass samples were collected on day 5 from late-exponential MIXO and HETERO cultures and triple-rinsed with distilled water. FTIR spectra were recorded using a Vertex 70 coupled with an HTS-XT microplate reader (Bruker, Germany). Sample aliquots of 10 μl were pipetted on a 384-well ZnSe microplate in at least 3 replicates. Absorption spectra were recorded in the frequency range of 4000–600 cm-1 with a spectral resolution of 4 cm -1 and 64 scans added. Only spectra within the absorption limits between 0.25 and 0.80 (where according to the Lambert–Bouger–Beer law, the concentration of a component is proportional to the intensity of the absorption band) were used for data analysis. Spectra were vector-normalized and data processed with Opus 7.5 (Bruker Optics GmbH, Ettlingen, Germany). The baseline of each spectrum was corrected using the rubber band method. The macromolecular composition of *G. sulphuraria* biomass was calculated as described previously [81]. The content of carbohydrates, nucleic-acids, proteins and lipids was expressed as percent of dry weight. The four reported macromolecular classes summed to 90%; no unmeasured fraction was imputed in the experimental results.

Glucose concentrations were determined by HPLC using an Agilent 1100 Series system (Santa Clara, California, USA) equipped with a Shodex SH1011 column (300 mm × 7.8 mm) and a refractive index detector. The column temperature was maintained at 50 °C. The mobile phase consisted of 0.005 mol/L H_2_SO_4_, with a flow rate of 0.6 mL/min and an injection volume of 5 µL. Xylose concentrations were determined by HPLC using an Agilent 1100 Series system (Santa Clara, California, USA) equipped with a Shodex SP0810 column (300 mm × 7.8 mm) and a refractive index detector. The column temperature was maintained at 80 °C. The mobile phase consisted of deionized H_2_O, with a flow rate of 0.6 mL/min and an injection volume of 5 µL. Compounds were identified by comparison with retention times of external standards. Quantification was performed automatically using Agilent ChemStation software based on calibration curves prepared from standard solutions of glucose and xylose.

### Stationary-phase substrate-demand experiment

For estimation of near-maintenance substrate demand, HET cultures were grown in half-strength ACM with approximately 20 g L⁻¹ nominal glucose and followed to early stationary phase. Dry biomass and residual glucose were measured on days 0, 3, 6, 7 and 8. Apparent maintenance-associated glucose demand between days 7 and 8 was calculated as (S7 − S8)/X /24, where S is glucose concentration (g L⁻¹) and X is the mean biomass concentration across the interval. The result was also converted to mmol glucose gDW⁻¹ h⁻¹ using a glucose molar mass of 180.156 g mol⁻¹. Because biomass decreased during the interval, this value is interpreted as an apparent substrate demand and not as a direct ATP-maintenance coefficient.

### Statistical Analysis

Replicate numbers vary by experiment and time point and are reported in the figure captions and Supplementary Data 7. Grouped data are expressed as mean ± standard deviation unless otherwise stated; Supplementary Data 7 provides the individual measurements, included n values and sample-SD calculations. Statistically significant differences between treatments were determined by one-way analysis of variance followed by Bonferroni post-hoc test. IBM SPSS Statistics for Windows, Version 29.0.2.0 (IBM Corp., Armonk, NY, USA) was used for statistical data analysis and figure creation, with p < 0.05 considered statistically significant.

### Genome-scale metabolic reconstruction and compartmentation curation

The *G. sulphuraria* genome-scale metabolic model was developed following established reconstruction principles [82] and curated at reaction, gene–protein–reaction (GPR), directionality and compartment levels. The draft network was based primarily on the *G. sulphuraria* MetaCyc/BioCyc reconstruction and genome annotation associated with NCBI BioProject PRJNA221242. Structural curation was performed in **ScrumPy** [83], while subsequent constraint-based simulations and optimization used the **COBRA Toolbox** in MATLAB [84].

Cytosolic reactions were derived mainly from the *G. sulphuraria* genome/MetaCyc annotation. Mitochondrial, plastid, thylakoid and peroxisomal reactions were assigned using G. *sulphuraria*-specific literature and, where applicable, mappings to the compartmentalized Chlamydomonas reinhardtii model iCre1355 [19]. Compartments were connected only through defined transport reactions.

Reaction-level provenance and curation decisions were retained in Supplementary Data 5, including reaction equations, compartment, GPR, bounds, subsystem, EC/KEGG identifiers, mapped iCre1355 reactions, implementation source, mapping confidence, unresolved metabolites and curation notes. The SBML file served as the executable model, whereas the workbook documented compartmentation, provenance and manual curation. These files are provided in Supplementary Data 5.

### Literature specific-growth-rate dataset

A condition-specific literature dataset of *G. sulphuraria* specific growth rates (μ) was compiled for AUTO, MIXO and HETERO cultivation. Only species-specific growth-rate observations were retained; biomass concentration, productivity, unresolved records, model-derived constraints and congeneric taxa were excluded. Reported μ values were retained in d⁻¹. When only doubling time (t_d, h) was available, μ was calculated as μ = ln(2) × 24 / t_d, with uncertainty propagated as SD_μ = μ × SD_td / t_d. Dilution rate was used as operational μ only for documented steady-state cultures. Records were classified as primary, steady-state, doubling-time-derived, secondary, or grouped/replicate-specific; unresolved information was recorded as NA. For model comparison, μ values were converted to h⁻¹ by division by 24. No pooled mean or universal μ_max was imposed; condition-specific values and trophic-state ranges defined the validation envelope. Each literature μ was tested for feasible growth under the corresponding trophic condition. The complete dataset, DOI mapping, uncertainty information and validation notes are provided in Supplementary Data 4.

### Biomass objective construction

Biomass objective functions were specified separately for AUTO, MIXO and HETERO following BOF-construction principles [85]. The AUTO BOF retained the literature-derived composition. For MIXO and HETERO, experimentally measured dry-mass fractions were retained without normalization: MIXO protein 0.4147, carbohydrate 0.3377, lipid 0.0701 and nucleic acids 0.0775 g gDW⁻¹; HETERO protein 0.4016, carbohydrate 0.3609, lipid 0.0610 and nucleic acids 0.0765 g gDW⁻¹. Each measured category was allocated among its constituent molecular species using relative proportions in TOTAL_BIOMASS.xlsx (Supplementary Data 2), preserving the measured category totals. Approximately 0.1 g gDW⁻¹ of dry biomass was not characterized by these measurements and was neither assigned to a hypothetical metabolite nor redistributed among measured categories. Pigments were represented independently according to model assumptions. This residual is a compositional coverage gap, not a measured ±10% statistical error bar. Accordingly, direct equivalence of one biomass-objective flux unit with 1 g measured total dry biomass remains a model-calibration assumption.

The lipid component was reconstructed at fatty-acid and lipid-class resolution [23, 86]. Total lipid was partitioned between membrane lipid and TAG, the membrane fraction was distributed among nine lipid classes, and trophic-state fatty-acid profiles were converted to molecular-species coefficients. Exact allocation parameters lacking direct experimental measurements were retained as explicit modelling assumptions and documented separately from measured inputs.

Genome sequence GCF_000341285.1 was used to determine DNA nucleotide composition and annotated coding sequences were used as the RNA composition proxy. GAM was incorporated into biomass formation, whereas NGAM = 1.7 mmol ATP gDW⁻¹ h⁻¹, derived from the stationary-phase experiment in this study, was imposed through a separate ATP-maintenance reaction.

Component evidence and calculations are provided in Supplementary Data 1. Reference BOFs and molecular proportions are in TOTAL_BIOMASS.xlsx (Supplementary Data 2; identical shared Input in Data 3). Its lipid-pool reaction excludes diacylglycerol as a separate input. The scripts in Data 3 apply measured MIXO and HETERO fractions at runtime.

### Model constraints, FBA and FVA

Simulations were performed separately for AUTO, MIXO and HETERO states using the corresponding trophic-state biomass objective. AUTO was defined as CO₂-supported growth without organic-carbon uptake; CO₂ availability was set to 6 mmol gDW⁻¹ h⁻¹, photon uptake was limited to 20 mmol gDW⁻¹ h⁻¹, and Rubisco flux was constrained to 0.01–1000 mmol gDW⁻¹ h⁻¹. MIXO combined one organic substrate with CO₂ and finite light; organic-carbon availability was normalized to 6 C-mmol gDW⁻¹ h⁻¹, photon uptake was limited to 50 mmol gDW⁻¹ h⁻¹, CO₂ uptake was permitted, and Rubisco was constrained to a minimum flux of 0.01 mmol gDW⁻¹ h⁻¹. HETERO permitted one organic substrate at a time, while external CO₂ uptake and Rubisco flux were closed. A small photon allowance of 0.2 mmol gDW⁻¹ h⁻¹ was retained under HETERO conditions to permit pigment-associated biomass reactions rather than to represent photosynthetic growth.

For substrate and product comparisons, alternative organic-carbon uptake routes were closed and only the tested carbon source was enabled. Growth simulations maximized the corresponding biomass reaction. HVP production was evaluated by maximizing each native product-output reaction while maintaining ≥50% of the substrate- and condition-specific maximum growth rate, and fixed-growth carbon yields were additionally calculated at μ = 0.060 h⁻¹. Targeted FVA used fixed biomass flux at μ = 0.060 h ¹ and a 105% minimum-uptake bound. Fluxes are reported in mmol gDW⁻¹ h⁻¹ unless stated otherwise. Reproducible scripts, inputs and numerical outputs are provided in Supplementary Data 3.

### Model quality control and reproducibility

The final GEM model version in SBML file format was evaluated using MEMOTE, a standardized framework for genome-scale metabolic model quality assessment [87]. Quality control included tests of stoichiometric consistency, elemental and charge balance, metabolite formulae, reaction directionality, gene associations, unbounded fluxes and biomass-precursor producibility. Carrier-associated formula corrections were evaluated separately from stoichiometric corrections.

Numerical results in Tables 1–3 and the HVP heatmap are archived outputs associated with the final 2,487-reaction SBML model. The MATLAB scripts assemble the experimental MIXO and HETERO BOFs in memory from measured mass fractions without normalizing their totals. Additional calibration is needed to equate absolute biomass flux with measured total dry biomass.

## Supporting information

Supplementary Data summary

## Data availability

The final SBML model, reconstruction spreadsheet, biomass-objective source tables, cultivation data and simulation scripts generated for this study are provided in Supplementary Data 1–7. The experimental measurements underlying Figures 1–4 are provided in Supplementary Data 7. Additionally model optimisation and analysis results files available in Zenodo (<u>10.5281/zenodo.23348921</u>). All Supplementary Data is also available in GitHub (https://github.com/BigDataInSilicoBiologyGroup/GAldieria-sulphuraria-GEM).

## Acknowledgements

This research was funded by the European Union (European Regional Development Fund) under project No. 1.1.1.3/1./24/A/133. This research was funded by the European Union (European Regional Development Fund) under project No 1.1.1.9/LZP/1/24/028. This research was funded by No. 23-11.17e/20/173/FACCE SURPLUS and supported by No. 23-11.17e/20/173 “Sustainable up-cycling of agricultural residues: modular cascading waste conversion system” (UpWaste) grant number 031B0934A and 031B0934B/).

## Author contributions

A.P. coordinated the overall study and manuscript preparation, performed the genome-scale metabolic modelling, model curation and computational analyses, and prepared the manuscript. S.K. coordinated and performed the experimental part. M.G. and K.S. performed biomass cultivation and biomass-related experimental analyses. R.S. performed HPLC analyses. A.V. contributed to experimental methodology and study design. D.P. contributed expertise in algal cultivation methodology and S.S. provided expertise on the experimental and modeling optimisation design set development. E.S. provided expertise on modeling optimisation design set development. All authors contributed to discussion of the results, reviewed the manuscript and approved the final version.

## Supplementary information

Supplementary Data 1 – BOF Build. Twenty-two component calculation workbooks and six explanatory documents covering protein, carbohydrate, lipid, RNA/DNA, pigment, GAM and NGAM biomass construction.

Supplementary Data 2 – TOTAL_BIOMASS. Literature-informed reference reactions and relative molecular compositions in TOTAL_BIOMASS.xlsx, with the BOF integration manual, macro-biomass description and experimental BOF methods note.

Supplementary Data 3 – Model Optimisation. MATLAB analyses 1–4, shared inputs, archived numerical outputs and HVP figures generated with the final SBML model.

Supplementary Data 4 – Growth Rates. Curated trophic-state-separated specific-growth-rate workbook and explanatory methods document.

Supplementary Data 5 – Model Reconstruction. The final SBML, GSM_COMP_KEGG_LINK.xlsx and its reading guide.

Supplementary Data 6 – MEMOTE. Quality report for the final SBML and the algal-model comparison workbook.

Supplementary Data 7 – Experimental Data. Raw and grouped experimental measurements underlying Figures 1–4, including endpoint growth, biomass and sugar kinetics, OD750 and cell-count measurements, biomass composition, and stationary-phase glucose-demand and NGAM calculations.

