## Supplementary Data summary for "Experimental trophic flexibility guides a compartmentalized genome-scale model of red microalgae *Galdieria sulphuraria*"

#### Contents

Supplementary Table S1 compares MEMOTE results. Supplementary Figures S1-S3 show the MIXO capacity heatmap and fixed-growth HETERO/MIXO carbon-yield plots. Supplementary Tables S2-S4 define HVP targets and summarize fixed-growth analyses. The closing page maps Supplementary Data 1-7 to the distributed files.

#### Model identity and quality-report scope

The final SBML in Supplementary Data 5 and the shared Inputs directory of Supplementary Data 3 contains 2,487 reactions, 2,270 metabolites and 1,050 gene products. The archived numerical outputs in Supplementary Data 3 and MEMOTE report in Supplementary Data 6 refer to this model.

The MEMOTE report evaluates the final SBML: 90.182% overall and 99.500% consistency, with 49 mass-unbalanced and 28 charge-unbalanced reactions.

#### Supplementary Table S1

MEMOTE comparison of the final G. sulphuraria GEM with published eukaryotic algal models for which numerical overall MEMOTE scores were reported. Different reported MEMOTE runs are shown for context; full provenance and caveats are in the comparison workbook in Supplementary Data 6.

| Organism / model | Genes | Reactions | Metabolites | MEMOTE total (%) | Consistency (%) |
|---|---|---|---|---|---|
| G. sulphuraria / final GEM | 1,050 | 2,487 | 2,270 | 90.182 | 99.500 |
| Nannochloropsis salina / iNS934 | 934 | 2,345 | 1,985 | 18.0 | not reported |
| Microchloropsis gaditana / iRJ1321 | 1,321 | 1,918 | 1,862 | 25.3 | not reported |
| Microchloropsis gaditana / AuCoMe | 3,474 | 2,729 | 3,076 | 17.7 | not reported |
| Microchloropsis gaditana B-31 / iMgadit23 | 889 | 2,330 | 1,977 | 67.0 | 55.7 |

The 90.182% entry applies to the supplied 2,487-reaction final SBML. Symbolic spent-photon carriers Photon_USED_Plas and Photon_USED_Thyl (formula R1; charge 0) support formal photon bookkeeping and are not chemical metabolites.

#### Supplementary Figure S1

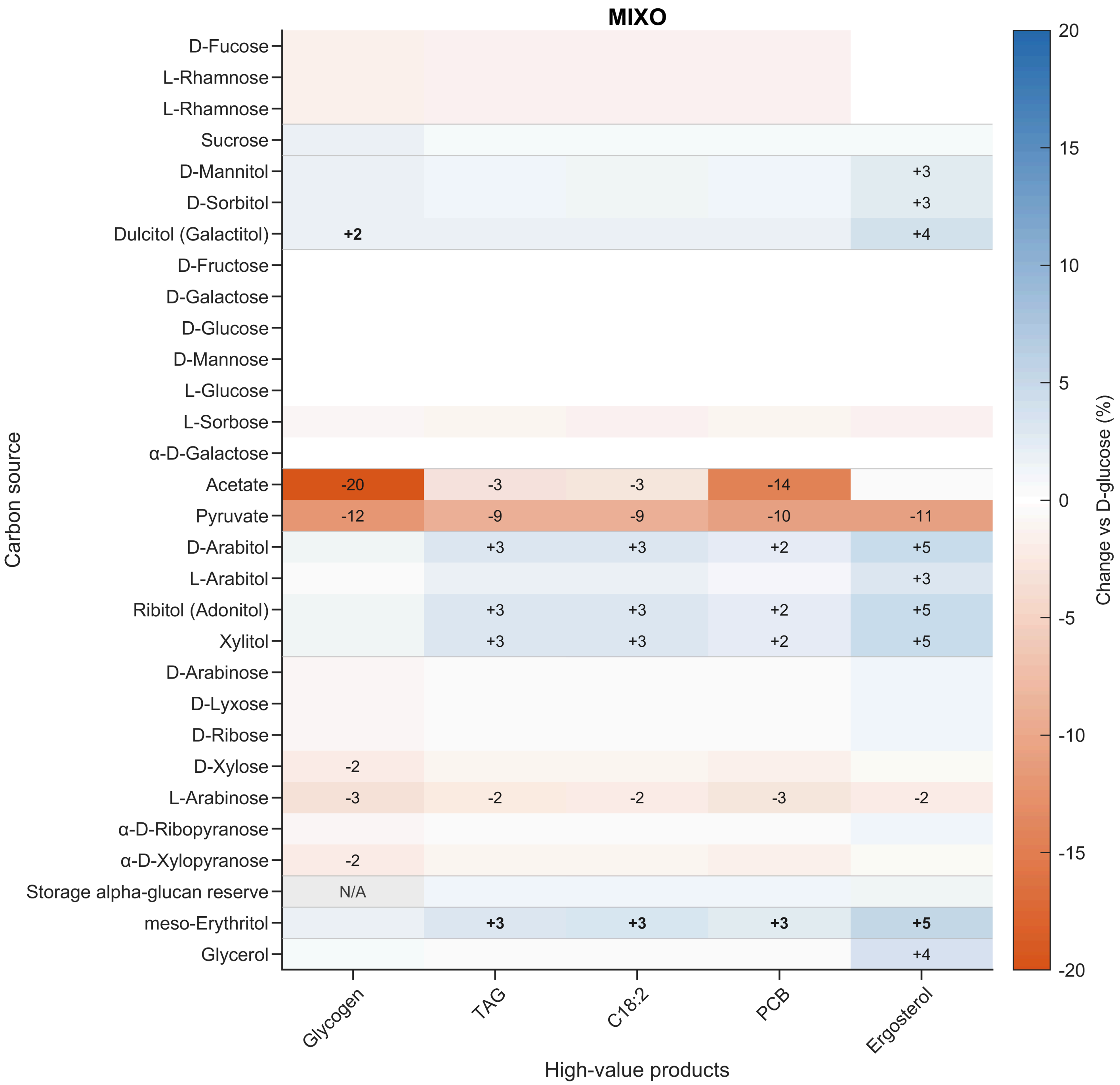


MIXO capacity change vs D-glucose at >=50% substrate-specific maximum growth.

### Supplementary Figure S2

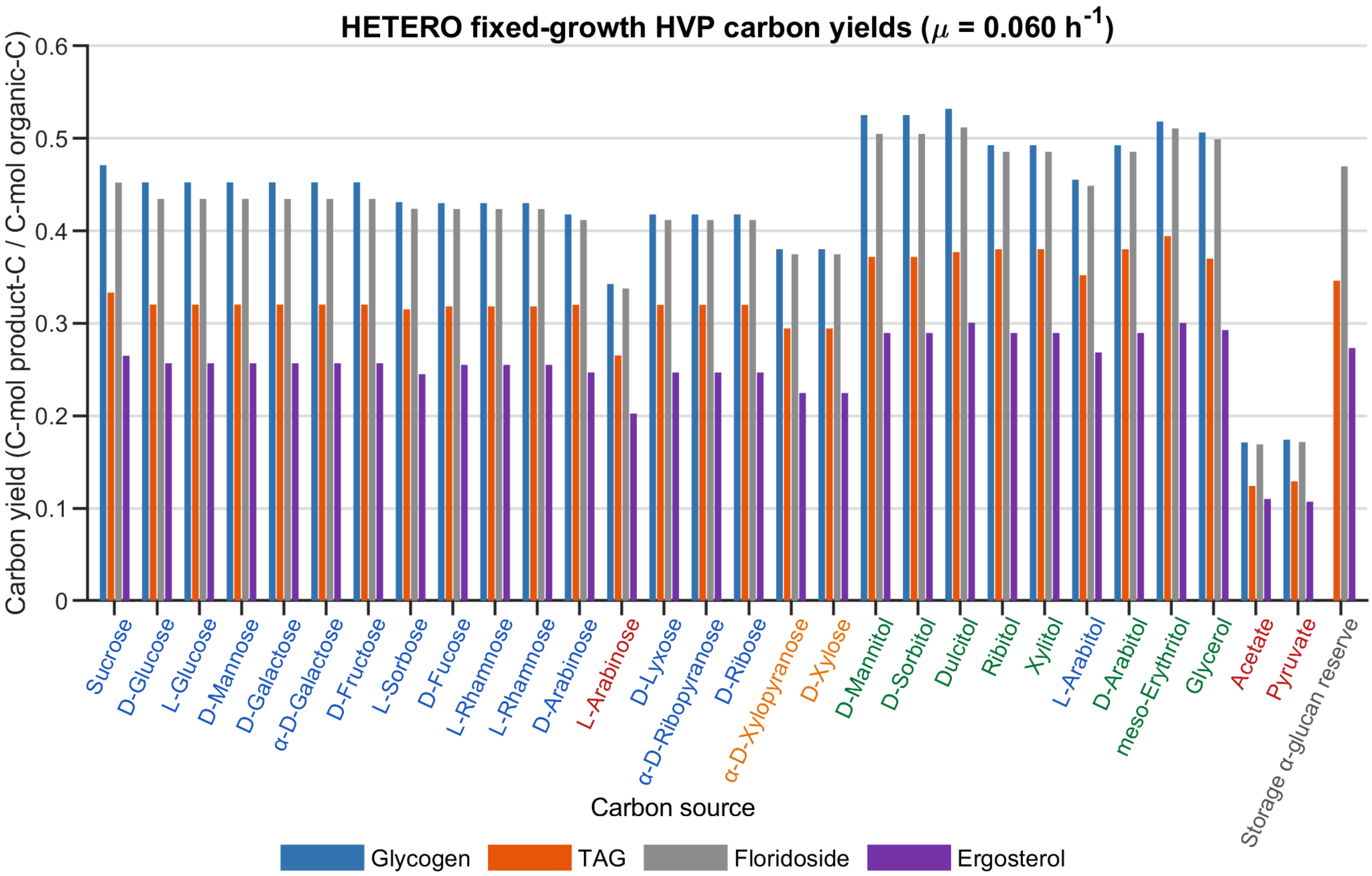


Fixed-growth (mu = 0.060 h-1) HETERO HVP carbon yields; organic-carbon basis.

#### Supplementary Figure S3

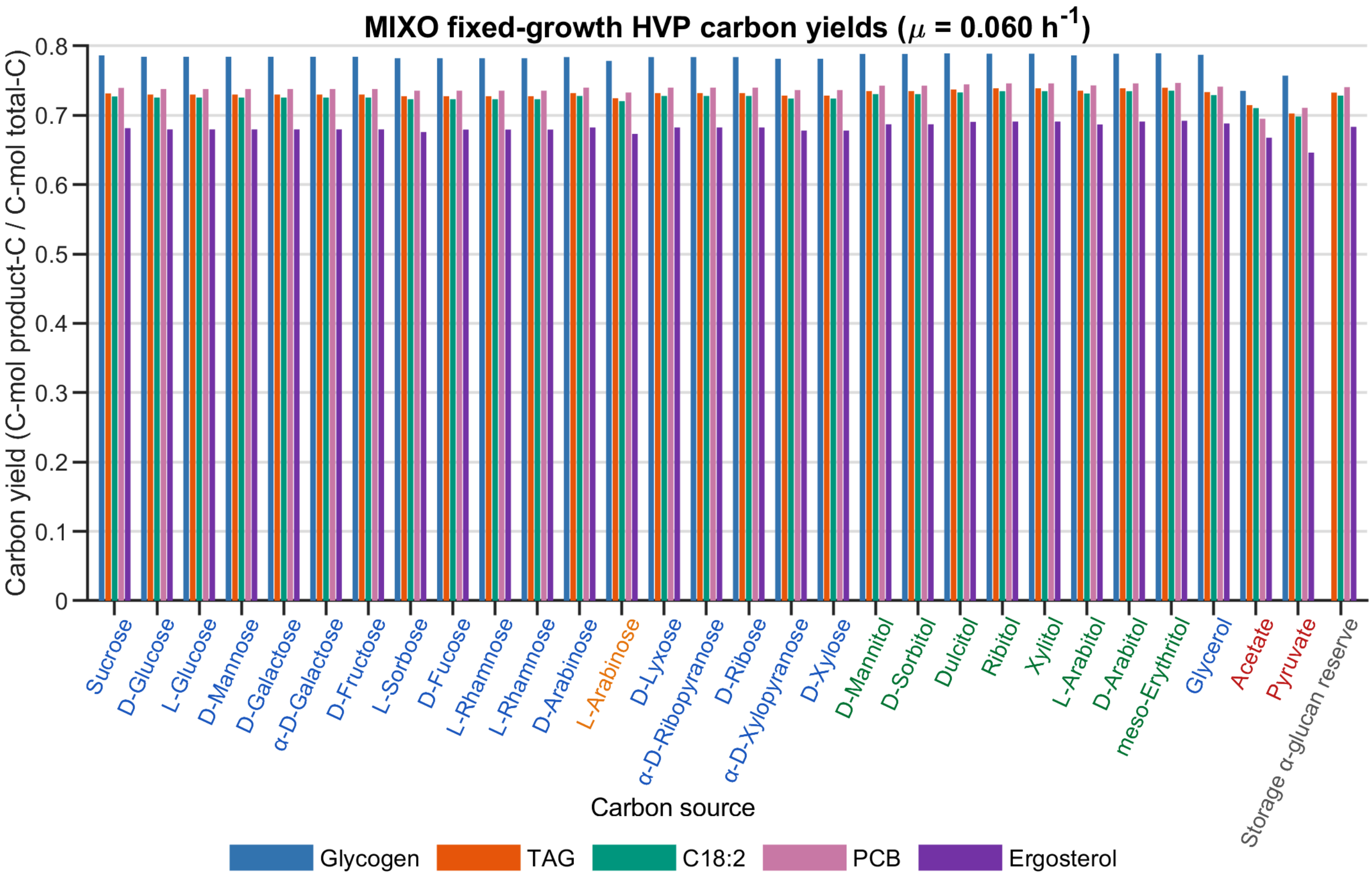


Fixed-growth (mu = 0.060 h-1) MIXO HVP total-carbon yields (organic C + CO2-C).

### Supplementary Table S2

Supplementary Table S2. High-value products prioritized for biological interpretation in each trophic state.

| HVP | Primary significance | AUTO | MIXO | HETERO | Key Galdieria DOI(s) |
|---|---|---|---|---|---|
| Glycogen | Storage/specialty carbohydrate | ✓ | ✓ | ✓ | 10.1016/j.ijbiomac.2016.04.051; 10.1016/j.biortech.2015.11.014 |
| TAG | Neutral lipid / lipid feedstock | — | ✓ | ✓ | 10.1016/j.biortech.2015.11.014 |
| C18:2 | Nutritional ω-6 PUFA | ✓ | ✓ | — | 10.1039/C2FO30198A |
| β-Carotene | Pigment; provitamin A | ✓ | — | — | 10.3389/fpls.2022.978246 |
| PCB | Phycocyanin chromophore | ✓ | ✓ | — | 10.1111/febs.70391; 10.3389/fpls.2022.978246 |
| Floridoside | Compatible solute | — | — | ✓ | 10.1111/nph.13108; 10.1186/s13568-016-0244-6 |
| Squalene | Sterol-pathway precursor | — | — | — | No direct quantitative Galdieria DOI validated |
| Ergosterol | Sterol; vitamin $D_2$ precursor | — | ✓ | ✓ | 10.3390/md18030169; 10.3390/ijms22031247 |
| Phytol | Chlorophyll-derived diterpene | ✓ | — | — | 10.3390/md18030169; 10.3390/ijms22031247 |

### Supplementary Table S3

Highest fixed-growth HETERO carbon yields at mu = 0.060 h-1.

Yields: C-mol product-C / C-mol organic carbon.

| HVP | Best substrate | Max yield | Glucose yield | Change (%) |
|---|---|---|---|---|
| Glycogen | Dulcitol | 0.532 | 0.453 | 17.575 |
| TAG | meso-Erythritol | 0.394 | 0.320 | 23.109 |
| Floridoside | Dulcitol | 0.512 | 0.435 | 17.796 |
| Ergosterol | Dulcitol | 0.301 | 0.257 | 17.032 |

Changes are based on underlying unrounded optimization results.

### Supplementary Table S4

MIXO fixed-growth carbon uptake at mu = 0.060 h-1.

Medians across the five selected HVP optimizations; C-mmol gDW-1 h-1.

| Substrate | Organic C | $CO_2$ C | Total C | $CO_2$ share (%) |
|---|---|---|---|---|
| D-Glucose | 6.000 | 2.241 | 8.241 | 27.194 |
| Dulcitol (Galactitol) | 6.000 | 2.467 | 8.467 | 29.135 |
| Ribitol (Adonitol) | 6.000 | 2.521 | 8.521 | 29.587 |
| meso-Erythritol | 6.000 | 2.548 | 8.548 | 29.808 |
| Glycerol | 6.000 | 2.346 | 8.346 | 28.113 |
| Acetate | 6.000 | 1.683 | 7.683 | 21.905 |
| Pyruvate | 6.000 | 1.482 | 7.482 | 19.811 |

Organic C and CO2 C are separate uptake streams; total C is their sum.

#### Supplementary Data package manifest

The numbering below is used throughout the manuscript and this Supplementary Information. The combined archive contains Supplementary Data 1-7.

| Package | Contents | Main-text use |
| --- | --- | --- |
| Supplementary Data 1 - BOF Build | Component workbooks and explanatory documents | BOF evidence, calculations, GAM and NGAM |
| Supplementary Data 2 - TOTAL_BIOMASS | Reference TOTAL_BIOMASS.xlsx and integration documents | Reference AUTO/MIXO/HETERO reactions and molecular distributions |
| Supplementary Data 3 - Model Optimisation | MATLAB Scripts 1-4, shared inputs, archived outputs and figures | Growth validation, substrate optimization, FVA and HVP analyses |
| Supplementary Data 4 - Growth Rates | Curated growth-rate workbook and explanatory document | Literature growth-validation envelope |
| Supplementary Data 5 - Model Reconstruction | Final SBML, mapping workbook and guide | Executable final model, compartmentation and provenance |
| Supplementary Data 6 - MEMOTE | HTML report for final SBML and comparison workbook | Final-model quality and Supplementary Table S1 |
| Supplementary Data 7 - Experimental Data | Raw and grouped measurements underlying Figures 1-4 | Endpoint growth, kinetics, cell counts, composition and NGAM |
