## Supplementary Data 1 for "Experimental trophic flexibility guides a compartmentalized genome-scale model of red microalgae *Galdieria sulphuraria*": Galdieria_Carbohydrate_Biomass.docx

Carbohydrate Biomass Pseudo‑Reaction Construction for Galdieria sulphuraria

Introduction

In genome‑scale metabolic models (GEMs), the biomass objective function (BOF) represents the overall biosynthetic demand required for cellular growth. The BOF is implemented as a pseudo‑reaction consuming precursor metabolites in proportions corresponding to experimentally determined biomass composition. Accurate representation of macromolecular composition is essential because the BOF defines metabolic flux distribution during flux balance analysis (FBA).

One of the major macromolecular components of algal biomass is carbohydrate. In extremophilic red microalgae such as Galdieria sulphuraria (order Cyanidiales), carbohydrates play two key physiological roles: (i) intracellular storage of carbon and energy, mainly as glycogen, and (ii) structural polysaccharides forming the cell wall matrix. Because carbohydrate metabolism is highly dependent on trophic growth mode, biomass composition varies significantly between autotrophic, mixotrophic, and heterotrophic conditions.

The objective of the present work was to construct trophic‑condition‑specific carbohydrate biomass pseudo‑reactions for Galdieria sulphuraria suitable for implementation in a genome‑scale metabolic model. The methodology integrates trophic-state-specific total carbohydrate fractions with a literature-constrained structural carbohydrate component representing cell-wall polysaccharides. Direct heterotrophic biomass-composition datasets from Pleissner et al. (2021, 2023, 2025), together with protein-constrained mass closure for autotrophy and physiological interpolation for mixotrophy, were used to define the total carbohydrate coefficients. The total pool was then partitioned into intracellular storage glycogen and structural wall carbohydrate.

### 1. Experimental Data Sources

Total carbohydrate coefficients were taken from the accompanying carbohydrate-fraction synthesis rather than from glycogen concentration alone. Direct heterotrophic measurements showed strong condition dependence: 0.20 g gDW⁻¹ in digestate/hydrolyzed-straw cultivation (Pleissner et al., 2021, DOI: 10.1016/j.biortech.2021.125477), 0.442 g gDW⁻¹ during fed-batch cultivation on hydrolyzed fish residues (Pleissner et al., 2023), and 0.038–0.436 g gDW⁻¹ across technical-scale medium systems (Pleissner et al., 2025). These observations support a practical heterotrophic BOF range of 0.10–0.36 g gDW⁻¹ for balanced growth conditions.

A direct autotrophic carbohydrate mass fraction was not available in the selected datasets. The autotrophic range was therefore estimated by mass closure using the updated protein fraction and a conservative residual allowance for lipids, pigments, nucleic acids, ash, cofactors, and other biomass components. This gave 0.16–0.23 g gDW⁻¹. Mixotrophy was treated as an intermediate physiological state between autotrophy and heterotrophy, giving a practical range of 0.20–0.25 g gDW⁻¹.

Sakurai et al. (2016, DOI: 10.1016/j.biortech.2015.11.014) remains relevant for identifying glycogen as the major intracellular storage carbohydrate of Galdieria sulphuraria, but its glycogen measurements are not used here as the sole basis for the total carbohydrate fraction.

### 2. Calculation and Partitioning of Total Carbohydrate Fractions

The trophic-state total carbohydrate coefficient was defined first. For the mean BOF, the midpoint of each recommended range was used:

Autotrophic mean = (0.16 + 0.23) / 2 = 0.195 g gDW⁻¹

Mixotrophic mean = (0.20 + 0.25) / 2 = 0.225 g gDW⁻¹

Heterotrophic mean = (0.10 + 0.36) / 2 = 0.230 g gDW⁻¹

Structural carbohydrate was treated as a subfraction of this total, not as an additional mass added on top. The intracellular storage-glycogen coefficient was therefore calculated as:

storage glycogen = total carbohydrate − structural carbohydrate

Using the central structural-carbohydrate estimate of 0.020 g gDW⁻¹ gives storage-glycogen coefficients of 0.175, 0.205, and 0.210 g gDW⁻¹ for autotrophic, mixotrophic, and heterotrophic biomass, respectively. Glycogen is represented as an anhydroglucose polymer with empirical repeat formula C6H10O5.

Table 1. Recommended total carbohydrate fractions used for trophic-state BOF construction.

| **Trophic state** | **Total carbohydrate range (g gDW⁻¹)** |
| --- | --- |
| Autotrophic | 0.16–0.23 |
| Mixotrophic | 0.20–0.25 |
| Heterotrophic | 0.10–0.36 |

### 3. Structural Carbohydrate Representation

The total-carbohydrate datasets quantify combined carbohydrate biomass but do not explicitly separate intracellular storage glycogen from but does not explicitly quantify structural polysaccharides associated with the cell wall. Direct measurements of the whole-cell structural-carbohydrate fraction of Galdieria sulphuraria are currently unavailable. The closest taxonomic evidence is provided by the related Cyanidiales red alga Cyanidium caldarium. Isolated cell-wall preparations of this species were protein-rich (approximately 50–55 % of wall material) and contained about 12–14 % hemicellulose and 3–4 % cellulose. These measurements demonstrate that Cyanidiales walls contain structural polysaccharides, but they do not report the cell-wall mass as a fraction of whole-cell dry weight and therefore cannot be used directly as a g gDW⁻¹ coefficient.

Mannan-containing wall polymers are also documented in red algae. In Bangia atropurpurea, the wall contains β-1,4-mannan together with β-1,3-xylan and cellulose. This evidence supports inclusion of a mannose-containing structural component, although it is not a quantitative measurement for G. sulphuraria.

As a broader whole-biomass reference, comparative analysis of ten microalgae reported that cell-wall-related polysaccharides contributed approximately 10 % of dry biomass on average, with substantial species-specific variation. Because the Cyanidiales wall is relatively protein-rich and has a lower polysaccharide proportion than many other microalgae, 0.10 g gDW⁻¹ was treated as an upper comparative value rather than the central coefficient.

A literature-constrained structural-carbohydrate coefficient of 0.020 g gDW⁻¹ was therefore selected for the central GSM biomass composition, with 0.010–0.030 g gDW⁻¹ recommended for sensitivity analysis. The central coefficient was provisionally partitioned into 0.016 g gDW⁻¹ structural glucan and 0.004 g gDW⁻¹ mannose-containing polymer. This 80:20 partition is a modelling allocation used to represent both wall-carbohydrate classes; it is not a directly measured glucan:mannan ratio for G. sulphuraria.

Table 2. Structural carbohydrate fractions used in the GSM model.

| **Structural carbohydrate** | **Fraction (g gDW⁻¹)** |
| --- | --- |
| Glucan-containing structural polysaccharides | 0.016 |
| Mannose-containing structural polysaccharides | 0.004 |

### 4. Integration of Storage and Structural Carbohydrates

The final carbohydrate biomass fraction for each trophic condition was obtained by partitioning the independently defined total carbohydrate coefficient into storage glycogen and structural wall carbohydrate. The central structural coefficient was fixed at 0.020 g gDW⁻¹ for all three trophic states because no condition-specific wall-composition measurements are available. It was provisionally divided into 0.016 g gDW⁻¹ glucan-containing structural polysaccharides and 0.004 g gDW⁻¹ mannose-containing structural polysaccharides.

The structural fraction was subtracted from each trophic-state total rather than added to it. This prevents double counting and preserves the final total carbohydrate means of 0.195 g gDW⁻¹ for autotrophy, 0.225 g gDW⁻¹ for mixotrophy, and 0.230 g gDW⁻¹ for heterotrophy. The corresponding intracellular storage-glycogen coefficients are 0.175, 0.205, and 0.210 g gDW⁻¹, respectively.

For uncertainty analysis, the total structural-carbohydrate pool may be varied from 0.010 to 0.030 g gDW⁻¹ while retaining the trophic-state total carbohydrate coefficient. Any increase or decrease in structural carbohydrate must therefore be balanced by the opposite change in storage glycogen.

Table 3. Final carbohydrate biomass fractions used in the metabolic model.

| **Condition** | **Storage glycogen** | **Structural glucan** | **Structural mannan** | **Total carbohydrate** |
| --- | --- | --- | --- | --- |
| Autotrophic | 0.175 | 0.016 | 0.004 | 0.195 |
| Mixotrophic | 0.205 | 0.016 | 0.004 | 0.225 |
| Heterotrophic | 0.210 | 0.016 | 0.004 | 0.230 |

### 5. Construction of the Carbohydrate Biomass Pseudo‑Reaction

In genome-scale metabolic modeling, biomass synthesis is represented by a pseudo-reaction that consumes metabolic precursors required for cell growth. The carbohydrate pseudo-reaction represents the synthesis of a generic carbohydrate biomass pool composed of intracellular storage glycogen and structural polysaccharides.

Autotrophic condition:
0.175 glycogen_storage[c] + 0.016 wall_glucan[c] + 0.004 wall_mannan[c] → carbohydrate_biomass[c]

Mixotrophic condition:
0.205 glycogen_storage[c] + 0.016 wall_glucan[c] + 0.004 wall_mannan[c] → carbohydrate_biomass[c]

Heterotrophic condition:
0.210 glycogen_storage[c] + 0.016 wall_glucan[c] + 0.004 wall_mannan[c] → carbohydrate_biomass[c]

These pseudo-reactions preserve the mean total carbohydrate coefficients of 0.195, 0.225, and 0.230 g gDW⁻¹. The structural coefficients are literature-constrained modelling estimates and should be evaluated using the recommended 0.010–0.030 g gDW⁻¹ sensitivity range. The carbohydrate biomass pool is subsequently incorporated into the overall biomass objective function together with lipid, protein, nucleotide, cofactor, pigment, and maintenance-energy terms.

### 6. Conclusion

The carbohydrate biomass pseudo-reaction for Galdieria sulphuraria was constructed by defining the trophic-state total carbohydrate fraction first and then partitioning that total into intracellular storage glycogen and structural wall polysaccharides. This approach prevents the structural fraction from being added on top of an already complete carbohydrate coefficient.

Direct heterotrophic biomass-composition measurements support a broad condition-dependent carbohydrate interval. For balanced GSM simulations, 0.10–0.36 g gDW⁻¹ was retained as the heterotrophic working range. Autotrophic and mixotrophic ranges of 0.16–0.23 and 0.20–0.25 g gDW⁻¹ were retained from the total-carbohydrate synthesis. Their mean BOF coefficients are therefore 0.195, 0.225, and 0.230 g gDW⁻¹ for autotrophic, mixotrophic, and heterotrophic growth, respectively.

Structural carbohydrate was assigned a central coefficient of 0.020 g gDW⁻¹, with a recommended sensitivity range of 0.010–0.030 g gDW⁻¹. This estimate was constrained by the protein-rich wall composition of Cyanidium caldarium (Bailey & Staehelin, 1968, DOI: 10.1099/00221287-54-2-269), evidence for β-mannan in Bangia atropurpurea (Umemoto & Araki, 2010, DOI: 10.1007/s10126-009-9196-z), and comparative measurements of cell-wall-related polysaccharides in microalgae (Bernaerts et al., 2018, DOI: 10.1016/j.algal.2018.03.017).

The mean structural pool was provisionally represented by 0.016 g gDW⁻¹ glucan-containing structural polysaccharides and 0.004 g gDW⁻¹ mannose-containing structural polysaccharides. These are broad model classes rather than definitive polymer assignments for G. sulphuraria. After subtraction of this structural pool, the mean storage-glycogen coefficients are 0.175, 0.205, and 0.210 g gDW⁻¹ for autotrophic, mixotrophic, and heterotrophic biomass, respectively.
