## Supplementary Data 1 for "Experimental trophic flexibility guides a compartmentalized genome-scale model of red microalgae *Galdieria sulphuraria*": Supplementary_GAM_NGAM.docx

Supplementary methods: GAM and NGAM

Galdieria sulphuraria biomass objective function and ATP-maintenance constraint

**Authoritative interpretation. GAM = 30 mmol ATP gDW⁻¹ is the implemented BOF coefficient. NGAM = 1.7 mmol ATP gDW⁻¹ h⁻¹ is a selected working constraint, supported by an approximate Galdieria stationary-phase estimate (≈1.7) and an algal cross-species benchmark (2.85), but it is not a direct Galdieria chemostat measurement.**

### 1. GAM and NGAM represent different energy terms

Growth-associated maintenance (GAM) is ATP hydrolysis coupled to formation of one gram of biomass. Non-growth-associated maintenance (NGAM) is a time-dependent ATP demand that persists independently of biomass production. They therefore have different units and must be implemented separately.

**qATP,maintenance = GAM × μ + NGAM**

| **Parameter** | **Meaning** | **Units** | **Model implementation** |
| --- | --- | --- | --- |
| GAM | Energy coupled to biomass synthesis and assembly | mmol ATP gDW⁻¹ | ATP hydrolysis in, or coupled to, the BOF |
| NGAM | Homeostasis, ion gradients, repair and turnover independent of growth | mmol ATP gDW⁻¹ h⁻¹ | Lower bound on a separate ATP-maintenance reaction |

### 2. Growth-associated maintenance

All three trophic-state GAM workbooks use the same selected coefficient because no condition-specific Galdieria GAM measurements are available. The implemented reaction is:

**30 ATP + 30 H₂O → 30 ADP + 30 Pi + 30 H⁺**

Thus, GAM = 30 mmol ATP gDW⁻¹. This coefficient is a central modeling assumption, not a direct experimental value. It is recorded as GAM_BIOMASS in the autotrophic, mixotrophic and heterotrophic GAM_Objective_Function_Final.xlsx workbooks.

### 3. Non-growth-associated maintenance

#### 3.1. Selected working value and implementation

For the current Galdieria reconstruction, the working value is:

**NGAM = 1.7 mmol ATP gDW⁻¹ h⁻¹**

This value is the chosen point within the study's model-dependent stationary-phase range. NGAM should be implemented through a separate cytosolic ATP-maintenance reaction, with both bounds fixed at 1.7 mmol gDW⁻¹ h⁻¹:

**ATP + H₂O → ADP + Pi + H⁺**

The GAM workbooks contain GAM_BIOMASS but do not currently encode a dedicated NGAM reaction or its lower bound. Consequently, the NGAM value must be declared in the full metabolic model or simulation configuration; it must not be added to the BOF coefficient.

#### 3.2. Previous Galdieria estimate

A later calculation used the near-stationary day 7–8 interval: biomass decreased from 5.12 to 4.96 g L⁻¹ while glucose decreased from 8.68 to 7.43 g L⁻¹. Using the mean biomass concentration (5.04 g L⁻¹), the apparent glucose demand was:

**(1.25 g L⁻¹ d⁻¹ ÷ 5.04 gDW L⁻¹) ÷ 180.16 × 1000 ÷ 24 = 0.057 mmol glucose gDW⁻¹ h⁻¹**

Assuming 30 mmol ATP generated per mmol glucose gives an apparent ATP-maintenance demand of approximately 1.7 mmol ATP gDW⁻¹ h⁻¹. The selected 1.7 value lies within the estimated 1.50–1.73 range, but this remains an NGAM-like estimate rather than a pure NGAM measurement because residual growth, storage turnover, stress metabolism and the true ATP yield were not independently resolved.

#### 3.3. DOI-supported algal benchmark

Kliphuis et al. determined ATP requirements from seven photoautotrophic chemostat experiments with Chlamydomonas reinhardtii. They reported 109 mmol ATP g⁻¹ for biomass formation and 2.85 mmol ATP g⁻¹ h⁻¹ for maintenance (DOI: 10.1007/s10811-011-9674-3). This validates the experimental method and provides an algal benchmark, but it does not directly determine the Galdieria parameter.

**Evidence boundary. The statement that algae generally have NGAM values of 0.5–3.0 mmol ATP gDW⁻¹ h⁻¹ is not retained, because the validated sources do not establish that interval as a universal algal range.**

### 4. Recommended reporting

| **Quantity** | **Value used** | **Evidence status** | **Required wording** |
| --- | --- | --- | --- |
| GAM | 30 mmol ATP gDW⁻¹ | Selected and implemented in all three BOFs | Model-selected central coefficient |
| NGAM | 1.7 mmol ATP gDW⁻¹ h⁻¹ | Selected working constraint | Not a direct Galdieria measurement |
| Galdieria apparent maintenance | ≈1.7 mmol ATP gDW⁻¹ h⁻¹ | Calculated from day 7–8 batch data | Approximate NGAM-like support only |
| C. reinhardtii maintenance | 2.85 mmol ATP g⁻¹ h⁻¹ | Experimentally determined by chemostat regression | Cross-species benchmark only |

Sensitivity analysis should report at least NGAM = 0, 1.7 and 2.85 mmol ATP gDW⁻¹ h⁻¹, while keeping the provenance of each scenario explicit. A future Galdieria-specific estimate should preferably be obtained from steady-state cultures at several growth rates by regressing ATP production or substrate demand against μ; GAM is represented by the slope and NGAM by the zero-growth intercept.

### 5. Validated references

Kliphuis AMJ, Klok AJ, Martens DE, Lamers PP, Janssen M, Wijffels RH (2012). Metabolic modeling of Chlamydomonas reinhardtii: energy requirements for photoautotrophic growth and maintenance. Journal of Applied Phycology 24:253–266. DOI: 10.1007/s10811-011-9674-3.

Lachance JC, Lloyd CJ, Monk JM, et al. (2019). BOFdat: generating biomass objective functions for genome-scale metabolic models from experimental data. PLoS Computational Biology 15:e1006971. DOI: 10.1371/journal.pcbi.1006971.

Feist AM, Palsson BØ (2010). The biomass objective function. Current Opinion in Microbiology 13:344–349. DOI: 10.1016/j.mib.2010.03.003.
