## Supplementary Data 1 for "Experimental trophic flexibility guides a compartmentalized genome-scale model of red microalgae *Galdieria sulphuraria*": Supplementary_Lipid_BOF.docx

Supplementary methods: trophic-state lipid biomass objective functions

**Galdieria sulphuraria — detailed revalidated calculation rationale**

Companion to the three self-contained trophic-state lipid BOF workbooks

### 1. Purpose and evidence boundary

This supplement explains how lipid biomass-objective-function (BOF) coefficients were constructed when no single experiment supplied a complete, trophic-state-resolved intact lipidome. A BOF represents the metabolites required to form one unit of biomass in defined proportions; its coefficients must therefore combine experimental composition with explicit assumptions when measurements are incomplete (Feist and Palsson, 2010; DOI: 10.1016/j.mib.2010.03.003).

Direct evidence was restricted to trophic-state lipid content, analytical lipid-pool observations and fatty-acid distributions reported for Galdieria by Sakurai et al. (2016; DOI: 10.1016/j.biortech.2015.11.014) and López et al. (2019; DOI: 10.1038/s41598-019-46645-3). The selected MEM/TAG splits, the complete nine-class membrane distributions, exclusion of DG and free fatty acids (FFA) from stable biomass, and the 180 g mol⁻¹ membrane-headgroup contribution are modeling assumptions. They are not claimed to be directly measured values and therefore have no direct experimental DOI.

### 2. Three-level calculation hierarchy

| **Level** | **Question answered** | **Output passed forward** | **Evidence status** |
| --- | --- | --- | --- |
| 1. Stable-pool allocation | How much of total modeled lipid is structural membrane lipid (MEM) versus storage triacylglycerol (TAG)? | fMEM,t and fTAG,t; each trophic-state column sums to 1 | Directional biological evidence plus explicit modeling allocation; Sakurai DOI: 10.1016/j.biortech.2015.11.014 |
| 2. Membrane-class allocation | How is the Level-1 MEM pool divided among nine modeled lipid classes? | fc,t for MGDG, DGDG, SQDG, PG, PE, PC, PI, PS and DGTS; each column sums to 1 | Autotrophic baseline informed by algal membrane biology; mixotrophic/heterotrophic shifts are estimates; no direct complete nine-class dataset |
| 3. BOF conversion | How are total lipid, stable-pool fractions, class fractions and fatty-acid profiles converted to mmol gDW⁻¹ coefficients? | Condition-specific lipid-species coefficients and aggregate BOF reactions | Measured/selected inputs plus transparent molecular-weight approximations; BOF method DOI: 10.1016/j.mib.2010.03.003 |

The hierarchy is directional: Level 1 must be fixed before Level 2, and both must be fixed before Level 3. Level-3 formulas are authoritative for model insertion; Level-1 and Level-2 workbooks document upstream allocation choices and must not be used as independent final BOFs.

### 3. Why Level 1 and Level 3 values differ

Level 1 and Level 3 values are expected to differ because they describe different quantities. Level 1 divides total lipid mass between MEM and TAG and is therefore dimensionless. Level 3 converts the allocated masses into molar, class-specific and molecular-species-specific BOF coefficients expressed in mmol gDW⁻¹. Numerical equality between the two levels is neither required nor meaningful because their units and denominators differ.

The conversion is applied in four steps:

1. Pool mass (g gDW⁻¹) = total lipid (g gDW⁻¹) × Level-1 pool fraction.
2. Pool amount (mmol gDW⁻¹) = pool mass (g gDW⁻¹) ÷ average pool molecular weight (g mol⁻¹) × 1000.
3. Membrane-class amount (mmol gDW⁻¹) = MEM amount × Level-2 class fraction.
4. Species coefficient (mmol gDW⁻¹) = class amount × normalized fatty-acid/species fraction.

Table S2. Relationship between Level-1 allocations and Level-3 pool amounts.

| **Trophic state** | **Total lipid (g gDW⁻¹)** | **Level-1 MEM/TAG** | **Allocated mass MEM/TAG (g gDW⁻¹)** | **Level-3 amount MEM/TAG (mmol gDW⁻¹)** |
| --- | --- | --- | --- | --- |
| Autotrophic | 0.1500 | 0.75 / 0.25 | 0.112500 / 0.037500 | 0.143760 / 0.043518 |
| Mixotrophic | 0.0425 | 0.65 / 0.35 | 0.027625 / 0.014875 | 0.035326 / 0.017177 |
| Heterotrophic | 0.0375 | 0.50 / 0.50 | 0.018750 / 0.018750 | 0.023937 / 0.021443 |

For example, the autotrophic Level-1 MEM fraction of 0.75 acts on 0.1500 g lipid gDW⁻¹, giving 0.1125 g MEM gDW⁻¹. Division by the calculated average membrane-lipid molecular weight and conversion from mol to mmol gives 0.143760 mmol MEM gDW⁻¹. Level 2 and the normalized species profile then distribute this amount among the modeled membrane classes and species.

### 4. Level 1 — allocation of stable lipid between MEM and TAG

#### 4.1. What Level 1 represents

Level 1 does not reproduce every chromatographic pool. It defines the stable lipid demand represented in the BOF. MEM represents structural lipids required to construct cellular and organellar membranes. TAG represents a stable neutral-lipid storage demand. DG and FFA are omitted from this stable two-pool allocation because DG is a central biosynthetic intermediate and FFA are acyl building blocks rather than a uniquely defined stable biomass compartment. This is a modeling decision, not a claim that DG or FFA were absent experimentally.

#### 4.2. Why trophic states differ

Sakurai et al. showed that the amounts and compositions of lipids change across autotrophic, mixotrophic and heterotrophic G. sulphuraria cultivation and that neutral-lipid accumulation is promoted by glucose in darkness (DOI: 10.1016/j.biortech.2015.11.014). López et al. independently showed trophic-state-dependent lipid content and fatty-acid composition in Galdieria sp. USBA-GBX-832 (DOI: 10.1038/s41598-019-46645-3). These studies support the direction of the allocation—greater structural-membrane dominance in light-grown cells and progressively greater storage-lipid representation with organic-carbon dependence—but they do not directly report the exact 0.75/0.25, 0.65/0.35 and 0.50/0.50 BOF fractions.

| **Stable modeled pool** | **Autotrophic** | **Mixotrophic** | **Heterotrophic** | **Rationale / evidence** |
| --- | --- | --- | --- | --- |
| MEM | 0.75 | 0.65 | 0.50 | Assumed monotonic decline in photosynthetic-membrane dominance from light-only to dark organic-carbon growth; direction supported by 10.1016/j.biortech.2015.11.014 and 10.1038/s41598-019-46645-3 |
| TAG | 0.25 | 0.35 | 0.50 | Complement to MEM within the stable modeled pool; increased storage representation is biologically consistent with trophic-state lipid remodeling; exact values are assumptions |
| DG | 0 | 0 | 0 | Excluded from stable BOF. The reported heterotrophic 43% analytical DG observation is retained as evidence, not added as a third stable pool; 10.1016/j.biortech.2015.11.014 |
| FFA | 0 | 0 | 0 | Excluded as a standalone stable pool; fatty-acid percentages remain inputs for reconstructing acyl composition |
| Closure | 1.00 | 1.00 | 1.00 | Required normalization: fMEM,t + fTAG,t = 1 |

#### 4.3. Why heterotrophic DG was not counted

The heterotrophic analytical dataset contains a large DG observation (43%; Sakurai et al., DOI: 10.1016/j.biortech.2015.11.014). Counting that value together with independently assigned MEM and TAG fractions would mix an intermediate pool with end-pool demands and could double-count lipid carbon because DG is a precursor to both glycerophospholipids/glycolipids and TAG. For the stable BOF, MEM and TAG were therefore renormalized to one and DG was set to zero.

### 5. Level 2 — allocation of MEM among nine lipid classes

#### 5.1. Class definitions and modeled compartments

The MEM pool was separated into plastid-associated glycolipids (MGDG, DGDG and SQDG), phospholipids (PG, PE, PC, PI and PS), and the betaine lipid DGTS. This creates a model-compatible set of class pseudo-metabolites while retaining the main functional distinction between photosynthetic and extraplastidial membranes. The complete nine-class pattern was not measured in one matched G. sulphuraria experiment; the table is therefore an internally consistent reconstruction.

| **Class** | **AUTO** | **MIXO** | **HETERO** | **Interpretation of the assumed trophic trend** |
| --- | --- | --- | --- | --- |
| MGDG | 0.16 | 0.14 | 0.12 | Reduced with decreasing photosynthetic-membrane contribution |
| DGDG | 0.23 | 0.21 | 0.19 | Largest glycolipid class; reduced progressively outside strict autotrophy |
| SQDG | 0.13 | 0.11 | 0.08 | Reduced with the modeled plastid-membrane fraction |
| PG | 0.07 | 0.07 | 0.07 | Held constant to avoid unsupported trophic redistribution |
| PE | 0.14 | 0.14 | 0.17 | Increased in heterotrophy as extraplastidial membrane representation rises |
| PC | 0.16 | 0.17 | 0.19 | Modestly increased from autotrophy to heterotrophy |
| PI | 0.04 | 0.04 | 0.04 | Held constant because condition-specific evidence was insufficient |
| PS | 0.02 | 0.02 | 0.03 | Small pool; slight heterotrophic increase assumed |
| DGTS | 0.05 | 0.10 | 0.11 | Higher modeled representation in organic-carbon conditions |
| Sum | 1.00 | 1.00 | 1.00 | Required closure within MEM |

#### 5.2. How the Level-2 values were assumed

The autotrophic column was used as the structural baseline because autotrophic biomass must retain the strongest photosynthetic-membrane demand. Mixotrophic values were placed between autotrophic and heterotrophic endpoints unless evidence was insufficient, in which case the fraction was held constant. Heterotrophic values reduced the three modeled plastid glycolipids and redistributed the released fraction primarily to PE, PC, PS and DGTS. This interpolation is constrained by three rules: all values are non-negative; each trophic-state column sums exactly to one; and the direction of the aggregate glycolipid-to-non-glycolipid shift agrees with the trophic remodeling observed in Galdieria lipid studies (DOIs: 10.1016/j.biortech.2015.11.014; 10.1038/s41598-019-46645-3).

No DOI is assigned to an individual Level-2 number because none of the exact class fractions is a direct literature measurement. The literature supports biological direction and the existence of trophic remodeling, not the precise numerical reconstruction.

### 6. Level 3 — conversion to condition-specific BOF coefficients

#### 6.1. Step 3.1: select total lipid content

| **Trophic state** | **Selected Ltotal,t (g lipid gDW⁻¹)** | **Source status** | **MEM mass** | **TAG mass** |
| --- | --- | --- | --- | --- |
| Autotrophic | 0.1500 | Selected central value informed by López et al.; 10.1038/s41598-019-46645-3 | 0.112500 | 0.037500 |
| Mixotrophic | 0.0425 | Selected central value informed by López et al.; 10.1038/s41598-019-46645-3 | 0.027625 | 0.014875 |
| Heterotrophic | 0.0375 | Selected central value informed by López et al.; 10.1038/s41598-019-46645-3 | 0.018750 | 0.018750 |

For trophic state t: mMEM,t = Ltotal,t × fMEM,t; mTAG,t = Ltotal,t × fTAG,t. The selected values are BOF inputs, not universal species constants. Strain, medium, temperature, light, growth phase and extraction method can change total lipid content (DOIs: 10.1016/j.biortech.2015.11.014; 10.1038/s41598-019-46645-3).

#### 6.2. Step 3.2: normalize fatty-acid profiles

Condition-specific fatty-acid distributions were transferred from the source tables in Sakurai et al. (DOI: 10.1016/j.biortech.2015.11.014) and López et al. (DOI: 10.1038/s41598-019-46645-3). When the included reported species did not sum exactly to 100%, each percentage was renormalized: pnorm,i = 100 × pi / Σpi. Renormalization preserves relative composition while preventing incomplete coverage or rounding from changing the assigned lipid-pool mass.

The notation Cn:d was interpreted chemically: n is the carbon number and d is the number of double bonds. Earlier workbook values that doubled d were corrected. For a free fatty acid represented as CnH(2n−2d)O2, the working molecular weight is MWFA,i = 12.011n + 1.008(2n−2d) + 2(15.999).

#### 6.3. Step 3.3: reconstruct TAG coefficients

TAG molecular species were represented using the condition-specific TAG fatty-acid distribution and an average glycerol/backbone correction. The allocated TAG mass is first converted to an effective molar basis, then multiplied by each normalized fatty-acid fraction. In generic form: cTAG,i,t = 1000 × mTAG,t × pTAG,i,t / MW̄TAG,t, where p is expressed as a fraction. The factor 1000 converts mol gDW⁻¹ to mmol gDW⁻¹. This reconstruction is necessary because FAME analysis measures derivatized fatty acids, not a complete set of intact TAG molecular species.

#### 6.4. Step 3.4: reconstruct membrane-class coefficients

For membrane class c and fatty acid i in trophic state t, the implemented structure is: cc,i,t = 1000 × mMEM,t × fc,t × pMEM,i,t / MW̄MEM,t. Here fc,t is the Level-2 fraction and pMEM,i,t is the normalized membrane fatty-acid fraction. MW̄MEM,t combines the weighted fatty-acid contribution with a single 180 g mol⁻¹ average headgroup/backbone contribution. The 180 g mol⁻¹ term is an explicit approximation and has no direct DOI; it is used because matched intact molecular species and class-specific molecular weights were unavailable for every class.

The same fatty-acid proxy distribution is applied across membrane classes within a trophic state. This assumes that class-specific acyl distributions are sufficiently similar for BOF-scale representation. It should not be interpreted as a lipidomics claim that MGDG, DGDG, SQDG, PG, PE, PC, PI, PS and DGTS contain identical acyl species.

#### 6.5. Step 3.5: assemble aggregate BOF reactions

Each class-specific reaction consumes its reconstructed species coefficients to form one class pseudo-metabolite. A final lipid-pool reaction combines the nine membrane-class pseudo-metabolites and TAG into the modeled total lipid pool. DG is excluded from the stable BOF and therefore has no reaction row in the current Reaction List. This layered pseudo-metabolite design separates biochemical class assembly from the final biomass demand, consistent with the general purpose of a BOF to represent biomass precursor requirements in defined proportions (Feist and Palsson, DOI: 10.1016/j.mib.2010.03.003).

#### 6.6. Interpretation of Level-3 model coefficients

- Level 1 is used only to assign total lipid mass to MEM and TAG; Level 2 is used only to divide MEM among the nine membrane-lipid classes.
- Only Level-3 coefficients are transferred as stoichiometric coefficients for individual lipid metabolites and aggregate lipid reactions.
- Glycerol-3-phosphate is not added as a separate BOF term because the reconstructed intact-lipid molecular weights already include the glycerol backbone; separate addition would double-count it.
- Exact unrounded coefficients are retained in Biomass_Lipids. Reaction List contains model-transfer text rounded to two significant digits, written with decimal commas and ordinary decimal notation; zero-coefficient compounds are omitted.

#### 6.7. Model-species coverage within Level 3

The calculated class amount in Calc and the sum of retained molecular species in Biomass_Lipids are related but not identical. The retained membrane species represent 97.7%, 98.3% and 97.8% of the calculated MEM amount in the autotrophic, mixotrophic and heterotrophic workbooks, respectively; retained TAG species represent 99.9%, 98.9% and 97.9%. This residual reflects analytical fatty-acid categories without a retained model-species row. It is distinct from the expected Level-1-to-Level-3 unit conversion and should be interpreted as model-species coverage, not as a failure of Level-1 mass closure.

### 7. Worked mass-flow example

For the mixotrophic calculation, Ltotal = 0.0425 g gDW⁻¹. Level 1 assigns 65% to MEM and 35% to TAG: mMEM = 0.0425 × 0.65 = 0.027625 g gDW⁻¹ and mTAG = 0.0425 × 0.35 = 0.014875 g gDW⁻¹. Level 2 assigns 14% of MEM to MGDG, so the mixotrophic MGDG mass basis is 0.027625 × 0.14 = 0.0038675 g gDW⁻¹ before division among fatty-acid species and conversion to mmol gDW⁻¹. The remaining eight membrane classes are calculated identically with their own Level-2 fractions. This example demonstrates that Level-2 fractions apply to MEM—not to total biomass and not directly to total lipid.

### 8. Assumptions, justification and required interpretation

| **Decision** | **Why it was necessary** | **How it must be reported** | **Recommended sensitivity test** |
| --- | --- | --- | --- |
| Trophic MEM/TAG fractions | No matched complete stable-pool mass balance was available | Model allocation informed by trophic remodeling; not direct measurement | Vary MEM/TAG by ±10–20 percentage points while preserving closure |
| Nine Level-2 class fractions | No complete matched nine-class dataset across all trophic states | Reconstructed, normalized class distribution | Replace with lipidomics when available; perturb glycolipid versus phospholipid blocks |
| DG excluded | Avoids treating a central intermediate as an independent stable end pool and avoids possible double counting | 43% heterotrophic DG remains source evidence (10.1016/j.biortech.2015.11.014) | Alternative BOF with explicit DG pool |
| FFA excluded | FFA profile informs acyl composition but is not a uniquely defined structural pool | FFA coefficient is zero | Alternative small FFA pool if absolute measurements become available |
| 180 g mol⁻¹ headgroup term | Complete class/species molecular weights unavailable | Single average molecular-weight approximation; no direct DOI | Use class-specific headgroup and backbone masses |
| Shared MEM acyl profile | Class-resolved acyl profiles unavailable | BOF-scale proxy, not intact lipidomics | Test class-specific profiles when measured |

### 9. Additional transparency, uncertainty and reproducibility information

#### 9.1. FAME-to-intact-lipid limitation

Fatty-acid methyl ester (FAME) analysis quantifies derivatized fatty-acyl chains after cleavage and methylation; it does not directly identify the original intact lipid molecular species. Consequently, a FAME profile omits the intact glycerol or other backbone, the class-defining polar headgroup, esterification water losses, the number and positional arrangement of acyl chains, sn-position, double-bond position and geometry unless separately resolved, and the pairing of multiple fatty acids within one lipid molecule. It also cannot by itself determine whether a measured acyl chain originated from TAG, DG, a phospholipid, a glycolipid or a betaine lipid. This is why the workbooks combine measured relative fatty-acid composition with explicit Level-1 pool allocation, Level-2 class allocation and Level-3 molecular-weight reconstruction. The resulting coefficients are BOF-compatible approximations, not a reconstructed intact lipidome. The source Galdieria lipid and fatty-acid measurements are described by Sakurai et al. (DOI: 10.1016/j.biortech.2015.11.014) and López et al. (DOI: 10.1038/s41598-019-46645-3).

#### 9.2. Reconciliation of total-lipid datasets and central-value selection

The older source description derived total lipid as lipid concentration divided by dry-biomass concentration. This gives Sakurai-based values of 0.055, 0.022 and 0.073 g lipid gDW⁻¹ for autotrophic, mixotrophic and heterotrophic cultivation, respectively (DOI: 10.1016/j.biortech.2015.11.014). The central Level-3 BOFs instead use 0.1500, 0.0425 and 0.0375 g gDW⁻¹, selected from the López-supported trophic comparison (DOI: 10.1038/s41598-019-46645-3). The two datasets were not averaged because they differ in strain, culture design, sampling state and analytical workflow. The selected López-informed values provide one internally consistent three-state set, while the Sakurai-derived values are retained as alternative total-lipid scenarios rather than discarded.

| **Trophic state** | **Sakurai-derived alternative** | **Selected central Level 3** | **Central/alternative ratio** | **Interpretation** |
| --- | --- | --- | --- | --- |
| Autotrophic | 0.055 | 0.1500 | 2.73 | Large method/strain sensitivity; use both in scenario analysis |
| Mixotrophic | 0.022 | 0.0425 | 1.93 | Central value is approximately twice the alternative |
| Heterotrophic | 0.073 | 0.0375 | 0.51 | Dataset ordering reverses; especially important uncertainty |

#### 9.3. Meaning and limitation of the 180 g mol⁻¹ term

The 180 g mol⁻¹ term in Inputs!B8 is not the molecular mass of a universal lipid headgroup. MGDG, DGDG, SQDG, PG, PE, PC, PI, PS and DGTS have chemically different headgroups and therefore different intact molecular weights. In the implemented equation, MW_MEM = 92.094 + 2×MW_FA − 2×18.015 + 180, the term acts as one average residual contribution added after the glycerol-plus-two-fatty-acid esterification calculation. It is used only because the available data do not provide matched class- and species-resolved intact-lipid masses. Therefore it should be described as an average molecular-weight proxy, replaced by class-specific values when lipidomics permit, and included in sensitivity analysis. It has no direct experimental DOI.

#### 9.4. Why one fatty-acid profile was applied to all nine membrane classes

The source data provide trophic-state fatty-acid distributions but not a complete matrix resolving every fatty acid within every one of the nine membrane classes under matched conditions (DOIs: 10.1016/j.biortech.2015.11.014; 10.1038/s41598-019-46645-3). Applying one condition-specific MEM profile to all classes preserves the measured aggregate acyl distribution, avoids inventing unsupported class-specific profiles, and keeps the BOF tractable. This assumption distributes each class mass across the same relative acyl pattern; it does not assert biochemical identity among classes. The approach is most appropriate for biomass-scale accounting and least appropriate for questions dependent on class-specific remodeling, organelle localization or particular molecular species.

#### 9.5. Equation-to-workbook reproducibility map

The following references point to the authoritative self-contained trophic-state workbooks: Autotroph_Lipid_Biomass.xlsx, Mixotroph_Lipid_Biomass.xlsx and Heterotroph_Lipid_Biomass.xlsx. All three use the same sheet layout; where a cell range is repeated, it applies to every trophic-state file. Evidence classification and modeling assumptions are documented in Sections 1, 4, 5, 8 and 9 of this supplement because the current workbooks do not contain a Sources_Assumptions sheet.

| **Step or equation** | **Workbook** | **Exact sheet and cells** | **Role** |
| --- | --- | --- | --- |
| Workbook validation dashboard | Corresponding trophic-state workbook | README_Checks!D4:F10 | Live closure and consistency checks; all display PASS |
| Level-1 MEM/TAG fractions and closure | Corresponding trophic-state workbook | 'Level 1'!A4:C5; 'Level 1'!A7:B8 | MEM and TAG fractions; sum and required value |
| Level-2 nine-class fractions and closure | Corresponding trophic-state workbook | 'Level 2'!A4:C12; 'Level 2'!A14:B15 | Nine class fractions; sum and required value |
| Core mass inputs and 180 g mol⁻¹ proxy | Corresponding trophic-state workbook | Inputs!A4:C8; Inputs!A12:C12 | Total lipid, TAG, DG, FFA, MEM and molecular-weight proxy inputs |
| Nine class fractions used by formulas | Corresponding trophic-state workbook | Inputs!A14:C23 | Formula-linked Level-2 fractions |
| FA identity and source percentages | Corresponding trophic-state workbook | Inputs!A26:H58 | Cn:d identities and MEM/DG/FFA/TAG source profiles |
| Profile totals and normalization factors | Corresponding trophic-state workbook | Inputs!A60:G62 | Reported percentage totals and 100/sum normalization factors |
| FA formulas, MWs and normalized profiles | Corresponding trophic-state workbook | Calc!A12:G43 | CnH(2n−2d)O2 approximation plus normalized MEM/DG/FFA/TAG percentages |
| Weighted FA and intact-pool MW proxies | Corresponding trophic-state workbook | Calc!A46:C52 | Weighted MEM/TAG FA MWs, intact MEM/TAG MWs and backbone adjustments |
| Pool mass allocation and molar conversion | Corresponding trophic-state workbook | Calc!A54:D59 | MEM/TAG masses, average MWs, molar amounts and mass closure |
| Class allocation and mmol conversion | Corresponding trophic-state workbook | Calc!A62:D72 | Level-2 fractions, class molar amounts and MEM closure |
| Exact molecular-species coefficients | Corresponding trophic-state workbook | Biomass_Lipids!A5:D14; A19:D27; A32:D40; A45:D53; A58:D66; A71:D79; A84:D92; A97:D105; A110:D118; A123:D132 | Exact MGDG, DGDG, SQDG, PG, PE, PC, PI, PS, DGTS and TAG coefficients |
| Authoritative DG treatment | Corresponding trophic-state workbook | Inputs!A6:C6; README_Checks!D10:F10 | DG input is zero and the validation dashboard confirms exclusion from the stable BOF; no DG reaction row is present |

#### 9.6. Final BOF coefficient summary for all trophic states

The table reports the sum of exact molecular-species coefficients for each non-DG class in Biomass_Lipids (mmol gDW⁻¹). Reaction List is a model-transfer representation rounded to two significant digits, whereas the values below are calculated from the unrounded coefficients. Because each aggregate class has a different reconstructed molecular-weight basis, these molar sums should not be added to infer mass without using the corresponding formulas.

| **Aggregate lipid class** | **AUTO mmol gDW⁻¹** | **MIXO mmol gDW⁻¹** | **HETERO mmol gDW⁻¹** |
| --- | --- | --- | --- |
| MGDG | 0.02247156 | 0.00486148 | 0.0028093 |
| DGDG | 0.03230287 | 0.00729222 | 0.00444807 |
| SQDG | 0.01825815 | 0.00381974 | 0.00187287 |
| PG | 0.00983131 | 0.00243074 | 0.00163876 |
| PE | 0.01966262 | 0.00486148 | 0.00397985 |
| PC | 0.02247156 | 0.00590323 | 0.00444807 |
| PI | 0.00561789 | 0.00138899 | 0.00093643 |
| PS | 0.00280895 | 0.0006945 | 0.00070233 |
| DGTS | 0.00702236 | 0.00347249 | 0.0025752 |
| TAG | 0.04347448 | 0.01698824 | 0.02099237 |
| DG | 0 | 0 | 0 |

Mass-basis check: the selected total-lipid inputs are 0.1500, 0.0425 and 0.0375 g gDW⁻¹. Their MEM/TAG allocations are respectively 0.112500/0.037500, 0.027625/0.014875 and 0.018750/0.018750 g gDW⁻¹. DG and FFA are zero in the stable central BOFs. The general BOF rationale follows Feist and Palsson (DOI: 10.1016/j.mib.2010.03.003).

### 10. Validated references

Sakurai T, Aoki M, Ju X, Ueda T, Nakamura Y, Fujiwara S, Umemura T, Tsuzuki M, Minoda A (2016). Profiling of lipid and glycogen accumulations under different growth conditions in the sulfothermophilic red alga Galdieria sulphuraria. Bioresource Technology 200:861–866. https://doi.org/10.1016/j.biortech.2015.11.014.

López G, Yate C, Ramos FA, Cala MP, Restrepo S, Baena S (2019). Production of polyunsaturated fatty acids and lipids from autotrophic, mixotrophic and heterotrophic cultivation of Galdieria sp. strain USBA-GBX-832. Scientific Reports 9:10791. https://doi.org/10.1038/s41598-019-46645-3.

Feist AM, Palsson BØ (2010). The biomass objective function. Current Opinion in Microbiology 13:344–349. https://doi.org/10.1016/j.mib.2010.03.003.
