## Supplementary Data 1 for "Experimental trophic flexibility guides a compartmentalized genome-scale model of red microalgae *Galdieria sulphuraria*": Galdieria_RNS_DNS_nucleotide.docx

Derivation of DNA (DNS) and RNA (RNS) Nucleotide Fractions for Biomass Objective Function Construction in Galdieria sulphuraria

This document describes the methodology used to determine organism‑specific nucleotide fractions for DNA (DNS) and RNA (RNS) in the thermoacidophilic red alga Galdieria sulphuraria. The resulting nucleotide composition values were used to parameterize the nucleic acid component of the biomass objective function (BOF) in a genome‑scale metabolic model.

### 1. Genome Data Source

Genome sequence data were obtained from the National Center for Biotechnology Information (NCBI) genome repository. All sequence files were downloaded from the publicly available dataset:
https://www.ncbi.nlm.nih.gov/datasets/genome/GCF_000341285.1/

This dataset contains the reference genome assembly for Galdieria sulphuraria and includes the genomic DNA FASTA files and annotated coding sequences (CDS) required for nucleotide composition analysis.

### 2. Reference Genome Publication

Schönknecht, G. et al. (2013). Gene transfer from bacteria and archaea facilitated evolution of an extremophilic eukaryote. Science.
DOI: https://doi.org/10.1126/science.1231707

### 3. Methodology

Nucleotide fractions were derived directly from genome sequence data. Two datasets were analyzed: (i) the complete genomic FASTA file and (ii) coding sequence (CDS) FASTA files extracted from the genome annotation. The genomic FASTA file was used to determine DNA nucleotide composition (DNS), while CDS sequences were used to estimate RNA nucleotide composition (RNS).

### 4. Sequence Processing

Genome FASTA sequences were scanned and occurrences of adenine (A), cytosine (C), guanine (G), and thymine (T) were counted. FASTA headers beginning with '>' were excluded. Ambiguous nucleotides (e.g., N) were ignored.

The total number of nucleotides was calculated as:
N_total = A + C + G + T

Nucleotide fractions were calculated using:
f_X = X / (A + C + G + T)

### 5. DNA Nucleotide Fractions (DNS)

A = 0.31220
T = 0.31081
C = 0.18970
G = 0.18728

### 6. Genome GC Content

The GC content of the genome assembly was calculated as:
GC = (G + C) / (A + T + G + C)

The GC content determined from the genome FASTA sequence was 0.37699 (37.7%).

### 7. RNA Nucleotide Fractions (RNS)

A = 0.30424
U = 0.29894
C = 0.17685
G = 0.21997

### 8. Nucleic Acid Biomass Fractions

Direct experimental measurements of RNA and DNA biomass fractions have not been reported specifically for Galdieria sulphuraria. Therefore, nucleic‑acid biomass fractions used in the biomass objective function were anchored to experimentally derived values reported for microalgae.

A comprehensive meta‑analysis by Finkel et al. (2016) analyzed published measurements of microalgal macromolecular composition and reported median nucleic‑acid biomass fractions of:

RNA = 5.7% of dry biomass (0.057 g gDW⁻¹)
DNA = 1.0% of dry biomass (0.010 g gDW⁻¹)

Source:
Finkel, Z.V. et al. (2016). Phylogenetic diversity in the macromolecular composition of microalgae. PLoS ONE.
DOI: https://doi.org/10.1371/journal.pone.0155977

These values represent experimentally measured nucleic‑acid content compiled across multiple microalgal species grown under nutrient‑sufficient exponential growth conditions.

It is important to note that ranges sometimes used in metabolic modeling studies (e.g., RNA 0.05–0.07 g gDW⁻¹ and DNA 0.01–0.015 g gDW⁻¹) represent modeling approximations centered around literature medians rather than directly measured intervals from this study.

### 9. Application in Biomass Objective Function

DNA precursor reaction:
0.31220 dATP + 0.31081 dTTP + 0.18970 dCTP + 0.18728 dGTP → DNA

RNA precursor reaction:
0.30424 ATP + 0.29894 UTP + 0.17685 CTP + 0.21997 GTP → RNA
