## Supplementary Data 1 for "Experimental trophic flexibility guides a compartmentalized genome-scale model of red microalgae *Galdieria sulphuraria*": Pigments_trophic_data.docx

Supplementary methods: trophic-state pigment biomass objective functions

*Galdieria sulphuraria — calculation rationale, evidence boundary and workbook map*

Companion to four condition-specific pigment BOF workbooks

### 1. Purpose and evidence boundary

This supplement documents how trophic-state pigment biomass-objective-function (BOF) coefficients were assembled for autotrophic, mixotrophic, glucose-heterotrophic and glycerol-heterotrophic growth. The BOF module represents pigment demand as four model metabolites: zeaxanthin, chlorophyll a, β-carotene and 3Z-phycocyanobilin.

The literature directly supports trophic regulation and relative pigment differences, but it does not provide one matched, complete, dry-weight-normalized dataset for all four compounds under all four modeled conditions. Therefore, published relative profiles, derived scaling and model-selected total pigment fractions are separated explicitly. The selected totals are BOF inputs aligned with the total-biomass reconstruction; they are not presented as new measurements.

### 2. Modeled pigment set and compartment mapping

Table S1. Pigments retained in the biomass module.

| **Pigment** | **Model metabolite** | **MW (g mol⁻¹)** | **Compartment** |
| --- | --- | --- | --- |
| Zeaxanthin | m_CPD1F-130_Thyl[t] | 568.87 | Thylakoid |
| Chlorophyll a | m_CHLOROPHYLL-A_Plas[p] | 893.49 | Plastid |
| β-Carotene | m_CPD1F-129_Thyl[t] | 536.87 | Thylakoid |
| 3Z-Phycocyanobilin | m_3Z-PHYCOCYANOBILIN_Plas[p] | 586.00 | Plastid |

Zeaxanthin and β-carotene are represented by the model identifiers CPD1F-130 and CPD1F-129, respectively. Chlorophyll a and 3Z-phycocyanobilin are represented explicitly. The final pseudo-reaction produces m_PIGMENT_TOTAL_POOL[c], which is incorporated into the higher-level BOF.

### 3. Biological evidence and trophic-state reconstruction

#### 3.1. Autotrophic reference

Autotrophy was retained as the pigment-rich reference state. Perez Saura et al. reported that phototrophic cells contained substantially more chlorophyll a, zeaxanthin and β-carotene than heterotrophic cells, while Abiusi et al. independently documented strong trophic changes in the cellular light-absorption cross section (DOIs: 10.3389/fpls.2022.978246; 10.1016/j.algal.2021.102513).

#### 3.2. Mixotrophic scaling

The mixotrophic profile was scaled from the autotrophic reference using the reported mean absorption-cross-section ratio 62/114 ≈ 0.54. This is a pigment-pool proxy: it supports the direction and approximate magnitude of the reduction, but it is not a compound-specific chemical assay for every pigment.

#### 3.3. Carbon-source-specific heterotrophy

Glucose and glycerol heterotrophy were kept separate because pigment repression depends on the organic carbon source. Perez Saura et al. reported approximately twofold phototrophic-to-glycerol differences for chlorophyll a and carotenoids and substantially stronger suppression with glucose; phycocyanin showed still larger changes (DOI: 10.3389/fpls.2022.978246). These ratios support condition-specific reconstruction rather than one universal heterotrophic pigment profile.

Table S2. Evidence status of the principal reconstruction inputs.

| Input or decision | Evidence class | Interpretation |
| --- | --- | --- |
| AUTO:glycerol-HET and AUTO:glucose-HET pigment ratios | Direct relative evidence | Supports carbon-source-specific trophic scaling; Pérez Saura et al. (2022). |
| AUTO:MIXO absorption-cross-section ratio | Direct proxy measurement | Supports approximate whole-pigment-pool scaling; Abiusi et al. (2021). |
| Selected total pigment fractions | Model-selected BOF inputs | Synchronized across the four workbooks, Macro_Biomass.docx and this supplement; not direct measurements. |
| 4.634% phycocyanobilin conversion | Implemented structural approximation | Three PCB chromophores per approximately 38 kDa αβ monomer; prevents double counting of the protein backbone. |
| Pigment molecular weights | Chemical constants used by the model | Convert allocated mass fractions to mmol gDW⁻¹. |

### 4. Avoiding phycocyanin protein double counting

C-phycocyanin is a phycobiliprotein. Adding the complete holoprotein to the pigment BOF while its amino-acid mass is already represented in the protein BOF would double-count the protein backbone. Accordingly, only the covalently attached chromophore is represented as a pigment term.

mPCB = mC-phycocyanin × fPCB

The implemented central conversion is fPCB = (3 × 587) / 38,000 = 0.0463421, or 4.634% of phycocyanin holoprotein mass. The 3–5% interval is retained only for sensitivity analysis. This structural conversion excludes the protein backbone and is not a matched dry-weight measurement. The source used dry-biomass normalization for chlorophyll and carotenoids, but soluble-protein normalization for phycocyanin.

### 5. Selected condition-specific inputs

Table S3. Pigment profiles and selected total-pigment fractions implemented in the workbooks.

| Condition | Zea | Chl a | β-Car | PCB | Profile total (mg gDW⁻¹) | Selected Lpig (g gDW⁻¹) |
| --- | --- | --- | --- | --- | --- | --- |
| Autotrophic | 0.65 | 8.25 | 0.45 | 3.186020 | 12.536020 | 0.0121 |
| Mixotrophic | 0.35 | 4.45 | 0.24 | 1.714658 | 6.754658 | 0.0065 |
| Heterotrophic (glucose) | 0.036 | 0.5 | 0.021 | 0.023171 | 0.580171 | 0.000580171 |
| Heterotrophic (glycerol) | 0.325 | 2.65 | 0.161 | 0.137868 | 3.273868 | 0.00485 |

The profile columns preserve the condition-specific relative composition. Lpig is the selected total pigment fraction applied in the fractional BOF reaction. Small differences between a profile sum and 1000 × Lpig are resolved by normalizing the profile before applying Lpig.

### 6. Conversion to model-ready BOF coefficients

#### 6.1. Step 1: normalize the pigment mass profile

wi,t = Di,t / Σj Dj,t

Di,t is the source-profile value in mg pigment gDW⁻¹ for pigment i and condition t. The normalized mass fraction wi,t preserves relative composition and sums to one.

#### 6.2. Step 2: apply the selected total pigment fraction

mi,t = Lpig,t × wi,t

Lpig,t is the model-selected pigment fraction in g pigment gDW⁻¹. The resulting mi,t is the mass of pigment i assigned per gDW.

#### 6.3. Step 3: convert mass to amount

ci,t = 1000 × Lpig,t × wi,t / MWi

ci,t is the fractional BOF coefficient in mmol gDW⁻¹ and MWi is in g mol⁻¹. The factor 1000 converts mol to mmol. These ci,t values, not the source-profile mg values, are used in the biomass-scaled reaction.

#### 6.4. Step 4: construct the composition-only reaction

xi,t = ci,t / Σj cj,t

xi,t is a normalized molar composition coefficient. The composition-only reaction is useful for inspecting relative pigment composition, but it does not encode the total pigment mass. The fractional reaction is authoritative for BOF integration.

### 7. Worked example

For autotrophic chlorophyll a, D = 8.25 mg gDW⁻¹ and the reconciled profile total is 12.536020 mg gDW⁻¹. With Lpig = 0.0121 g gDW⁻¹ and MW = 893.49 g mol⁻¹:

cChl a,AUTO = 1000 × 0.0121 × (8.25/12.536020) / 893.49 = 0.008912303 mmol gDW⁻¹

The same calculation is applied to the other three pigments and to each modeled condition.

### 8. Final BOF coefficients

Table S4. Exact condition-specific pigment coefficients from Calc!E2:E5 (mmol gDW⁻¹).

| Condition | Zea | Chl a | β-Car | PCB | Σ coefficients |
| --- | --- | --- | --- | --- | --- |
| Autotrophic | 0.001102874 | 0.008912303 | 0.000809038 | 0.005247791 | 0.016072007 |
| Mixotrophic | 0.000592059 | 0.004792701 | 0.000430182 | 0.002815723 | 0.008630664 |
| Heterotrophic (glucose) | 0.000063283 | 0.000559603 | 0.000039116 | 0.000039541 | 0.000701543 |
| Heterotrophic (glycerol) | 0.000846352 | 0.004393765 | 0.000444260 | 0.000348534 | 0.006032911 |

The sum of coefficients is an amount total (mmol gDW⁻¹), not the selected pigment mass fraction. It differs among conditions because the total pigment mass and the molecular-weight-weighted composition both differ.

### 9. Aggregate reaction structure and model interpretation

For condition t, the biomass-scaled pigment reaction has the following generic structure:

cZea,t Zea + cChl,t Chl a + cβCar,t β-Car + cPCB,t PCB → m_PIGMENT_TOTAL_POOL[c]

- Use Reaction List!C3 for the condition-specific fractional reaction incorporated into the BOF.
- Use Reaction List!C2 only as a normalized composition representation; it omits the selected total pigment fraction.
- The optional sink in Reaction List!C4 is not the biomass synthesis reaction.
- The pigment-pool pseudo-metabolite is a bookkeeping construct; its coefficient is determined by the higher-level total BOF architecture.
- Do not add whole C-phycocyanin as a separate pigment metabolite when its protein backbone is already included in the protein fraction.

### 10. Uncertainty and required interpretation

Table S5. Main uncertainties and recommended sensitivity analyses.

| Uncertain element | Why uncertainty remains | Recommended test |
| --- | --- | --- |
| Selected Lpig values | Central BOF totals are model selections, not one matched four-condition measurement. | Vary each total across its documented evidence envelope while preserving within-condition composition. |
| AUTO:MIXO scaling | Absorption cross section is a whole-pool optical proxy. | Test alternative mixotrophic scale factors around 0.54. |
| Heterotrophic scaling | Relative ratios depend on carbon source and normalization basis. | Retain separate glucose and glycerol scenarios; do not average them by default. |
| PCB conversion | The implemented central structural factor is 4.634%. | Test 3–5% while keeping the protein backbone excluded; do not substitute the sensitivity bounds into the central workbook. |
| Compound mapping | Model identifiers abstract pigment localization. | Use thylakoid for zeaxanthin/β-carotene, plastid for chlorophyll a/PCB and cytosol only for the aggregate pigment pool. |

### 11. Equation-to-workbook reproducibility map

All four condition-specific workbooks share the same six-sheet structure: Inputs, Calc, Reaction List, Metabolite List, Sources_Assumptions and VALIDATION_STATUS. The ranges below apply to every workbook. Formula cells retain full precision; Reaction List stores model-transfer text.

Table S6. Exact workbook locations for reproducing the pigment BOF calculations.

| Step or quantity | Sheet and cells | Role |
| --- | --- | --- |
| Selected total pigment fraction Lpig,t | Inputs!A2:C2 | Condition-specific BOF mass input (g gDW⁻¹). |
| Pigment identity, model ID and molecular weight | Inputs!A4:C8 | Four retained pigments and MW values. |
| Condition-specific profile Di,t | Inputs!D4:D8 | Profile in mg gDW⁻¹; D8 is formula-derived PCB. |
| Normalized mass fractions wi,t | Inputs!E4:E8 | D_i/ΣD; header plus four formulas. |
| Profile and carotenoid totals | Inputs!A11:D11 | Closure and reporting checks. |
| PCB structural conversion | Inputs!A14:C19 | Chromophore count, MWs, implemented factor and holoprotein proxy. |
| Fractional coefficients ci,t | Calc!A1:E5 | Identity mapping and mmol gDW⁻¹ calculation. |
| Composition-only molar fractions xi,t | Calc!F1:F5 | Normalized molar composition. |
| Coefficient total | Calc!A8:B8 | Sum of Calc!E2:E5. |
| Composition-only reaction | Reaction List!A2:H2 | Normalized molar reaction; Lpig not applied. |
| Fractional BOF reaction | Reaction List!A3:H3 | Model-ready reaction with Lpig applied. |
| Pigment-pool sink | Reaction List!A4:H4 | Optional sink; not the biomass synthesis equation. |
| Metabolite identities and compartments | Metabolite List!A1:F6 | Thylakoid/plastid assignments plus cytosolic aggregate pool. |
| Evidence and assumptions | Sources_Assumptions!A1:D7 | Evidence boundary and implemented assumptions in all four workbooks. |
| Release status | VALIDATION_STATUS!A1:B9 | Current filename, role and validation status in all four workbooks. |

The glucose-heterotrophic workbook is now included and follows the same calculation and metadata structure as the other three workbooks.

### 12. Validated references

Perez Saura P, Adamo S, Bounnit T, et al. (2022). Cell adaptation of the extremophilic red microalga Galdieria sulphuraria to heterotrophic growth on glycerol. Frontiers in Plant Science 13:978246. https://doi.org/10.3389/fpls.2022.978246.

Abiusi F, Trompetter E, Hoenink H, Wijffels RH, Janssen M (2021). Autotrophic and mixotrophic biomass production of the acidophilic Galdieria sulphuraria ACUF 64. Algal Research 60:102513. https://doi.org/10.1016/j.algal.2021.102513.

Barone R, De Napoli L, Mayol L, et al. (2020). Autotrophic and heterotrophic growth conditions modify biomolecule production in the microalga Galdieria sulphuraria (Cyanidiophyceae, Rhodophyta). Marine Drugs 18:169. https://doi.org/10.3390/md18030169.

Feist AM, Palsson BØ (2010). The biomass objective function. Current Opinion in Microbiology 13:344–349. https://doi.org/10.1016/j.mib.2010.03.003.
