## Supplementary Data 1 for "Experimental trophic flexibility guides a compartmentalized genome-scale model of red microalgae *Galdieria sulphuraria*": Galdieria_Protein_Biomass_Methods.docx

Calculation of Protein (Amino Acid) Fractions for Biomass Objective Function in Galdieria sulphuraria

This document describes the construction of protein biomass pseudoreactions for the Galdieria sulphuraria model. Literature amino-acid profiles were converted to composition-only coefficients (mmol per gram protein residue), and the publication-selected profiles were calibrated before applying the measured protein fractions in the biomass objective.

### 1. Data Sources

Autotrophic and mixotrophic amino‑acid compositions were obtained from:
Graziani et al., 2023. Amino acid profile and protein bioaccessibility of two Galdieria sulphuraria strains. DOI: 10.1016/j.ifset.2023.103287.

Heterotrophic amino‑acid compositions were obtained from:
Abiusi et al., 2024. Amino acid content of Galdieria sulphuraria cultivated heterotrophically at different growth phases. (Table 3).

### 2. Amino Acid Composition Data

For autotrophic and mixotrophic conditions, amino‑acid concentrations were reported in mg amino acid per g protein for three strains (SAG 108.79, ACUF 064, and Abiusi 064). Separate biomass reactions were constructed for each strain under each trophic condition.

For heterotrophic conditions, amino‑acid concentrations were reported for two strains (SAG 108.79 and ACUF 074) across three growth phases:
• Exponential (Exp)
• Early stationary (ES)
• Stationary phase (S1)
The average amino‑acid fraction for each phase was calculated across the two strains.

### 3. Treatment of Missing Amino Acids

Tryptophan (Trp) values were not detected (n.d.) in the heterotrophic dataset. To maintain completeness of the protein composition, Trp was assumed to represent 1% of the total amino‑acid pool for each condition. The value was calculated as:

Trp = 0.01 × (sum of all reported amino acids for that condition).

### 4. Conversion to Biomass Stoichiometric Coefficients

For each publication-selected profile, the reported amino-acid values (mg g⁻¹ protein) were converted to raw mmol coefficients and then rescaled within the protein category. Let aᵢ be the source mmol coefficient after the original two-significant-digit conversion and Mᵢ the amino-acid molecular weight. The long-polymer peptide-residue mass of the source vector is S = Σ[aᵢ(Mᵢ − 18.01528)]/1000 g. The calibrated composition-only coefficient is cᵢ = aᵢ/S mmol per gram protein residue. This preserves all relative amino-acid proportions. The HETERO exponential and MIXO SAG 108.79 profiles are the publication-selected profiles; other profiles remain historical reference variants in the source workbooks.

### 5. Biomass Reaction Construction

The calibrated publication pool reaction is Σ(cᵢ × AAᵢ) → m_PROTEIN_TOTAL_POOL[c] + [Σcᵢ] m_WATER_Cyto[c]. The water term uses a long-chain peptide approximation (one condensation water per incorporated residue; terminal-group correction is not resolved). Thus the calibrated composition-only reaction represents approximately one gram of protein residue, and the final measured protein fraction is applied once in the biomass assembly. The fractional workbook illustration uses its historical literature protein fraction rather than the measured publication fraction.

### 6. Generated Biomass Reactions

The workbooks contain paired composition-only (*_F) and historical reference-fraction (*_G) reactions for each source profile. Only the publication-selected HETERO exponential and MIXO SAG 108.79 pairs were recalibrated here; all other source variants remain unchanged. In the publication model, the calibrated *_F protein reaction supplies the pool and the measured protein fraction is applied once in the runtime BOF. The *_G illustrations already include their workbook reference fraction and must not be used in parallel.

### 7. Application in Genome‑Scale Modeling

For the publication analyses, use the calibrated HETERO exponential or MIXO SAG 108.79 composition-only reaction and apply its measured protein fraction once in the runtime BOF: 0.4016 g gDW⁻¹ for HETERO and 0.4147 g gDW⁻¹ for MIXO. The *_G source-workbook examples retain their historical reference fractions (0.335 and 0.575) and are not used in parallel. AUTO and the other source profiles are unchanged reference variants. The measured four-category total remains 0.9000 g gDW⁻¹; the unmeasured 0.1000 is not redistributed.
