## Supplementary Data 2 for "Experimental trophic flexibility guides a compartmentalized genome-scale model of red microalgae *Galdieria sulphuraria*": Experimental_BOF_nonrenormalized_methods_note.pdf

### Experimental BOF reconstruction — publication methods note

The baseline TOTAL_BIOMASS.xlsx resource preserves the literature-informed reference BOFs and the relative molecular distributions of amino acids, carbohydrate species, lipid species and RNA/DNA components. In the updated MATLAB analyses, experimental MIXO and HETERO mass fractions are applied to these relative reference distributions without rescaling the measured macromolecular pool totals to 1.000 g/gDW.

Measured MIXO: protein 0.4147, carbohydrate 0.3377, lipid 0.0701, nucleic acids 0.0775 g/gDW. Measured HETERO: protein 0.4016, carbohydrate 0.3609, lipid 0.0610, nucleic acids 0.0765 g/gDW. Both macromolecular totals equal 0.9000 g/gDW. Reference-derived pigment contributions are handled separately and their potential overlap with measured categories requires checking.

The remaining approximately 10% of dry mass is compositionally unresolved, not a measured ±10% statistical error bar. The missing composition is not redistributed among measured pools. A unit of BOF flux has not independently been proven to correspond to exactly 1 g of total dry biomass; absolute specific-growth-rate interpretation remains subject to biomass-mass calibration. AUTO uses the literature-informed AUTO reference BOF.

All four analysis scripts and their executable inputs and output workbooks are contained in Supplementary Data 3. This note describes the implemented analytical BOF, not a replacement of the unchanged SBML model reactions.
