## Supplementary Data 2 for "Experimental trophic flexibility guides a compartmentalized genome-scale model of red microalgae *Galdieria sulphuraria*": Galdieria_BOF_Integration_User_Manual.docx

**User Manual for Integrating Biomass Objective Functions into the Galdieria Genome-Scale Model**

*TOTAL_BIOMASS workbook → COBRA Toolbox MATLAB model → SBML-FBC*

Scope: autotrophic, mixotrophic and heterotrophic BOFs; macromolecular pools; GAM/NGAM; validation; export and round-trip testing

### 1. Purpose and scope

This manual describes the complete procedure for transferring biomass-composition data from the TOTAL_BIOMASS workbook into an existing Galdieria sulphuraria COBRA Toolbox model, saving the integrated MATLAB model, and exporting validated SBML Level 3 FBC files. The workflow is intended for modular biomass objective functions containing separate protein, carbohydrate, lipid, RNA, DNA and pigment pools for autotrophic, mixotrophic and heterotrophic growth.

The procedure is designed to prevent the most common BOF integration errors: applying mass fractions twice, omitting intermediate pool reactions, mixing trophic-state reactions, introducing inconsistent metabolite identifiers, combining GAM and NGAM, exporting an objective-free SBML file, and accepting an SBML export without testing the round trip.

**Core rule:** Every biomass fraction must be applied exactly once. A macromolecular coefficient must be placed either in the precursor-pool reaction or in the final biomass-assembly reaction, but not in both.

### 2. Workflow overview

| **Step** | **Action** | **Expected result** |
| --- | --- | --- |
| 1 | Freeze the authoritative workbook | Confirm the latest TOTAL_BIOMASS version, source sheets, trophic means, metabolite identifiers and final reactions. |
| 2 | Choose one BOF architecture | Use normalized pool reactions plus a fraction-weighted final BOF, or fraction-scaled pool reactions plus an unweighted assembly reaction. |
| 3 | Prepare a clean reaction-transfer table | Create one row per reaction with COBRA-compatible identifiers, reaction strings, bounds, objective coefficient and subsystem. |
| 4 | Load and verify the base model | Use readCbModel, confirm compartments, inspect current objectives and remove only obsolete BOF reactions. |
| 5 | Add missing pseudo-metabolites | Add macromolecular pools and final biomass metabolites before inserting reactions. |
| 6 | Insert pool and assembly reactions | Add all reactions required for AUTO, MIXO and HETERO conditions. |
| 7 | Implement GAM and NGAM correctly | Keep GAM inside biomass synthesis and NGAM as a separate constrained ATP-maintenance reaction. |
| 8 | Build condition-specific model variants | Activate one trophic objective and apply the correct uptake, light, gas and nutrient constraints. |
| 9 | Validate production and stoichiometry | Test individual pools, the final BOF, bounds, objective, duplicates, blocked reactions and mass/charge balance. |
| 10 | Save, export and reload | Save MAT files, export SBML-FBC, reload each SBML file and confirm that growth and objective are preserved. |

### 3. Required files and software

| **Required item** | **Purpose** |
| --- | --- |
| Authoritative TOTAL_BIOMASS workbook | Source of macromolecular composition, pool reactions, final BOFs, GAM and trophic-state means. |
| Current Galdieria model (.mat or .xml) | Base network to which BOF reactions will be added. |
| MATLAB | Execution environment. |
| COBRA Toolbox | Model loading, reaction manipulation, optimization and export. |
| Linear-programming solver | GLPK, CPLEX, Gurobi or another COBRA-compatible solver. |
| libSBML support | Required by COBRA Toolbox for reliable SBML import and export. |
| Versioned output folder | Stores scripts, logs, MAT files, SBML files and validation summaries. |

initCobraToolbox(false);
changeCobraSolver('glpk', 'LP');

% Confirm that COBRA Toolbox is available.
which readCbModel
which addReaction
which writeCbModel
which optimizeCbModel

### 4. Understand the BOF architecture before editing the model

A modular BOF contains at least two reaction layers. The first layer produces generic macromolecular pools. The second layer assembles those pools into a trophic-state-specific biomass product. If the final biomass reaction consumes a pool metabolite, the reaction that produces that pool must also be included in the model.

#### 4.1. Layer 1: macromolecular pool reactions

| **Pool** | **Typical product metabolite** | **Examples of consumed precursors** |
| --- | --- | --- |
| Protein | m_PROTEIN_TOTAL_POOL[c] | Twenty amino acids and, where used, protein-associated cofactors. |
| Carbohydrate | m_CARBOHYDRATE_TOTAL_POOL[c] | Storage glycogen, structural glucan and structural mannan residue. |
| Lipid | m_LIPID_TOTAL_POOL[c] | State-specific membrane and storage lipid metabolites. |
| RNA | m_RNA_TOTAL_POOL[c] | Ribonucleoside triphosphates or RNA nucleotide pool. |
| DNA | m_DNA_TOTAL_POOL[c] | Deoxyribonucleoside triphosphates or DNA nucleotide pool. |
| Pigment | m_PIGMENT_TOTAL_POOL[c] | Chlorophyll a, zeaxanthin, beta-carotene and phycocyanobilin. |

#### 4.2. Layer 2: final biomass assembly

The final reaction consumes the macromolecular pools, adds growth-associated ATP hydrolysis, and forms a trophic-state-specific biomass product. Only the chosen final biomass reaction should carry the active objective coefficient during a simulation.

m_PROTEIN_TOTAL_POOL[c]
+ m_CARBOHYDRATE_TOTAL_POOL[c]
+ m_LIPID_TOTAL_POOL[c]
+ m_RNA_TOTAL_POOL[c]
+ m_DNA_TOTAL_POOL[c]
+ m_PIGMENT_TOTAL_POOL[c]
+ GAM ATP and water
-> ADP + Pi + H+ + m_TOTAL_BIOMASS_STATE[c]

#### 4.3. Choose one coefficient architecture

| **Architecture** | **Pool reaction** | **Final assembly** |
| --- | --- | --- |
| A. Normalized pool (recommended) | Defines internal composition of one unit of the pool. | Consumes the pool using its g gDW^-1 macromolecular fraction. |
| B. Fraction-scaled pool | Already contains the full trophic-state macromolecular amount. | Consumes exactly 1 unit of the already-scaled pool. |

**Do not mix architectures:** If a pool reaction already represents 0.195 g carbohydrate gDW^-1 and the final BOF also multiplies the carbohydrate pool by 0.195, the carbohydrate demand is applied twice.

### 5. Freeze and document the authoritative TOTAL_BIOMASS data

Before any MATLAB work, record the workbook version, source sheet, source row, selected scenario and date. The workbook should distinguish minimum, maximum and mean BOFs and should identify the active trophic state. Use the mean BOF for the standard model unless a sensitivity or uncertainty analysis is being performed.

| **Field** | **AUTO** | **MIXO** | **HETERO** | **Required check** |
| --- | --- | --- | --- | --- |
| Protein mean | from Master Composition | from Master Composition | from Master Composition | Fractions reflect current protein evidence. |
| Carbohydrate mean | 0.195 | 0.225 | 0.230 | Structural carbohydrate is part of the total, not added on top. |
| Lipid mean | from Master Composition | from Master Composition | from Master Composition | Use the current level-3 lipid BOF. |
| RNA mean | from Master Composition | from Master Composition | from Master Composition | Same nucleotide convention across states. |
| DNA mean | from Master Composition | from Master Composition | from Master Composition | Same nucleotide convention across states. |
| Pigment mean | 0.0121 | 0.0065 | selected heterotrophic mean | Use phycocyanobilin, not whole phycocyanin. |
| GAM | 30 | 30 | 30 | mmol ATP gDW^-1 in final BOF. |
| NGAM | separate | separate | separate | Do not place NGAM inside the biomass reaction. |

### 6. Prepare the reaction-transfer table

Create a clean transfer sheet containing only the reactions that will be inserted. The safest approach is to copy the authoritative reaction rows into a new sheet named BOF_Transfer. Each reaction must use exactly the metabolite identifiers found in the model. Do not rely on display names.

| **Column** | **Required content** | **Example** | **Validation** |
| --- | --- | --- | --- |
| Abbreviation | Unique reaction ID | TOTAL_BIOMASS_AUTO_MEAN | Must not duplicate an active reaction. |
| Description | Human-readable explanation | Autotrophic mean total biomass objective | State and scenario must be explicit. |
| Reaction | COBRA reaction string | A + B -> C | All metabolite IDs must exist or be added. |
| GPR | Normally blank for BOF pseudo-reactions |  | Do not invent genes for assembly reactions. |
| Lower bound | Usually 0 | 0 | Biomass synthesis is irreversible. |
| Upper bound | Usually 1000 | 1000 | Use model-wide bound convention. |
| Objective | 1 only for the selected objective; otherwise 0 | 0 or 1 | Only one objective in each condition model. |
| Subsystem | Biomass synthesis | Biomass synthesis | Use one consistent subsystem name. |
| Source workbook | Exact source file/version | TOTAL_BIOMASS_recalculated_vXX.xlsx | Preserve provenance. |

### 7. Standardize identifiers and compartments

Reaction strings must use the exact metabolite IDs stored in model.mets. A metabolite name is not an identifier. Compartment suffixes and SBML compartment symbols must be consistent. In the current Galdieria convention, cytosol, mitochondrion, plastid, thylakoid and peroxisome are represented by c, m, p, t and x, respectively.

| **Compartment** | **Symbol** | **Example metabolite ID** |
| --- | --- | --- |
| Cytosol | c | m_ATP_Cyto[c] |
| Mitochondrion | m | m_ATP_Mito[m] |
| Plastid | p | m_CHLOROPHYLL-A_Plas[p] |
| Thylakoid | t | m_CPD1F-130_Thyl[t] |
| Peroxisome | x | model-specific [x] metabolite |

```
requiredMets = {
    'm_PROTEIN_TOTAL_POOL[c]'
    'm_CARBOHYDRATE_TOTAL_POOL[c]'
    'm_LIPID_TOTAL_POOL[c]'
    'm_RNA_TOTAL_POOL[c]'
    'm_DNA_TOTAL_POOL[c]'
    'm_PIGMENT_TOTAL_POOL[c]'
};

missingMets = setdiff(requiredMets, model.mets);
disp(missingMets);
```

### 8. Load and inspect the base COBRA model

initCobraToolbox(false);
changeCobraSolver('glpk', 'LP');

model = readCbModel('COMBINED_FULL_METAB_DATA_SBML_5.xml');

fprintf('Metabolites: %d\n', numel(model.mets));
fprintf('Reactions: %d\n', numel(model.rxns));
fprintf('Genes: %d\n', numel(model.genes));

verificationBefore = verifyModel(model);
currentObjectives = model.rxns(model.c ~= 0);
disp('Current objective reactions:');
disp(currentObjectives);

Save an untouched backup before editing. Every integration script should create a new model variable rather than overwriting the only copy of the base model.

baseModel = model;
save('Galdieria_base_before_BOF.mat', 'baseModel', '-v7.3');

### 9. Identify and remove obsolete BOF reactions

Search broadly, but remove narrowly. A text search for BIOMASS, TOTAL_POOL or the old reaction IDs is useful for discovery; however, only explicitly reviewed obsolete reactions should be deleted. Do not remove valid precursor pathways or exchange reactions merely because their names contain biomass-related terms.

```
candidateRxns = model.rxns(contains(upper(model.rxns), ...
    {'BIOMASS','TOTAL_POOL','PIGMENT_TOTAL','CARBOHYDRATE_TOTAL'}));
disp(candidateRxns);

rxnsToRemove = {
    'OLD_TOTAL_BIOMASS_AUTO'
    'OLD_TOTAL_BIOMASS_MIXO'
    'OLD_TOTAL_BIOMASS_HETERO'
};

existingToRemove = intersect(rxnsToRemove, model.rxns);
if ~isempty(existingToRemove)
    model = removeRxns(model, existingToRemove);
end
```

### 10. Add missing pool and biomass pseudo-metabolites

Pseudo-metabolites connect the modular BOF layers. They may intentionally have no ordinary elemental formula, but they should have clear names and charges. Real biochemical precursors must retain valid chemical formulas and charges.

```
if ~ismember('m_CARBOHYDRATE_TOTAL_POOL[c]', model.mets)
    model = addMetabolite(model, ...
        'm_CARBOHYDRATE_TOTAL_POOL[c]', ...
        'metName', 'Total carbohydrate biomass pool', ...
        'metFormula', '', ...
        'Charge', 0);
end

if ~ismember('m_TOTAL_BIOMASS_AUTO_MEAN[c]', model.mets)
    model = addMetabolite(model, ...
        'm_TOTAL_BIOMASS_AUTO_MEAN[c]', ...
        'metName', 'Autotrophic mean biomass', ...
        'metFormula', '', ...
        'Charge', 0);
end
```

**Recommended annotation:** Mark model-created pool metabolites and pseudo-reactions in Notes or annotation fields so they are not mistaken for MetaCyc or KEGG compounds.

### 11. Add macromolecular pool reactions

Add the macromolecular pool reactions before the final BOF. The example below uses the normalized-pool architecture. Replace the example coefficients with the authoritative values from the current workbook.

#### 11.1. Carbohydrate pool example

The total carbohydrate fraction is partitioned into intracellular storage glycogen and structural wall carbohydrate. For the current mean autotrophic composition, the documented mass partition is 0.175 glycogen, 0.016 structural glucan and 0.004 structural mannan within a total of 0.195 g gDW^-1. When converting polymer mass to mmol of anhydrohexose repeat units, use approximately 162.14 g mol^-1.

```
model = addReaction(model, ...
    'CARBOHYDRATE_TOTAL_POOL_BIO_AUTOTROPH_NOFRAC', ...
    'reactionFormula', ...
    ['1.08 m_STARCH_Cyto[c] + ' ...
     '0.099 m_1-3-beta-D-Glucans_Cyto[c] + ' ...
     '0.025 m_MANNAN-RESIDUE_Cyto[c] -> ' ...
     'm_CARBOHYDRATE_TOTAL_POOL[c]'], ...
    'lowerBound', 0, ...
    'upperBound', 1000, ...
    'subSystem', 'Biomass synthesis');
```

**Mannan donor:** Do not place GDP-mannose directly into the final carbohydrate pool without releasing GDP. Prefer a mannan-residue or structural-mannan pseudo-metabolite connected through a separate polymerization reaction.

#### 11.2. Pigment pool example

```
model = addReaction(model, ...
    'PIGMENT_TOTAL_POOL_BIO_AUTOTROPH_NOFRAC', ...
    'reactionFormula', ...
    ['0.054 m_CPD1F-130_Thyl[t] + ' ...
     '0.68 m_CHLOROPHYLL-A_Plas[p] + ' ...
     '0.037 m_CPD1F-129_Thyl[t] + ' ...
     '0.23 m_3Z-PHYCOCYANOBILIN_Plas[p] -> ' ...
     'm_PIGMENT_TOTAL_POOL[c]'], ...
    'lowerBound', 0, ...
    'upperBound', 1000, ...
    'subSystem', 'Biomass synthesis');
```

Whole phycocyanin must not be consumed by the pigment BOF because its apoprotein is already represented in the protein pool. The pigment reaction should contain only the phycocyanobilin chromophore contribution.

#### 11.3. Other pools

Use the same pattern for protein, lipid, RNA and DNA. For each pool reaction, confirm that all required precursors can be synthesized in the relevant compartment and that no coefficient has already been multiplied by the final macromolecular mass fraction.

### 12. Add final trophic-state biomass reactions

Create one final mean BOF for each trophic condition. Minimum and maximum variants may also be retained for sensitivity analysis, but they should not be active objectives in the same simulation.

```
autoBOF = [ ...
    '0.615 m_PROTEIN_TOTAL_POOL[c] + ' ...
    '0.195 m_CARBOHYDRATE_TOTAL_POOL[c] + ' ...
    '0.150 m_LIPID_TOTAL_POOL[c] + ' ...
    '0.057 m_RNA_TOTAL_POOL[c] + ' ...
    '0.010 m_DNA_TOTAL_POOL[c] + ' ...
    '0.0121 m_PIGMENT_TOTAL_POOL[c] + ' ...
    '30 m_ATP_Cyto[c] + 30 m_WATER_Cyto[c] -> ' ...
    '30 m_ADP_Cyto[c] + 30 m_Pi_Cyto[c] + ' ...
    '30 m_PROTON_Cyto[c] + m_TOTAL_BIOMASS_AUTO_MEAN[c]'];

model = addReaction(model, ...
    'TOTAL_BIOMASS_AUTO_MEAN', ...
    'reactionFormula', autoBOF, ...
    'lowerBound', 0, ...
    'upperBound', 1000, ...
    'subSystem', 'Biomass synthesis');
```

**Use authoritative coefficients:** The numbers above illustrate the implementation pattern. Replace any coefficient that differs from the latest Master Composition sheet.

### 13. Implement GAM and NGAM

#### 13.1. Growth-associated maintenance (GAM)

GAM is proportional to biomass formation and therefore belongs inside every final biomass reaction. With GAM = 30 mmol ATP gDW^-1, each unit of biomass flux hydrolyses 30 ATP.

30 m_ATP_Cyto[c] + 30 m_WATER_Cyto[c]
-> 30 m_ADP_Cyto[c] + 30 m_Pi_Cyto[c] + 30 m_PROTON_Cyto[c]

#### 13.2. Non-growth-associated maintenance (NGAM)

NGAM is independent of biomass formation and must be represented by a separate ATP hydrolysis reaction with a fixed flux for the publication analyses. The existing model reaction is R_NGAM_ATP_Cyto, and the implemented value is 1.7 mmol ATP gDW^-1 h^-1.

ngamRxn = 'R_NGAM_ATP_Cyto';
assert(ismember(ngamRxn, model.rxns), 'NGAM reaction missing');
model = changeRxnBounds(model, ngamRxn, 1.7, 'l');
model = changeRxnBounds(model, ngamRxn, 1.7, 'u');

### 14. Create condition-specific model variants

Keep one master model containing all valid trophic BOFs. Create separate model variables for AUTO, MIXO and HETERO simulations. Set one objective, close the unused final BOFs, and apply the matching environmental constraints.

```
allBOFs = {
    'TOTAL_BIOMASS_AUTO_MEAN'
    'TOTAL_BIOMASS_MIXO_MEAN'
    'TOTAL_BIOMASS_HETERO_MEAN'
};

modelAuto = model;
modelMixo = model;
modelHeto = model;

modelAuto = changeObjective(modelAuto, 'TOTAL_BIOMASS_AUTO_MEAN');
modelMixo = changeObjective(modelMixo, 'TOTAL_BIOMASS_MIXO_MEAN');
modelHeto = changeObjective(modelHeto, 'TOTAL_BIOMASS_HETERO_MEAN');

modelAuto = changeRxnBounds(modelAuto, ...
    {'TOTAL_BIOMASS_MIXO_MEAN','TOTAL_BIOMASS_HETERO_MEAN'}, 0, 'u');
modelMixo = changeRxnBounds(modelMixo, ...
    {'TOTAL_BIOMASS_AUTO_MEAN','TOTAL_BIOMASS_HETERO_MEAN'}, 0, 'u');
modelHeto = changeRxnBounds(modelHeto, ...
    {'TOTAL_BIOMASS_AUTO_MEAN','TOTAL_BIOMASS_MIXO_MEAN'}, 0, 'u');
```

| **Condition** | **Organic carbon** | **Light** | **Inorganic carbon / gases** |
| --- | --- | --- | --- |
| Autotrophic | Closed unless specifically tested | Open at defined photon uptake | CO2 and O2 according to cultivation |
| Mixotrophic | Open for measured substrate | Open | CO2/O2 according to experiment |
| Heterotrophic | Open for measured glucose/glycerol uptake | Closed | O2 open for aerobic growth; CO2 secretion allowed |

### 15. Validate the integrated BOF before export

#### 15.1. Structural model verification

```
verificationAfter = verifyModel(model);

assert(numel(unique(model.rxns)) == numel(model.rxns), ...
    'Duplicate reaction IDs found.');
assert(numel(unique(model.mets)) == numel(model.mets), ...
    'Duplicate metabolite IDs found.');
assert(all(model.lb <= model.ub), ...
    'At least one lower bound exceeds its upper bound.');

disp('Objective reactions:');
disp(model.rxns(model.c ~= 0));
```

#### 15.2. Test each macromolecular pool

A zero final growth rate can be caused by only one unavailable pool. Test each pool reaction independently under the same environmental constraints before testing the final BOF.

```
testRxns = {
    'PROTEIN_TOTAL_POOL_BIO_AUTOTROPH_NOFRAC'
    'CARBOHYDRATE_TOTAL_POOL_BIO_AUTOTROPH_NOFRAC'
    'LIPID_TOTAL_POOL_BIO_AUTOTROPH_NOFRAC'
    'RNA_TOTAL_POOL_BIO_AUTOTROPH_NOFRAC'
    'DNA_TOTAL_POOL_BIO_AUTOTROPH_NOFRAC'
    'PIGMENT_TOTAL_POOL_BIO_AUTOTROPH_NOFRAC'
};

for i = 1:numel(testRxns)
    testModel = changeObjective(modelAuto, testRxns{i});
    sol = optimizeCbModel(testModel, 'max');
    fprintf('%-60s  %g\n', testRxns{i}, sol.f);
end
```

#### 15.3. Test final growth

solutionAuto = optimizeCbModel(modelAuto, 'max');
solutionMixo = optimizeCbModel(modelMixo, 'max');
solutionHeto = optimizeCbModel(modelHeto, 'max');

fprintf('AUTO growth: %.8g h^-1\n', solutionAuto.f);
fprintf('MIXO growth: %.8g h^-1\n', solutionMixo.f);
fprintf('HETERO growth: %.8g h^-1\n', solutionHeto.f);

#### 15.4. Mass and charge balance

Run the COBRA mass/charge balance check and review genuine biochemical reactions separately from deliberate pool pseudo-reactions. Generic carriers such as ACP, BCCP, tRNA, flavodoxin and protein pools require conserved pseudo-elements or curated formulas. Do not hide an incorrect reaction stoichiometry by inventing an artificial metabolite formula.

```
[massImbalance, chargeImbalance, ...
 imBalancedMass, imBalancedCharge, imBalancedRxnBool] = ...
    checkMassChargeBalance(model);
```

#### 15.5. Energy and cycle checks

Close all external energy sources and test whether ATP maintenance can still carry flux. A positive ATPM flux without substrate, light or electron donor indicates an energy-generating cycle or an incorrectly reversible reaction.

cycleTest = model;
% Close carbon, photons and other energy inputs using the model-specific exchange IDs.
cycleTest = changeObjective(cycleTest, 'R_NGAM_ATP_Cyto');
cycleSol = optimizeCbModel(cycleTest, 'max');
fprintf('ATP without energy input: %g\n', cycleSol.f);

### 16. Troubleshooting guide

| **Symptom** | **Likely cause** | **Corrective action** |
| --- | --- | --- |
| Final biomass flux is zero | One pool cannot be produced; missing transport; wrong compartment; unused BOF closed incorrectly. | Optimize every pool separately; inspect blocked metabolites and transport routes. |
| Growth is much smaller than expected | Fraction applied twice; uptake bound too low; GAM/NGAM too high. | Audit coefficient architecture and compare with measured uptake units. |
| Growth is unrealistically high | Unused carbon sources remain open; NGAM not constrained; energy-generating cycle. | Close all unintended exchanges and test ATP production without substrate. |
| addReaction reports missing metabolites | Reaction string contains display names or outdated IDs. | Use exact model.mets identifiers or add the required metabolite first. |
| Duplicate reaction ID error | Old BOF was not removed. | Use intersect and removeRxns before insertion. |
| SBML reload loses objective | Objective field was not set or export did not preserve FBC objective. | Run changeObjective immediately before export and inspect reloaded model.c. |
| Mass-balance report flags BOF pools | Pseudo-metabolites have no elemental formula. | Separate expected pseudo-reaction warnings from real biochemical imbalances. |
| Mannan reaction is unbalanced | GDP-mannose is consumed without GDP release. | Use a mannan-residue pool or explicit GDP-producing polymerization reaction. |
| Pigment protein is double counted | Whole phycocyanin was included in pigment BOF. | Use phycocyanobilin only; leave apoprotein in the protein pool. |
| MAT and XML models produce different growth | Round-trip altered bounds, objective, IDs or formulas. | Compare reaction count, bounds, objective and growth before and after export. |

### 17. Save MATLAB models and export SBML

Save the master model and each condition-specific variant. Use explicit names containing the trophic state, scenario and version. Do not overwrite the base network.

```
save('Galdieria_TOTAL_BIOMASS_master_vXX.mat', 'model', '-v7.3');
save('Galdieria_AUTO_mean_vXX.mat', 'modelAuto', '-v7.3');
save('Galdieria_MIXO_mean_vXX.mat', 'modelMixo', '-v7.3');
save('Galdieria_HETERO_mean_vXX.mat', 'modelHeto', '-v7.3');

writeCbModel(modelAuto, ...
    'format', 'sbml', ...
    'fileName', 'Galdieria_AUTO_mean_vXX.xml');
writeCbModel(modelMixo, ...
    'format', 'sbml', ...
    'fileName', 'Galdieria_MIXO_mean_vXX.xml');
writeCbModel(modelHeto, ...
    'format', 'sbml', ...
    'fileName', 'Galdieria_HETERO_mean_vXX.xml');
```

### 18. Mandatory SBML round-trip validation

Every exported SBML file must be reloaded with readCbModel. Compare the number of reactions and metabolites, the objective, the biomass flux and the key bounds. Export is not complete until the reloaded model behaves like the MATLAB model.

```
roundTrip = readCbModel('Galdieria_AUTO_mean_vXX.xml', ...
    'fileType', 'SBML');

verifyRoundTrip = verifyModel(roundTrip);
roundTripSol = optimizeCbModel(roundTrip, 'max');

fprintf('Before export: %.10g\n', solutionAuto.f);
fprintf('After export:  %.10g\n', roundTripSol.f);

fprintf('Original reactions: %d\n', numel(modelAuto.rxns));
fprintf('Reloaded reactions: %d\n', numel(roundTrip.rxns));

disp('Reloaded objective:');
disp(roundTrip.rxns(roundTrip.c ~= 0));
```

**Acceptance criterion:** The pre-export and post-reload optimal biomass fluxes should agree within numerical solver tolerance, and the intended trophic-state BOF must remain the only active objective.

### 19. Version control and provenance

| **File** | **Recommended content** |
| --- | --- |
| integrate_TOTAL_BIOMASS_vXX.m | Complete reproducible integration script. |
| BOF_Transfer_vXX.xlsx | Clean list of inserted reactions and source provenance. |
| Galdieria_TOTAL_BIOMASS_master_vXX.mat | Master model with all trophic BOFs. |
| Galdieria_AUTO/MIXO/HETERO_mean_vXX.mat | Condition-specific MATLAB models. |
| Galdieria_AUTO/MIXO/HETERO_mean_vXX.xml | Validated SBML-FBC files. |
| BOF_validation_vXX.txt or .xlsx | Pool-production tests, growth rates, objective, balance results and round-trip comparison. |
| CHANGELOG_vXX.txt | Old reaction IDs removed, new IDs added, coefficient source and unresolved issues. |

### 20. Final acceptance checklist

- Authoritative workbook version and source sheets are recorded.
- One coefficient architecture is used consistently.
- All macromolecular pool reactions required by the final BOF are present.
- Every metabolite ID matches model.mets and the intended compartment.
- Only explicitly obsolete BOF reactions were removed.
- GAM is inside the final BOF and NGAM is a separate ATP hydrolysis reaction.
- Only one trophic-state biomass reaction is the active objective.
- Unused trophic-state BOFs are closed in each condition model.
- Condition-specific exchange and light constraints have been applied.
- Every macromolecular pool can carry positive flux.
- The final BOF carries positive, biologically plausible flux.
- No unintended ATP-generating cycle remains.
- Mass/charge warnings have been classified as real chemistry or deliberate pseudo-reactions.
- MAT files were saved before SBML export.
- Each SBML file was reloaded and produced the same growth result within tolerance.
- The integration script, validation output and change log were archived.

### 21. Recommended implementation decision for the current Galdieria model

For the current project, use the normalized-pool architecture: each macromolecular reaction defines one normalized composition unit, and the final AUTO, MIXO or HETERO BOF applies the macromolecular mass fraction once. Retain all three trophic mean BOFs in the master model, activate one objective per simulation, keep GAM = 30 mmol ATP gDW^-1 in the final BOF, and keep NGAM = 1.7 mmol ATP gDW^-1 h^-1 as a separate constrained reaction. Export and validate one SBML file for each trophic state.

### 22. Key COBRA Toolbox functions used

| **Function** | **Use in this workflow** |
| --- | --- |
| readCbModel | Load MAT or SBML into a COBRA model structure. |
| verifyModel | Check structural consistency of model fields. |
| addMetabolite | Add missing pool or biomass pseudo-metabolites. |
| addReaction | Insert macromolecular and final biomass reactions. |
| removeRxns | Remove reviewed obsolete BOF reactions. |
| changeObjective | Set the active trophic-state BOF. |
| changeRxnBounds | Apply uptake, maintenance and objective bounds. |
| optimizeCbModel | Run FBA and pool-production tests. |
| checkMassChargeBalance | Identify stoichiometric formula and charge inconsistencies. |
| writeCbModel | Export a COBRA model to SBML-FBC. |
| save | Store reproducible MATLAB model files. |

Official software references: COBRA Toolbox documentation (https://opencobra.github.io/cobratoolbox/) and the SBML Level 3 Flux Balance Constraints package specification. Function signatures can vary by COBRA Toolbox version; confirm the installed version using help functionName before execution.
