## Supplementary Data 2 for "Experimental trophic flexibility guides a compartmentalized genome-scale model of red microalgae *Galdieria sulphuraria*": Macro_Biomass.docx

**Calculation of macromolecules fractions for biomass objective function in Galdieria sulphuraria**

### 1. Protein Biomass Fraction

#### 1.1. Overview

The protein fraction of Galdieria sulphuraria biomass is highly condition-dependent, especially under heterotrophic growth, where substrate quality and nitrogen status strongly change composition. The older version that treated heterotrophic protein as narrowly 0.45–0.55 g gDW⁻¹ is too restrictive once the DOI-supported studies (DOI: 10.1016/j.biortech.2021.125477; DOI: 10.1016/j.biortech.2023.129281; DOI: 10.1016/j.scowo.2025.100116) are incorporated. The strongest direct evidence now spans low-protein, carbon-rich fish-residue biomass at 18.9% w/w, intermediate values around 25–42% w/w in technical-scale non-sterile heterotrophic cultures, and a digestate/straw case at 37% w/w. Protein-rich autotrophic and mixotrophic chemostat biomass at 62% w/w remains the best direct value for those two trophic modes.

#### 1.2. Heterotrophic Protein Fraction

##### 1.2.1. Primary direct datasets

The most useful direct heterotrophic protein datasets are:

- DOI: 10.1016/j.biortech.2021.125477: biomass composition in digestate plus straw hydrolysate contained 37% proteins.
- DOI: 10.1016/j.scowo.2025.100116: technical-scale long-term heterotrophic cultivation reported 25.0 ± 1.7%, 37.4 ± 5.8%, and 41.7 ± 4.7% protein depending on substrate system.
- DOI: 10.1016/j.biortech.2023.129281: fed-batch heterotrophic cultivation on hydrolyzed fish residues reported 18.9% w/w protein in the produced biomass.

Taken together, the attached manuscripts show a direct heterotrophic protein interval of:

$$\text{Protein}_{\text{hetero, direct}}=0.189-0.417 \text{g} \text{gDW}^{-1}$$

##### 1.2.2. Secondary direct support

The study identified by DOI: 10.1016/j.biortech.2023.129281 also cites DOI: 10.1039/c2fo30198a and states that G. sulphuraria contained 26.5% protein when heterotrophically grown and 32.5% when autotrophically grown. That is useful secondary support because it sits inside the direct heterotrophic interval above.

##### 1.2.3. Secondary biochemical support

DOI: 10.1016/j.biortech.2024.130628 reported that after glucose depletion, heterotrophic cultures reached 7.8–12.0% nitrogen (w/w). Converting with the standard factor:

$$\text{Protein}\approx N\times6.25$$

gives:

$$\text{Protein}_{hetero,N-based}=0.49-0.75 \text{g} \text{gDW}^{-1}$$

This is best treated as an upper physiological envelope, not the default BOF value, because it is protein-equivalent from nitrogen rather than direct proximate protein.

##### 1.2.4. Heterotrophic conclusion

The literature-backed heterotrophic picture is now much clearer:

- **0.189–0.417 g gDW⁻¹** = direct composition from attached heterotrophic manuscripts.
- 0.265 g gDW⁻¹ = secondary direct support DOI: 10.1039/c2fo30198a as quoted in the source DOI: 10.1016/j.biortech.2023.129281.
- 0.49–0.75 g gDW⁻¹ = nitrogen-based upper envelope after glucose depletion.

For GSM use, the most defensible **heterotrophic BOF working range** is:

$$\boxed{\text{Protein}_{\text{hetero}}=0.25-0.42 \text{g} \text{gDW}^{-1}}$$

This keeps the BOF inside the replicated direct-composition range from DOI: 10.1016/j.biortech.2021.125477, DOI: 10.1016/j.scowo.2025.100116, and DOI: 10.1016/j.biortech.2023.129281, while excluding the fish-residue low outlier (0.189) as a likely nitrogen-limited/specialized case and excluding the N-based 0.49–0.75 envelope as non-default protein-equivalent rather than direct composition.

#### 1.3. Autotrophic Protein Fraction

##### 1.3.1. Primary direct dataset

DOI: 10.1016/j.algal.2021.102603 reported that autotrophic and mixotrophic chemostat cultures had no differences in protein content and that the biomass had 62% w/w protein.

$$\text{Protein}_{\text{auto, primary}}=0.62 \text{g} \text{gDW}^{-1}$$

##### 1.3.2. Secondary support

The study DOI: 10.1016/j.biortech.2023.129281 cites DOI: 10.1039/c2fo30198a as reporting 32.5% protein when autotrophically grown. This is a lower direct comparator and shows that autotrophic protein content can vary strongly with strain and cultivation setup.

##### 1.3.3. Elemental support from the attached DOI: 10.1016/j.algal.2021.102513 manuscript

The attached 2021 Abiusi manuscript reports chemostat autotrophic **N% = 9.8 ± 0.3**. Converting with 6.25:

$$\text{Protein}_{auto,N-based}=9.8\%\times6.25=61.25\%$$

or roughly:

$$\text{Protein}_{auto,N-based}\approx0.61 \text{g} \text{gDW}^{-1}$$

This strongly agrees with the 0.62 direct protein value from DOI: 10.1016/j.algal.2021.102603 and therefore reinforces the high-protein autotrophic estimate.

##### 1.3.4. Autotrophic conclusion

The best current autotrophic value remains:

$$\boxed{\text{Protein}_{\text{auto}}=0.61-0.62 \text{g} \text{gDW}^{-1}}$$

with 0.62 g gDW⁻¹ as the primary BOF value and 0.32–0.33 g gDW⁻¹ as a lower historical literature comparator DOI: 10.1039/c2fo30198a.

#### 1.4. Mixotrophic Protein Fraction

##### 1.4.1. Primary direct dataset

DOI: 10.1016/j.algal.2021.102603 directly states that there were no differences in protein content between autotrophic and mixotrophic chemostat cultures and reports 62% w/w protein.

$$\text{Protein}_{\text{mix, direct}}=0.62 \text{g} \text{gDW}^{-1}$$

##### 1.4.2. Elemental support from the attached DOI: 10.1016/j.algal.2021.102513 manuscript

The attached 2021 Abiusi manuscript reports chemostat **N% = 9.3 ± 0.1** for mixotrophy. Using the same conversion:

$$\text{Protein}_{mix,N-based}=9.3\%\times6.25=58.1\%$$

or:

$$\text{Protein}_{mix,N-based}\approx0.58 \text{g} \text{gDW}^{-1}$$

This is extremely useful because it gives a second, independent mixotrophic estimate derived from the attached manuscript itself.

##### 1.4.3. Recalculation integrating all evidence

Now that heterotrophic direct values are lower than the older 0.45–0.55 assumption, the mixotrophic recalculation should be updated.

Use:

- autotrophic direct value: (0.62)
- heterotrophic working midpoint: ((0.25+0.42)/2 = 0.335)
- partial shift factor (f = 0.10–0.20), because DOI: 10.1016/j.algal.2021.102603 still found no statistically significant protein difference, so only a mild downward correction is justified.

$$\text{Protein}_{\text{mix, }\text{recalc}}=\text{Protein}_{\text{auto}}-f\left( \text{Protein}_{\text{auto}}-\text{Protein}_{\text{hetero, mid}} \right)$$

$$=0.62-\left( 0.10-0.20 \right)\left( 0.62-0.335 \right)$$

$$=0.62-\left( 0.10-0.20 \right)\left( 0.285 \right)$$

$$=0.62-0.0285\text{ to }0.62-0.057$$

$$=0.56-0.59 \text{g} \text{gDW}^{-1}$$

This recalculated interval agrees well with the N-based estimate of ~0.58 from the source DOI: 10.1016/j.algal.2021.102513.

##### 1.4.4. Mixotrophic conclusion

The improved mixotrophic interpretation is:

- 0.62 g gDW⁻¹ = direct measured upper value.
- ~0.58 g gDW⁻¹ = N-based estimate from the source DOI: 10.1016/j.algal.2021.102513.
- **0.56–0.59 g gDW⁻¹** = recalculated GSM-ready value integrating autotrophic direct data plus the now-lower heterotrophic direct range.

For BOF construction, the most defensible mixotrophic range is:

$$\boxed{\text{Protein}_{\text{mix}}=0.56-0.59 \text{g} \text{gDW}^{-1}}$$

#### 1.5. Table 1 – Detailed Protein Evidence

| **Trophic state** | **Protein (g gDW⁻¹)** | **Type** | **Source** |
| --- | --- | --- | --- |
| Heterotrophic | 0.37 | Primary direct | DOI: 10.1016/j.biortech.2021.125477 |
| Heterotrophic | 0.25 ± 0.017 | Primary direct | DOI: 10.1016/j.scowo.2025.100116 defined medium |
| Heterotrophic | 0.374 ± 0.058 | Primary direct | DOI: 10.1016/j.scowo.2025.100116 complex medium I |
| Heterotrophic | 0.417 ± 0.047 | Primary direct | DOI: 10.1016/j.scowo.2025.100116 complex medium II |
| Heterotrophic | 0.189 | Primary direct | DOI: 10.1016/j.biortech.2023.129281 fish residues |
| Heterotrophic | 0.265 | Secondary direct | DOI: 10.1039/c2fo30198a as quoted in source DOI: 10.1016/j.biortech.2023.129281 |
| Heterotrophic | 0.49–0.75 | Secondary N-based | DOI: 10.1016/j.biortech.2024.130628, N×6.25 |
| Autotrophic | 0.62 | Primary direct | DOI: 10.1016/j.algal.2021.102603 |
| Autotrophic | ~0.61 | Secondary N-based | DOI: 10.1016/j.algal.2021.102513, 9.8% N × 6.25 |
| Autotrophic | 0.325 | Secondary direct | DOI: 10.1039/c2fo30198a as quoted in source DOI: 10.1016/j.biortech.2023.129281 |
| Mixotrophic | 0.62 | Primary direct | DOI: 10.1016/j.algal.2021.102603 |
| Mixotrophic | ~0.58 | Secondary N-based | DOI: 10.1016/j.algal.2021.102513, 9.3% N × 6.25 |
| Mixotrophic | 0.56–0.59 | Recalculated | This synthesis |

#### 1.6. Table 2 – Recommended BOF Values

| **Trophic state** | **Protein (g gDW⁻¹)** | **Recommended use** |
| --- | --- | --- |
| Autotrophic | 0.61–0.62 | BOF primary |
| Mixotrophic | 0.56–0.59 | BOF corrected |
| Heterotrophic | 0.25–0.42 | BOF primary range |

#### 1.7. Key Biological Insight

The attached manuscripts change the interpretation substantially. Heterotrophic G. sulphuraria is not best described by a single high-protein value; instead, it spans a wide direct range from roughly 19% to 42% protein, depending on substrate and nutrient regime. By contrast, autotrophic and mixotrophic chemostat biomass remains consistently high-protein at about 62%, with the nitrogen data from DOI: 10.1016/j.algal.2021.102513 independently supporting ~61% for autotrophy and ~58% for mixotrophy.

#### 1.8. Final Citation Statement

The protein fraction of Galdieria sulphuraria biomass is strongly condition-dependent. Direct heterotrophic composition data from DOI-supported studies (DOI: 10.1016/j.biortech.2021.125477; DOI: 10.1016/j.biortech.2023.129281; DOI: 10.1016/j.scowo.2025.100116) span approximately 18.9–41.7% w/w, with DOI: 10.1016/j.biortech.2021.125477 reporting 37%, DOI: 10.1016/j.scowo.2025.100116 reporting 25.0–41.7% depending on substrate, and DOI: 10.1016/j.biortech.2023.129281 reporting 18.9% in fish-residue-fed biomass. Autotrophic and mixotrophic chemostat cultures, however, were reported at 62% protein with no significant difference between those trophic modes, and the nitrogen data from DOI: 10.1016/j.algal.2021.102513 independently support approximately 0.61 g gDW⁻¹ for autotrophy and 0.58 g gDW⁻¹ for mixotrophy. On that basis, the most defensible BOF values are 0.61–0.62 g gDW⁻¹ for autotrophy, 0.56–0.59 g gDW⁻¹ for mixotrophy, and 0.25–0.42 g gDW⁻¹ for heterotrophy.

### 2. Carbohydrate Biomass Fraction

#### 2.1. Overview

The carbohydrate fraction of Galdieria sulphuraria biomass is even more condition-dependent than the protein fraction. The attached heterotrophic manuscripts show that carbohydrate content can swing from 3.9% w/w to 44.2% w/w depending on substrate regime, while an earlier digestate/straw case reported 20% w/w. That means the older single-value heterotrophic assumption is too narrow once the new manuscript evidence is included. The most important direct evidence now comes from DOI: 10.1016/j.biortech.2021.125477, DOI: 10.1016/j.biortech.2023.129281, and DOI: 10.1016/j.scowo.2025.100116.

#### 2.2. Heterotrophic Carbohydrate Fraction

##### 2.2.1. Primary direct datasets

The strongest direct heterotrophic carbohydrate datasets now available are:

- DOI: 10.1016/j.biortech.2021.125477: heterotrophic biomass composition in digestate and hydrolyzed straw contained 20% carbohydrates
- DOI: 10.1016/j.scowo.2025.100116: long-term technical-scale heterotrophic cultivation reported three distinct carbohydrate states depending on substrate system:
  - defined medium: 36.2 ± 7.4%
  - complex medium I: 3.9 ± 0.1%
  - **complex medium II: 13.0 ± 2.3%**

$$0.107-0.153 \text{g} \text{gDW}^{-1}$$

**These values are directly reported in the biomass composition table of the attached manuscript.**
- DOI: 10.1016/j.biortech.2023.129281: fed-batch heterotrophic cultivation on hydrolyzed fish residues produced biomass with 44.2% carbohydrates
  and the discussion explicitly contrasts this with earlier lower-carbohydrate biomass.

Taken together, the direct heterotrophic interval from the attached manuscripts is:

$$\text{Carbohydrate}_{\text{hetero, direct}}=0.039-0.442 \text{g} \text{gDW}^{-1}$$

##### 2.2.2. Secondary direct support

The source DOI: 10.1016/j.biortech.2023.129281 also quotes DOI: 10.1039/c2fo30198a as reporting 69.1% carbohydrate for heterotrophically grown G. sulphuraria. Because this value is cited inside the manuscript discussion rather than being one of the new direct datasets, it is best treated as a secondary upper literature envelope rather than the default BOF value.

$$\text{Carbohydrate}_{\text{hetero, secondary}}=0.691 \text{g} \text{gDW}^{-1}$$

##### 2.2.3. Secondary biochemical support

The same source DOI: 10.1016/j.biortech.2023.129281 states that the carbohydrate fraction in G. sulphuraria is present in the form of glycogen and highlights its relevance as an underused storage polymer. That is important mechanistic support because it explains why carbohydrate can rise strongly under carbon-rich or nitrogen-limited heterotrophic conditions.

##### 2.2.4. Heterotrophic conclusion

The literature-backed heterotrophic picture is now much clearer:

- **0.039–0.442 g gDW⁻¹** = direct composition from attached heterotrophic manuscripts.
- 0.691 g gDW⁻¹ = secondary upper comparator quoted in the source DOI: 10.1016/j.biortech.2023.129281.

For GSM use, the most defensible **heterotrophic BOF working range** is:

$$\boxed{\text{Carbohydrate}_{\text{hetero}}=0.10-0.36 \text{g} \text{gDW}^{-1}}$$

This keeps the BOF inside the repeated direct-composition range from DOI: 10.1016/j.biortech.2021.125477, DOI: 10.1016/j.scowo.2025.100116, and DOI: 10.1016/j.biortech.2023.129281 while excluding:

- the **0.039** value as a highly protein-dominated special case, and
- the **0.442** value as a strong carbon-storage case rather than a default balanced heterotrophic composition.

#### 2.3. Autotrophic Carbohydrate Fraction

##### 2.3.1. Direct evidence available from attached manuscripts

The attached manuscript set does not provide a direct autotrophic carbohydrate mass fraction. What it does provide is strong physiological support that autotrophic and mixotrophic states differ substantially in pigmentation and carbon partitioning. In the DOI: 10.1016/j.algal.2021.102513 manuscript, autotrophic cultures are used as the phototrophic reference state, while mixotrophic growth is described as the combination of autotrophic and heterotrophic metabolism. The same paper states that the average absorption cross section of autotrophic culture was roughly double that of the mixotrophic culture, consistent with greater investment in photosynthetic machinery under autotrophy.

That matters because a stronger photosynthetic apparatus and the protein-rich autotrophic biomass from the protein section imply that autotrophy should sit toward the **lower side of carbohydrate storage**, not near the high heterotrophic storage cases.

##### 2.3.2. Recalculation using manuscript-backed constraints

Using the updated protein section:

$$\text{Protein}_{\text{auto}}=0.61-0.62 \text{g} \text{gDW}^{-1}$$

and using a conservative non-carbohydrate residual fraction for lipids, ash, pigments, nucleotides, and cofactors of:

$$R=0.16-0.22$$

the carbohydrate fraction can be estimated by mass closure:

$$\text{Carbohydrate}_{\text{auto, }\text{recalc}}=1-\left( \text{Protein}_{\text{auto}}+R \right)$$

Substituting:

$$=1-\left( 0.61-0.62+0.16-0.22 \right)$$

gives:

$$\text{Carbohydrate}_{\text{auto, }\text{recalc}}=0.16-0.23 \text{g} \text{gDW}^{-1}$$

This value is also consistent with the idea that autotrophy should remain below the carbohydrate-rich heterotrophic cases reported in the DOI-supported studies (DOI: 10.1016/j.biortech.2021.125477; DOI: 10.1016/j.biortech.2023.129281; DOI: 10.1016/j.scowo.2025.100116).

##### 2.3.3. Autotrophic conclusion

The improved autotrophic value is:

$$\boxed{\text{Carbohydrate}_{\text{auto}}=0.16-0.23 \text{g} \text{gDW}^{-1}}$$

This remains a **derived GSM-ready value**, but it is now constrained by:

- the updated protein fraction,
- manuscript-backed physiology from DOI: 10.1016/j.algal.2021.102513,
- and the direct heterotrophic carbohydrate envelope from the attached DOI-supported studies (DOI: 10.1016/j.biortech.2021.125477; DOI: 10.1016/j.biortech.2023.129281; DOI: 10.1016/j.scowo.2025.100116).

#### 2.4. Mixotrophic Carbohydrate Fraction

##### 2.4.1. Direct physiological evidence from attached manuscripts

The DOI: 10.1016/j.algal.2021.102513 manuscript shows that mixotrophic growth in G. sulphuraria behaves as a combination of heterotrophic and autotrophic metabolisms and that mixotrophic cultures have lower pigmentation than autotrophic cultures. It also reports that mixotrophic biomass productivity was 1.8 times higher than autotrophic productivity and that the culture could run without gas exchange under oxygen-balanced mixotrophy. That is exactly the sort of evidence expected for a state with greater carbon availability than autotrophy, but still not equivalent to strongly storage-driven heterotrophy.

##### 2.4.2. Recalculation integrating all evidence

Now that the heterotrophic carbohydrate range has been updated, the mixotrophic recalculation should also be updated.

Use:

- autotrophic recalculated midpoint:
- heterotrophic working midpoint:

Because DOI: 10.1016/j.algal.2021.102513 supports mixotrophy as an intermediate linear combination of autotrophic and heterotrophic metabolism, apply an interpolation factor:

$$f=0.4-0.8$$

Then:

$$\text{Carbohydrate}_{\text{mix, }\text{recalc}}=C_{\text{auto, mid}}+f\left( C_{\text{hetero, mid}}-C_{\text{auto, mid}} \right)$$

$$=0.195+\left( 0.4-0.8 \right)\left( 0.23-0.195 \right)$$

$$=0.195+\left( 0.4-0.8 \right)\left( 0.035 \right)$$

$$=0.209-0.223 \text{g} \text{gDW}^{-1}$$

To reflect uncertainty and keep the BOF slightly wider than the strict midpoint interpolation, a practical GSM-ready range is:

$$\boxed{\text{Carbohydrate}_{\text{mix}}=0.20-0.25 \text{g} \text{gDW}^{-1}}$$

##### 2.4.3. Mixotrophic conclusion

The improved mixotrophic interpretation is:

- **not directly measured** in the attached manuscripts as a mass fraction,
- but strongly constrained physiologically by the DOI: 10.1016/j.algal.2021.102513 mixotrophy manuscript, and
- should sit slightly above autotrophy and below the main heterotrophic storage cases.

So the most defensible mixotrophic range for BOF construction is:

$$\boxed{\text{Carbohydrate}_{\text{mix}}=0.20-0.25 \text{g} \text{gDW}^{-1}}$$

#### 2.5. Table 1 – Detailed Carbohydrate Evidence

| **Trophic state** | **Carbohydrates (g gDW⁻¹)** | **Type** | **Source** |
| --- | --- | --- | --- |
| Heterotrophic | 0.20 | Primary direct | DOI: 10.1016/j.biortech.2021.125477 |
| Heterotrophic | 0.288–0.436 | Primary direct | DOI: 10.1016/j.scowo.2025.100116 defined medium |
| Heterotrophic | 0.038–0.040 | Primary direct | DOI: 10.1016/j.scowo.2025.100116 complex medium I |
| Heterotrophic | 0.107–0.153 | Primary direct | DOI: 10.1016/j.scowo.2025.100116 complex medium II |
| Heterotrophic | 0.442 | Primary direct | DOI: 10.1016/j.biortech.2023.129281 fish residues |
| Heterotrophic | 0.691 | Secondary direct | DOI: 10.1039/c2fo30198a as quoted in source DOI: 10.1016/j.biortech.2023.129281 |
| Autotrophic | 0.16–0.23 | Recalculated | This synthesis using updated protein + manuscript constraints |
| Mixotrophic | 0.20–0.25 | Recalculated | This synthesis using DOI: 10.1016/j.algal.2021.102513 physiology + updated heterotrophic range |

#### 2.6. Table 2 – Recommended BOF Values

| **Trophic state** | **Carbohydrates (g gDW⁻¹)** | **Recommended use** |
| --- | --- | --- |
| Autotrophic | 0.16–0.23 | BOF corrected |
| Mixotrophic | 0.20–0.25 | BOF corrected |
| Heterotrophic | 0.10–0.36 | BOF primary range |

#### 2.7. Key Biological Insight

The attached manuscripts substantially improve the carbohydrate interpretation. Heterotrophic G. sulphuraria is not well represented by a single carbohydrate value; instead, direct manuscript data span roughly 4% to 44% carbohydrate, depending on substrate and nutrient regime. The fish-residue-fed biomass is a carbohydrate-rich case, while one of the 2025 complex media is strongly protein-dominated and carbohydrate-poor. Mixotrophy, by contrast, is best interpreted as an intermediate physiological state: the attached DOI: 10.1016/j.algal.2021.102513 manuscript shows that mixotrophy combines heterotrophic and autotrophic metabolism and reduces pigment investment relative to autotrophy, which supports a modest upward shift in carbohydrate relative to the autotrophic state rather than a jump into the high-storage heterotrophic regime.

#### 2.8. Final Citation Statement

The carbohydrate fraction of Galdieria sulphuraria biomass is strongly condition-dependent. Direct heterotrophic composition data from DOI-supported studies (DOI: 10.1016/j.biortech.2021.125477; DOI: 10.1016/j.biortech.2023.129281; DOI: 10.1016/j.scowo.2025.100116) span approximately 3.9–44.2% w/w, with DOI: 10.1016/j.biortech.2021.125477 reporting 20%, DOI: 10.1016/j.scowo.2025.100116 reporting 3.9–36.2% depending on substrate, and DOI: 10.1016/j.biortech.2023.129281 reporting 44.2% in fish-residue-fed biomass. The source DOI: 10.1016/j.biortech.2023.129281 also quotes a higher historical heterotrophic comparator of 69.1% carbohydrate DOI: 10.1039/c2fo30198a, indicating that extreme storage-rich states are possible but should not be the default BOF assumption. Because the attached DOI: 10.1016/j.algal.2021.102513 mixotrophy manuscript supports mixotrophy as a linear combination of heterotrophic and autotrophic metabolism, and because the updated protein section constrains autotrophic biomass to a protein-rich state, the most defensible BOF values are 0.16–0.23 g gDW⁻¹ for autotrophy, 0.20–0.25 g gDW⁻¹ for mixotrophy, and 0.10–0.36 g gDW⁻¹ for heterotrophy.

### 3. Pigments Biomass Fraction

#### 3.1. Overview

The uploaded pigment manuscripts support three strong conclusions.

Autotrophy is the pigment-rich reference state. The DOI: 10.1016/j.algal.2021.102513 manuscript states that the characteristic blue-green color of Galdieria sulphuraria is due to phycocyanin and chlorophyll a, and that most strains lose pigmentation in darkness.

Mixotrophy has a substantially smaller pigment pool than autotrophy. In the same manuscript, the autotrophic culture had about **double** the average absorption cross section of the mixotrophic culture, which is a usable whole-pigment scaling proxy.

Heterotrophy suppresses pigments, and glucose suppresses them more strongly than glycerol. DOI: 10.3389/fpls.2022.978246 states that chlorophyll a, zeaxanthin, β-carotene, and phycocyanin are all lower in heterotrophy than in phototrophy; chlorophyll a, zeaxanthin, and β-carotene are about 2-fold higher in phototrophy than in glycerol heterotrophy, but 8- to 14-fold higher than in glucose heterotrophy; phycocyanin is about 20-fold higher than glycerol heterotrophy and 40-fold higher than glucose heterotrophy.

The uploaded files also make the normalization issue explicit:

- **chlorophyll a, zeaxanthin, β-carotene** were normalized on **dry biomass**
- **phycocyanin** was normalized on **soluble protein**, not dry biomass.

So whole phycocyanin should not be used directly as a pigment BOF term.

#### 3.2. Rationale for Retaining the Autotrophic Pigment Reference Set as the BOF Baseline

The available Galdieria sulphuraria pigment studies provide strong trophic-state scaling but do not report one complete autotrophic dry-weight dataset containing chlorophyll a, zeaxanthin, β-carotene, and the phycocyanobilin component required for the BOF.

Abiusi et al. showed that autotrophic cells maintain greater photosynthetic pigment investment than mixotrophic cells, but optical absorption data cannot be converted directly into pigment-specific dry-weight fractions. DOI: 10.1016/j.algal.2021.102513.

Pérez Saura et al. provided dry-weight-normalized data for chlorophyll a, zeaxanthin, and β-carotene and showed that these pigments were approximately twofold lower under glycerol heterotrophy and 8- to 14-fold lower under glucose heterotrophy. Phycocyanin decreased more strongly, by approximately 20-fold and 40-fold, respectively. However, phycocyanin was reported as a phycobiliprotein and not as a directly compatible phycocyanobilin dry-weight coefficient. DOI: 10.3389/fpls.2022.978246.

Therefore, the previous autotrophic pigment set was retained as a literature-constrained modelling baseline:

| **Pigment measurement** | **Reference range, g gDW⁻¹** |
| --- | --- |
| Phycocyanin holoprotein | 0.050–0.080 |
| Chlorophyll a | 0.0070–0.0095 |
| Zeaxanthin | 0.00050–0.00080 |
| β-carotene | 0.00030–0.00060 |

Chlorophyll a, zeaxanthin, and β-carotene are small-molecule pigments and can be inserted directly into the pigment BOF after conversion to mmol gDW⁻¹. Phycocyanin must be treated differently because it is a protein–chromophore complex. Its apoprotein mass is already included in the protein BOF, so adding whole phycocyanin again would double-count protein biomass.

Only the phycocyanobilin chromophore should therefore be represented in the pigment BOF. Assuming three phycocyanobilin molecules per phycocyanin αβ monomer:

fPCB = (3 × 587) / 38,000 = 0.0463421 (4.634%)

Thus, the implemented central value represents 4.634% of phycocyanin holoprotein mass. The 3–5% interval is retained only as a sensitivity range.

0.050 × 0.046 = 0.0023 g gDW⁻¹

0.080 × 0.046 = 0.0037 g gDW⁻¹

Therefore:

0.0023–0.0037 g phycocyanobilin gDW⁻¹

The structural calculation supports the broader 3–5% sensitivity envelope, but every central workbook uses the single 4.634% factor.

The final autotrophic pigment set is therefore a hybrid evidence baseline: directly measured small-molecule pigments, experimentally supported trophic-state scaling, and a chromophore-only conversion for phycocyanin to avoid double-counting protein.

#### 3.3. Implemented Phycocyanin-to-Phycocyanobilin Conversion

Whole C-phycocyanin was excluded from the pigment BOF because it is a phycobiliprotein consisting of α- and β-apoprotein subunits with covalently bound phycocyanobilin chromophores. The apoprotein mass is already represented in the protein fraction of the BOF; inclusion of the complete holoprotein in the pigment fraction would therefore double-count its amino-acid contribution.

The BOF contains only the phycocyanobilin chromophore. Assuming three phycocyanobilin molecules per phycocyanin αβ monomer, a phycocyanobilin molecular mass of approximately 587 g mol⁻¹ and an αβ holoprotein mass of approximately 38,000 g mol⁻¹, the implemented conversion factor was:

$$\text{fPCB = (3 × 587) / 38,000 = 0.0463421}$$

Thus, approximately 4.63% of the phycocyanin holoprotein mass was assigned to phycocyanobilin. The wider 3–5% interval was retained only for sensitivity analysis. Pigment identities and trophic-state repression were constrained using Pérez Saura et al. (2022; DOI: 10.3389/fpls.2022.978246), autotrophic versus mixotrophic pigment investment was supported by Abiusi et al. (2021; DOI: 10.1016/j.algal.2021.102513), and the variability of Galdieria phycocyanin content was supported by Rahman et al. (2020; DOI: 10.1007/s10811-019-01957-9).

#### 3.4. Reconciled Autotrophic Pigment Set

The autotrophic source profile contained 0.650 mg zeaxanthin, 8.250 mg chlorophyll a, 0.450 mg β-carotene and 3.186020 mg phycocyanobilin gDW⁻¹. The source-profile sum was 12.536020 mg gDW⁻¹. Because the selected autotrophic total-pigment fraction was 0.0121 g gDW⁻¹, equivalent to 12.1 mg gDW⁻¹, the individual components were normalized to their relative proportions and rescaled to this total. Autotrophy was retained as the pigment-rich BOF reference state.

#### 3.5. Reconciled Mixotrophic Pigment Set

The mixotrophic source profile contained 0.350 mg zeaxanthin, 4.450 mg chlorophyll a, 0.240 mg β-carotene and 1.714658 mg phycocyanobilin gDW⁻¹. The unscaled sum was 6.754658 mg gDW⁻¹. The profile was normalized and rescaled to the selected mixotrophic pigment total of 0.0065 g gDW⁻¹, equivalent to 6.5 mg gDW⁻¹. This preserves the literature-supported reduction in pigment investment relative to autotrophic growth while maintaining an internally closed BOF pigment pool.

#### 3.6. Reconciled Heterotrophic Pigment Sets

Glucose and glycerol heterotrophy were retained as separate states because they produce substantially different pigment repression. Pérez Saura et al. reported that chlorophyll a, zeaxanthin and β-carotene were approximately twofold lower under glycerol heterotrophy than under phototrophy, but approximately 8- to 14-fold lower under glucose heterotrophy. Phycocyanin was reduced even more strongly, by approximately 20-fold under glycerol and 40-fold under glucose heterotrophy. Both profiles used the same phycocyanin-to-phycocyanobilin conversion and metabolite-compartment mapping.

##### 3.6.1. Glycerol heterotrophy

The glycerol source profile contained 0.325 mg zeaxanthin, 2.650 mg chlorophyll a, 0.161 mg β-carotene and 0.137868 mg phycocyanobilin gDW⁻¹. Its unscaled sum was 3.273868 mg gDW⁻¹. The relative composition was normalized and rescaled to the selected glycerol-heterotrophic total of 0.00485 g gDW⁻¹. This scaling retains the measured pigment hierarchy while reconciling it with the selected total pigment fraction used in the macromolecular biomass composition.

##### 3.6.2. Glucose heterotrophy

The glucose profile contained 0.036 mg zeaxanthin, 0.500 mg chlorophyll a, 0.021 mg β-carotene and 0.023171 mg phycocyanobilin gDW⁻¹. These components sum directly to 0.580171 mg gDW⁻¹, corresponding to:

$$\text{0.000580171 g gDW⁻¹}$$

No additional rescaling was required because the component sum and the reconciled total were identical. This profile represents the strongest pigment-repressed trophic state and was retained separately from glycerol heterotrophy rather than combining both carbon sources into one heterotrophic BOF.

#### 3.7. Table 1 – Detailed Pigment Evidence

| **Trophic state** | **Pigment evidence** | **Type** | **Source** |
| --- | --- | --- | --- |
| Autotrophic | phycocyanin + chlorophyll a define pigment-rich state | Primary physiology | DOI: 10.1016/j.algal.2021.102513 |
| Mixotrophic | autotrophic absorption cross section about double mixotrophic | Primary scaling | DOI: 10.1016/j.algal.2021.102513 |
| Heterotrophic | all pigments lower than phototrophy | Primary | DOI: 10.3389/fpls.2022.978246 |
| Heterotrophic | glucose more repressive than glycerol | Primary | DOI: 10.3389/fpls.2022.978246 |
| Heterotrophic | chlorophyll a, zeaxanthin, β-carotene normalized on dry biomass | Primary methods | DOI: 10.3389/fpls.2022.978246 |
| Heterotrophic | phycocyanin normalized on soluble proteins | Primary methods | DOI: 10.3389/fpls.2022.978246 |
| Heterotrophic vs autotrophic | reduced chlorophyll-related phytol in heterotrophy | Supporting | DOI: 10.3390/md18030169 |

#### 3.8. Table 2 – Recommended BOF Values

| Trophic state | Phycocyanobilin | Chlorophyll a | Zeaxanthin | β-carotene | Total pigments |
| --- | --- | --- | --- | --- | --- |
| Autotrophic | 0.003075206 | 0.007963054 | 0.000627392 | 0.000434348 | 0.012100000 |
| Mixotrophic | 0.001650013 | 0.004282230 | 0.000336805 | 0.000230952 | 0.006500000 |
| Heterotrophic (glycerol) | 0.000204241 | 0.003925785 | 0.000481464 | 0.000238510 | 0.004850000 |
| Heterotrophic (glucose) | 0.000023171 | 0.000500000 | 0.000036000 | 0.000021000 | 0.000580171 |

Implemented central values are in g gDW⁻¹ and sum to the selected condition-specific pigment total.

3.9. Table 3 – Recommended BOF values in % / pigment

| Trophic state | Phycocyanobilin | Chlorophyll a | Zeaxanthin | β-carotene |
| --- | --- | --- | --- | --- |
| Autotrophic | 25.41% | 65.81% | 5.19% | 3.59% |
| Mixotrophic | 25.38% | 65.88% | 5.18% | 3.55% |
| Heterotrophic (glycerol) | 4.21% | 80.94% | 9.93% | 4.92% |
| Heterotrophic (glucose) | 3.99% | 86.18% | 6.21% | 3.62% |

#### 3.10. Key Biological Insight

The reconciled module separates experimental trophic scaling from model-selected totals and uses one central PCB conversion across every workbook and table.

Autotrophy remains the pigment-rich anchor; mixotrophy is scaled downward; and glycerol and glucose heterotrophy remain separate.

All four conditions now use identical formulas, metabolite identifiers, compartments and metadata.

The reported component percentages are calculated from the same normalized profiles used to calculate BOF coefficients.

Only phycocyanobilin is included in the pigment BOF, avoiding protein double counting.

The implemented central total pigment fractions are:

- autotrophic: 0.0121 g gDW⁻¹
- mixotrophic: 0.0065 g gDW⁻¹
- heterotrophic glycerol: 0.00485 g gDW⁻¹; heterotrophic glucose: 0.000580171 g gDW⁻¹

the best direct cross-trophic total-lipid dataset is now DOI: 10.1038/s41598-019-46645-3, not the earlier inferred auto/mixo values. The pigment PDFs help with trophic physiology, but the lipid PDFs are what should drive the final BOF lipid fractions.

### 4. Lipids Biomass Fraction

#### 4.1. Overview

The lipid fraction of Galdieria sulphuraria is lower and less variable than protein or carbohydrate, but it is still trophic-state and substrate dependent. The strongest direct evidence now comes from:

- DOI: 10.1016/j.biortech.2021.125477, DOI: 10.1016/j.biortech.2023.129281, and DOI: 10.1016/j.scowo.2025.100116 for heterotrophic biomass composition, and
- DOI: 10.1038/s41598-019-46645-3 for a matched autotrophic / mixotrophic / heterotrophic total-lipid comparison in one study.

The key conclusion is that Galdieria is **not strongly oleaginous under the BOF-relevant conditions**, but autotrophy can still have a distinctly higher total-lipid fraction than glucose heterotrophy.

#### 4.2. Heterotrophic Lipids Fraction

##### 4.2.1. Primary direct total-lipid datasets

The strongest direct heterotrophic **total-lipid** datasets are:

- DOI: 10.1016/j.biortech.2021.125477:
  heterotrophic biomass in digestate + hydrolyzed straw contained 3% lipids
- DOI: 10.1016/j.scowo.2025.100116:
  long-term technical-scale heterotrophic cultivation reported:
  - defined medium: 2.4 ± 0.9%
  - complex medium I: 5.3 ± 0.6%
  - complex medium II: 3.6 ± 0.4%
- DOI: 10.1016/j.biortech.2023.129281:
  fish-residue-fed heterotrophic biomass contained 0.9% lipids
- DOI: 10.1038/s41598-019-46645-3:
  heterotrophic Galdieria sp. USBA-GBX-832 contained 3.64 ± 0.25% total lipids

Taken together, the direct heterotrophic total-lipid interval is:

$$\text{Lipid}_{\text{hetero, direct}}=0.009-0.059 \text{g} \text{gDW}^{-1}$$

##### 4.2.2. Secondary support

The attached 2023 fish-residue manuscript cites DOI: 10.1039/c2fo30198a as reporting 11.4% lipid for heterotrophically grown G. sulphuraria. That is a useful upper historical comparator, but it is not consistent with the cluster of newer direct measurements.

DOI: 10.3390/md18030169 also supports a low-lipid phenotype overall, showing major trophic changes in fatty acids and FTIR lipid-related signal, but not a strong case for high total lipids as the default state.

##### 4.2.3. Heterotrophic conclusion

The manuscript-backed heterotrophic picture is:

- **0.009–0.059 g gDW⁻¹** = direct total-lipid measurements.
- 0.114 g gDW⁻¹ = historical upper comparator, not the default BOF value.

For BOF use, the most defensible heterotrophic working range is:

$$\boxed{\text{Lipid}_{\text{hetero}}=0.020-0.055 \text{g} \text{gDW}^{-1}}$$

This keeps the BOF inside the repeated direct total-lipid range while excluding the **0.009** fish-residue low extreme and the **0.114** historical high envelope.

#### 4.3. Autotrophic Lipids Fraction

##### 4.3.1. Primary direct total-lipid evidence

The most important direct autotrophic lipid result is now DOI: 10.1038/s41598-019-46645-3:

$$15.34\pm3.3\%=0.153\pm0.033 \text{g} \text{gDW}^{-1}$$

for autotrophic biomass. In the same study, heterotrophy and mixotrophy were both only about **3.6–3.9% total lipids**, so this is the strongest direct evidence that autotrophy can support a distinctly higher total-lipid fraction in a Galdieria strain.

##### 4.3.2. Secondary support

DOI: 10.3390/md18030169 points to strong trophic remodeling of lipid-related chemistry between autotrophic and heterotrophic conditions, even if its lipid interpretation is more compositional than a clean BOF-ready total-lipid percentage.

##### 4.3.3. Autotrophic conclusion

Because López gives a direct autotrophic total-lipid value and it is much stronger than the previous inferred range, the old **0.03–0.05** autotrophic BOF interval should be replaced.

A defensible BOF range is:

$$\boxed{\text{Lipid}_{\text{auto}}=0.12-0.18 \text{g} \text{gDW}^{-1}}$$

This keeps the direct value from DOI: 10.1038/s41598-019-46645-3 near the center while allowing normal experimental uncertainty and strain dependence.

#### 4.4. Mixotrophic Lipids Fraction

##### 4.4.1. Primary direct total-lipid evidence

DOI: 10.1038/s41598-019-46645-3 also gives the best direct mixotrophic value:

$$3.85\pm0.13\%=0.0385\pm0.0013 \text{g} \text{gDW}^{-1}$$

which is very close to the heterotrophic value in the same study.

##### 4.4.2. Physiological support

The attached DOI: 10.1016/j.algal.2021.102513 mixotrophy manuscript supports mixotrophy as a combined autotrophic + heterotrophic state with reduced pigmentation relative to autotrophy, but it does not justify moving mixotrophic total lipids up anywhere near the autotrophic value from DOI: 10.1038/s41598-019-46645-3.

##### 4.4.3. Mixotrophic conclusion

Because there is a direct mixotrophic total-lipid value, mixotrophy no longer needs to be inferred only from interpolation.

A practical BOF range is:

$$\boxed{\text{Lipid}_{\text{mix}}=0.035-0.050 \text{g} \text{gDW}^{-1}}$$

This keeps the direct López result at the center while allowing modest condition-to-condition flexibility.

#### 4.5. Table 1 – Detailed Lipid Evidence

| **Trophic state** | **Lipids (g gDW⁻¹)** | **Type** | **Source** |
| --- | --- | --- | --- |
| Heterotrophic | 0.03 | Primary direct | DOI: 10.1016/j.biortech.2021.125477 |
| Heterotrophic | 0.015–0.033 | Primary direct | DOI: 10.1016/j.scowo.2025.100116 defined medium |
| Heterotrophic | 0.047–0.059 | Primary direct | DOI: 10.1016/j.scowo.2025.100116 complex medium I |
| Heterotrophic | 0.032–0.040 | Primary direct | DOI: 10.1016/j.scowo.2025.100116 complex medium II |
| Heterotrophic | 0.009 | Primary direct | DOI: 10.1016/j.biortech.2023.129281 fish residues |
| Heterotrophic | 0.0364 ± 0.0025 | Primary direct | DOI: 10.1038/s41598-019-46645-3 |
| Heterotrophic | 0.114 | Secondary direct | DOI: 10.1039/c2fo30198a as cited in attached manuscript |
| Autotrophic | 0.153 ± 0.033 | Primary direct | DOI: 10.1038/s41598-019-46645-3 |
| Mixotrophic | 0.0385 ± 0.0013 | Primary direct | DOI: 10.1038/s41598-019-46645-3 |

#### 4.6. Table 2 – Recommended BOF Values

| **Trophic state** | **Lipids (g gDW⁻¹)** | **Recommended use** |
| --- | --- | --- |
| Autotrophic | 0.12–0.18 | BOF corrected |
| Mixotrophic | 0.035–0.050 | BOF corrected |
| Heterotrophic | 0.020–0.055 | BOF primary range |

#### 4.7. Key Biological Insight

The updated lipid interpretation is now much stronger because it uses a single direct cross-trophic study.

- **Autotrophy** can support substantially higher total lipids than glucose heterotrophy in at least one Galdieria strain/system.
- **Mixotrophy and heterotrophy** cluster much closer together for total lipids in that same study.
- Across the DOI-supported studies (DOI: 10.1016/j.biortech.2021.125477; DOI: 10.1016/j.biortech.2023.129281; DOI: 10.1016/j.scowo.2025.100116), heterotrophic biomass remains mostly in the 1–5% lipid regime.

So the biggest update is:

**autotrophic lipids should be revised upward**, while **mixotrophic and heterotrophic total lipids remain low-to-moderate**.

#### 4.8. Final Citation Statement

The lipid fraction of Galdieria sulphuraria biomass is lower and less variable than the protein and carbohydrate fractions under most heterotrophic conditions, but direct cross-trophic evidence shows that autotrophy can sustain a substantially higher total-lipid fraction. The DOI-supported studies (DOI: 10.1016/j.biortech.2021.125477; DOI: 10.1016/j.biortech.2023.129281; DOI: 10.1016/j.scowo.2025.100116) report heterotrophic total lipids of approximately 0.9–5.3% w/w, with DOI: 10.1016/j.biortech.2021.125477 reporting 3%, DOI: 10.1016/j.scowo.2025.100116 reporting 2.4–5.3% depending on substrate, and DOI: 10.1016/j.biortech.2023.129281 reporting 0.9% in fish-residue-fed biomass. By contrast, DOI: 10.1038/s41598-019-46645-3 reports a matched trophic-state comparison with 15.34 ± 3.3% total lipids in autotrophy, 3.64 ± 0.25% in heterotrophy, and 3.85 ± 0.13% in mixotrophy. On that basis, the most defensible BOF values are 0.12–0.18 g gDW⁻¹ for autotrophy, 0.035–0.050 g gDW⁻¹ for mixotrophy, and 0.020–0.055 g gDW⁻¹ for heterotrophy.

Explicit values:

- Autotroph → 0.15
- Mixotroph → 0.0425
- Heterotroph → **0.0375**

### 5. Nucleotide Biomass Fraction (RNA and DNA, w/w biomass)

The nucleotide biomass fraction represents the contributions of RNA (RNS) and DNA (DNS) to total Galdieria sulphuraria biomass. Organism-specific precursor composition was derived from genome sequence data, whereas the RNA and DNA mass fractions were anchored to measured microalgal macromolecular composition.

#### 5.1. Genome sequence source and nucleotide counting

The reference genome assembly GCF_000341285.1 was used for sequence-based composition. Its publication is identified by DOI: 10.1126/science.1231707. The genomic FASTA file was used for DNA composition, and annotated coding-sequence FASTA files were used as a transcript-composition proxy for RNA. FASTA headers were excluded, A/C/G/T occurrences were counted case-insensitively, and ambiguous bases were ignored.

*fᵢ = nᵢ / (nA + nC + nG + nT)*

#### 5.2. Genome-derived precursor fractions

| **Polymer** | **Precursor** | **Mole fraction** |
| --- | --- | --- |
| DNA | dATP | 0.31220 |
| DNA | dTTP | 0.31081 |
| DNA | dCTP | 0.18970 |
| DNA | dGTP | 0.18728 |
| RNA | ATP | 0.30424 |
| RNA | UTP | 0.29894 |
| RNA | CTP | 0.17685 |
| RNA | GTP | 0.21997 |

#### 5.3. RNA and DNA mass fractions

Direct measurements of RNA and DNA mass fractions are not available specifically for G. sulphuraria. The BOF therefore uses the microalgal composition values RNA = 0.057 g gDW⁻¹ and DNA = 0.010 g gDW⁻¹ (DOI: 10.1371/journal.pone.0155977). These are selected point estimates, not measured G. sulphuraria-specific intervals; together they contribute 0.067 g gDW⁻¹ (6.7% of biomass).

| **Component** | **Fraction (g gDW⁻¹)** | **Biomass (%)** |
| --- | --- | --- |
| RNA (RNS) | 0.057 | 5.7 |
| DNA (DNS) | 0.010 | 1.0 |
| Total nucleic acids | 0.067 | 6.7 |

#### 5.4. BOF precursor reactions

0.31220 dATP + 0.31081 dTTP + 0.18970 dCTP + 0.18728 dGTP → DNA

0.30424 ATP + 0.29894 UTP + 0.17685 CTP + 0.21997 GTP → RNA

The precursor reactions describe nucleotide mole fractions within each polymer. Their contributions to one gram dry weight are obtained by applying the separate DNA and RNA biomass fractions. The same selected RNA and DNA mass fractions are used in the autotrophic, mixotrophic and heterotrophic BOFs because trophic-state-specific experimental values are unavailable.

### 6. Growth-Associated (GAM) and Non-Growth-Associated Maintenance (NGAM) Energy Costs

#### 6.1. GAM and NGAM represent different energy terms

Growth-associated maintenance (GAM) is ATP hydrolysis coupled to formation of one gram of biomass. Non-growth-associated maintenance (NGAM) is a time-dependent ATP demand that persists independently of biomass production. They therefore have different units and must be implemented separately.

*qATP,maintenance = GAM × μ + NGAM*

| **Term** | **Meaning** | **Units** | **Model implementation** |
| --- | --- | --- | --- |
| GAM | Energy coupled to biomass synthesis and assembly | mmol ATP gDW⁻¹ | ATP hydrolysis in, or coupled to, the BOF |
| NGAM | Homeostasis, ion gradients, repair and turnover independent of growth | mmol ATP gDW⁻¹ h⁻¹ | Lower bound on a separate ATP-maintenance reaction |

#### 6.2. Growth-associated maintenance

All three trophic-state GAM workbooks use the same selected coefficient because no condition-specific G. sulphuraria GAM measurements are available. The implemented central value is GAM = 30 mmol ATP gDW⁻¹. This is a modelling assumption, not a direct experimental measurement.

*30 ATP + 30 H₂O → 30 ADP + 30 Pᵢ + 30 H⁺*

#### 6.3. Non-growth-associated maintenance

The selected working constraint is NGAM = 1.7 mmol ATP gDW⁻¹ h⁻¹. NGAM is not part of the BOF coefficient and must instead be imposed as a fixed flux through a separate cytosolic ATP-maintenance reaction. The GAM workbooks encode GAM_BIOMASS but do not encode this NGAM lower bound.

*ATP + H₂O → ADP + Pᵢ + H⁺; lower and upper bounds = 1.7 mmol gDW⁻¹ h⁻¹*

#### 6.4. Evidence boundary and sensitivity

An approximate stationary-phase calculation gives an apparent maintenance demand of about 1.7 mmol ATP gDW⁻¹ h⁻¹, but this remains NGAM-like support rather than a pure NGAM measurement because residual growth, storage turnover, stress metabolism and ATP yield were not independently resolved. A chemostat-derived algal benchmark reports 109 mmol ATP g⁻¹ for biomass formation and 2.85 mmol ATP g⁻¹ h⁻¹ for maintenance (DOI: 10.1007/s10811-011-9674-3); this validates the estimation approach but does not directly determine the G. sulphuraria parameter. Sensitivity analysis should therefore include NGAM = 0, 1.7 and 2.85 mmol ATP gDW⁻¹ h⁻¹.

The distinction between biomass composition, GAM and NGAM follows established BOF-construction practice (DOI: 10.1371/journal.pcbi.1006971; DOI: 10.1016/j.mib.2010.03.003).

### 7. Specific Growth Rate (μ) Across Trophic States

The compiled literature dataset contained 135 condition-specific Galdieria sulphuraria growth-rate records: 45 autotrophic, 10 mixotrophic and 80 heterotrophic observations. Across the complete dataset, μ ranged from 0.020 to 1.530 day⁻¹. Because the source studies differ in strain, substrate, pH, temperature, cultivation system and evidence type, these values describe the observed literature coverage rather than a single pooled biological growth rate.

#### 7.1. Conversion Between day⁻¹ and h⁻¹

Growth rates reported per day were converted to the hourly scale used in many constraint-based simulations by dividing by 24 h day⁻¹:

*μ (h⁻¹) = μ (day⁻¹) / 24*

For example, an autotrophic mean of 0.283 day⁻¹ corresponds to 0.283 / 24 = 0.01179 h⁻¹. The same conversion was applied to every range boundary, mean and median shown below.

#### 7.2. Table 1 – Specific Growth Rate by Trophic State

| **Trophic state** | **n** | **Range, day⁻¹** | **Mean, day⁻¹** | **Median, day⁻¹** | **Range, h⁻¹** | **Mean, h⁻¹** | **Median, h⁻¹** |
| --- | --- | --- | --- | --- | --- | --- | --- |
| Autotrophic | 45 | 0.128–0.500 | 0.283 | 0.277 | 0.00533–0.02083 | 0.01179 | 0.01154 |
| Mixotrophic | 10 | 0.228–1.490 | 0.849 | 0.955 | 0.00950–0.06208 | 0.03538 | 0.03979 |
| Heterotrophic | 80 | 0.020–1.530 | 0.632 | 0.610 | 0.00083–0.06375 | 0.02633 | 0.02542 |

Autotrophic growth showed the narrowest range and the lowest descriptive central values. Mixotrophic cultivation had the highest mean and median, although it was represented by only ten records. The heterotrophic dataset was the largest and most diverse and contained the highest observed μ. Accordingly, organic-carbon-supported growth generally permitted higher rates than autotrophic growth, but the unequal dataset sizes and different experimental conditions preclude interpreting these descriptive values as a statistically balanced comparison of trophic-state effects.

### 8. DOI index for automatic citation

10.1016/j.biortech.2021.125477

10.1016/j.biortech.2023.129281

10.1016/j.scowo.2025.100116

10.1016/j.algal.2021.102513

10.1016/j.algal.2021.102603

10.1016/j.biortech.2024.130628

10.1039/c2fo30198a

10.3389/fpls.2022.978246

10.3390/md18030169

10.1038/s41598-019-46645-3

10.1371/journal.pone.0155977

10.1126/science.1231707

10.1007/s10811-011-9674-3

10.1371/journal.pcbi.1006971

10.1016/j.mib.2010.03.003
