## Supplementary Data 3 for "Experimental trophic flexibility guides a compartmentalized genome-scale model of red microalgae *Galdieria sulphuraria*": S1_Galdieria_publication_growth_validation.m

```
function S1_Galdieria_publication_growth_validation()
% Galdieria sulphuraria growth-validation analysis.
% Inputs are loaded automatically from the shared Inputs folder.
% Output: 1_Galdieria_publication_growth_validation.xlsx.
scriptDir = fileparts(mfilename('fullpath'));
if isempty(scriptDir), scriptDir = pwd; end
supplementDir = fileparts(fileparts(scriptDir));
inputDir = fullfile(supplementDir,'Inputs');

cfg.modelFile = fullfile(inputDir,'Galdieria_sulphuraria_SBML_Final.xml');
cfg.substrateFile = fullfile(inputDir,'Model_Substrate_list.xlsx');
cfg.biomassFile = fullfile(inputDir,'TOTAL_BIOMASS.xlsx');
cfg.substrateSheet = 'Sole_substrate_ScrumPy';
cfg.resultsFile = fullfile(scriptDir,'1_Galdieria_publication_growth_validation.xlsx');

cfg.scriptName = '1_Galdieria_publication_growth_validation';

cfg.NGAM = 1.7;
cfg.GAM = 30;

cfg.screenCarbon = 6;              % C-mmol gDW^-1 h^-1
cfg.validationMaxCarbon = 120;     % non-binding opening cap

cfg.heteroPhotonCalibrationMu = 0.06;
cfg.expectedPhotonPerPigment = 0.58;
cfg.photonCalibrationTolerance = 0.02;

% Shared trophic-state constraints used by Scripts 1-4.
cfg.o2MaxUptake = 1000;
cfg.heteroPhotonMaxUptake = 0.2;
cfg.mixoPhotonMaxUptake = 50;
cfg.mixoCO2MaxUptake = cfg.validationMaxCarbon;
cfg.rubiscoMinFluxMIXO = 0.01;
cfg.basalMediumLB = -1000;
cfg.basalMediumUB = 1000;
cfg.o2DummyLB = 0;
cfg.o2DummyUB = 1000;

cfg.MW.glucose = 180.156;
cfg.MW.xylose  = 150.130;


cfg.exp.MIXO.protein       = 0.4147;
cfg.exp.MIXO.carbohydrate  = 0.3377;
cfg.exp.MIXO.nucleic       = 0.0775;
cfg.exp.MIXO.lipid         = 0.0701;

cfg.exp.HETERO.protein      = 0.4016;
cfg.exp.HETERO.carbohydrate = 0.3609;
cfg.exp.HETERO.nucleic      = 0.0765;
cfg.exp.HETERO.lipid        = 0.0610;

cfg.reference.MIXO.rnaFraction = ...
    0.06222707/(0.06222707+0.01091703);

cfg.reference.HETERO.rnaFraction = ...
    0.085/(0.085+0.015);

cfg.reference.HETERO.pigmentMass = 0.00485;

cfg.reference.MIXO.PCB_mg  = 1.714658;
cfg.reference.MIXO.CHLA_mg = 4.450000;
cfg.reference.MIXO.BETA_mg = 0.240000;
cfg.reference.MIXO.ZEA_mg  = 0.350000;

cfg.reference.MIXO.pigmentMass = ...
    (cfg.reference.MIXO.PCB_mg + ...
     cfg.reference.MIXO.CHLA_mg + ...
     cfg.reference.MIXO.BETA_mg + ...
     cfg.reference.MIXO.ZEA_mg)/1000;

if exist('initCobraToolbox','file') ~= 2
    error(['COBRA Toolbox is not on the MATLAB path. Install/initialize ' ...
           'COBRA Toolbox before running this supplementary validation.']);
end

if ~isfile(cfg.modelFile)
    error('Required model file not found: %s',cfg.modelFile);
end

if ~isfile(cfg.substrateFile)
    error('Required substrate workbook not found: %s',cfg.substrateFile);
end

if ~isfile(cfg.biomassFile)
    error('Required biomass workbook not found: %s',cfg.biomassFile);
end

quietWarningState = warning;
warning('off','all');
try
    evalc('initCobraToolbox(false);');
catch ME
    warning(quietWarningState);
    rethrow(ME);
end
warning(quietWarningState);

quietWarningState = warning;
warning('off','all');
try
    evalc('baseModel = readCbModel(cfg.modelFile);');
catch ME
    warning(quietWarningState);
    rethrow(ME);
end
warning(quietWarningState);

cfg = mapModel_pub(baseModel,cfg);

% Match Script 2 exactly: close every pre-existing biomass output before
% constructing the publication experimental BOFs on one shared base model.
[baseModel,~] = closeOriginalBiomassOutputs_pub_s1(baseModel);

[baseModel,pigHET] = addPigmentPool_pub(baseModel,'HETERO',cfg);
[baseModel,pigMIX] = addPigmentPool_pub(baseModel,'MIXO',cfg);

cfg.pigment.HETERO = pigHET;
cfg.pigment.MIXO = pigMIX;

% Exact-core metabolite resolution is required here; substring matching can
% mis-map ADP_CYTO to DADP/3-5-ADP and change growth predictions.
[baseModel,bofHET] = createExperimentalBOF_pub( ...
    baseModel,'HETERO',cfg.exp.HETERO, ...
    cfg.reference.HETERO.rnaFraction, ...
    cfg.reference.HETERO.pigmentMass, ...
    cfg.pigment.HETERO.poolMet,cfg);

[baseModel,bofMIX] = createExperimentalBOF_pub( ...
    baseModel,'MIXO',cfg.exp.MIXO, ...
    cfg.reference.MIXO.rnaFraction, ...
    cfg.reference.MIXO.pigmentMass, ...
    cfg.pigment.MIXO.poolMet,cfg);

cfg.HETERO.biomassRxn = bofHET.sinkRxn;
cfg.HETERO.sinkRxn = bofHET.sinkRxn;
cfg.HETERO.assemblyRxn = bofHET.assemblyRxn;
cfg.HETERO.pigmentRxn = cfg.pigment.HETERO.rxn;

cfg.MIXO.biomassRxn = bofMIX.sinkRxn;
cfg.MIXO.sinkRxn = bofMIX.sinkRxn;
cfg.MIXO.assemblyRxn = bofMIX.assemblyRxn;
cfg.MIXO.pigmentRxn = cfg.pigment.MIXO.rxn;

% Use exactly the same HETERO/MIXO medium, biomass-output isolation and
% pigment-producer isolation as Script 2.
modelHETnative = prepareSharedCondition_pub(baseModel,'HETERO',cfg);
modelMIX = prepareSharedCondition_pub(baseModel,'MIXO',cfg);

[alphaPhotonPerPigment,photonPerBiomass] = ...
    calibrateHeteroPhoton_pub(modelHETnative,cfg);

cfg.HETERO.photonPerPigment = alphaPhotonPerPigment;
cfg.HETERO.photonPerBiomass = photonPerBiomass;

if abs(cfg.HETERO.photonPerPigment - ...
        cfg.expectedPhotonPerPigment) > cfg.photonCalibrationTolerance

    error(['Calculated HETERO photon/pigment coefficient %.8g differs ' ...
           'from expected structural value %.8g. Validation stopped.'], ...
           cfg.HETERO.photonPerPigment, ...
           cfg.expectedPhotonPerPigment);
end

% Equality coupling is identical to Script 2: photon uptake is proportional
% to the HETERO biomass flux and cannot provide additional free energy.
modelHET = addPhotonBiomassCoupling_pub( ...
    modelHETnative,cfg.photon.rxn,cfg.HETERO.biomassRxn, ...
    cfg.HETERO.photonPerBiomass);

growthConditions = buildGrowthConditions_pub();

growthInternal = measuredMuValidation_pub( ...
    modelHET,modelMIX,growthConditions,cfg);

capacityInternal = substrateCapacityUnique_pub( ...
    modelHET,modelMIX,cfg);

[summaryReport,growthReport,capacityReport,settingsReport] = ...
    buildSimplifiedReports_pub( ...
        growthInternal,capacityInternal,cfg);

if isfile(cfg.resultsFile)
    delete(cfg.resultsFile);
end

writeReportSheet_pub(cfg.resultsFile,'SUMMARY', ...
    'Publication model validation - simplified supplementary report', ...
    ['Only results directly used in the manuscript model-evaluation ' ...
     'paragraph are retained. Detailed implementation settings remain ' ...
     'in this reproducibility MATLAB script.'], ...
    {'Reported result','Value','Meaning'},summaryReport);

writeReportSheet_pub(cfg.resultsFile,'GROWTH_VALIDATION', ...
    'Measured-growth validation', ...
    ['Measured apparent mu was fixed as biomass flux. The model then ' ...
     'calculated the minimum organic-carbon demand needed to support that growth.'], ...
    {'Experiment','Condition','Mode','Carbon source','Measurement', ...
     'Measured mu (d^-1)','Measured mu (h^-1)','Feasible', ...
     'Minimum carbon demand (C-mmol gDW^-1 h^-1)','Model result'}, ...
    growthReport);

writeReportSheet_pub(cfg.resultsFile,'SUBSTRATE_CAPACITY', ...
    'Unique substrate-capacity simulations', ...
    ['Five unique model conditions represent seven experimental substrate-' ...
     'condition tests. Glucose and xylose capacities were compared at the ' ...
     'same 6 C-mmol gDW^-1 h^-1 carbon supply.'], ...
    {'Condition','Mode','Carbon source','Experimental test IDs', ...
     'Tests represented','Feasible','Maximum growth (h^-1)', ...
     'Theoretical yield (g g^-1)','Experimental yield (g g^-1)', ...
     'Interpretation'},capacityReport);

writeReportSheet_pub(cfg.resultsFile,'VALIDATION_SETTINGS', ...
    'Validation settings needed to interpret the report', ...
    ['Only major constraints are shown here. Exact biomass coefficients, ' ...
     'reaction IDs and implementation details are contained directly in ' ...
     'this reproducibility MATLAB script.'], ...
    {'Setting','Value','Why it matters'},settingsReport);

fprintf('Growth validation ................. DONE\n');
fprintf('Created: 1_Galdieria_publication_growth_validation.xlsx\n');

end


function cfg = mapModel_pub(model,cfg)

cfg.ngam.rxn = requireRxn_pub(model, ...
    {'R_NGAM_ATP_Cyto','NGAM_ATP_Cyto'});

cfg.photon.rxn = requireRxn_pub(model, ...
    {'R_Photon_Cyto_tx','Photon_Cyto_tx'});

cfg.co2.rxn = requireRxn_pub(model, ...
    {'R_CO2_Cyto_tx','CO2_Cyto_tx'});
cfg.o2.rxn = requireRxn_pub(model, ...
    {'R_O2_Cyto_tx','O2_Cyto_tx'});
cfg.o2Dummy.rxn = requireRxn_pub(model, ...
    {'R_O2_Dummy_Cyto_tx','O2_Dummy_Cyto_tx'});
cfg.rubisco.rxn = requireRxn_pub(model, { ...
    'R_RIBULOSE-BISPHOSPHATE-CARBOXYLASE-RXN_Plas', ...
    'RIBULOSE-BISPHOSPHATE-CARBOXYLASE-RXN_Plas'});

cfg.basal.Fe2 = requireRxn_pub(model,{'R_FE+2_Cyto_tx','FE+2_Cyto_tx'});
cfg.basal.Mg2 = requireRxn_pub(model,{'R_MG+2_Cyto_tx','MG+2_Cyto_tx'});
cfg.basal.NH4 = requireRxn_pub(model,{'R_NH4_Cyto_tx','NH4_Cyto_tx'});
cfg.basal.NO3 = requireRxn_pub(model,{'R_NO3_Cyto_tx','NO3_Cyto_tx'});
cfg.basal.H   = requireRxn_pub(model,{'R_PROTON_Cyto_tx','PROTON_Cyto_tx'});
cfg.basal.Pi  = requireRxn_pub(model,{'R_Pi_Cyto_tx','Pi_Cyto_tx'});
cfg.basal.SO4 = requireRxn_pub(model,{'R_SO4_Cyto_tx','SO4_Cyto_tx'});
cfg.basal.H2O = requireRxn_pub(model,{'R_WATER_Cyto_tx','WATER_Cyto_tx'});

% The final trophic-state SBML no longer contains the legacy generic
% HETERO pigment-pool producer. Publication pigment pools are rebuilt from
% TOTAL_BIOMASS.xlsx below, so this legacy reaction is optional only.
cfg.pigment.originalRxn = '';
legacyPigmentCandidates = { ...
    'R_PIGMENT_TOTAL_POOL_BIO_HETEROTROPH_GLUCOSE_NOFRAC_bm', ...
    'PIGMENT_TOTAL_POOL_BIO_HETEROTROPH_GLUCOSE_NOFRAC_bm'};
for kk = 1:numel(legacyPigmentCandidates)
    rLegacy = resolveRxn_pub(model,legacyPigmentCandidates{kk});
    if ~isempty(rLegacy)
        cfg.pigment.originalRxn = rLegacy;
        break;
    end
end

cfg.pigment.PCB  = findMet_pub(model,{'3Z-PHYCOCYANOBILIN_PLAS'});
cfg.pigment.CHLA = findMet_pub(model,{'CHLOROPHYLL-A_PLAS','CHLOROPHYLL-A_THYL'});
cfg.pigment.BETA = findMet_pub(model,{'CPD1F-129_THYL'});
cfg.pigment.ZEA  = findMet_pub(model,{'CPD1F-130_THYL'});

pidx = find(strcmp(model.rxns,cfg.photon.rxn),1);

cfg.photon.nativeLB = model.lb(pidx);
cfg.photon.nativeUB = model.ub(pidx);

if abs(cfg.photon.nativeLB - (-0.2)) > 1e-12 || ...
        abs(cfg.photon.nativeUB) > 1e-12

    error(['Unexpected native photon bounds. Expected ' ...
           'R_Photon_Cyto_tx = [-0.2,0].']);
end

T = readtable(cfg.substrateFile,'Sheet',cfg.substrateSheet, ...
    'VariableNamingRule','preserve');

vars = string(T.Properties.VariableNames);

subCol = find(strcmpi(vars,'Substrate_ID'),1);

modelCol = find( ...
    contains(lower(vars),'metacyc') & ...
    contains(lower(vars),'model exchange'),1);

if isempty(subCol) || isempty(modelCol)
    error('Required substrate workbook columns were not found.');
end

subs = string(T{:,subCol});
ids  = string(T{:,modelCol});

cfg.substrateMap.allCarbonRxns = {};

for k = 1:numel(ids)
    if ~ismissing(ids(k)) && strlength(strtrim(ids(k))) > 0
        r = resolveRxn_pub(model,char(ids(k)));
        if ~isempty(r)
            cfg.substrateMap.allCarbonRxns{end+1,1} = r; %#ok<AGROW>
        end
    end
end

cfg.substrateMap.allCarbonRxns = ...
    unique(cfg.substrateMap.allCarbonRxns,'stable');

% Close every carbon-containing boundary exactly as in Script 2, not only
% the substrate-list exchanges. This prevents hidden alternative carbon input.
cfg.carbonBoundaryRxns = identifyCarbonBoundaryRxns_pub_s1(model,cfg);
cfg.carbonBoundaryRxns = unique([ ...
    string(cfg.carbonBoundaryRxns(:)); ...
    string(cfg.substrateMap.allCarbonRxns(:)); ...
    string(cfg.co2.rxn)],'stable');
cfg.carbonBoundaryRxns = cellstr(cfg.carbonBoundaryRxns);

rowG = find(strcmpi(strtrim(subs),'D-Glucose'),1);
rowX = find(strcmpi(strtrim(subs),'D-Xylose'),1);

if isempty(rowG) || isempty(rowX)
    error('D-Glucose or D-Xylose row missing from substrate workbook.');
end

cfg.substrateMap.glucose = ...
    resolveRxn_pub(model,char(ids(rowG)));

cfg.substrateMap.xylose = ...
    resolveRxn_pub(model,char(ids(rowX)));

if isempty(cfg.substrateMap.glucose)
    error('D-Glucose model boundary could not be resolved.');
end

if isempty(cfg.substrateMap.xylose)
    error('D-Xylose model boundary could not be resolved.');
end

xText = upper(string(cfg.substrateMap.xylose));

if ~contains(xText,'BETA') && isfield(model,'rxnNames')
    ix = find(strcmp(model.rxns,cfg.substrateMap.xylose),1);
    xText = xText + " " + upper(string(model.rxnNames{ix}));
end

if ~contains(xText,'BETA')
    error('D-Xylose did not resolve to the beta-D-xylose boundary.');
end

end

function [model2,info] = addPigmentPool_pub(model,mode,cfg)
% Rebuild the condition-specific pigment pool from TOTAL_BIOMASS.xlsx.
% Publication totals are applied later in the top-level BOF; this reaction
% supplies the detailed pigment composition only.

model2 = model;
mode = upper(string(mode));

if mode == "HETERO"
    poolMet = 'm_PIGMENT_TOTAL_POOL_EXP_HETERO_PUB[c]';
    rxn = 'R_PIGMENT_TOTAL_POOL_EXP_HETERO_PUB_bm';
    sheetName = 'Reaction List Heto';
    workbookRxn = 'PIGMENT_TOTAL_POOL_BIO_HETEROTROPH_GLUCOSE_NOFRAC';
elseif mode == "MIXO"
    poolMet = 'm_PIGMENT_TOTAL_POOL_EXP_MIXO_PUB[c]';
    rxn = 'R_PIGMENT_TOTAL_POOL_EXP_MIXO_PUB_bm';
    sheetName = 'Reaction List Mixo';
    workbookRxn = 'PIGMENT_TOTAL_POOL_BIO_MIXOTROPH_NOFRAC';
else
    error('Unsupported pigment mode: %s',mode);
end

formula = readWorkbookReactionFormula_pub_s1( ...
    cfg.biomassFile,sheetName,workbookRxn);

cPCB  = coefficientForWorkbookMet_pub_s1(formula,'m_3Z-PHYCOCYANOBILIN_Plas[p]');
try
    cCHLA = coefficientForWorkbookMet_pub_s1(formula,'m_CHLOROPHYLL-A_Plas[p]');
catch
    cCHLA = coefficientForWorkbookMet_pub_s1(formula,'m_CHLOROPHYLL-A_Thyl[t]');
end
cBETA = coefficientForWorkbookMet_pub_s1(formula,'m_CPD1F-129_Thyl[t]');
cZEA  = coefficientForWorkbookMet_pub_s1(formula,'m_CPD1F-130_Thyl[t]');

coeff = [cPCB;cCHLA;cBETA;cZEA];
if any(~isfinite(coeff)) || sum(coeff) <= 0
    error('Invalid pigment composition in TOTAL_BIOMASS.xlsx for %s.',mode);
end
coeff = coeff/sum(coeff);

if ~ismember(poolMet,model2.mets)
    model2 = addMetabolite(model2,poolMet, ...
        'metName',sprintf('%s publication pigment total pool',mode), ...
        'metFormula','','Charge',0);
end

if ismember(rxn,model2.rxns)
    model2 = removeRxns(model2,{rxn});
end

model2 = addReaction(model2,rxn, ...
    'metaboliteList',{ ...
        cfg.pigment.PCB
        cfg.pigment.CHLA
        cfg.pigment.BETA
        cfg.pigment.ZEA
        poolMet}, ...
    'stoichCoeffList',[-coeff;1], ...
    'lowerBound',0,'upperBound',1000, ...
    'subSystem','Biomass pigment pool');
model2 = padConstraintColumns_pub_s1(model2);

info.poolMet = poolMet;
info.rxn = rxn;
info.workbookSheet = sheetName;
info.workbookReaction = workbookRxn;

end

function model2 = closeOriginalPigmentPool_pub(model,cfg)

model2 = model;

idx = find(strcmp(model2.rxns,cfg.pigment.originalRxn),1);

if ~isempty(idx)
    model2.lb(idx) = 0;
    model2.ub(idx) = 0;
end

end

function [model2,info] = createExperimentalBOF_pub( ...
    model,mode,expComp,rnaFraction,pigmentMass,pigmentPoolMet,cfg)

model2 = model;
mode = upper(string(mode));

poolMets = resolveModePoolMets_pub_s1(model2,mode);
protein = poolMets.protein;
carb    = poolMets.carb;
lipid   = poolMets.lipid;
rna     = poolMets.rna;
dna     = poolMets.dna;

ATP = findMet_pub(model2,{'ATP_CYTO'});
H2O = findMet_pub(model2,{'WATER_CYTO'});
ADP = findMet_pub(model2,{'ADP_CYTO'});
PI  = findMet_pub(model2,{'PI_CYTO'});
H   = findMet_pub(model2,{'PROTON_CYTO'});

raw = [ ...
    expComp.protein
    expComp.carbohydrate
    expComp.lipid
    expComp.nucleic*rnaFraction
    expComp.nucleic*(1-rnaFraction)
    pigmentMass
];

rawSum = sum(raw);

if rawSum <= 0
    error('Invalid raw biomass composition for %s.',mode);
end

% Experimental fractions are g per gDW; do not normalize the measured 90%.
% Within-pool amino-acid, lipid, carbohydrate and nucleotide profiles
% remain defined by the mode-specific TOTAL_BIOMASS.xlsx pool reactions.
coeff = raw;
if abs(sum(raw(1:5))-0.9) > 1e-8 || any(~isfinite(coeff)) || any(coeff<0)
    error('Invalid measured macromolecular mass fractions for %s.',mode);
end
if rawSum > 1+1e-8
    error('Experimental BOF exceeds 1 g/gDW for %s.',mode);
end

if mode == "HETERO"

    biomassMet  = 'm_TOTAL_BIOMASS_EXP_HETERO_PUB[c]';
    assemblyRxn = 'R_TOTAL_BIOMASS_EXP_HETERO_PUB_bm';
    sinkRxn     = 'R_TOTAL_BIOMASS_EXP_HETERO_PUB_SINK_bm_tx';

elseif mode == "MIXO"

    biomassMet  = 'm_TOTAL_BIOMASS_EXP_MIXO_PUB[c]';
    assemblyRxn = 'R_TOTAL_BIOMASS_EXP_MIXO_PUB_bm';
    sinkRxn     = 'R_TOTAL_BIOMASS_EXP_MIXO_PUB_SINK_bm_tx';

else
    error('Unsupported BOF mode: %s',mode);
end

if ~ismember(biomassMet,model2.mets)
    model2 = addMetabolite(model2,biomassMet, ...
        'metName',sprintf('Experimental %s biomass',mode), ...
        'metFormula','','Charge',0);
end

model2 = addReaction(model2,assemblyRxn, ...
    'metaboliteList',{ ...
        protein
        carb
        lipid
        rna
        dna
        pigmentPoolMet
        ATP
        H2O
        ADP
        PI
        H
        biomassMet}, ...
    'stoichCoeffList',[ ...
        -coeff
        -cfg.GAM
        -cfg.GAM
         cfg.GAM
         cfg.GAM
         cfg.GAM
         1], ...
    'lowerBound',0,'upperBound',1000, ...
    'subSystem','Biomass synthesis');
model2 = padConstraintColumns_pub_s1(model2);

model2 = addReaction(model2,sinkRxn, ...
    'metaboliteList',{biomassMet}, ...
    'stoichCoeffList',-1, ...
    'lowerBound',0,'upperBound',1000, ...
    'subSystem','Biomass sink');
model2 = padConstraintColumns_pub_s1(model2);

model2 = changeObjective(model2,sinkRxn);

info.assemblyRxn = assemblyRxn;
info.sinkRxn = sinkRxn;

end

function mets = resolveModePoolMets_pub_s1(model,mode)
% Resolve detailed condition-specific macromolecular pools. The latest SBML
% contains AUTO/MIXO/HETERO pool metabolites built from TOTAL_BIOMASS.xlsx.
% Generic pool names remain as a compatibility fallback for older models.

mode = upper(string(mode));
switch mode
    case "AUTO"
        tag = 'AUTO';
    case "MIXO"
        tag = 'MIXO';
    case "HETERO"
        tag = 'HETERO';
    otherwise
        error('Unsupported biomass mode: %s',mode);
end

mets.protein = findMet_pub(model,{ ...
    sprintf('PROTEIN_TOTAL_POOL_%s_CYTO',tag), ...
    'PROTEIN_TOTAL_POOL_CYTO'});
mets.carb = findMet_pub(model,{ ...
    sprintf('CARBOHYDRATE_TOTAL_POOL_%s_CYTO',tag), ...
    'CARBOHYDRATE_TOTAL_POOL_CYTO'});
mets.lipid = findMet_pub(model,{ ...
    sprintf('LIPID_TOTAL_POOL_%s_CYTO',tag), ...
    'LIPID_TOTAL_POOL_CYTO'});
mets.rna = findMet_pub(model,{ ...
    sprintf('RNA_TOTAL_POOL_%s_CYTO',tag), ...
    'RNA_TOTAL_POOL_CYTO'});
mets.dna = findMet_pub(model,{ ...
    sprintf('DNA_TOTAL_POOL_%s_CYTO',tag), ...
    'DNA_TOTAL_POOL_CYTO'});
end

function formula = readWorkbookReactionFormula_pub_s1(file,sheet,abbr)
C = readcell(file,'Sheet',sheet);
formula = "";
for i = 1:size(C,1)
    for j = 1:size(C,2)
        v = stringCell_pub_s1(C{i,j});
        if strcmpi(strtrim(v),string(abbr))
            for k = 1:size(C,2)
                s = stringCell_pub_s1(C{i,k});
                if contains(s,"->") || contains(s,"<=>") || contains(s,"<->")
                    formula = s;
                    return;
                end
            end
        end
    end
end
error('Reaction formula not found in %s / %s for %s',file,sheet,abbr);
end

function c = coefficientForWorkbookMet_pub_s1(formula,metToken)
lhs = split(string(formula),"->");
if numel(lhs) < 2
    error('Invalid workbook reaction formula for %s.',metToken);
end
terms = split(strtrim(lhs(1))," + ");
target = strtrim(string(metToken));
for i = 1:numel(terms)
    term = strtrim(terms(i));
    tok = regexp(char(term),'^([0-9eE+\-.]+)\s+(.+)$','tokens','once');
    if isempty(tok)
        continue;
    end
    met = strtrim(string(tok{2}));
    if strcmpi(met,target)
        c = str2double(strrep(tok{1},',','.'));
        if ~isfinite(c)
            error('Invalid coefficient for %s in workbook BOF.',metToken);
        end
        return;
    end
end
error('Metabolite %s not found in workbook BOF formula.',metToken);
end

function s = stringCell_pub_s1(v)
if isempty(v)
    s = "";
elseif isstring(v) || ischar(v)
    s = string(v);
elseif isnumeric(v) || islogical(v)
    s = string(v);
else
    try
        s = string(v);
    catch
        s = "";
    end
end
end

function model2 = closeAlternativeBiomass_pub(model,keepRxns)

model2 = model;

ids = upper(string(model2.rxns));
candidates = find(contains(ids,'TOTAL_BIOMASS'));

for j = candidates(:)'
    r = model2.rxns{j};

    if ~ismember(r,keepRxns)
        model2.lb(j) = 0;
        model2.ub(j) = 0;
    end
end

end

function model2 = fixNGAM_pub(model,cfg)

model2 = model;

idx = find(strcmp(model2.rxns,cfg.ngam.rxn),1);

if isempty(idx)
    error('NGAM reaction missing.');
end

model2.lb(idx) = cfg.NGAM;
model2.ub(idx) = cfg.NGAM;

end

function m = prepareSharedCondition_pub(m,mode,cfg)
% Shared publication trophic-state setup; mirrors Script 2.
mode = upper(string(mode));

m = closeMappedCarbon_pub(m,cfg);

fields = {'Fe2','Mg2','NH4','NO3','H','Pi','SO4','H2O'};
for k = 1:numel(fields)
    r = cfg.basal.(fields{k});
    m = setDirectBounds_pub_s1(m,r,cfg.basalMediumLB,cfg.basalMediumUB);
end

nidx = find(strcmp(m.rxns,cfg.ngam.rxn),1);
m.lb(nidx) = cfg.NGAM;
m.ub(nidx) = cfg.NGAM;

if mode == "HETERO"
    bof = cfg.HETERO;
elseif mode == "MIXO"
    bof = cfg.MIXO;
else
    error('Unsupported trophic mode: %s',mode);
end

m = selectOnlyBOF_pub_s1(m,bof);
m = selectOnlyPigmentProducer_pub_s1(m,bof.pigmentRxn);

m = setUptake_pub(m,cfg.o2.rxn,cfg.o2MaxUptake);

if mode == "HETERO"
    m = setUptake_pub(m,cfg.photon.rxn,cfg.heteroPhotonMaxUptake);
    m = setDirectBounds_pub_s1(m,cfg.o2Dummy.rxn,0,0);
    m = closeUptake_pub(m,cfg.co2.rxn);
    m = setDirectBounds_pub_s1(m,cfg.rubisco.rxn,0,0);
elseif mode == "MIXO"
    m = setUptake_pub(m,cfg.photon.rxn,cfg.mixoPhotonMaxUptake);
    m = setDirectBounds_pub_s1(m,cfg.o2Dummy.rxn,cfg.o2DummyLB,cfg.o2DummyUB);
    m = openUptakePreserve_pub_s1(m,cfg.co2.rxn,cfg.mixoCO2MaxUptake);
    m = setDirectBounds_pub_s1(m,cfg.rubisco.rxn,cfg.rubiscoMinFluxMIXO,1000);
else
    error('Unsupported trophic mode: %s',mode);
end
end

function m = setDirectBounds_pub_s1(m,rxn,lb,ub)
idx = find(strcmp(m.rxns,rxn),1);
if isempty(idx), error('Reaction not found: %s',rxn); end
m.lb(idx)=lb; m.ub(idx)=ub;
end

function m = openUptakePreserve_pub_s1(m,rxn,q)
idx = find(strcmp(m.rxns,rxn),1);
if isempty(idx), error('Reaction not found: %s',rxn); end
s = full(m.S(:,idx)); nz=find(abs(s)>1e-12);
if numel(nz)~=1, error('Expected one-metabolite boundary reaction: %s',rxn); end
if s(nz)<0
    m.lb(idx)=-abs(q);
else
    m.ub(idx)=abs(q);
end
end

function [m,auditT] = closeOriginalBiomassOutputs_pub_s1(m)
ids = string(m.rxns);
if isfield(m,'rxnNames')
    names = string(m.rxnNames);
else
    names = repmat("",numel(m.rxns),1);
end
normIds = strings(numel(ids),1);
normNames = strings(numel(names),1);
for k=1:numel(ids)
    normIds(k)=upper(regexprep(char(ids(k)),'[^A-Z0-9]',''));
    normNames(k)=upper(regexprep(char(names(k)),'[^A-Z0-9]',''));
end
rawLower=lower(ids); rawNameLower=lower(names);
isBmTx = endsWith(rawLower,"_bm_tx") | endsWith(rawNameLower,"_bm_tx") | ...
         contains(normIds,"BMTX") | contains(normNames,"BMTX");
isTotal = contains(normIds,"TOTALBIOMASS") | contains(normNames,"TOTALBIOMASS");
idx=find(isBmTx | isTotal);
oldLB=m.lb(idx); oldUB=m.ub(idx); oldC=zeros(numel(idx),1);
if isfield(m,'c') && numel(m.c)==numel(m.rxns)
    oldC=m.c(idx); m.c(idx)=0;
end
m.lb(idx)=0; m.ub(idx)=0;
auditT=table(ids(idx),names(idx),oldLB,oldUB,oldC, ...
    'VariableNames',{'Reaction_ID','Reaction_name','Original_LB','Original_UB','Original_objective'});
end

function m = selectOnlyBOF_pub_s1(m,bof)
ids=string(m.rxns); names=repmat("",numel(ids),1);
if isfield(m,'rxnNames'), names=string(m.rxnNames); end
normIds=strings(numel(ids),1); normNames=strings(numel(names),1);
for k=1:numel(ids)
    normIds(k)=upper(regexprep(char(ids(k)),'[^A-Z0-9]',''));
    normNames(k)=upper(regexprep(char(names(k)),'[^A-Z0-9]',''));
end
raw=lower(ids); rawName=lower(names);
cand=find(contains(normIds,"TOTALBIOMASS") | contains(normNames,"TOTALBIOMASS") | ...
          contains(normIds,"BMTX") | contains(normNames,"BMTX") | ...
          endsWith(raw,"_bm_tx") | endsWith(rawName,"_bm_tx"));
keep=string({bof.assemblyRxn,bof.sinkRxn});
for j=cand(:)'
    if ~any(string(m.rxns{j})==keep)
        m.lb(j)=0; m.ub(j)=0;
        if isfield(m,'c') && numel(m.c)>=j, m.c(j)=0; end
    end
end
aidx=find(strcmp(m.rxns,bof.assemblyRxn),1);
sidx=find(strcmp(m.rxns,bof.sinkRxn),1);
if isempty(aidx) || isempty(sidx), error('Selected BOF reactions are missing.'); end
m.lb(aidx)=0; m.ub(aidx)=max(1000,m.ub(aidx));
m.lb(sidx)=0; m.ub(sidx)=max(1000,m.ub(sidx));
m=changeObjective(m,bof.sinkRxn);
end

function m = selectOnlyPigmentProducer_pub_s1(m,keepRxn)
ids=string(m.rxns); normIds=strings(numel(ids),1);
for k=1:numel(ids)
    normIds(k)=upper(regexprep(char(ids(k)),'[^A-Z0-9]',''));
end
closeIdx=find(contains(normIds,"PIGMENTTOTALPOOLBIO") | ...
              contains(normIds,"PIGMENTTOTALPOOLSINK") | ...
              contains(normIds,"PIGMENTTOTALPOOLEXP"));
for j=closeIdx(:)'
    m.lb(j)=0; m.ub(j)=0;
    if isfield(m,'c') && numel(m.c)>=j, m.c(j)=0; end
end
idx=find(strcmp(m.rxns,keepRxn),1);
if isempty(idx), error('Selected experimental pigment producer is missing: %s',keepRxn); end
m.lb(idx)=0; m.ub(idx)=1000;
end

function m = restoreMixoInorganicCarbon_pub_s1(m,cfg)
m = openUptakePreserve_pub_s1(m,cfg.co2.rxn,cfg.mixoCO2MaxUptake);
m = setDirectBounds_pub_s1(m,cfg.rubisco.rxn,cfg.rubiscoMinFluxMIXO,1000);
end

function m = padConstraintColumns_pub_s1(m)
if ~isfield(m,'C') || isempty(m.C), return; end
n=numel(m.rxns); nc=size(m.C,2);
if nc<n
    m.C(:,nc+1:n)=0;
elseif nc>n
    error('model.C has more columns than reactions after publication BOF construction.');
end
end

function model2 = addPhotonBiomassCoupling_pub(model,photonRxn,biomassRxn,k)
% Equality form used by Script 2.
model2 = ensureConstraintFields_pub(model);
pidx = find(strcmp(model2.rxns,photonRxn),1);
bidx = find(strcmp(model2.rxns,biomassRxn),1);
if isempty(pidx) || isempty(bidx)
    error('Photon or biomass reaction missing for HETERO coupling.');
end
s = full(model2.S(:,pidx)); nz=find(abs(s)>1e-12);
if numel(nz)~=1, error('Expected one-metabolite photon boundary.'); end
if s(nz)<0, uptakeSign=-1; else, uptakeSign=1; end
row = sparse(1,numel(model2.rxns));
row(pidx)=1;
row(bidx)=-uptakeSign*k;
model2.C(end+1,:)=row;
model2.d(end+1,1)=0;
model2.dsense(end+1,1)='E';
model2.ctrs{end+1,1}='HETERO_PHOTON_BIOMASS_COUPLING_PUB';
end

function rxns = identifyCarbonBoundaryRxns_pub_s1(model,cfg)
rxns = strings(0,1);
for j=1:numel(model.rxns)
    nz=find(abs(full(model.S(:,j)))>1e-12);
    if numel(nz)~=1, continue; end
    mi=nz(1); nC=NaN;
    if isfield(model,'metFormulas') && numel(model.metFormulas)>=mi
        nC=formulaCarbonCount_pub_s1(string(model.metFormulas{mi}));
    end
    if isnan(nC) && (any(strcmp(model.rxns{j},cfg.substrateMap.allCarbonRxns)) || ...
            strcmp(model.rxns{j},cfg.co2.rxn))
        nC=1;
    end
    if isfinite(nC) && nC>0
        rxns(end+1,1)=string(model.rxns{j}); %#ok<AGROW>
    end
end
end

function nC = formulaCarbonCount_pub_s1(formula)
nC=NaN; f=char(strtrim(string(formula)));
if isempty(f), return; end
toks=regexp(f,'([A-Z][a-z]?)([0-9]*)','tokens');
if isempty(toks), return; end
nC=0;
for k=1:numel(toks)
    if strcmp(toks{k}{1},'C')
        if isempty(toks{k}{2}), nC=nC+1;
        else, nC=nC+str2double(toks{k}{2}); end
    end
end
end

function model2 = setPhotonBounds_pub(model,lb,ub,cfg)

model2 = model;

idx = find(strcmp(model2.rxns,cfg.photon.rxn),1);

if isempty(idx)
    error('Photon boundary reaction missing.');
end

model2.lb(idx) = lb;
model2.ub(idx) = ub;

end

function [alpha,k] = calibrateHeteroPhoton_pub(model,cfg)

m = closeMappedCarbon_pub(model,cfg);

m = setUptake_pub( ...
    m,cfg.substrateMap.glucose,cfg.validationMaxCarbon/6);

bidx = find(strcmp(m.rxns,cfg.HETERO.biomassRxn),1);

m.lb(bidx) = cfg.heteroPhotonCalibrationMu;
m.ub(bidx) = cfg.heteroPhotonCalibrationMu;

pidx = find(strcmp(m.rxns,cfg.photon.rxn),1);

m.c(:) = 0;
m.c(pidx) = 1;

sol = optimizeCbModel(m,'max');

if ~isfield(sol,'stat') || sol.stat ~= 1
    error('HETERO photon calibration is infeasible.');
end

minimumPhoton = max(0,-sol.x(pidx));

k = minimumPhoton/cfg.heteroPhotonCalibrationMu;

gidx = find(strcmp(m.rxns,cfg.pigment.HETERO.rxn),1);
pigmentFlux = sol.x(gidx);

if pigmentFlux <= 1e-12
    error('HETERO pigment-pool flux is zero during photon calibration.');
end

alpha = minimumPhoton/pigmentFlux;

end

function model2 = addPhotonPigmentCoupling_pub( ...
    model,photonRxn,pigmentRxn,alpha)

model2 = ensureConstraintFields_pub(model);

pidx = find(strcmp(model2.rxns,photonRxn),1);
gidx = find(strcmp(model2.rxns,pigmentRxn),1);

row = sparse(1,numel(model2.rxns));

row(pidx) = 1;
row(gidx) = alpha;

model2.C(end+1,:) = row;
model2.d(end+1,1) = 0;
model2.dsense(end+1,1) = 'E';
model2.ctrs{end+1,1} = 'HETERO_PHOTON_PIGMENT_COUPLING_PUB';

end


function T = buildGrowthConditions_pub()

Experiment = [ ...
    "Mixed-sugar kinetics 1"
    "Mixed-sugar kinetics 1"
    "Mixed-sugar kinetics 2"
    "Mixed-sugar kinetics 2"
    "Glucose-supported kinetics"
    "Glucose-supported kinetics"
    "Glucose-supported kinetics"
    "Glucose-supported kinetics"
];

Condition = [ ...
    "KIN1_MIX_MIX"
    "KIN1_HET_MIX"
    "KIN2_MIX_MIX"
    "KIN2_HET_MIX"
    "GLC_OD_MIX"
    "GLC_OD_HET"
    "GLC_CELL_MIX"
    "GLC_CELL_HET"
];

Mode = [ ...
    "MIXO"
    "HETERO"
    "MIXO"
    "HETERO"
    "MIXO"
    "HETERO"
    "MIXO"
    "HETERO"
];

Carbon = [ ...
    "MIXED"
    "MIXED"
    "MIXED"
    "MIXED"
    "GLUCOSE"
    "GLUCOSE"
    "GLUCOSE"
    "GLUCOSE"
];

Measurement_method = [ ...
    "Two-point apparent mu, days 2-4"
    "Two-point apparent mu, days 2-4"
    "Log-linear fit, days 1-5"
    "Log-linear fit, days 1-5"
    "OD750-derived mu, days 3-5"
    "OD750-derived mu, days 3-5"
    "Cell-count-derived mu, days 3-5"
    "Cell-count-derived mu, days 3-5"
];

Measured_mu_d_1 = [1.73;1.76;1.23;1.44;1.48;1.65;1.54;1.65];

Observed_order = [ ...
    "NOT_MEASURED"
    "NOT_MEASURED"
    "GLUCOSE_FIRST_THEN_XYLOSE"
    "GLUCOSE_FIRST_THEN_XYLOSE"
    "NOT_APPLICABLE"
    "NOT_APPLICABLE"
    "NOT_APPLICABLE"
    "NOT_APPLICABLE"
];

T = table( ...
    Experiment,Condition,Mode,Carbon,Measurement_method, ...
    Measured_mu_d_1,Observed_order);

end


function Growth = measuredMuValidation_pub( ...
    modelHET,modelMIX,T,cfg)

n = height(T);

Experiment = T.Experiment;
Condition = T.Condition;
Mode = T.Mode;
Carbon = T.Carbon;
Measurement_method = T.Measurement_method;

Measured_mu_d_1 = T.Measured_mu_d_1;
Measured_mu_h_1 = Measured_mu_d_1/24;

Feasible = false(n,1);
Minimum_carbon_demand = nan(n,1);
Glucose_uptake = nan(n,1);
Xylose_uptake = nan(n,1);
Preferred_carbon = strings(n,1);
Observed_order = T.Observed_order;
Order_agreement = nan(n,1);
MIX_light_at_native_lower_bound = false(n,1);

for i = 1:n

    mode = upper(string(Mode(i)));
    carbon = upper(string(Carbon(i)));
    mu = Measured_mu_h_1(i);

    if mode == "HETERO"
        m = modelHET;
        biomassRxn = cfg.HETERO.biomassRxn;
        pigmentRxn = cfg.pigment.HETERO.rxn;
    else
        m = modelMIX;
        biomassRxn = cfg.MIXO.biomassRxn;
        pigmentRxn = cfg.pigment.MIXO.rxn;
    end

    m = prepareValidationCarbon_pub(m,carbon,cfg);
    if mode == "MIXO"
        m = restoreMixoInorganicCarbon_pub_s1(m,cfg);
    end

    bidx = find(strcmp(m.rxns,biomassRxn),1);
    m.lb(bidx) = mu;
    m.ub(bidx) = mu;

    sol = minimumCarbonSolve_pub(m,carbon,cfg);

    if ~sol.feasible
        continue;
    end

    Feasible(i) = true;
    Minimum_carbon_demand(i) = sol.carbon;
    Glucose_uptake(i) = sol.qGlc;
    Xylose_uptake(i) = sol.qXyl;

    if sol.qGlc > 1e-9 && sol.qXyl <= 1e-9
        Preferred_carbon(i) = "GLUCOSE";
    elseif sol.qXyl > 1e-9 && sol.qGlc <= 1e-9
        Preferred_carbon(i) = "XYLOSE";
    elseif sol.qGlc > 1e-9 && sol.qXyl > 1e-9
        Preferred_carbon(i) = "BOTH";
    else
        Preferred_carbon(i) = "NONE";
    end

    if Observed_order(i) == "GLUCOSE_FIRST_THEN_XYLOSE"
        Order_agreement(i) = double(Preferred_carbon(i)=="GLUCOSE");
    end

    pidx = find(strcmp(m.rxns,cfg.photon.rxn),1);
    nidx = find(strcmp(m.rxns,cfg.ngam.rxn),1);
    gidx = find(strcmp(m.rxns,pigmentRxn),1);

    photonFlux = sol.solution.x(pidx);

    if abs(sol.solution.x(nidx)-cfg.NGAM) > 1e-7
        error('NGAM flux differs from fixed value in %s.',Condition(i));
    end

    if mode == "HETERO"
        pigmentFlux = sol.solution.x(gidx);

        if pigmentFlux > 1e-12
            observedCoupling = (-photonFlux)/pigmentFlux;

            if abs(observedCoupling - ...
                    cfg.HETERO.photonPerPigment) > 1e-7
                error('HETERO photon/pigment coupling failed in %s.', ...
                    Condition(i));
            end
        end
    else
        MIX_light_at_native_lower_bound(i) = ...
            abs(photonFlux-cfg.photon.nativeLB) < 1e-8;
    end
end

Growth = table( ...
    Experiment,Condition,Mode,Carbon,Measurement_method, ...
    Measured_mu_d_1,Measured_mu_h_1,Feasible, ...
    Minimum_carbon_demand,Glucose_uptake,Xylose_uptake, ...
    Preferred_carbon,Observed_order,Order_agreement, ...
    MIX_light_at_native_lower_bound);

end

function out = minimumCarbonSolve_pub(m,carbon,cfg)

out.feasible = false;
out.carbon = NaN;
out.qGlc = NaN;
out.qXyl = NaN;
out.solution = [];

[rxns,C] = testedCarbonRxns_pub(carbon,cfg);

m.c(:) = 0;

for k = 1:numel(rxns)
    idx = find(strcmp(m.rxns,rxns{k}),1);
    m.c(idx) = -C(k);
end

solution = optimizeCbModel(m,'min');

if ~isfield(solution,'stat') || solution.stat ~= 1
    return;
end

qGlc = getUptake_pub(solution,m,cfg.substrateMap.glucose);
qXyl = getUptake_pub(solution,m,cfg.substrateMap.xylose);

out.feasible = true;
out.qGlc = qGlc;
out.qXyl = qXyl;
out.carbon = 6*qGlc + 5*qXyl;
out.solution = solution;

end


function Capacity = substrateCapacityUnique_pub( ...
    modelHET,modelMIX,cfg)

Unique_condition = [ ...
    "MIX_GLC"
    "MIX_XYL"
    "HET_GLC"
    "HET_XYL"
    "HET_MIX"
];

Mode = [ ...
    "MIXO"
    "MIXO"
    "HETERO"
    "HETERO"
    "HETERO"
];

Carbon = [ ...
    "GLUCOSE"
    "XYLOSE"
    "GLUCOSE"
    "XYLOSE"
    "MIXED"
];

Experimental_test_IDs = [ ...
    "SCREEN1_MIX_GLC"
    "SCREEN1_MIX_XYL"
    "SCREEN1_HET_GLC; SCREEN2_HET_GLC"
    "SCREEN1_HET_XYL; SCREEN2_HET_XYL"
    "SCREEN2_HET_MIX"
];

Experimental_tests_represented = [1;1;2;2;1];

Experimental_endpoint_yield = [NaN;NaN;0.442;0.397;0.417];

n = numel(Unique_condition);

Feasible = false(n,1);
Capacity_mu_h_1 = nan(n,1);
Theoretical_yield = nan(n,1);
Glucose_uptake = nan(n,1);
Xylose_uptake = nan(n,1);
MIX_light_at_native_lower_bound = false(n,1);

for i = 1:n

    mode = upper(string(Mode(i)));
    carbon = upper(string(Carbon(i)));

    if mode == "HETERO"
        m = modelHET;
        biomassRxn = cfg.HETERO.biomassRxn;
    else
        m = modelMIX;
        biomassRxn = cfg.MIXO.biomassRxn;
    end

    m = setScreenCarbon_pub(m,carbon,cfg);
    if mode == "MIXO"
        m = restoreMixoInorganicCarbon_pub_s1(m,cfg);
    end
    m = changeObjective(m,biomassRxn);

    sol = optimizeCbModel(m,'max');

    if ~isfield(sol,'stat') || sol.stat ~= 1
        continue;
    end

    Feasible(i) = true;
    Capacity_mu_h_1(i) = sol.f;

    qGlc = getUptake_pub(sol,m,cfg.substrateMap.glucose);
    qXyl = getUptake_pub(sol,m,cfg.substrateMap.xylose);

    Glucose_uptake(i) = qGlc;
    Xylose_uptake(i) = qXyl;

    substrateMass = ...
        qGlc*cfg.MW.glucose/1000 + ...
        qXyl*cfg.MW.xylose/1000;

    if substrateMass > 0
        Theoretical_yield(i) = sol.f/substrateMass;
    end

    pidx = find(strcmp(m.rxns,cfg.photon.rxn),1);
    nidx = find(strcmp(m.rxns,cfg.ngam.rxn),1);

    photonFlux = sol.x(pidx);

    if abs(sol.x(nidx)-cfg.NGAM) > 1e-7
        error('NGAM flux differs from fixed value in %s.',Unique_condition(i));
    end

    if mode == "MIXO"
        MIX_light_at_native_lower_bound(i) = ...
            abs(photonFlux-cfg.photon.nativeLB) < 1e-8;
    end
end

Capacity = table( ...
    Unique_condition,Mode,Carbon,Experimental_test_IDs, ...
    Experimental_tests_represented,Feasible,Capacity_mu_h_1, ...
    Theoretical_yield,Glucose_uptake,Xylose_uptake, ...
    Experimental_endpoint_yield,MIX_light_at_native_lower_bound);

end


function [Summary,GrowthReport,CapacityReport,Settings] = ...
    buildSimplifiedReports_pub(G,C,cfg)

Model_result = repmat("Measured growth feasible",height(G),1);

for i=1:height(G)
    if contains(G.Carbon(i),"MIXED")
        if G.Preferred_carbon(i)=="GLUCOSE" && ...
                G.Order_agreement(i)==1
            Model_result(i) = ...
                "Glucose selected; agrees with observed glucose-first depletion";
        elseif G.Preferred_carbon(i)=="GLUCOSE"
            Model_result(i) = ...
                "Glucose selected in minimum-carbon solution";
        elseif G.Preferred_carbon(i)=="XYLOSE"
            Model_result(i) = ...
                "Xylose selected in minimum-carbon solution";
        elseif G.Preferred_carbon(i)=="BOTH"
            Model_result(i) = ...
                "Both sugars used in minimum-carbon solution";
        else
            Model_result(i) = ...
                "No organic carbon required at fixed growth";
        end
    end
end

Carbon_source = G.Carbon;
Carbon_source(Carbon_source=="MIXED") = "Mixed glucose + xylose";
Carbon_source(Carbon_source=="GLUCOSE") = "Glucose";

GrowthReport = table( ...
    G.Experiment,G.Condition,G.Mode,Carbon_source,G.Measurement_method, ...
    G.Measured_mu_d_1,G.Measured_mu_h_1,G.Feasible, ...
    G.Minimum_carbon_demand,Model_result);

Interpretation = strings(height(C),1);

for i=1:height(C)
    if C.Unique_condition(i)=="MIX_GLC"
        Interpretation(i)="Glucose supports slightly higher capacity than xylose";
    elseif C.Unique_condition(i)=="MIX_XYL"
        Interpretation(i)="Xylose growth feasible";
    elseif C.Unique_condition(i)=="HET_GLC"
        Interpretation(i)="Higher capacity and theoretical yield than xylose";
    elseif C.Unique_condition(i)=="HET_XYL"
        Interpretation(i)="Xylose growth feasible";
    else
        Interpretation(i)="Mixed feed growth feasible";
    end
end

CapCarbon = C.Carbon;
CapCarbon(CapCarbon=="GLUCOSE") = "Glucose";
CapCarbon(CapCarbon=="XYLOSE") = "Xylose";
CapCarbon(CapCarbon=="MIXED") = "Glucose + xylose";

CapacityReport = table( ...
    C.Unique_condition,C.Mode,CapCarbon,C.Experimental_test_IDs, ...
    C.Experimental_tests_represented,C.Feasible,C.Capacity_mu_h_1, ...
    C.Theoretical_yield,C.Experimental_endpoint_yield,Interpretation);

experimentalTests = sum(C.Experimental_tests_represented);
feasibleTests = sum(C.Experimental_tests_represented(C.Feasible));

muMin = min(G.Measured_mu_h_1);
muMax = max(G.Measured_mu_h_1);

carbonMin = min(G.Minimum_carbon_demand);
carbonMax = max(G.Minimum_carbon_demand);

kin2mask = ismember(G.Condition,["KIN2_MIX_MIX","KIN2_HET_MIX"]);
orderMatched = sum(G.Order_agreement(kin2mask)==1);

r = C(C.Unique_condition=="HET_GLC",:);
hetGlcMu = r.Capacity_mu_h_1(1);
hetGlcYield = r.Theoretical_yield(1);

r = C(C.Unique_condition=="HET_XYL",:);
hetXylMu = r.Capacity_mu_h_1(1);
hetXylYield = r.Theoretical_yield(1);

r = C(C.Unique_condition=="MIX_GLC",:);
mixGlcMu = r.Capacity_mu_h_1(1);

r = C(C.Unique_condition=="MIX_XYL",:);
mixXylMu = r.Capacity_mu_h_1(1);

mixMask = G.Mode=="MIXO";
mixAtPhotonBound = sum(G.MIX_light_at_native_lower_bound(mixMask));

Reported_result = [ ...
    "Experimental substrate-condition tests feasible"
    "Quantitative growth measurements"
    "Measured apparent mu range"
    "Minimum model carbon demand range"
    "Mixed-sugar ordered-use agreement"
    "HET glucose capacity / theoretical yield"
    "HET xylose capacity / theoretical yield"
    "MIX glucose / xylose capacity"
    "NGAM"
    "MIX light limitation"
];

Value = [ ...
    string(feasibleTests) + " of " + string(experimentalTests)
    string(height(G)) + " measurements from " + ...
        string(numel(unique(G.Experiment))) + " experiments"
    sprintf('%.4f-%.4f h^-1',muMin,muMax)
    sprintf('%.2f-%.2f C-mmol gDW^-1 h^-1',carbonMin,carbonMax)
    string(orderMatched) + " of 2 KIN2 conditions"
    sprintf('%.4f h^-1 / %.3f g g^-1',hetGlcMu,hetGlcYield)
    sprintf('%.4f h^-1 / %.3f g g^-1',hetXylMu,hetXylYield)
    sprintf('%.4f / %.4f h^-1',mixGlcMu,mixXylMu)
    sprintf('%.1f mmol ATP gDW^-1 h^-1',cfg.NGAM)
    sprintf("Finite MIXO photon cap = %.0f mmol gDW^-1 h^-1",cfg.mixoPhotonMaxUptake)
];

Meaning = [ ...
    "All tested growth conditions were feasible"
    "KIN1, KIN2 and glucose-supported kinetics"
    "Experimental mu fixed as biomass flux"
    "Calculated minimum organic-carbon requirement"
    "HET KIN2 selected glucose; MIX KIN2 needed no organic carbon"
    "Standardized 6 C-mmol supply"
    "Standardized 6 C-mmol supply"
    "Standardized 6 C-mmol supply"
    "Fixed in all simulations"
    "Same MIXO light constraint as Script 2"
];

Summary = table(Reported_result,Value,Meaning);

Setting = [ ...
    "Model"
    "Biomass objectives"
    "NGAM"
    "GAM"
    "Measured-growth validation"
    "Mixed glucose + xylose"
    "Capacity comparison"
    "MIX light"
    "HET light"
];

SettingValue = [ ...
    "Final supplied SBML model"
    "Condition-specific MIXO and HETERO experimental BOFs"
    sprintf('%.1f mmol ATP gDW^-1 h^-1',cfg.NGAM)
    sprintf('%.0f mmol ATP gDW^-1 biomass',cfg.GAM)
    "Fix measured mu, then minimize organic-carbon uptake"
    "Both substrates available; no forced uptake ratio"
    sprintf('%.0f C-mmol gDW^-1 h^-1',cfg.screenCarbon)
    sprintf('Photon uptake capped at %.0f mmol gDW^-1 h^-1', ...
        cfg.mixoPhotonMaxUptake)
    sprintf('Pigment-linked photon requirement; coupling = %.2f', ...
        cfg.HETERO.photonPerPigment)
];

Why_it_matters = [ ...
    "Galdieria_sulphuraria_SBML_Final.xml"
    "Exact coefficients are defined directly in this reproduction script"
    "Fixed LB = UB"
    "Included in biomass objective"
    "Substrate uptake is model-derived, not experimentally fitted"
    "Model may select the energetically preferred substrate"
    "Same carbon supply used for glucose and xylose comparison"
    "Same finite MIXO photon cap as Script 2"
    "Retains mandatory small HET light requirement"
];

Settings = table(Setting,SettingValue,Why_it_matters);

end


function writeReportSheet_pub(file,sheetName,titleText,noteText,headers,T)

writecell({titleText},file,'Sheet',sheetName,'Range','A1');
writecell({noteText},file,'Sheet',sheetName,'Range','A2');
writecell(headers,file,'Sheet',sheetName,'Range','A4');

data = table2cell(T);

if ~isempty(data)
    writecell(data,file,'Sheet',sheetName,'Range','A5');
end

end


function m = prepareValidationCarbon_pub(m,carbon,cfg)

m = closeMappedCarbon_pub(m,cfg);

switch upper(string(carbon))

    case "GLUCOSE"
        m = setUptake_pub( ...
            m,cfg.substrateMap.glucose, ...
            cfg.validationMaxCarbon/6);

    case "XYLOSE"
        m = setUptake_pub( ...
            m,cfg.substrateMap.xylose, ...
            cfg.validationMaxCarbon/5);

    case "MIXED"
        m = setUptake_pub( ...
            m,cfg.substrateMap.glucose, ...
            cfg.validationMaxCarbon/6);

        m = setUptake_pub( ...
            m,cfg.substrateMap.xylose, ...
            cfg.validationMaxCarbon/5);

    otherwise
        error('Unsupported carbon condition: %s',carbon);
end

end

function m = setScreenCarbon_pub(m,carbon,cfg)

m = closeMappedCarbon_pub(m,cfg);

switch upper(string(carbon))

    case "GLUCOSE"
        m = setUptake_pub( ...
            m,cfg.substrateMap.glucose, ...
            cfg.screenCarbon/6);

    case "XYLOSE"
        m = setUptake_pub( ...
            m,cfg.substrateMap.xylose, ...
            cfg.screenCarbon/5);

    case "MIXED"
        m = setUptake_pub( ...
            m,cfg.substrateMap.glucose, ...
            (cfg.screenCarbon/2)/6);

        m = setUptake_pub( ...
            m,cfg.substrateMap.xylose, ...
            (cfg.screenCarbon/2)/5);

    otherwise
        error('Unsupported carbon condition: %s',carbon);
end

end

function [rxns,C] = testedCarbonRxns_pub(carbon,cfg)

switch upper(string(carbon))

    case "GLUCOSE"
        rxns = {cfg.substrateMap.glucose};
        C = 6;

    case "XYLOSE"
        rxns = {cfg.substrateMap.xylose};
        C = 5;

    case "MIXED"
        rxns = { ...
            cfg.substrateMap.glucose, ...
            cfg.substrateMap.xylose};
        C = [6 5];

    otherwise
        error('Unsupported carbon condition: %s',carbon);
end

end

function m = closeMappedCarbon_pub(m,cfg)

for k = 1:numel(cfg.carbonBoundaryRxns)
    r = cfg.carbonBoundaryRxns{k};
    if ismember(r,m.rxns)
        m = closeUptake_pub(m,r);
    end
end

end

function m = closeUptake_pub(m,rxn)

idx = find(strcmp(m.rxns,rxn),1);

if isempty(idx)
    return;
end

s = full(m.S(:,idx));
nz = find(abs(s)>0);

if numel(nz) ~= 1
    error('Expected one-metabolite boundary reaction: %s',rxn);
end

if s(nz) < 0
    m.lb(idx) = max(m.lb(idx),0);
else
    m.ub(idx) = min(m.ub(idx),0);
end

end

function m = setUptake_pub(m,rxn,q)

idx = find(strcmp(m.rxns,rxn),1);

if isempty(idx)
    error('Reaction not found: %s',rxn);
end

s = full(m.S(:,idx));
nz = find(abs(s)>0);

if numel(nz) ~= 1
    error('Expected one-metabolite boundary reaction: %s',rxn);
end

if s(nz) < 0
    m.lb(idx) = -abs(q);
    m.ub(idx) = 0;
else
    m.lb(idx) = 0;
    m.ub(idx) = abs(q);
end

end

function q = getUptake_pub(sol,m,rxn)

q = 0;

idx = find(strcmp(m.rxns,rxn),1);

if isempty(idx)
    return;
end

s = full(m.S(:,idx));
nz = find(abs(s)>0);

if numel(nz) ~= 1
    return;
end

v = sol.x(idx);

if s(nz) < 0
    q = max(0,-v);
else
    q = max(0,v);
end

end

function m = ensureConstraintFields_pub(m)

if ~isfield(m,'C') || isempty(m.C)

    m.C = sparse(0,numel(m.rxns));
    m.d = zeros(0,1);
    m.dsense = char(zeros(0,1));
    m.ctrs = cell(0,1);

end

end


function r = requireRxn_pub(model,candidates)

r = '';

for k = 1:numel(candidates)

    r = resolveRxn_pub(model,candidates{k});

    if ~isempty(r)
        return;
    end
end

error('Required reaction not found: %s',strjoin(candidates,' / '));

end

function r = resolveRxn_pub(model,q)

r = '';

idx = find(strcmp(model.rxns,string(q)),1);

if isempty(idx) && isfield(model,'rxnNames')
    idx = find(strcmpi(string(model.rxnNames),string(q)),1);
end

if isempty(idx)

    nq = normalizeID_pub(q);

    nr = string(cellfun( ...
        @normalizeID_pub,model.rxns, ...
        'UniformOutput',false));

    idx = find(strcmp(nr,nq),1);
end

if ~isempty(idx)
    r = model.rxns{idx};
end

end

function s = normalizeID_pub(x)

s = upper(regexprep(char(string(x)),'[^A-Za-z0-9]',''));

if startsWith(s,'R') && strlength(string(s)) > 4
    s = char(extractAfter(string(s),1));
end

end

function met = findMet_pub(model,tokens)
% Resolve metabolites by exact normalized biochemical core first.
% This is intentionally identical in principle to Script 2 and prevents
% ambiguous substring matches such as ADP_CYTO -> DADP_Cyto or 3-5-ADP.

ids = string(model.mets);
normIds = strings(numel(ids),1);
for j = 1:numel(ids)
    normIds(j) = string(normalizeMetCore_pub_s1(ids(j)));
end

if isfield(model,'metNames')
    names = string(model.metNames);
    normNames = strings(numel(names),1);
    for j = 1:numel(names)
        normNames(j) = string(normalizeMetCore_pub_s1(names(j)));
    end
else
    names = repmat("",numel(ids),1);
    normNames = repmat("",numel(ids),1);
end

for k = 1:numel(tokens)
    target = string(normalizeMetCore_pub_s1(tokens{k}));
    idx = find(normIds == target);
    if isempty(idx)
        idx = find(normNames == target);
    end
    idx = unique(idx,'stable');
    if numel(idx) == 1
        met = model.mets{idx};
        return;
    elseif numel(idx) > 1
        exactId = find(normIds(idx) == target);
        if numel(exactId) == 1
            met = model.mets{idx(exactId)};
            return;
        end
        error('Ambiguous exact metabolite mapping for %s: %s', ...
            char(string(tokens{k})),strjoin(cellstr(ids(idx)),', '));
    end
end

% Conservative fallback only when no exact normalized core exists.
labels = upper(ids);
if isfield(model,'metNames')
    labels = labels + " " + upper(names);
end
for k = 1:numel(tokens)
    rawToken = upper(string(tokens{k}));
    idx = find(contains(labels,rawToken));
    idx = unique(idx,'stable');
    if numel(idx) == 1
        met = model.mets{idx};
        return;
    elseif numel(idx) > 1
        error('Ambiguous fallback metabolite mapping for %s: %s', ...
            char(rawToken),strjoin(cellstr(ids(idx)),', '));
    end
end

error('Required metabolite not found: %s',strjoin(tokens,', '));
end

function s = normalizeMetCore_pub_s1(x)
s = upper(strtrim(char(string(x))));
s = regexprep(s,'^M_','');
s = regexprep(s,'\[[A-Z]+\]$','');
s = regexprep(s,'[^A-Z0-9]','');
end
```
