## Supplementary Data 3 for "Experimental trophic flexibility guides a compartmentalized genome-scale model of red microalgae *Galdieria sulphuraria*": S2_Galdieria_publication_substrate_optimisation.m

```
function [summaryResults,longResults,cfg] = S2_Galdieria_publication_substrate_optimisation(modelInput, substrateFile, biomassFile)
% Galdieria sulphuraria sole-substrate optimisation analysis.
% Inputs are loaded automatically from the shared Inputs folder unless supplied.
% Output: 2_Galdieria_publication_substrate_optimisation.xlsx.
scriptDir = fileparts(mfilename('fullpath'));
if isempty(scriptDir), scriptDir = pwd; end
supplementDir = fileparts(fileparts(scriptDir));
inputDir = fullfile(supplementDir,'Inputs');

if nargin < 1 || isempty(modelInput)
    modelInput = fullfile(inputDir,'Galdieria_sulphuraria_SBML_Final.xml');
end
if nargin < 2 || isempty(substrateFile)
    substrateFile = fullfile(inputDir,'Model_Substrate_list.xlsx');
end
if nargin < 3 || isempty(biomassFile)
    biomassFile = fullfile(inputDir,'TOTAL_BIOMASS.xlsx');
end

ensureCobraReady_pub();
ensureLPSolverSelected_pub();

if isstruct(modelInput)
    baseModel = modelInput;
    cfgModelSource_pub = "COBRA struct supplied by user";
elseif ischar(modelInput) || isstring(modelInput)
    modelFile = char(string(modelInput));
    if ~isfile(modelFile)
        error('Model file not found: %s',modelFile);
    end
    [~,~,ext] = fileparts(modelFile);
    if strcmpi(ext,'.mat')
        baseModel = loadCobraModelFromMat_pub(modelFile);
        cfgModelSource_pub = "MATLAB COBRA model: " + string(modelFile);
    elseif any(strcmpi(ext,{'.xml','.sbml'}))
        [baseModel,loaderUsed] = loadSBMLModel_pub(modelFile);
        cfgModelSource_pub = string(loaderUsed) + ": " + string(modelFile);
    else
        error('Unsupported model file extension: %s. Use .xml, .sbml, .mat, or a COBRA struct.',ext);
    end
else
    error('modelInput must be a COBRA model struct or a .mat/.xml/.sbml model filename.');
end

requiredFields = {'rxns','mets','S','lb','ub'};
for k = 1:numel(requiredFields)
    if ~isfield(baseModel,requiredFields{k})
        error('Input model is missing required field: %s',requiredFields{k});
    end
end

substrateFile = char(string(substrateFile));
biomassFile = char(string(biomassFile));
if ~isfile(substrateFile)
    error('Substrate workbook not found: %s',substrateFile);
end
if ~isfile(biomassFile)
    error('Biomass workbook not found: %s',biomassFile);
end


cfg.scriptDir = scriptDir;
cfg.substrateFile = substrateFile;
cfg.biomassFile = biomassFile;
cfg.modelSource = cfgModelSource_pub;
cfg.substrateSheet = 'Sole_substrate_ScrumPy';
cfg.resultsFile = fullfile(scriptDir,'2_Galdieria_publication_substrate_optimisation.xlsx');

cfg.referenceMu_d = 1.44;                 % d^-1, KIN2_HET_MIX
cfg.referenceMu_h = cfg.referenceMu_d/24; % = 0.060 h^-1
cfg.referenceMuCondition = 'KIN2_HET_MIX';
cfg.referenceMuMethod = 'Log-linear fit, days 1-5';
assert(abs(cfg.referenceMu_h-0.060) < 1e-12);

cfg.NGAM = 1.7;
cfg.GAM = 30;
cfg.screenCarbon = 6;          % C-mmol gDW^-1 h^-1
cfg.validationMaxCarbon = 120; % non-binding carbon opening for min-demand solve
cfg.fvaCarbonTolerance = 0.05; % <= 105% of minimum total carbon

cfg.o2MaxUptake = 1000;
cfg.autoPhotonProbeMax = 20;
cfg.autoNativePhotonDiagnostic = 0.2;
cfg.heteroPhotonMaxUptake = 0.2;

cfg.mixoPhotonMaxUptake = 50;

cfg.autoCO2CarbonCap = cfg.screenCarbon;
cfg.mixoCO2MaxUptake = cfg.validationMaxCarbon;
cfg.rubiscoMinFluxAUTO = 0.01;
cfg.rubiscoMinFluxMIXO = 0.01;

cfg.expectedPhotonPerPigment = 0.58;
cfg.photonCalibrationTolerance = 0.02;
cfg.heteroPhotonCalibrationMu = cfg.referenceMu_h;

cfg.growthTol = 1e-8;
cfg.uptakeTol = 1e-9;
cfg.biomassFractionForUseTest = 0.99;
cfg.autoPhotonCalibrationFraction = 0.99;
cfg.failureProbeSubstrateMultiplier = 10;
cfg.strictCarbonBoundaryClosure = true;
cfg.carbonBoundaryExemptRxns = {};
cfg.autoBoundaryRescueCarbon = cfg.screenCarbon;
cfg.autoDiagnosticFluxTol = 1e-9;
cfg.autoDiagnosticDemandUB = 1000;
cfg.mixoCO2ScanFractions = [0.99 0.90 0.50 0.10];

cfg.basalMediumLB = -1000;
cfg.basalMediumUB = 1000;
cfg.o2DummyLB = 0;
cfg.o2DummyUB = 1000;

cfg.exp.MIXO.protein       = 0.4147;
cfg.exp.MIXO.carbohydrate  = 0.3377;
cfg.exp.MIXO.nucleic       = 0.0775;
cfg.exp.MIXO.lipid         = 0.0701;

cfg.exp.HETERO.protein      = 0.4016;
cfg.exp.HETERO.carbohydrate = 0.3609;
cfg.exp.HETERO.nucleic      = 0.0765;
cfg.exp.HETERO.lipid        = 0.0610;

cfg.reference.MIXO.rnaFraction = ...
    0.06222707/(0.06222707+0.01091703);
cfg.reference.HETERO.rnaFraction = ...
    0.085/(0.085+0.015);

cfg.reference.HETERO.pigmentMass = 0.00485;
cfg.reference.MIXO.PCB_mg  = 1.714658;
cfg.reference.MIXO.CHLA_mg = 4.450000;
cfg.reference.MIXO.BETA_mg = 0.240000;
cfg.reference.MIXO.ZEA_mg  = 0.350000;
cfg.reference.MIXO.pigmentMass = ...
    (cfg.reference.MIXO.PCB_mg + cfg.reference.MIXO.CHLA_mg + ...
     cfg.reference.MIXO.BETA_mg + cfg.reference.MIXO.ZEA_mg)/1000;

T = readtable(cfg.substrateFile,'Sheet',cfg.substrateSheet, ...
    'VariableNamingRule','preserve');

requiredCols = {'Substrate_ID','Class','Growth (1995)', ...
                'MetaCyc / model exchange ID'};
for k = 1:numel(requiredCols)
    if ~ismember(requiredCols{k},T.Properties.VariableNames)
        error('Required workbook column not found: %s',requiredCols{k});
    end
end

substrateName = string(T.('Substrate_ID'));
substrateClass = string(T.('Class'));
expGrowthText = string(T.('Growth (1995)'));
exchangeText = string(T.('MetaCyc / model exchange ID'));

if ismember('Modeling notes',T.Properties.VariableNames)
    modelNotes = string(T.('Modeling notes'));
else
    modelNotes = repmat("",height(T),1);
end

resolvedExchange = strings(height(T),1);
carbonAtoms = nan(height(T),1);
maxSubstrateUptake = nan(height(T),1);
expectedGrowth = nan(height(T),1);

for i = 1:height(T)
    resolvedExchange(i) = string(resolveRxn_pub(baseModel,exchangeText(i),true));
    if strlength(resolvedExchange(i)) == 0
        warning('Substrate exchange could not be resolved: %s -> %s', ...
            substrateName(i),exchangeText(i));
    end
    carbonAtoms(i) = inferCarbonAtoms_pub(substrateName(i),substrateClass(i));
    maxSubstrateUptake(i) = cfg.screenCarbon / carbonAtoms(i);
    expectedGrowth(i) = parseGrowthPhenotype_pub(expGrowthText(i));
end

cfg.allMappedCarbonRxns = cellstr(unique( ...
    resolvedExchange(strlength(resolvedExchange)>0),'stable'));

duplicateExchange = false(height(T),1);
for i = 1:height(T)
    if strlength(resolvedExchange(i)) > 0
        duplicateExchange(i) = sum(resolvedExchange == resolvedExchange(i)) > 1;
    end
end

cfg.ngamRxn = requireRxn_pub(baseModel, ...
    {'R_NGAM_ATP_Cyto','NGAM_ATP_Cyto','ATPM_NGAM_Cyto'});
cfg.photonRxn = requireRxn_pub(baseModel, ...
    {'R_Photon_Cyto_tx','Photon_Cyto_tx'});
cfg.co2Rxn = requireRxn_pub(baseModel, ...
    {'R_CO2_Cyto_tx','CO2_Cyto_tx'});
cfg.o2Rxn = requireRxn_pub(baseModel, ...
    {'R_O2_Cyto_tx','O2_Cyto_tx'});
cfg.o2DummyRxn = requireRxn_pub(baseModel, ...
    {'R_O2_Dummy_Cyto_tx','O2_Dummy_Cyto_tx'});

cfg.basalRxns.Fe2 = requireRxn_pub(baseModel, ...
    {'R_FE+2_Cyto_tx','FE+2_Cyto_tx'});
cfg.basalRxns.Mg2 = requireRxn_pub(baseModel, ...
    {'R_MG+2_Cyto_tx','MG+2_Cyto_tx'});
cfg.basalRxns.NH4 = requireRxn_pub(baseModel, ...
    {'R_NH4_Cyto_tx','NH4_Cyto_tx'});
cfg.basalRxns.NO3 = requireRxn_pub(baseModel, ...
    {'R_NO3_Cyto_tx','NO3_Cyto_tx'});
cfg.basalRxns.H = requireRxn_pub(baseModel, ...
    {'R_PROTON_Cyto_tx','PROTON_Cyto_tx'});
cfg.basalRxns.Pi = requireRxn_pub(baseModel, ...
    {'R_Pi_Cyto_tx','Pi_Cyto_tx'});
cfg.basalRxns.SO4 = requireRxn_pub(baseModel, ...
    {'R_SO4_Cyto_tx','SO4_Cyto_tx'});
cfg.basalRxns.H2O = requireRxn_pub(baseModel, ...
    {'R_WATER_Cyto_tx','WATER_Cyto_tx'});

cfg.rubiscoRxn = requireRxn_pub(baseModel, { ...
    'R_RIBULOSE-BISPHOSPHATE-CARBOXYLASE-RXN_Plas', ...
    'RIBULOSE-BISPHOSPHATE-CARBOXYLASE-RXN_Plas'});

[cfg.carbonBoundaryRxns,carbonBoundaryAudit] = ...
    identifyCarbonBoundaryRxns_pub(baseModel,cfg);
cfg.carbonBoundaryRxns = unique([ ...
    string(cfg.carbonBoundaryRxns(:)); ...
    string(cfg.allMappedCarbonRxns(:)); ...
    string(cfg.co2Rxn)],'stable');
cfg.carbonBoundaryRxns = cellstr(cfg.carbonBoundaryRxns);

[baseModel,originalBofAudit] = closeOriginalBiomassOutputs_pub(baseModel);
cfg.preexistingClosedBiomassRxns = cellstr(string(originalBofAudit.Reaction_ID));

[baseModel,cfg,workbookMeanAudit] = implementWorkbookMeanBOFs_pub( ...
    baseModel,cfg,cfg.biomassFile);

autoBOF = cfg.AUTO;

[baseModel,cfg,experimentalBofAudit] = ...
    implementExperimentalValidationBOFs_pub(baseModel,cfg);

cfg.AUTO = autoBOF;


modelAUTO = prepareCondition_pub(baseModel,'AUTO',cfg);
modelMIXO = prepareCondition_pub(baseModel,'MIXO',cfg);
modelHETERO_uncoupled = prepareCondition_pub(baseModel,'HETERO',cfg);

assertBasalMedium_pub(modelAUTO,cfg,'AUTO');
assertBasalMedium_pub(modelMIXO,cfg,'MIXO');
assertBasalMedium_pub(modelHETERO_uncoupled,cfg,'HETERO');

assertO2DummyBounds_pub(modelAUTO,cfg.o2DummyRxn,0,1000,'AUTO');
assertO2DummyBounds_pub(modelMIXO,cfg.o2DummyRxn,0,1000,'MIXO');
assertO2DummyBounds_pub(modelHETERO_uncoupled,cfg.o2DummyRxn,0,0,'HETERO');

rowGlc = find(strcmpi(strtrim(substrateName),'D-Glucose'),1);
if isempty(rowGlc) || strlength(resolvedExchange(rowGlc))==0
    error('D-Glucose is required for HETERO photon calibration.');
end

[alpha,kPhoton,heteroPhotonCalibration] = calibrateHeteroPhoton_pub( ...
    modelHETERO_uncoupled,char(resolvedExchange(rowGlc)), ...
    cfg.validationMaxCarbon/carbonAtoms(rowGlc),cfg);

cfg.HETERO.photonPerPigment = alpha;
cfg.HETERO.photonPerBiomass = kPhoton;

if abs(alpha-cfg.expectedPhotonPerPigment) > cfg.photonCalibrationTolerance
    error(['HETERO photon/pigment calibration %.8g differs from expected ' ...
           '%.8g beyond tolerance %.8g.'], ...
        alpha,cfg.expectedPhotonPerPigment,cfg.photonCalibrationTolerance);
end

modelHETERO = addPhotonBiomassCoupling_pub( ...
    modelHETERO_uncoupled,cfg.photonRxn,cfg.HETERO.sinkRxn, ...
    cfg.HETERO.photonPerBiomass,'HETERO_PHOTON_BIOMASS_COUPLING_PUB');

assertCarbonBoundaryState_pub(modelAUTO,{cfg.co2Rxn},cfg,'AUTO_BASE');
assertCarbonBoundaryState_pub(modelMIXO,{cfg.co2Rxn},cfg,'MIXO_BASE');
assertCarbonBoundaryState_pub(modelHETERO,{},cfg,'HETERO_BASE');

assertBiomassOutputState_pub(modelAUTO,cfg.AUTO.sinkRxn,cfg,'AUTO');
assertBiomassOutputState_pub(modelMIXO,cfg.MIXO.sinkRxn,cfg,'MIXO');
assertBiomassOutputState_pub(modelHETERO,cfg.HETERO.sinkRxn,cfg,'HETERO');

basalMediumAudit = buildBasalMediumAudit_pub( ...
    baseModel,modelAUTO,modelMIXO,modelHETERO,cfg);
conditionAudit = buildConditionAudit_pub( ...
    modelAUTO,modelMIXO,modelHETERO,cfg);

autoCapacity = capacityOne_pub( ...
    modelAUTO,cfg.AUTO.sinkRxn,'AUTO','CO2',cfg.co2Rxn,1,cfg);

n = height(T);
mixoCapacity = repmat(emptyFixedResult_pub(),n,1);
heteroCapacity = repmat(emptyFixedResult_pub(),n,1);

for i=1:n
    if strlength(resolvedExchange(i))==0
        mixoCapacity(i) = failedFixedResult_pub('MIXO',substrateName(i),'MISSING_EXCHANGE');
        heteroCapacity(i) = failedFixedResult_pub('HETERO',substrateName(i),'MISSING_EXCHANGE');
        continue;
    end

    r = char(resolvedExchange(i));

    m = modelMIXO;
    m = openUptake_pub(m,r,cfg.screenCarbon/carbonAtoms(i),false);
    assertCarbonBoundaryState_pub(m,{cfg.co2Rxn,r},cfg,'MIXO_CAPACITY');
    mixoCapacity(i) = capacityOne_pub( ...
        m,cfg.MIXO.sinkRxn,'MIXO',substrateName(i),r,carbonAtoms(i),cfg);

    h = modelHETERO;
    h = openUptake_pub(h,r,cfg.screenCarbon/carbonAtoms(i),false);
    assertCarbonBoundaryState_pub(h,{r},cfg,'HETERO_CAPACITY');
    heteroCapacity(i) = capacityOne_pub( ...
        h,cfg.HETERO.sinkRxn,'HETERO',substrateName(i),r,carbonAtoms(i),cfg);
end

capacityResults = buildCapacityTable_pub( ...
    autoCapacity,mixoCapacity,heteroCapacity,substrateName,resolvedExchange, ...
    carbonAtoms,expectedGrowth,expGrowthText);

autoFixed = fixedMuCarbonAnalysis_pub( ...
    modelAUTO,cfg.AUTO.sinkRxn,'AUTO','CO2',cfg.co2Rxn,1,cfg);

mixoFixed = repmat(emptyFixedResult_pub(),n,1);
heteroFixed = repmat(emptyFixedResult_pub(),n,1);

for i=1:n
    if strlength(resolvedExchange(i))==0
        mixoFixed(i) = failedFixedResult_pub('MIXO',substrateName(i),'MISSING_EXCHANGE');
        heteroFixed(i) = failedFixedResult_pub('HETERO',substrateName(i),'MISSING_EXCHANGE');
        continue;
    end

    r = char(resolvedExchange(i));
    qOpen = cfg.validationMaxCarbon/carbonAtoms(i);

    m = modelMIXO;
    m = openUptake_pub(m,r,qOpen,false);
    assertCarbonBoundaryState_pub(m,{cfg.co2Rxn,r},cfg,'MIXO_FIXED_MU');
    mixoFixed(i) = fixedMuCarbonAnalysis_pub( ...
        m,cfg.MIXO.sinkRxn,'MIXO',substrateName(i),r,carbonAtoms(i),cfg);

    h = modelHETERO;
    h = openUptake_pub(h,r,qOpen,false);
    assertCarbonBoundaryState_pub(h,{r},cfg,'HETERO_FIXED_MU');
    heteroFixed(i) = fixedMuCarbonAnalysis_pub( ...
        h,cfg.HETERO.sinkRxn,'HETERO',substrateName(i),r,carbonAtoms(i),cfg);
end

fixedMuResults = buildFixedMuTable_pub( ...
    autoFixed,mixoFixed,heteroFixed,substrateName,resolvedExchange,carbonAtoms);

carbonFVA = buildCarbonFVATable_pub( ...
    autoFixed,mixoFixed,heteroFixed,substrateName,resolvedExchange);

bofProduction = buildBOFProduction_pub(cfg);
bofSourceAudit = buildBOFSourceAudit_pub( ...
    cfg,workbookMeanAudit,experimentalBofAudit);

summaryResults = buildPublicationSummary_pub( ...
    substrateName,substrateClass,expGrowthText,resolvedExchange,carbonAtoms, ...
    duplicateExchange,expectedGrowth,autoCapacity,autoFixed, ...
    mixoCapacity,heteroCapacity,mixoFixed,heteroFixed,cfg);

publicationComparison = buildPublicationComparison_pub( ...
    substrateName,substrateClass,expGrowthText,resolvedExchange,carbonAtoms, ...
    autoCapacity,autoFixed,mixoCapacity,heteroCapacity,heteroFixed, ...
    expectedGrowth,cfg);

suppSubstrates = buildSupplementarySubstrateTable_pub( ...
    substrateName,substrateClass,expGrowthText,resolvedExchange,carbonAtoms, ...
    autoCapacity,autoFixed,mixoCapacity,heteroCapacity,heteroFixed, ...
    expectedGrowth,cfg);

minimalSupplementTable = buildMinimalSupplementTable_pub(suppSubstrates);

longResults = fixedMuResults;

Setting = [ ...
    "Reference fixed mu"
    "Reference mu provenance"
    "Reference mu measurement"
    "AUTO BOF"
    "MIXO BOF"
    "HETERO BOF"
    "NGAM"
    "GAM"
    "Capacity carbon supply"
    "Minimum-carbon opening cap"
    "Carbon FVA tolerance"
    "MIXO light treatment"
    "HETERO light treatment"
    "AUTO/MIXO O2 dummy"
    "HETERO O2 dummy"
];

Value = [ ...
    sprintf('%.6f h^-1',cfg.referenceMu_h)
    sprintf('%.2f d^-1 / 24; %s',cfg.referenceMu_d,cfg.referenceMuCondition)
    string(cfg.referenceMuMethod)
    "TOTAL_BIOMASS_AUTO_MEAN from TOTAL_BIOMASS.xlsx"
    "Same experimental modular BOF as publication validation"
    "Same experimental modular BOF as publication validation"
    sprintf('%.1f mmol ATP gDW^-1 h^-1',cfg.NGAM)
    sprintf('%.0f mmol ATP gDW^-1 biomass',cfg.GAM)
    sprintf('%.0f C-mmol gDW^-1 h^-1',cfg.screenCarbon)
    sprintf('%.0f C-mmol gDW^-1 h^-1',cfg.validationMaxCarbon)
    sprintf('%.0f%% above minimum total carbon',100*cfg.fvaCarbonTolerance)
    "Finite model light constraint selected from feasibility analysis"
    sprintf('Pigment-linked; photon/pigment = %.4f',cfg.HETERO.photonPerPigment)
    "0..1000"
    "0..0"
];

Note = [ ...
    "Standardized cross-condition biomass flux"
    "Measured HETERO KIN2 mixed-sugar apparent growth rate"
    "Previous cultivation-validation dataset"
    "AUTO composition source"
    "Experimental measured 90% retained unnormalized; pigment separately accounted"
    "Experimental measured 90% retained unnormalized; pigment separately accounted"
    "Fixed LB = UB"
    "Included in each BOF"
    "Sole-substrate capacity comparison"
    "Non-binding cap used only while minimizing required carbon"
    "Same 5% tolerance principle as prior targeted FVA"
    "Exact numerical light flux retained for computational reproducibility only"
    "Same structural coupling used in previous validation"
    "Required for light-state oxygen balancing"
    ""
];

settings = table(Setting,Value,Note);

if isfile(cfg.resultsFile)
    delete(cfg.resultsFile);
end

sheetName = 'SOLE_SUBSTRATE_VALIDATION';

note1 = sprintf(['AUTO uses CO2 as the only carbon source. Maximum growth = %.4f h^-1; ' ...
    'at growth = %.3f h^-1, minimum CO2 demand = %.4f C-mmol gDW^-1 h^-1.'], ...
    autoCapacity.MaxMu_h,cfg.referenceMu_h,autoFixed.MinimumTotalCarbon);

note2 = sprintf(['A common reference growth rate of %.3f h^-1 was used for the ' ...
    'minimum-carbon and flux-variability (FVA) comparisons (%s).'], ...
    cfg.referenceMu_h,cfg.referenceMuMethod);

note3 = sprintf(['HETERO uses one organic substrate at a time; MIXO combines the same ' ...
    'substrate with CO2. Capacity-screen carbon supply = %.0f C-mmol gDW^-1 h^-1.'], ...
    cfg.screenCarbon);

note4 = ['Percentages are model-internal comparisons: HETERO values are compared with ' ...
    'D-glucose and MIXO growth is compared with HETERO. Published values are shown as ' ...
    'independent experimental evidence, not as prediction-error measurements.'];

writecell({"Sole-substrate validation across trophic states"}, ...
    cfg.resultsFile,'Sheet',sheetName,'Range','A1');
writecell({note1},cfg.resultsFile,'Sheet',sheetName,'Range','A2');
writecell({note2},cfg.resultsFile,'Sheet',sheetName,'Range','A3');
writecell({note3},cfg.resultsFile,'Sheet',sheetName,'Range','A4');
writecell({note4},cfg.resultsFile,'Sheet',sheetName,'Range','A5');

publicHeaders = { ...
    'Substrate', ...
    'Literature growth', ...
    'Published quantitative evidence', ...
    'DOI', ...
    'HETERO maximum growth rate (h^-1)', ...
    'HETERO growth vs glucose (%)', ...
    'HETERO carbon demand at growth 0.060 h^-1 (C-mmol gDW^-1 h^-1)', ...
    'HETERO carbon demand vs glucose (%)', ...
    'HETERO substrate uptake range at growth 0.060 h^-1 (mmol gDW^-1 h^-1)', ...
    'MIXO maximum growth rate (h^-1)', ...
    'MIXO growth increase vs HETERO (%)', ...
    'CO2 share of MIXO external carbon (%)', ...
    'Model vs literature' ...
    };
writecell(publicHeaders,cfg.resultsFile,'Sheet',sheetName,'Range','A7');
writecell(table2cell(minimalSupplementTable),cfg.resultsFile, ...
    'Sheet',sheetName,'Range','A8');

fprintf('Substrate optimisation ............ DONE\n');
fprintf('Created: 2_Galdieria_publication_substrate_optimisation.xlsx\n');

end


function [m,cfg,auditT] = implementExperimentalValidationBOFs_pub(m,cfg)

    [m,pigH] = addExperimentalPigmentPool_pub(m,'HETERO',cfg);
    [m,pigM] = addExperimentalPigmentPool_pub(m,'MIXO',cfg);

    [m,bofH,aH] = createExperimentalBOFHybrid_pub( ...
        m,'HETERO',cfg.exp.HETERO,cfg.reference.HETERO.rnaFraction, ...
        cfg.reference.HETERO.pigmentMass,pigH.poolMet,pigH.rxn,cfg);

    [m,bofM,aM] = createExperimentalBOFHybrid_pub( ...
        m,'MIXO',cfg.exp.MIXO,cfg.reference.MIXO.rnaFraction, ...
        cfg.reference.MIXO.pigmentMass,pigM.poolMet,pigM.rxn,cfg);

    cfg.HETERO = bofH;
    cfg.MIXO = bofM;

    auditT = [aH;aM];
end

function [m,info] = addExperimentalPigmentPool_pub(m,mode,cfg)
    % Detailed pigment composition is read from TOTAL_BIOMASS.xlsx.
    mode = upper(string(mode));
    pig = cfg.resolvedPigmentMets;

    if mode=="HETERO"
        poolMet = 'm_S2_PIGMENT_TOTAL_POOL_EXP_HETERO_PUB[c]';
        rxn = 'S2_PIGMENT_TOTAL_POOL_EXP_HETERO_PUB_PRODUCER_bm';
        sheetName = 'Reaction List Heto';
        workbookRxn = 'PIGMENT_TOTAL_POOL_BIO_HETEROTROPH_GLUCOSE_NOFRAC';
    elseif mode=="MIXO"
        poolMet = 'm_S2_PIGMENT_TOTAL_POOL_EXP_MIXO_PUB[c]';
        rxn = 'S2_PIGMENT_TOTAL_POOL_EXP_MIXO_PUB_PRODUCER_bm';
        sheetName = 'Reaction List Mixo';
        workbookRxn = 'PIGMENT_TOTAL_POOL_BIO_MIXOTROPH_NOFRAC';
    else
        error('Unsupported experimental pigment mode: %s',mode);
    end

    formula = readWorkbookReactionFormula_pub( ...
        cfg.biomassFile,sheetName,workbookRxn);

    cZea = coefficientForWorkbookMet_pub(formula,'m_CPD1F-130_Thyl[t]');
    try
        cChl = coefficientForWorkbookMet_pub(formula,'m_CHLOROPHYLL-A_Plas[p]');
    catch
        cChl = coefficientForWorkbookMet_pub(formula,'m_CHLOROPHYLL-A_Thyl[t]');
    end
    cBeta = coefficientForWorkbookMet_pub(formula,'m_CPD1F-129_Thyl[t]');
    cPCB = coefficientForWorkbookMet_pub(formula,'m_3Z-PHYCOCYANOBILIN_Plas[p]');

    coeff = [cPCB;cChl;cBeta;cZea]; % PCB, Chl-a, beta-carotene, zeaxanthin
    if any(~isfinite(coeff)) || sum(coeff)<=0
        error('Invalid pigment composition for %s in TOTAL_BIOMASS.xlsx.',mode);
    end
    coeff = coeff/sum(coeff);
    source = "TOTAL_BIOMASS.xlsx / " + string(workbookRxn);

    if ismember(rxn,m.rxns)
        m = removeRxns(m,{rxn});
    end
    if ~ismember(poolMet,m.mets)
        m = addMetabolite(m,poolMet, ...
            'metName',sprintf('Experimental %s pigment pool',mode), ...
            'metFormula','','Charge',0);
    end

    m = addReaction(m,rxn, ...
        'reactionName',sprintf('Experimental %s pigment pool from TOTAL_BIOMASS.xlsx',mode), ...
        'metaboliteList',{pig.pcb;pig.chla;pig.beta;pig.zea;poolMet}, ...
        'stoichCoeffList',[-coeff;1], ...
        'lowerBound',0,'upperBound',1000, ...
        'subSystem','Biomass pigment pool');
    m = padConstraintColumns_pub(m);

    info.poolMet = poolMet;
    info.rxn = rxn;
    info.coeff = coeff;
    info.source = source;
end

function [m,bof,auditT] = createExperimentalBOFHybrid_pub( ...
    m,mode,expComp,rnaFraction,pigmentMass,pigmentPoolMet,pigmentRxn,cfg)

    mode = upper(string(mode));
    mets = resolveModeBOFMets_pub(m,mode);

    raw = [ ...
        expComp.protein
        expComp.carbohydrate
        expComp.lipid
        expComp.nucleic*rnaFraction
        expComp.nucleic*(1-rnaFraction)
        pigmentMass
    ];

    represented = sum(raw);
    if represented<=0
        error('Invalid experimental BOF raw mass for %s.',mode);
    end
    % Preserve experimental g/gDW fractions. Detailed pool profiles come
    % from the mode-specific reference reactions in TOTAL_BIOMASS.xlsx.
    coeff = raw;
    if abs(sum(raw(1:5))-0.9)>1e-8 || any(~isfinite(coeff)) || any(coeff<0) || represented>1+1e-8
        error('Invalid unnormalized experimental BOF for %s.',mode);
    end

    newMet = sprintf('m_S2_TOTAL_BIOMASS_EXP_%s_PUB[c]',mode);
    assembly = sprintf('S2_TOTAL_BIOMASS_EXP_%s_PUB_bm',mode);
    sink = sprintf('S2_TOTAL_BIOMASS_EXP_%s_PUB_SINK_bm_tx',mode);

    if ismember(assembly,m.rxns), m=removeRxns(m,{assembly}); end
    if ismember(sink,m.rxns), m=removeRxns(m,{sink}); end
    if ~ismember(newMet,m.mets)
        m=addMetabolite(m,newMet, ...
            'metName',sprintf('Experimental %s total biomass publication analysis',mode), ...
            'metFormula','','Charge',0);
    end

    m=addReaction(m,assembly, ...
        'reactionName',sprintf('Experimental %s BOF; same composition as publication validation',mode), ...
        'metaboliteList',{ ...
            mets.protein;mets.carb;mets.lipid;mets.rna;mets.dna;pigmentPoolMet; ...
            mets.atp;mets.water;mets.adp;mets.pi;mets.proton;newMet}, ...
        'stoichCoeffList',[ ...
            -coeff;-cfg.GAM;-cfg.GAM;cfg.GAM;cfg.GAM;cfg.GAM;1], ...
        'lowerBound',0,'upperBound',1000, ...
        'subSystem','Biomass synthesis');
    m=padConstraintColumns_pub(m);

    m=addReaction(m,sink, ...
        'reactionName',sprintf('Experimental %s biomass sink',mode), ...
        'metaboliteList',{newMet}, ...
        'stoichCoeffList',-1, ...
        'lowerBound',0,'upperBound',1000, ...
        'subSystem','Biomass sink');
    m=padConstraintColumns_pub(m);

    bof.mode=char(mode);
    bof.assemblyRxn=assembly;
    bof.sinkRxn=sink;
    bof.biomassMet=newMet;
    bof.pigmentRxn=pigmentRxn;
    bof.pigmentMet=pigmentPoolMet;
    bof.workbookReaction='EXPERIMENTAL_PUBLICATION_VALIDATION';
    bof.workbookSheet='Experimental cultivation data';
    bof.workbookPigmentReaction='EXPERIMENTAL_PUBLICATION_VALIDATION';
    bof.sourceFormula='Built from exact raw experimental BOF inputs used in publication validation';
    bof.pigmentSourceFormula='Condition-specific pigment profile used in publication validation';
    bof.proteinCoeff=coeff(1);
    bof.carbohydrateCoeff=coeff(2);
    bof.lipidCoeff=coeff(3);
    bof.rnaCoeff=coeff(4);
    bof.dnaCoeff=coeff(5);
    bof.pigmentCoeff=coeff(6);
    bof.GAM=cfg.GAM;

    comp=["Protein";"Carbohydrate";"Lipid";"RNA";"DNA";"Pigment"];
    auditT=table( ...
        repmat(mode,6,1),comp,raw,coeff, ...
        repmat(represented,6,1), ...
        repmat(max(0,1-represented),6,1), ...
        repmat(cfg.GAM,6,1), ...
        'VariableNames',{'Mode','Component','Raw_g_gDW', ...
        'BOFCoefficient_g_per_gDW','RawModeledMassSum', ...
        'UnresolvedRawMass','GAM_mmol_per_gDW'});
end

function out = capacityOne_pub(m,biomassRxn,mode,substrate,subRxn,nC,cfg)
    mode = upper(string(mode));
    out = emptyFixedResult_pub();
    out.Mode = string(mode);
    out.Substrate = string(substrate);
    out.Exchange = string(subRxn);
    out.CarbonAtoms = nC;

    m = setObjective_pub(m,biomassRxn);
    sol = optimizeCbModel(m,'max');

    out.Feasible = isfield(sol,'stat') && sol.stat==1 && ...
        isfield(sol,'x') && ~isempty(sol.x);

    if ~out.Feasible
        out.Status = "INFEASIBLE";
        return;
    end

    out.MaxMu_h = max(0,sol.f);
    out.PredictedGrowth = out.MaxMu_h > cfg.growthTol;
    out.Status = "OPTIMAL";

    if mode=="AUTO"
        out.CO2Uptake = uptakeMagnitudeFromSolution_pub(m,cfg.co2Rxn,sol.x);
        out.SubstrateUptake = NaN;
        out.TotalCarbon = out.CO2Uptake;
    else
        out.SubstrateUptake = uptakeMagnitudeFromSolution_pub(m,subRxn,sol.x);
        if mode=="MIXO"
            out.CO2Uptake = uptakeMagnitudeFromSolution_pub(m,cfg.co2Rxn,sol.x);
        else
            out.CO2Uptake = 0;
        end
        out.TotalCarbon = nC*out.SubstrateUptake + out.CO2Uptake;
    end

    out.PhotonUptake = uptakeMagnitudeFromSolution_pub(m,cfg.photonRxn,sol.x);
    didx=find(strcmp(m.rxns,cfg.o2DummyRxn),1);
    if ~isempty(didx), out.O2DummyFlux=sol.x(didx); end
end

function out = fixedMuCarbonAnalysis_pub( ...
    m,biomassRxn,mode,substrate,subRxn,nC,cfg)
    mode = upper(string(mode));

    out = emptyFixedResult_pub();
    out.Mode = string(mode);
    out.Substrate = string(substrate);
    out.Exchange = string(subRxn);
    out.CarbonAtoms = nC;
    out.FixedMu_h = cfg.referenceMu_h;

    if mode=="AUTO"
        m=openUptake_pub(m,cfg.co2Rxn,cfg.validationMaxCarbon,true);
    elseif mode=="MIXO"
        m=openUptake_pub(m,cfg.co2Rxn,cfg.validationMaxCarbon,true);
    end

    bidx=find(strcmp(m.rxns,biomassRxn),1);
    if isempty(bidx), error('Biomass sink not found: %s',biomassRxn); end
    m.lb(bidx)=cfg.referenceMu_h;
    m.ub(bidx)=cfg.referenceMu_h;

    [sol,minTotal] = minimizeExternalCarbon_pub(m,mode,subRxn,nC,cfg);
    out.Feasible = ~isempty(sol);
    out.FixedMuFeasible = out.Feasible;

    if ~out.Feasible
        out.Status="FIXED_MU_INFEASIBLE";
        return;
    end

    out.Status="FIXED_MU_FEASIBLE";
    out.MinimumTotalCarbon=minTotal;
    out.SubstrateUptake=NaN;
    out.CO2Uptake=0;

    if mode=="AUTO"
        out.CO2Uptake=uptakeMagnitudeFromSolution_pub(m,cfg.co2Rxn,sol.x);
    else
        out.SubstrateUptake=uptakeMagnitudeFromSolution_pub(m,subRxn,sol.x);
        if mode=="MIXO"
            out.CO2Uptake=uptakeMagnitudeFromSolution_pub(m,cfg.co2Rxn,sol.x);
        end
    end

    if mode=="AUTO"
        out.TotalCarbon = max(0,out.CO2Uptake);
    else
        out.TotalCarbon = max(0,nC*out.SubstrateUptake) + max(0,out.CO2Uptake);
    end

    if mode=="MIXO"
        [solOrg,minOrg] = minimizeOrganicCarbon_pub(m,subRxn,nC);
        if ~isempty(solOrg)
            out.MinimumOrganicCarbon=minOrg;
        end
    elseif mode=="HETERO"
        out.MinimumOrganicCarbon=minTotal;
    else
        out.MinimumOrganicCarbon=0;
    end

    out.PhotonUptake=uptakeMagnitudeFromSolution_pub(m,cfg.photonRxn,sol.x);
    didx=find(strcmp(m.rxns,cfg.o2DummyRxn),1);
    if ~isempty(didx), out.O2DummyFlux=sol.x(didx); end

    orgC=0;
    if mode~="AUTO" && isfinite(out.SubstrateUptake)
        orgC=nC*out.SubstrateUptake;
    end
    co2C=max(0,out.CO2Uptake);
    denom=orgC+co2C;
    if denom>cfg.uptakeTol
        out.OrganicCarbonFraction=orgC/denom;
        out.CO2CarbonFraction=co2C/denom;
    end

    mFVA=addTotalCarbonUpperConstraint_pub( ...
        m,mode,subRxn,nC,minTotal*(1+cfg.fvaCarbonTolerance),cfg);

    if mode=="AUTO"
        [mn,mx]=manualFluxRange_pub(mFVA,cfg.co2Rxn);
        out.CO2FVAmin=uptakeRangeMin_pub(m,cfg.co2Rxn,mn,mx);
        out.CO2FVAmax=uptakeRangeMax_pub(m,cfg.co2Rxn,mn,mx);
    else
        [mn,mx]=manualFluxRange_pub(mFVA,subRxn);
        out.SubstrateFVAmin=uptakeRangeMin_pub(m,subRxn,mn,mx);
        out.SubstrateFVAmax=uptakeRangeMax_pub(m,subRxn,mn,mx);

        if mode=="MIXO"
            [mn,mx]=manualFluxRange_pub(mFVA,cfg.co2Rxn);
            out.CO2FVAmin=uptakeRangeMin_pub(m,cfg.co2Rxn,mn,mx);
            out.CO2FVAmax=uptakeRangeMax_pub(m,cfg.co2Rxn,mn,mx);
        else
            out.CO2FVAmin=0;
            out.CO2FVAmax=0;
        end
    end
end

function [sol,minC] = minimizeExternalCarbon_pub(m,mode,subRxn,nC,cfg)
    mode = upper(string(mode));
    m.c(:)=0;

    if mode=="AUTO"
        idx=find(strcmp(m.rxns,cfg.co2Rxn),1);
        m.c(idx)=-uptakeFluxSign_pub(m,cfg.co2Rxn);
        if uptakeFluxSign_pub(m,cfg.co2Rxn)<0, m.c(idx)=-1; else, m.c(idx)=1; end
    else
        idx=find(strcmp(m.rxns,subRxn),1);
        if uptakeFluxSign_pub(m,subRxn)<0, m.c(idx)=-nC; else, m.c(idx)=nC; end

        if mode=="MIXO"
            cidx=find(strcmp(m.rxns,cfg.co2Rxn),1);
            if uptakeFluxSign_pub(m,cfg.co2Rxn)<0, m.c(cidx)=-1; else, m.c(cidx)=1; end
        end
    end

    s=optimizeCbModel(m,'min');
    if ~isfield(s,'stat') || s.stat~=1 || ~isfield(s,'x') || isempty(s.x)
        sol=[]; minC=NaN; return;
    end

    sol=s;
    minC=max(0,s.f);
end

function [sol,minOrg] = minimizeOrganicCarbon_pub(m,subRxn,nC)
    m.c(:)=0;
    idx=find(strcmp(m.rxns,subRxn),1);
    if uptakeFluxSign_pub(m,subRxn)<0, m.c(idx)=-nC; else, m.c(idx)=nC; end
    s=optimizeCbModel(m,'min');
    if ~isfield(s,'stat') || s.stat~=1 || ~isfield(s,'x') || isempty(s.x)
        sol=[]; minOrg=NaN; return;
    end
    sol=s;
    minOrg=max(0,s.f);
end

function m = addTotalCarbonUpperConstraint_pub(m,mode,subRxn,nC,maxC,cfg)
    mode = upper(string(mode));
    m=ensureConstraintFields_pub(m);
    row=sparse(1,numel(m.rxns));

    if mode=="AUTO"
        idx=find(strcmp(m.rxns,cfg.co2Rxn),1);
        row(idx)=uptakeFluxSign_pub(m,cfg.co2Rxn);
    else
        idx=find(strcmp(m.rxns,subRxn),1);
        row(idx)=uptakeFluxSign_pub(m,subRxn)*nC;
        if mode=="MIXO"
            cidx=find(strcmp(m.rxns,cfg.co2Rxn),1);
            row(cidx)=uptakeFluxSign_pub(m,cfg.co2Rxn);
        end
    end

    m.C(end+1,:)=row;
    m.d(end+1,1)=maxC;
    m.dsense(end+1,1)='L';
    m.ctrs{end+1,1}=sprintf('TOTAL_CARBON_LE_%.12g',maxC);
end

function [mn,mx] = manualFluxRange_pub(m,rxn)
    idx=find(strcmp(m.rxns,rxn),1);
    if isempty(idx), mn=NaN; mx=NaN; return; end

    a=m; a.c(:)=0; a.c(idx)=1;
    s=optimizeCbModel(a,'min');
    if isfield(s,'stat') && s.stat==1, mn=s.f; else, mn=NaN; end

    b=m; b.c(:)=0; b.c(idx)=1;
    s=optimizeCbModel(b,'max');
    if isfield(s,'stat') && s.stat==1, mx=s.f; else, mx=NaN; end
end

function q = uptakeRangeMin_pub(m,rxn,mn,mx)
    if ~isfinite(mn) || ~isfinite(mx), q=NaN; return; end
    if uptakeFluxSign_pub(m,rxn)<0
        q=max(0,-mx);
    else
        q=max(0,mn);
    end
end

function q = uptakeRangeMax_pub(m,rxn,mn,mx)
    if ~isfinite(mn) || ~isfinite(mx), q=NaN; return; end
    if uptakeFluxSign_pub(m,rxn)<0
        q=max(0,-mn);
    else
        q=max(0,mx);
    end
end

function r = emptyFixedResult_pub()
    r=struct( ...
        'Mode',"",'Substrate',"",'Exchange',"",'CarbonAtoms',NaN, ...
        'Status',"",'Feasible',false,'PredictedGrowth',false, ...
        'MaxMu_h',NaN,'FixedMu_h',NaN,'FixedMuFeasible',false, ...
        'MinimumTotalCarbon',NaN,'MinimumOrganicCarbon',NaN, ...
        'SubstrateUptake',NaN,'CO2Uptake',NaN,'TotalCarbon',NaN, ...
        'OrganicCarbonFraction',NaN,'CO2CarbonFraction',NaN, ...
        'SubstrateFVAmin',NaN,'SubstrateFVAmax',NaN, ...
        'CO2FVAmin',NaN,'CO2FVAmax',NaN, ...
        'PhotonUptake',NaN,'O2DummyFlux',NaN);
end

function r = failedFixedResult_pub(mode,substrate,status)
    r=emptyFixedResult_pub();
    r.Mode=string(mode);
    r.Substrate=string(substrate);
    r.Status=string(status);
end

function T = buildCapacityTable_pub( ...
    auto,mixo,hetero,substrateName,resolvedExchange,carbonAtoms,expectedGrowth,expText)

    n=numel(substrateName);
    Mode=["AUTO";repmat("MIXO",n,1);repmat("HETERO",n,1)];
    Substrate=["CO2";substrateName;substrateName];
    Exchange=[string(auto.Exchange);resolvedExchange;resolvedExchange];
    Carbon_atoms=[1;carbonAtoms;carbonAtoms];
    Carbon_supply_Cmmol=[6;repmat(6,n,1);repmat(6,n,1)];
    Max_mu_h=[auto.MaxMu_h;[mixo.MaxMu_h]';[hetero.MaxMu_h]'];
    Feasible=[auto.Feasible;[mixo.Feasible]';[hetero.Feasible]'];
    Predicted_growth=[auto.PredictedGrowth;[mixo.PredictedGrowth]';[hetero.PredictedGrowth]'];
    Organic_uptake=[NaN;[mixo.SubstrateUptake]';[hetero.SubstrateUptake]'];
    CO2_uptake=[auto.CO2Uptake;[mixo.CO2Uptake]';[hetero.CO2Uptake]'];

    Experimental_growth=["NOT_SCORED_AUTO";expText;expText];
    Expected_growth=[NaN;expectedGrowth;expectedGrowth];
    Agreement=strings(1+2*n,1);

    Agreement(1)="AUTO_REFERENCE";
    for i=1:n
        Agreement(1+i)=agreementText_pub(expectedGrowth(i),mixo(i).PredictedGrowth);
        Agreement(1+n+i)=agreementText_pub(expectedGrowth(i),hetero(i).PredictedGrowth);
    end

    T=table(Mode,Substrate,Exchange,Carbon_atoms,Carbon_supply_Cmmol, ...
        Max_mu_h,Feasible,Predicted_growth,Organic_uptake,CO2_uptake, ...
        Experimental_growth,Expected_growth,Agreement);
end

function T = buildFixedMuTable_pub( ...
    auto,mixo,hetero,substrateName,resolvedExchange,carbonAtoms)

    n=numel(substrateName);
    Mode=["AUTO";repmat("MIXO",n,1);repmat("HETERO",n,1)];
    Substrate=["CO2";substrateName;substrateName];
    Exchange=[string(auto.Exchange);resolvedExchange;resolvedExchange];
    Carbon_atoms=[1;carbonAtoms;carbonAtoms];

    Fixed_mu_h=[auto.FixedMu_h;[mixo.FixedMu_h]';[hetero.FixedMu_h]'];
    Feasible=[auto.FixedMuFeasible;[mixo.FixedMuFeasible]';[hetero.FixedMuFeasible]'];
    Minimum_total_C=[auto.MinimumTotalCarbon;[mixo.MinimumTotalCarbon]';[hetero.MinimumTotalCarbon]'];
    Minimum_organic_C=[auto.MinimumOrganicCarbon;[mixo.MinimumOrganicCarbon]';[hetero.MinimumOrganicCarbon]'];
    Organic_uptake_minC=[auto.SubstrateUptake;[mixo.SubstrateUptake]';[hetero.SubstrateUptake]'];
    CO2_uptake_minC=[auto.CO2Uptake;[mixo.CO2Uptake]';[hetero.CO2Uptake]'];
    Organic_C_fraction=[auto.OrganicCarbonFraction;[mixo.OrganicCarbonFraction]';[hetero.OrganicCarbonFraction]'];
    CO2_C_fraction=[auto.CO2CarbonFraction;[mixo.CO2CarbonFraction]';[hetero.CO2CarbonFraction]'];
    Photon_uptake=[auto.PhotonUptake;[mixo.PhotonUptake]';[hetero.PhotonUptake]'];
    Status=[string(auto.Status);string({mixo.Status})';string({hetero.Status})'];

    T=table(Mode,Substrate,Exchange,Carbon_atoms,Fixed_mu_h,Feasible, ...
        Minimum_total_C,Minimum_organic_C,Organic_uptake_minC,CO2_uptake_minC, ...
        Organic_C_fraction,CO2_C_fraction,Photon_uptake,Status);
end

function T = buildCarbonFVATable_pub(auto,mixo,hetero,substrateName,resolvedExchange)
    n=numel(substrateName);
    Mode=["AUTO";repmat("MIXO",n,1);repmat("HETERO",n,1)];
    Substrate=["CO2";substrateName;substrateName];
    Exchange=[string(auto.Exchange);resolvedExchange;resolvedExchange];

    Organic_uptake_min=[NaN;[mixo.SubstrateFVAmin]';[hetero.SubstrateFVAmin]'];
    Organic_uptake_max=[NaN;[mixo.SubstrateFVAmax]';[hetero.SubstrateFVAmax]'];
    CO2_uptake_min=[auto.CO2FVAmin;[mixo.CO2FVAmin]';[hetero.CO2FVAmin]'];
    CO2_uptake_max=[auto.CO2FVAmax;[mixo.CO2FVAmax]';[hetero.CO2FVAmax]'];
    Fixed_mu_h=[auto.FixedMu_h;[mixo.FixedMu_h]';[hetero.FixedMu_h]'];
    Minimum_total_C=[auto.MinimumTotalCarbon;[mixo.MinimumTotalCarbon]';[hetero.MinimumTotalCarbon]'];

    T=table(Mode,Substrate,Exchange,Fixed_mu_h,Minimum_total_C, ...
        Organic_uptake_min,Organic_uptake_max,CO2_uptake_min,CO2_uptake_max);
end

function T = buildBOFProduction_pub(cfg)
    modes=["AUTO";"MIXO";"HETERO"];
    comps=["Protein";"Carbohydrate";"Lipid";"RNA";"DNA";"Pigment"];

    Mode=strings(18,1);
    Component=strings(18,1);
    BOF_coefficient=nan(18,1);
    Production_at_mu_0_060=nan(18,1);
    BOF_source=strings(18,1);

    row=0;
    for i=1:3
        if modes(i)=="AUTO"
            b=cfg.AUTO;
            src="TOTAL_BIOMASS_AUTO_MEAN";
        elseif modes(i)=="MIXO"
            b=cfg.MIXO;
            src="Experimental MIXO BOF from publication validation";
        else
            b=cfg.HETERO;
            src="Experimental HETERO BOF from publication validation";
        end

        c=[b.proteinCoeff;b.carbohydrateCoeff;b.lipidCoeff; ...
           b.rnaCoeff;b.dnaCoeff;b.pigmentCoeff];

        for j=1:6
            row=row+1;
            Mode(row)=modes(i);
            Component(row)=comps(j);
            BOF_coefficient(row)=c(j);
            Production_at_mu_0_060(row)=c(j)*cfg.referenceMu_h;
            BOF_source(row)=src;
        end
    end

    T=table(Mode,Component,BOF_coefficient,Production_at_mu_0_060,BOF_source);
end

function T = buildBOFSourceAudit_pub(cfg,meanAudit,expAudit)
    autoRow=meanAudit(strcmpi(string(meanAudit.Mode),'AUTO'),:);

    Source=strings(height(autoRow)+height(expAudit),1);
    Mode=strings(height(autoRow)+height(expAudit),1);
    Detail=strings(height(autoRow)+height(expAudit),1);

    Mode(1)=string(autoRow.Mode(1));
    Source(1)="TOTAL_BIOMASS.xlsx";
    Detail(1)="TOTAL_BIOMASS_AUTO_MEAN + autotrophic NOFRAC pigment composition";

    for i=1:height(expAudit)
        Mode(1+i)=string(expAudit.Mode(i));
        Source(1+i)="1_Galdieria_publication_growth_validation";
        Detail(1+i)=sprintf('%s measured=%.8g BOF=%.8g', ...
            char(expAudit.Component(i)),expAudit.Raw_g_gDW(i), ...
            expAudit.BOFCoefficient_g_per_gDW(i));
    end

    T=table(Mode,Source,Detail);
end

function T = buildPublicationSummary_pub( ...
    substrateName,substrateClass,expGrowthText,resolvedExchange,carbonAtoms, ...
    duplicateExchange,expectedGrowth,autoCapacity,autoFixed, ...
    mixoCapacity,heteroCapacity,mixoFixed,heteroFixed,cfg)

    n=numel(substrateName);

    AUTO_reference_mu_max=repmat(autoCapacity.MaxMu_h,n,1);
    AUTO_reference_min_CO2_at_mu=repmat(autoFixed.MinimumTotalCarbon,n,1);

    HETERO_mu_max=[heteroCapacity.MaxMu_h]';
    HETERO_growth=[heteroCapacity.PredictedGrowth]';
    HETERO_fixed_mu_feasible=[heteroFixed.FixedMuFeasible]';
    HETERO_min_C_at_mu=[heteroFixed.MinimumTotalCarbon]';
    HETERO_substrate_FVA_min=[heteroFixed.SubstrateFVAmin]';
    HETERO_substrate_FVA_max=[heteroFixed.SubstrateFVAmax]';

    MIXO_mu_max=[mixoCapacity.MaxMu_h]';
    MIXO_growth=[mixoCapacity.PredictedGrowth]';
    MIXO_fixed_mu_feasible=[mixoFixed.FixedMuFeasible]';
    MIXO_min_total_C_at_mu=[mixoFixed.MinimumTotalCarbon]';
    MIXO_min_organic_C_at_mu=[mixoFixed.MinimumOrganicCarbon]';
    MIXO_CO2_fraction_minC=[mixoFixed.CO2CarbonFraction]';
    MIXO_organic_FVA_min=[mixoFixed.SubstrateFVAmin]';
    MIXO_organic_FVA_max=[mixoFixed.SubstrateFVAmax]';
    MIXO_CO2_FVA_min=[mixoFixed.CO2FVAmin]';
    MIXO_CO2_FVA_max=[mixoFixed.CO2FVAmax]';

    HETERO_agreement=strings(n,1);
    MIXO_agreement=strings(n,1);
    for i=1:n
        HETERO_agreement(i)=agreementText_pub(expectedGrowth(i),HETERO_growth(i));
        MIXO_agreement(i)=agreementText_pub(expectedGrowth(i),MIXO_growth(i));
    end

    Reference_mu_h=repmat(cfg.referenceMu_h,n,1);

    T=table(substrateName,substrateClass,expGrowthText,resolvedExchange, ...
        carbonAtoms,duplicateExchange,Reference_mu_h, ...
        AUTO_reference_mu_max,AUTO_reference_min_CO2_at_mu, ...
        HETERO_mu_max,HETERO_growth,HETERO_agreement, ...
        HETERO_fixed_mu_feasible,HETERO_min_C_at_mu, ...
        HETERO_substrate_FVA_min,HETERO_substrate_FVA_max, ...
        MIXO_mu_max,MIXO_growth,MIXO_agreement,MIXO_fixed_mu_feasible, ...
        MIXO_min_total_C_at_mu,MIXO_min_organic_C_at_mu,MIXO_CO2_fraction_minC, ...
        MIXO_organic_FVA_min,MIXO_organic_FVA_max,MIXO_CO2_FVA_min,MIXO_CO2_FVA_max, ...
        'VariableNames',{ ...
        'Substrate','Class','Growth_1995','Exchange','Carbon_atoms', ...
        'Duplicate_exchange_mapping','Reference_mu_h', ...
        'AUTO_reference_max_mu_h','AUTO_min_CO2_Cmmol_at_reference_mu', ...
        'HETERO_max_mu_h','HETERO_growth','HETERO_growth_agreement_1995', ...
        'HETERO_reference_mu_feasible','HETERO_min_Cmmol_at_reference_mu', ...
        'HETERO_substrate_FVA_min_mmol','HETERO_substrate_FVA_max_mmol', ...
        'MIXO_max_mu_h','MIXO_growth','MIXO_growth_agreement_1995', ...
        'MIXO_reference_mu_feasible','MIXO_min_total_Cmmol_at_reference_mu', ...
        'MIXO_min_organic_Cmmol_at_reference_mu','MIXO_CO2_C_fraction_minC', ...
        'MIXO_organic_FVA_min_mmol','MIXO_organic_FVA_max_mmol', ...
        'MIXO_CO2_FVA_min_mmol','MIXO_CO2_FVA_max_mmol'});
end


function T = buildSupplementarySubstrateTable_pub( ...
    substrateName,substrateClass,expGrowthText,resolvedExchange,carbonAtoms, ...
    autoCapacity,autoFixed,mixoCapacity,heteroCapacity,heteroFixed, ...
    expectedGrowth,cfg)

    n = numel(substrateName);

    Substrate = string(substrateName);
    Class = string(substrateClass);
    Growth_1995 = string(expGrowthText);
    Exchange = string(resolvedExchange);
    Carbon_atoms = carbonAtoms;

    AUTO_mu_max = repmat(autoCapacity.MaxMu_h,n,1);
    AUTO_min_CO2_at_mu_0_060 = repmat(autoFixed.MinimumTotalCarbon,n,1);

    HETERO_mu_max = [heteroCapacity.MaxMu_h]';
    HETERO_growth_agreement = strings(n,1);
    for i=1:n
        HETERO_growth_agreement(i) = ...
            agreementText_pub(expectedGrowth(i),heteroCapacity(i).PredictedGrowth);
    end
    HETERO_min_C_at_mu_0_060 = [heteroFixed.MinimumTotalCarbon]';
    HETERO_FVA_min = [heteroFixed.SubstrateFVAmin]';
    HETERO_FVA_max = [heteroFixed.SubstrateFVAmax]';

    MIXO_mu_max = [mixoCapacity.MaxMu_h]';
    MIXO_growth_agreement = strings(n,1);
    for i=1:n
        MIXO_growth_agreement(i) = ...
            agreementText_pub(expectedGrowth(i),mixoCapacity(i).PredictedGrowth);
    end

    MIXO_organic_uptake = [mixoCapacity.SubstrateUptake]';
    MIXO_CO2_uptake = [mixoCapacity.CO2Uptake]';
    MIXO_organic_C = MIXO_organic_uptake .* carbonAtoms;

    totalC = MIXO_organic_C + MIXO_CO2_uptake;
    MIXO_CO2_fraction = nan(n,1);
    ok = isfinite(totalC) & totalC > cfg.uptakeTol;
    MIXO_CO2_fraction(ok) = MIXO_CO2_uptake(ok)./totalC(ok);

    MIXO_to_HETERO_mu_ratio = nan(n,1);
    okHet = isfinite(HETERO_mu_max) & HETERO_mu_max > cfg.growthTol;
    MIXO_to_HETERO_mu_ratio(okHet) = ...
        MIXO_mu_max(okHet)./HETERO_mu_max(okHet);

    Delta_mu_MIXO_minus_HETERO = MIXO_mu_max - HETERO_mu_max;

    Literature_parent_substrate = strings(n,1);
    Experimental_support = strings(n,1);
    Evidence_level = strings(n,1);
    DOI = strings(n,1);
    Literature_note = strings(n,1);

    for i=1:n
        [Literature_parent_substrate(i),Experimental_support(i), ...
            Evidence_level(i),DOI(i),Literature_note(i)] = ...
            literatureSupport_pub(Substrate(i),Exchange(i));
    end

    T = table( ...
        Substrate,Class,Growth_1995,Exchange,Carbon_atoms, ...
        Literature_parent_substrate,Experimental_support,Evidence_level,DOI,Literature_note, ...
        AUTO_mu_max,AUTO_min_CO2_at_mu_0_060, ...
        HETERO_mu_max,HETERO_growth_agreement,HETERO_min_C_at_mu_0_060, ...
        HETERO_FVA_min,HETERO_FVA_max, ...
        MIXO_mu_max,MIXO_growth_agreement,MIXO_organic_uptake, ...
        MIXO_CO2_uptake,MIXO_CO2_fraction, ...
        MIXO_to_HETERO_mu_ratio,Delta_mu_MIXO_minus_HETERO);
end


function T = buildMinimalSupplementTable_pub(S)

    n = height(S);

    Substrate = string(S.Substrate);
    Reported_growth = string(S.Growth_1995);

    Published_evidence = strings(n,1);
    Evidence_DOI = strings(n,1);
    for i=1:n
        [Published_evidence(i),Evidence_DOI(i)] = ...
            publishedEvidence_pub(Substrate(i),string(S.Exchange(i)));
    end

    HETERO_mu_max_h_1 = S.HETERO_mu_max;
    HETERO_agreement = string(S.HETERO_growth_agreement);
    HETERO_min_C_Cmmol_gDW_h_at_mu_0_060 = S.HETERO_min_C_at_mu_0_060;

    HETERO_FVA_mmol_gDW_h = strings(n,1);
    for i=1:n
        a = S.HETERO_FVA_min(i);
        b = S.HETERO_FVA_max(i);
        if isfinite(a) && isfinite(b)
            HETERO_FVA_mmol_gDW_h(i) = sprintf('%.3f-%.3f',a,b);
        else
            HETERO_FVA_mmol_gDW_h(i) = "";
        end
    end

    MIXO_mu_max_h_1 = S.MIXO_mu_max;
    MIXO_agreement = string(S.MIXO_growth_agreement);
    MIXO_CO2_external_C_pct = 100*S.MIXO_CO2_fraction;

    g = find(strcmpi(strtrim(Substrate),"D-Glucose"),1);
    if isempty(g)
        error('D-Glucose row is required for publication percentage comparisons.');
    end

    glcMu = HETERO_mu_max_h_1(g);
    glcC  = HETERO_min_C_Cmmol_gDW_h_at_mu_0_060(g);

    HETERO_mu_vs_glucose_pct = nan(n,1);
    HETERO_C_demand_vs_glucose_pct = nan(n,1);
    MIXO_increase_vs_HETERO_pct = nan(n,1);

    for i=1:n
        if isfinite(HETERO_mu_max_h_1(i)) && isfinite(glcMu) && glcMu>0
            HETERO_mu_vs_glucose_pct(i) = ...
                100*(HETERO_mu_max_h_1(i)/glcMu - 1);
        end

        if isfinite(HETERO_min_C_Cmmol_gDW_h_at_mu_0_060(i)) && ...
                isfinite(glcC) && glcC>0
            HETERO_C_demand_vs_glucose_pct(i) = ...
                100*(HETERO_min_C_Cmmol_gDW_h_at_mu_0_060(i)/glcC - 1);
        end

        if isfinite(HETERO_mu_max_h_1(i)) && HETERO_mu_max_h_1(i)>0 && ...
                isfinite(MIXO_mu_max_h_1(i))
            MIXO_increase_vs_HETERO_pct(i) = ...
                100*(MIXO_mu_max_h_1(i)/HETERO_mu_max_h_1(i) - 1);
        end
    end

    Model_agreement = HETERO_agreement + " / " + MIXO_agreement;

    T = table( ...
        Substrate,Reported_growth,Published_evidence,Evidence_DOI, ...
        HETERO_mu_max_h_1,HETERO_mu_vs_glucose_pct, ...
        HETERO_min_C_Cmmol_gDW_h_at_mu_0_060, ...
        HETERO_C_demand_vs_glucose_pct,HETERO_FVA_mmol_gDW_h, ...
        MIXO_mu_max_h_1,MIXO_increase_vs_HETERO_pct, ...
        MIXO_CO2_external_C_pct,Model_agreement);
end


function [evidence,doi] = publishedEvidence_pub(substrate,exchange)

    s = lower(strtrim(string(substrate)));
    e = lower(strtrim(string(exchange)));

    evidence = "";
    doi = "";

    if contains(s,"starch")
        evidence = "Storage substrate; not scored as sole-carbon phenotype";
        return;
    end

    if strcmpi(substrate,"Sucrose")
        evidence = "Batch pure sugars: mu = 1.08-1.15 d^-1 (glucose/fructose/sucrose)";
        doi = "10.1002/bit.20417";
        return;
    end

    if strcmpi(substrate,"D-Glucose")
        evidence = "mu = 1.10 +/- 0.03 d^-1 (074W); t_d = 50 h (074W)";
        doi = "10.3389/fpls.2022.978246; 10.1080/09670269910001736322";
        return;
    end

    if strcmpi(substrate,"L-Glucose")
        evidence = "Qualitative parent/stereoisomer evidence; no robust substrate-specific mu retained";
        doi = "10.1093/oxfordjournals.pcp.a078803";
        return;
    end

    if strcmpi(substrate,"D-Mannose")
        evidence = "t_d = 50 h";
        doi = "10.1080/09670269910001736322";
        return;
    end

    if strcmpi(substrate,"D-Galactose") || contains(s,"galactose")
        evidence = "t_d = 50 h (parent D-galactose)";
        doi = "10.1080/09670269910001736322";
        return;
    end

    if strcmpi(substrate,"D-Fructose")
        evidence = "t_d = 57 h; independent batch mu = 1.08 d^-1";
        doi = "10.1080/09670269910001736322; 10.1002/bit.20417";
        return;
    end

    if strcmpi(substrate,"L-Sorbose")
        evidence = "t_d = 170 h";
        doi = "10.1080/09670269910001736322";
        return;
    end

    if strcmpi(substrate,"D-Fucose")
        evidence = "t_d = 840 h";
        doi = "10.1080/09670269910001736322";
        return;
    end

    if strcmpi(substrate,"L-Rhamnose")
        evidence = "t_d = 450 h (parent L-rhamnose)";
        doi = "10.1080/09670269910001736322";
        if contains(e,"beta")
            evidence = evidence + "; anomer-specific growth not resolved";
        end
        return;
    end

    if strcmpi(substrate,"D-Arabinose") || strcmpi(substrate,"L-Arabinose") || ...
            strcmpi(substrate,"D-Lyxose")
        evidence = "Pentose growth/pathway demonstrated; approx. 50 h doubling for tested pentoses except L-xylose";
        doi = "10.1104/pp.114.1.231";
        return;
    end

    if strcmpi(substrate,"CPD-15829") || strcmpi(substrate,"D-Ribose")
        evidence = "t_d = 450 h (parent D-ribose)";
        doi = "10.1080/09670269910001736322";
        return;
    end

    if strcmpi(substrate,"CPD-25028") || strcmpi(substrate,"D-Xylose")
        evidence = "t_d = 282 h; modern study: glucose specific growth rate was 51% higher than xylose";
        doi = "10.1080/09670269910001736322; 10.1016/j.algal.2022.102752";
        return;
    end

    if strcmpi(substrate,"D-Mannitol")
        evidence = "t_d = 60 h";
        doi = "10.1080/09670269910001736322";
        return;
    end

    if strcmpi(substrate,"D-Sorbitol")
        evidence = "t_d = 60 h; direct mixotrophic enhancement demonstrated";
        doi = "10.1080/09670269910001736322; 10.1111/nph.17359";
        return;
    end

    if contains(s,"dulcitol") || contains(s,"galactitol")
        evidence = "t_d = 120 h (dulcitol/galactitol)";
        doi = "10.1080/09670269910001736322";
        return;
    end

    if contains(s,"ribitol") || contains(s,"adonitol")
        evidence = "t_d = 130 h (adonitol/ribitol)";
        doi = "10.1080/09670269910001736322";
        return;
    end

    if strcmpi(substrate,"Xylitol")
        evidence = "t_d = 65 h";
        doi = "10.1080/09670269910001736322";
        return;
    end

    if strcmpi(substrate,"L-Arabitol") || strcmpi(substrate,"D-Arabitol")
        evidence = "t_d = 60 h";
        doi = "10.1080/09670269910001736322";
        return;
    end

    if strcmpi(substrate,"meso-Erythritol")
        evidence = "Direct growth in the 27-carbon-source screen; no robust quantitative mu retained";
        doi = "10.1093/oxfordjournals.pcp.a078803";
        return;
    end

    if strcmpi(substrate,"Glycerol")
        evidence = "mu = 1.08 +/- 0.03 d^-1; doubling time = 0.64 +/- 0.02 d";
        doi = "10.3389/fpls.2022.978246";
        return;
    end

    if strcmpi(substrate,"Acetate")
        evidence = "AA F=1.25: biomass productivity = 1.18 +/- 0.05 g L^-1 d^-1; utilization is concentration/regime dependent";
        doi = "10.1021/acssuschemeng.5c02338; 10.1016/j.algal.2022.102752";
        return;
    end

    if strcmpi(substrate,"Pyruvate")
        evidence = "No convincing direct sole-pyruvate growth-rate dataset identified";
        doi = "10.1104/pp.010553";
        return;
    end

    evidence = "Qualitative Galdieria growth support";
    doi = "10.1093/oxfordjournals.pcp.a078803";
end

function label = evidenceLabel_pub(level)

    x = upper(strtrim(string(level)));

    switch x
        case "DIRECT_QUANTITATIVE"
            label = "Direct quantitative";
        case "DIRECT_GROWTH"
            label = "Direct growth";
        case "DIRECT_PATHWAY"
            label = "Direct pathway";
        case "REGIME_DEPENDENT"
            label = "Regime dependent";
        case "PARENT_SUBSTRATE"
            label = "Parent-substrate evidence";
        case "INSUFFICIENT_DIRECT"
            label = "Insufficient direct growth evidence";
        case "NOT_SCORED_STORAGE"
            label = "Storage; not scored";
        otherwise
            label = x;
    end
end

function [parent,support,level,doi,note] = literatureSupport_pub(substrate,exchange)

    s = lower(strtrim(string(substrate)));
    e = lower(strtrim(string(exchange)));

    parent = string(substrate);
    support = "";
    level = "";
    doi = "";
    note = "";

    if contains(s,"starch")
        parent = "Floridean starch";
        support = "Storage substrate; not used as a scored sole-growth phenotype";
        level = "NOT_SCORED_STORAGE";
        doi = "";
        note = "Excluded from qualitative phenotype score.";
        return;
    end

    if strcmpi(substrate,"Sucrose")
        parent = "Sucrose";
        support = "Direct Galdieria growth; quantitative growth data available";
        level = "DIRECT_QUANTITATIVE";
        doi = "10.1002/bit.20417";
        return;
    end

    if strcmpi(substrate,"D-Glucose")
        parent = "D-Glucose";
        support = "Strong direct Galdieria growth evidence with quantitative data";
        level = "DIRECT_QUANTITATIVE";
        doi = "10.1080/09670269910001736322; 10.1002/bit.20417";
        return;
    end

    if strcmpi(substrate,"L-Glucose")
        parent = "Glucose";
        support = "Shared model exchange with D-glucose; literature support is not exchange-stereochemistry specific here";
        level = "PARENT_SUBSTRATE";
        doi = "10.1093/oxfordjournals.pcp.a078803";
        note = "D/L-glucose are not independently resolved by the shared model exchange.";
        return;
    end

    if strcmpi(substrate,"D-Mannose")
        parent = "D-Mannose";
        support = "Direct Galdieria growth and dedicated mannose-metabolism evidence";
        level = "DIRECT_QUANTITATIVE";
        doi = "10.1016/S0031-9422(97)00081-2; 10.1080/09670269910001736322";
        return;
    end

    if strcmpi(substrate,"D-Galactose")
        parent = "D-Galactose";
        support = "Direct Galdieria growth with quantitative doubling data; modern mixotrophic support";
        level = "DIRECT_QUANTITATIVE";
        doi = "10.1080/09670269910001736322; 10.1007/s10811-023-03012-0";
        return;
    end

    if contains(s,string(char(945))+"-d-galactose") || contains(s,"alpha-d-galactose")
        parent = "D-Galactose";
        support = "Parent-substrate growth evidence; anomer-specific utilization not independently resolved";
        level = "PARENT_SUBSTRATE";
        doi = "10.1080/09670269910001736322";
        note = "Literature generally reports D-galactose rather than the alpha anomer separately.";
        return;
    end

    if strcmpi(substrate,"D-Fructose")
        parent = "D-Fructose";
        support = "Direct Galdieria growth with quantitative data";
        level = "DIRECT_QUANTITATIVE";
        doi = "10.1002/bit.20417; 10.1080/09670269910001736322";
        return;
    end

    if strcmpi(substrate,"L-Sorbose")
        parent = "L-Sorbose";
        support = "Direct Galdieria growth with quantitative doubling data";
        level = "DIRECT_QUANTITATIVE";
        doi = "10.1080/09670269910001736322";
        return;
    end

    if strcmpi(substrate,"D-Fucose")
        parent = "D-Fucose";
        support = "Direct Galdieria growth, reported as slow; anomer not separately resolved";
        level = "PARENT_SUBSTRATE";
        doi = "10.1080/09670269910001736322";
        return;
    end

    if strcmpi(substrate,"L-Rhamnose")
        parent = "L-Rhamnose";
        support = "Direct Galdieria growth with quantitative doubling data";
        level = "DIRECT_QUANTITATIVE";
        doi = "10.1080/09670269910001736322";
        if contains(e,"beta")
            note = "Same parent-substrate evidence is used for the beta-model exchange; anomer-specific growth was not resolved.";
            level = "PARENT_SUBSTRATE";
        end
        return;
    end

    if strcmpi(substrate,"D-Arabinose")
        parent = "D-Arabinose";
        support = "Direct pentose-utilization and pathway evidence in Galdieria";
        level = "DIRECT_PATHWAY";
        doi = "10.1104/pp.114.1.231";
        return;
    end

    if strcmpi(substrate,"L-Arabinose")
        parent = "L-Arabinose";
        support = "Direct Galdieria growth screen plus pentose uptake/metabolism evidence";
        level = "DIRECT_GROWTH";
        doi = "10.1093/oxfordjournals.pcp.a078803; 10.1104/pp.114.1.231";
        return;
    end

    if strcmpi(substrate,"D-Lyxose")
        parent = "D-Lyxose";
        support = "Direct pentose metabolism/growth evidence in Galdieria";
        level = "DIRECT_PATHWAY";
        doi = "10.1104/pp.114.1.231";
        return;
    end

    if strcmpi(substrate,"CPD-15829")
        parent = "D-Ribose";
        support = "Parent-substrate evidence for D-ribose; alpha-ribopyranose exchange not independently tested";
        level = "PARENT_SUBSTRATE";
        doi = "10.1080/09670269910001736322; 10.1104/pp.114.1.231";
        note = "Model identifier CPD-15829 corresponds to an alpha-D-ribopyranose representation.";
        return;
    end

    if strcmpi(substrate,"D-Ribose")
        parent = "D-Ribose";
        support = "Direct Galdieria growth with quantitative data and pentose-pathway evidence";
        level = "DIRECT_QUANTITATIVE";
        doi = "10.1080/09670269910001736322; 10.1104/pp.114.1.231";
        return;
    end

    if strcmpi(substrate,"CPD-25028")
        parent = "D-Xylose";
        support = "Parent-substrate evidence for D-xylose; alpha-xylose exchange not independently tested";
        level = "PARENT_SUBSTRATE";
        doi = "10.1016/j.algal.2022.102752; 10.1080/09670269910001736322";
        note = "Model identifier CPD-25028 corresponds to an alpha-D-xylose representation.";
        return;
    end

    if strcmpi(substrate,"D-Xylose")
        parent = "D-Xylose";
        support = "Direct Galdieria growth with quantitative doubling data and modern hetero/mixotrophic validation";
        level = "DIRECT_QUANTITATIVE";
        doi = "10.1016/j.algal.2022.102752; 10.1080/09670269910001736322";
        return;
    end

    if strcmpi(substrate,"D-Mannitol")
        parent = "D-Mannitol";
        support = "Direct Galdieria growth with quantitative data and dedicated polyol-metabolism evidence";
        level = "DIRECT_QUANTITATIVE";
        doi = "10.1007/s004250050153; 10.1080/09670269910001736322";
        return;
    end

    if strcmpi(substrate,"D-Sorbitol")
        parent = "D-Sorbitol";
        support = "Strong direct Galdieria mixotrophic evidence and quantitative growth data";
        level = "DIRECT_QUANTITATIVE";
        doi = "10.1111/nph.17359; 10.1080/09670269910001736322";
        return;
    end

    if contains(s,"dulcitol") || contains(s,"galactitol")
        parent = "Galactitol (Dulcitol)";
        support = "Direct Galdieria growth with quantitative doubling data";
        level = "DIRECT_QUANTITATIVE";
        doi = "10.1080/09670269910001736322";
        note = "Galactitol and dulcitol are synonyms.";
        return;
    end

    if contains(s,"ribitol") || contains(s,"adonitol")
        parent = "Ribitol (Adonitol)";
        support = "Direct Galdieria growth with quantitative doubling data";
        level = "DIRECT_QUANTITATIVE";
        doi = "10.1080/09670269910001736322";
        note = "Ribitol and adonitol are synonyms.";
        return;
    end

    if strcmpi(substrate,"Xylitol")
        parent = "Xylitol";
        support = "Direct Galdieria growth with quantitative data and dedicated polyol-metabolism evidence";
        level = "DIRECT_QUANTITATIVE";
        doi = "10.1007/s004250050153; 10.1080/09670269910001736322";
        return;
    end

    if strcmpi(substrate,"L-Arabitol")
        parent = "L-Arabitol";
        support = "Direct Galdieria growth with quantitative data and polyol-metabolism evidence";
        level = "DIRECT_QUANTITATIVE";
        doi = "10.1007/s004250050153; 10.1080/09670269910001736322";
        return;
    end

    if strcmpi(substrate,"D-Arabitol")
        parent = "D-Arabitol";
        support = "Direct Galdieria growth with quantitative data and polyol-metabolism evidence";
        level = "DIRECT_QUANTITATIVE";
        doi = "10.1007/s004250050153; 10.1080/09670269910001736322";
        return;
    end

    if strcmpi(substrate,"meso-Erythritol")
        parent = "Erythritol";
        support = "Experimentally used Galdieria carbon source; direct quantitative support is weaker than for major sugars/polyols";
        level = "DIRECT_GROWTH";
        doi = "10.1093/oxfordjournals.pcp.a078803; 10.1101/076208";
        return;
    end

    if strcmpi(substrate,"Glycerol")
        parent = "Glycerol";
        support = "Strong modern quantitative Galdieria growth evidence";
        level = "DIRECT_QUANTITATIVE";
        doi = "10.3389/fpls.2022.978246; 10.1016/j.enzmictec.2005.05.010";
        return;
    end

    if strcmpi(substrate,"Acetate")
        parent = "Acetate / acetic acid";
        support = "Experimental Galdieria utilization supported, but strongly concentration- and regime-dependent";
        level = "REGIME_DEPENDENT";
        doi = "10.1021/acssuschemeng.5c02338; 10.1111/nph.17359; 10.1016/j.algal.2022.102752";
        note = "Interpret growth/no-growth only in the context of concentration and cultivation regime.";
        return;
    end

    if strcmpi(substrate,"Pyruvate")
        parent = "Pyruvate";
        support = "Galdieria pathway/transport context exists, but convincing direct sole-pyruvate growth data are insufficient";
        level = "INSUFFICIENT_DIRECT";
        doi = "10.1104/pp.010553";
        note = "Do not describe pyruvate as experimentally validated sole-substrate growth.";
        return;
    end

    parent = string(substrate);
    support = "No curated Galdieria literature mapping assigned";
    level = "UNMAPPED";
    doi = "";
    note = "Review before publication.";
end

function T = buildPublicationCases_pub(S,cfg)

    wanted = [ ...
        "D-Glucose"
        "D-Xylose"
        "meso-Erythritol"
        "Glycerol"
        "Acetate"
        "Pyruvate"
    ];

    idxOut = zeros(0,1);
    for i=1:numel(wanted)
        idx = find(strcmpi(strtrim(string(S.Substrate)),wanted(i)),1);
        if ~isempty(idx)
            idxOut(end+1,1) = idx; %#ok<AGROW>
        end
    end

    T = S(idxOut,:);

    Interpretation = strings(height(T),1);
    for i=1:height(T)
        s = string(T.Substrate(i));

        if strcmpi(s,"D-Glucose")
            Interpretation(i) = ...
                "Experimental reference; lower HETERO carbon demand than xylose";
        elseif strcmpi(s,"D-Xylose")
            Interpretation(i) = ...
                "Experimental reference; slightly higher HETERO carbon demand than glucose";
        elseif strcmpi(s,"meso-Erythritol")
            Interpretation(i) = ...
                "Lowest HETERO carbon demand; reported phenotype is slow growth";
        elseif strcmpi(s,"Glycerol")
            Interpretation(i) = ...
                "No HETERO growth; MIXO growth predominantly CO2-supported";
        elseif strcmpi(s,"Acetate")
            Interpretation(i) = ...
                "High HETERO carbon demand; lower MIXO CO2 contribution than sugars";
        elseif strcmpi(s,"Pyruvate")
            Interpretation(i) = ...
                "High HETERO carbon demand; distinct central-carbon entry point";
        end
    end

    T.Interpretation = Interpretation;
end

function T = buildCompactBOFReference_pub(cfg)

    Component = ["Protein";"Carbohydrate";"Lipid";"RNA";"DNA";"Pigment"];

    AUTO_coefficient = [ ...
        cfg.AUTO.proteinCoeff
        cfg.AUTO.carbohydrateCoeff
        cfg.AUTO.lipidCoeff
        cfg.AUTO.rnaCoeff
        cfg.AUTO.dnaCoeff
        cfg.AUTO.pigmentCoeff
    ];

    MIXO_coefficient = [ ...
        cfg.MIXO.proteinCoeff
        cfg.MIXO.carbohydrateCoeff
        cfg.MIXO.lipidCoeff
        cfg.MIXO.rnaCoeff
        cfg.MIXO.dnaCoeff
        cfg.MIXO.pigmentCoeff
    ];

    HETERO_coefficient = [ ...
        cfg.HETERO.proteinCoeff
        cfg.HETERO.carbohydrateCoeff
        cfg.HETERO.lipidCoeff
        cfg.HETERO.rnaCoeff
        cfg.HETERO.dnaCoeff
        cfg.HETERO.pigmentCoeff
    ];

    AUTO_demand_at_mu_0_060 = AUTO_coefficient*cfg.referenceMu_h;
    MIXO_demand_at_mu_0_060 = MIXO_coefficient*cfg.referenceMu_h;
    HETERO_demand_at_mu_0_060 = HETERO_coefficient*cfg.referenceMu_h;

    T = table(Component,AUTO_coefficient,MIXO_coefficient,HETERO_coefficient, ...
        AUTO_demand_at_mu_0_060,MIXO_demand_at_mu_0_060, ...
        HETERO_demand_at_mu_0_060);
end

function T = buildMethodsSummary_pub(S,autoCapacity,autoFixed,cfg)

    scored = ~strcmpi(strtrim(string(S.Growth_1995)),"Storage");

    hetMatch = strcmpi(string(S.HETERO_growth_agreement),"MATCH");
    mixMatch = strcmpi(string(S.MIXO_growth_agreement),"MATCH");

    nScored = sum(scored);
    nHetMatch = sum(scored & hetMatch);
    nMixMatch = sum(scored & mixMatch);

    scoredRows = find(scored);
    scoredExchange = string(S.Exchange(scoredRows));
    [~,ia] = unique(scoredExchange,'stable');
    representativeRows = scoredRows(ia);
    nUnique = numel(representativeRows);

    nHetUnique = sum(strcmpi( ...
        string(S.HETERO_growth_agreement(representativeRows)),"MATCH"));
    nMixUnique = sum(strcmpi( ...
        string(S.MIXO_growth_agreement(representativeRows)),"MATCH"));

    Metric = [ ...
        "Reference biomass flux"
        "Reference biomass-flux provenance"
        "Organic-carbon capacity supply"
        "AUTO reference"
        "AUTO mu_max"
        "AUTO minimum CO2 at reference mu"
        "AUTO BOF"
        "MIXO BOF"
        "HETERO BOF"
        "NGAM"
        "GAM"
        "Scored substrate entries"
        "Unique scored exchange conditions"
        "HETERO phenotype agreement"
        "MIXO phenotype agreement"
        "HETERO unique-condition agreement"
        "MIXO unique-condition agreement"
        "Starch"
        "HETERO mismatch"
        "MIXO carbon-partition basis"
        "Literature evidence"
        "Anomer interpretation"
        "Light treatment"
    ];

    Value = [ ...
        sprintf('%.6f h^-1',cfg.referenceMu_h)
        sprintf('%.2f d^-1 / 24; %s; %s', ...
            cfg.referenceMu_d,cfg.referenceMuCondition,cfg.referenceMuMethod)
        sprintf('%.0f C-mmol gDW^-1 h^-1',cfg.screenCarbon)
        "CO2-only; repeated beside each substrate only for comparison"
        sprintf('%.6f h^-1',autoCapacity.MaxMu_h)
        sprintf('%.6f C-mmol gDW^-1 h^-1',autoFixed.MinimumTotalCarbon)
        "TOTAL_BIOMASS_AUTO_MEAN"
        "Experimental MIXO BOF from previous cultivation validation"
        "Experimental HETERO BOF from previous cultivation validation"
        sprintf('%.1f mmol ATP gDW^-1 h^-1',cfg.NGAM)
        sprintf('%.0f mmol ATP gDW^-1 biomass',cfg.GAM)
        sprintf('%d',nScored)
        sprintf('%d',nUnique)
        sprintf('%d/%d',nHetMatch,nScored)
        sprintf('%d/%d',nMixMatch,nScored)
        sprintf('%d/%d',nHetUnique,nUnique)
        sprintf('%d/%d',nMixUnique,nUnique)
        "Storage; excluded from qualitative phenotype score"
        "Glycerol"
        "Standardized 6 C-mmol biomass-maximizing capacity solution"
        "Curated Galdieria-specific evidence and DOI fields are included for each substrate"
        "Alpha/beta model exchanges inherit parent-substrate evidence unless independently resolved experimentally"
        "Finite condition-specific model light; exact numerical constraint retained in code"
    ];

    T = table(Metric,Value);
end

function T = buildPublicationComparison_pub( ...
    substrateName,substrateClass,expGrowthText,resolvedExchange,carbonAtoms, ...
    autoCapacity,autoFixed,mixoCapacity,heteroCapacity,heteroFixed, ...
    expectedGrowth,cfg)

    n = numel(substrateName);

    Substrate = string(substrateName);
    Class = string(substrateClass);
    Growth_phenotype = string(expGrowthText);
    Carbon_atoms = carbonAtoms;
    Exchange = string(resolvedExchange);

    AUTO_mu_max = repmat(autoCapacity.MaxMu_h,n,1);
    AUTO_CO2_uptake_at_mu_max = repmat(autoCapacity.CO2Uptake,n,1);
    AUTO_min_CO2_at_reference_mu = ...
        repmat(autoFixed.MinimumTotalCarbon,n,1);

    HETERO_mu_max = [heteroCapacity.MaxMu_h]';
    HETERO_min_C_at_reference_mu = [heteroFixed.MinimumTotalCarbon]';
    HETERO_substrate_FVA_min = [heteroFixed.SubstrateFVAmin]';
    HETERO_substrate_FVA_max = [heteroFixed.SubstrateFVAmax]';

    HETERO_phenotype_agreement = strings(n,1);
    for i=1:n
        HETERO_phenotype_agreement(i) = ...
            agreementText_pub(expectedGrowth(i),heteroCapacity(i).PredictedGrowth);
    end

    MIXO_mu_max = [mixoCapacity.MaxMu_h]';
    MIXO_organic_uptake_at_mu_max = [mixoCapacity.SubstrateUptake]';
    MIXO_CO2_uptake_at_mu_max = [mixoCapacity.CO2Uptake]';

    MIXO_organic_C_at_mu_max = ...
        MIXO_organic_uptake_at_mu_max .* carbonAtoms;

    MIXO_total_external_C_at_mu_max = ...
        MIXO_organic_C_at_mu_max + MIXO_CO2_uptake_at_mu_max;

    MIXO_CO2_fraction_of_external_C = nan(n,1);
    ok = isfinite(MIXO_total_external_C_at_mu_max) & ...
         MIXO_total_external_C_at_mu_max > cfg.uptakeTol;
    MIXO_CO2_fraction_of_external_C(ok) = ...
        MIXO_CO2_uptake_at_mu_max(ok) ./ MIXO_total_external_C_at_mu_max(ok);

    MIXO_phenotype_agreement = strings(n,1);
    for i=1:n
        MIXO_phenotype_agreement(i) = ...
            agreementText_pub(expectedGrowth(i),mixoCapacity(i).PredictedGrowth);
    end

    Delta_mu_MIXO_minus_HETERO = MIXO_mu_max - HETERO_mu_max;

    MIXO_to_HETERO_mu_ratio = nan(n,1);
    validHet = isfinite(HETERO_mu_max) & HETERO_mu_max > cfg.growthTol;
    MIXO_to_HETERO_mu_ratio(validHet) = ...
        MIXO_mu_max(validHet) ./ HETERO_mu_max(validHet);

    Delta_mu_MIXO_minus_AUTO = MIXO_mu_max - AUTO_mu_max;

    HETERO_C_demand_to_AUTO_CO2_demand = nan(n,1);
    validDemand = isfinite(HETERO_min_C_at_reference_mu) & ...
                  isfinite(AUTO_min_CO2_at_reference_mu) & ...
                  AUTO_min_CO2_at_reference_mu > cfg.uptakeTol;
    HETERO_C_demand_to_AUTO_CO2_demand(validDemand) = ...
        HETERO_min_C_at_reference_mu(validDemand) ./ ...
        AUTO_min_CO2_at_reference_mu(validDemand);

    Interpretation = strings(n,1);
    for i=1:n
        hetGrows = isfinite(HETERO_mu_max(i)) && ...
                   HETERO_mu_max(i) > cfg.growthTol;
        mixGrows = isfinite(MIXO_mu_max(i)) && ...
                   MIXO_mu_max(i) > cfg.growthTol;

        if ~hetGrows && mixGrows
            Interpretation(i) = "MIXO rescues HETERO failure";
        elseif mixGrows && MIXO_mu_max(i) > AUTO_mu_max(i) + cfg.growthTol
            Interpretation(i) = "MIXO mu exceeds AUTO reference";
        elseif mixGrows && hetGrows && ...
               MIXO_mu_max(i) > HETERO_mu_max(i) + cfg.growthTol
            Interpretation(i) = "MIXO mu exceeds HETERO only";
        elseif mixGrows
            Interpretation(i) = ...
                "MIXO growth without clear capacity advantage";
        else
            Interpretation(i) = "No MIXO growth";
        end
    end

    T = table( ...
        Substrate,Class,Growth_phenotype,Carbon_atoms, ...
        AUTO_mu_max,AUTO_CO2_uptake_at_mu_max,AUTO_min_CO2_at_reference_mu, ...
        HETERO_mu_max,HETERO_phenotype_agreement, ...
        HETERO_min_C_at_reference_mu,HETERO_substrate_FVA_min, ...
        HETERO_substrate_FVA_max, ...
        MIXO_mu_max,MIXO_phenotype_agreement, ...
        MIXO_organic_uptake_at_mu_max,MIXO_organic_C_at_mu_max, ...
        MIXO_CO2_uptake_at_mu_max,MIXO_CO2_fraction_of_external_C, ...
        Delta_mu_MIXO_minus_HETERO,MIXO_to_HETERO_mu_ratio, ...
        Delta_mu_MIXO_minus_AUTO,HETERO_C_demand_to_AUTO_CO2_demand, ...
        Interpretation,Exchange, ...
        'VariableNames',{ ...
        'Substrate', ...
        'Class', ...
        'Growth_phenotype', ...
        'Carbon_atoms', ...
        'AUTO_mu_max_h_1', ...
        'AUTO_CO2_uptake_at_mu_max', ...
        'AUTO_min_CO2_at_reference_mu', ...
        'HETERO_mu_max_h_1', ...
        'HETERO_phenotype_agreement', ...
        'HETERO_min_C_at_reference_mu', ...
        'HETERO_substrate_FVA_min', ...
        'HETERO_substrate_FVA_max', ...
        'MIXO_mu_max_h_1', ...
        'MIXO_phenotype_agreement', ...
        'MIXO_organic_uptake_at_mu_max', ...
        'MIXO_organic_C_at_mu_max', ...
        'MIXO_CO2_uptake_at_mu_max', ...
        'MIXO_CO2_fraction_of_external_C', ...
        'Delta_mu_MIXO_minus_HETERO', ...
        'MIXO_to_HETERO_mu_ratio', ...
        'Delta_mu_MIXO_minus_AUTO', ...
        'HETERO_C_demand_to_AUTO_CO2_demand', ...
        'Interpretation', ...
        'Exchange'});
end

function writePublicationComparisonSheet_pub(fileName,T,cfg)

    sheet = 'PUBLICATION_COMPARISON';

    titleLine = { ...
        'Cross-trophic sole-substrate comparison'};

    noteLine1 = { ...
        ['AUTO values are the same CO2-only reference repeated for each ' ...
         'substrate row; they do not represent autotrophic utilization ' ...
         'of the listed organic substrate.']};

    noteLine2 = { ...
        ['HETERO fixed-mu carbon demand/FVA and AUTO fixed-mu CO2 demand ' ...
         'are retained. MIXO fixed-mu partition is intentionally omitted; ' ...
         'MIXO columns use the valid standardized 6 C-mmol capacity screen.']};

    writecell(titleLine,fileName,'Sheet',sheet,'Range','A1');
    writecell(noteLine1,fileName,'Sheet',sheet,'Range','A2');
    writecell(noteLine2,fileName,'Sheet',sheet,'Range','A3');

    writetable(T,fileName,'Sheet',sheet,'Range','A4');

    Metric = [ ...
        "AUTO reference"
        "Reference fixed mu"
        "Capacity organic-carbon supply"
        "HETERO fixed-mu carbon metric"
        "MIXO carbon-partition metric"
        "AUTO BOF"
        "MIXO BOF"
        "HETERO BOF"
    ];

    Definition = [ ...
        "Single CO2-only AUTO state repeated beside each substrate row"
        sprintf('%.6f h^-1; measured KIN2 HETERO mixed-sugar reference',cfg.referenceMu_h)
        sprintf('%.0f C-mmol gDW^-1 h^-1',cfg.screenCarbon)
        "Minimum carbon demand plus <=105% minimum-carbon FVA"
        "Biomass-maximizing standardized capacity solution"
        "TOTAL_BIOMASS_AUTO_MEAN"
        "Experimental BOF from publication cultivation validation"
        "Experimental BOF from publication cultivation validation"
    ];

    Units_or_note = [ ...
        "AUTO is not simulated on each organic substrate"
        "Used as standardized experimentally anchored comparison flux"
        "Organic substrate; CO2 may additionally enter MIXO"
        "C-mmol gDW^-1 h^-1 and mmol substrate gDW^-1 h^-1"
        "CO2 fraction calculated from external organic C + external CO2"
        "TOTAL_BIOMASS.xlsx"
        "Same composition as previous experimental validation"
        "Same composition as previous experimental validation"
    ];

    provenance = table(Metric,Definition,Units_or_note);
    writetable(provenance,fileName,'Sheet','PUBLICATION_COMPARISON_NOTES');
end

function m = prepareCondition_pub(baseModel,mode,cfg)
    mode = upper(string(mode));
    m = baseModel;

    for k = 1:numel(cfg.carbonBoundaryRxns)
        m = closeUptake_pub(m,cfg.carbonBoundaryRxns{k});
    end

    m = applyBasalMedium_pub(m,cfg);

    idx = find(strcmp(m.rxns,cfg.ngamRxn),1);
    m.lb(idx) = cfg.NGAM;
    m.ub(idx) = cfg.NGAM;

    if mode == "AUTO"
        m = setUptakeOnly_pub(m,cfg.photonRxn,cfg.autoPhotonProbeMax);
    elseif mode == "MIXO"
        m = setUptakeOnly_pub(m,cfg.photonRxn,cfg.mixoPhotonMaxUptake);
    elseif mode == "HETERO"
        m = setUptakeOnly_pub(m,cfg.photonRxn,cfg.heteroPhotonMaxUptake);
    else
        error('Unsupported trophic mode: %s',mode);
    end

    m = setUptakeOnly_pub(m,cfg.o2Rxn,cfg.o2MaxUptake);

    if mode == "AUTO" || mode == "MIXO"
        m = setDirectBounds_pub(m,cfg.o2DummyRxn,cfg.o2DummyLB,cfg.o2DummyUB);
    else
        m = setDirectBounds_pub(m,cfg.o2DummyRxn,0,0);
    end

    if mode == "AUTO"
        bof = cfg.AUTO;
    elseif mode == "MIXO"
        bof = cfg.MIXO;
    else
        bof = cfg.HETERO;
    end
    m = selectOnlyBOF_pub(m,bof);
    m = selectOnlyWorkbookPigmentProducer_pub(m,bof.pigmentRxn);

    if mode == "AUTO"
        m = openUptake_pub(m,cfg.co2Rxn,cfg.autoCO2CarbonCap,true);
        m = setRubisco_pub(m,cfg.rubiscoRxn,cfg.rubiscoMinFluxAUTO,1000);
    elseif mode == "MIXO"
        m = openUptake_pub(m,cfg.co2Rxn,cfg.mixoCO2MaxUptake,true);
        m = setRubisco_pub(m,cfg.rubiscoRxn,cfg.rubiscoMinFluxMIXO,1000);
    else
        m = closeUptake_pub(m,cfg.co2Rxn);
        m = setRubisco_pub(m,cfg.rubiscoRxn,0,0);
    end
end

function m = setDirectBounds_pub(m,rxn,lb,ub)
    idx = find(strcmp(m.rxns,rxn),1);
    if isempty(idx)
        error('Reaction not found while setting direct bounds: %s',rxn);
    end
    m.lb(idx) = lb;
    m.ub(idx) = ub;
end

function m = applyBasalMedium_pub(m,cfg)
    fields = {'Fe2','Mg2','NH4','NO3','H','Pi','SO4','H2O'};
    for k = 1:numel(fields)
        r = cfg.basalRxns.(fields{k});
        m = setDirectBounds_pub(m,r,cfg.basalMediumLB,cfg.basalMediumUB);
    end
end

function assertBasalMedium_pub(m,cfg,label)
    fields = {'Fe2','Mg2','NH4','NO3','H','Pi','SO4','H2O'};
    for k = 1:numel(fields)
        r = cfg.basalRxns.(fields{k});
        [lb,ub] = boundsOf_pub(m,r);
        if abs(lb-cfg.basalMediumLB)>1e-12 || abs(ub-cfg.basalMediumUB)>1e-12
            error('%s: basal-medium bounds incorrect for %s: [%g,%g], expected [%g,%g].', ...
                label,r,lb,ub,cfg.basalMediumLB,cfg.basalMediumUB);
        end
    end
end

function assertO2DummyBounds_pub(m,rxn,expectedLB,expectedUB,label)
    [lb,ub] = boundsOf_pub(m,rxn);
    if abs(lb-expectedLB)>1e-12 || abs(ub-expectedUB)>1e-12
        error('%s: O2 dummy bounds [%g,%g], expected [%g,%g].', ...
            label,lb,ub,expectedLB,expectedUB);
    end
end

function T = buildBasalMediumAudit_pub(baseModel,modelAUTO,modelMIXO,modelHETERO,cfg)
    roles = ["Fe2+";"Mg2+";"NH4+";"NO3-";"H+";"Pi";"SO4(2-)";"H2O";"O2 dummy"];
    rxns = string({cfg.basalRxns.Fe2;cfg.basalRxns.Mg2;cfg.basalRxns.NH4; ...
        cfg.basalRxns.NO3;cfg.basalRxns.H;cfg.basalRxns.Pi; ...
        cfg.basalRxns.SO4;cfg.basalRxns.H2O;cfg.o2DummyRxn});

    n = numel(rxns);
    Original_LB=nan(n,1); Original_UB=nan(n,1);
    AUTO_LB=nan(n,1); AUTO_UB=nan(n,1);
    MIXO_LB=nan(n,1); MIXO_UB=nan(n,1);
    HETERO_LB=nan(n,1); HETERO_UB=nan(n,1);
    Rule=strings(n,1);

    for i=1:n
        [Original_LB(i),Original_UB(i)] = boundsOf_pub(baseModel,char(rxns(i)));
        [AUTO_LB(i),AUTO_UB(i)] = boundsOf_pub(modelAUTO,char(rxns(i)));
        [MIXO_LB(i),MIXO_UB(i)] = boundsOf_pub(modelMIXO,char(rxns(i)));
        [HETERO_LB(i),HETERO_UB(i)] = boundsOf_pub(modelHETERO,char(rxns(i)));

        if rxns(i)==string(cfg.o2DummyRxn)
            Rule(i)="AUTO/MIXO light-condition sink 0..1000; HETERO 0..0";
        else
            Rule(i)="Light_Advance basal medium explicitly reset to -1000..1000";
        end
    end

    T=table(roles,rxns,Original_LB,Original_UB, ...
        AUTO_LB,AUTO_UB,MIXO_LB,MIXO_UB,HETERO_LB,HETERO_UB,Rule, ...
        'VariableNames',{'Medium_component','Reaction_ID','Original_SBML_LB', ...
        'Original_SBML_UB','AUTO_LB','AUTO_UB','MIXO_LB','MIXO_UB', ...
        'HETERO_LB','HETERO_UB','Applied_rule'});
end

function T = buildO2DummyUsageAudit_pub(autoRun,mixoRuns,heteroRuns,cfg)
    n = 1 + numel(mixoRuns) + numel(heteroRuns);
    Mode=strings(n,1); Substrate=strings(n,1);
    O2_Dummy_LB=nan(n,1); O2_Dummy_UB=nan(n,1);
    O2_Dummy_flux=nan(n,1); Mu_h=nan(n,1); Growth=false(n,1);

    row=1;
    Mode(row)="AUTO"; Substrate(row)="CO2_ONLY";
    O2_Dummy_LB(row)=cfg.o2DummyLB; O2_Dummy_UB(row)=cfg.o2DummyUB;
    O2_Dummy_flux(row)=autoRun.O2DummySinkFlux;
    Mu_h(row)=autoRun.Mu_h; Growth(row)=autoRun.PredictedGrowth;

    for i=1:numel(mixoRuns)
        row=row+1;
        Mode(row)="MIXO"; Substrate(row)=mixoRuns(i).Substrate;
        O2_Dummy_LB(row)=cfg.o2DummyLB; O2_Dummy_UB(row)=cfg.o2DummyUB;
        O2_Dummy_flux(row)=mixoRuns(i).O2DummySinkFlux;
        Mu_h(row)=mixoRuns(i).Mu_h; Growth(row)=mixoRuns(i).PredictedGrowth;
    end

    for i=1:numel(heteroRuns)
        row=row+1;
        Mode(row)="HETERO"; Substrate(row)=heteroRuns(i).Substrate;
        O2_Dummy_LB(row)=0; O2_Dummy_UB(row)=0;
        O2_Dummy_flux(row)=heteroRuns(i).O2DummySinkFlux;
        Mu_h(row)=heteroRuns(i).Mu_h; Growth(row)=heteroRuns(i).PredictedGrowth;
    end

    T=table(Mode,Substrate,O2_Dummy_LB,O2_Dummy_UB,O2_Dummy_flux,Mu_h,Growth);
end

function T = buildO2DummySensitivity_pub( ...
    modelAUTO,modelMIXO,substrateName,resolvedExchange,carbonAtoms, ...
    maxSubstrateUptake,autoRun,mixoRuns,cfg)

    nValid = sum(strlength(resolvedExchange)>0);
    n = 1+nValid;

    Mode=strings(n,1); Substrate=strings(n,1); Exchange=strings(n,1);
    Mu_with_dummy=nan(n,1); Mu_dummy_closed=nan(n,1);
    Delta_mu=nan(n,1); Growth_with_dummy=false(n,1);
    Growth_dummy_closed=false(n,1); Dummy_required_for_growth=false(n,1);
    Dummy_flux_with_dummy=nan(n,1);

    row=1;
    Mode(row)="AUTO"; Substrate(row)="CO2_ONLY"; Exchange(row)=string(cfg.co2Rxn);
    Mu_with_dummy(row)=autoRun.Mu_h;
    Growth_with_dummy(row)=autoRun.PredictedGrowth;
    Dummy_flux_with_dummy(row)=autoRun.O2DummySinkFlux;

    m=modelAUTO;
    m=setDirectBounds_pub(m,cfg.o2DummyRxn,0,0);
    [rClosed,~]=optimizeAutoAdaptive_pub(m,cfg.AUTO.sinkRxn,cfg);
    Mu_dummy_closed(row)=rClosed.Mu_h;
    Growth_dummy_closed(row)=rClosed.PredictedGrowth;
    Delta_mu(row)=Mu_with_dummy(row)-Mu_dummy_closed(row);
    Dummy_required_for_growth(row)=Growth_with_dummy(row) && ~Growth_dummy_closed(row);

    for i=1:numel(substrateName)
        if strlength(resolvedExchange(i))==0
            continue;
        end
        row=row+1;
        r=char(resolvedExchange(i));

        Mode(row)="MIXO";
        Substrate(row)=substrateName(i);
        Exchange(row)=string(r);
        Mu_with_dummy(row)=mixoRuns(i).Mu_h;
        Growth_with_dummy(row)=mixoRuns(i).PredictedGrowth;
        Dummy_flux_with_dummy(row)=mixoRuns(i).O2DummySinkFlux;

        m=modelMIXO;
        m=openUptake_pub(m,r,maxSubstrateUptake(i),false);
        m=setDirectBounds_pub(m,cfg.o2DummyRxn,0,0);

        assertCarbonBoundaryState_pub(m,{cfg.co2Rxn,r},cfg,'MIXO_O2_DUMMY_CLOSED');
        assertBiomassOutputState_pub(m,cfg.MIXO.sinkRxn,cfg,'MIXO_O2_DUMMY_CLOSED');

        rr=optimizeState_pub(m,cfg.MIXO.sinkRxn,r,carbonAtoms(i),cfg, ...
            'MIXO_O2_DUMMY_CLOSED',substrateName(i));

        Mu_dummy_closed(row)=rr.Mu_h;
        Growth_dummy_closed(row)=rr.PredictedGrowth;
        Delta_mu(row)=Mu_with_dummy(row)-Mu_dummy_closed(row);
        Dummy_required_for_growth(row)=Growth_with_dummy(row) && ~Growth_dummy_closed(row);
    end

    T=table(Mode,Substrate,Exchange,Mu_with_dummy,Mu_dummy_closed,Delta_mu, ...
        Growth_with_dummy,Growth_dummy_closed,Dummy_required_for_growth, ...
        Dummy_flux_with_dummy);
end

function run = optimizeState_pub(m,biomassRxn,subRxn,nC,cfg,mode,substrateLabel)
    run = emptyRun_pub();
    run.Mode = string(mode);
    run.Substrate = string(substrateLabel);
    run.BiomassRxn = string(biomassRxn);
    run.SubstrateRxn = string(subRxn);
    run.CarbonAtoms = nC;

    m = setObjective_pub(m,biomassRxn);
    sol = optimizeCbModel(m,'max');

    if ~isfield(sol,'stat') || sol.stat ~= 1 || isempty(sol.f) || isempty(sol.x)
        run.Status = "INFEASIBLE";
        run.Mu_h = 0;
        run.PredictedGrowth = false;
        return;
    end

    run.Status = "OPTIMAL";
    run.Mu_h = max(0,sol.f);
    run.PredictedGrowth = run.Mu_h > cfg.growthTol;
    run.CO2Uptake_mmol = uptakeMagnitudeFromSolution_pub(m,cfg.co2Rxn,sol.x);
    run.PhotonUptake = uptakeMagnitudeFromSolution_pub(m,cfg.photonRxn,sol.x);

    didx = find(strcmp(m.rxns,cfg.o2DummyRxn),1);
    if ~isempty(didx)
        run.O2DummySinkFlux = sol.x(didx);
    end

    if ~isempty(subRxn)
        run.SubstrateUptake_mmol = uptakeMagnitudeFromSolution_pub(m,subRxn,sol.x);
        run.SubstrateUptake_Cmmol = run.SubstrateUptake_mmol * nC;

        if run.PredictedGrowth
            [minReq,maxPos] = substrateUseRangeAtBiomass_pub( ...
                m,biomassRxn,subRxn, ...
                cfg.biomassFractionForUseTest*run.Mu_h);
            run.MinRequiredUptakeAtFraction_mmol = minReq;
            run.MaxPossibleUptakeAtFraction_mmol = maxPos;
            run.CanConsumeAtFraction = maxPos > cfg.uptakeTol;
        end
    end
end

function [minReq,maxPos] = substrateUseRangeAtBiomass_pub(m,biomassRxn,subRxn,targetMu)
    minReq = NaN;
    maxPos = NaN;

    bidx = find(strcmp(m.rxns,biomassRxn),1);
    if isempty(bidx)
        return;
    end

    m.lb(bidx) = targetMu;
    m.ub(bidx) = targetMu;

    sidx = find(strcmp(m.rxns,subRxn),1);
    if isempty(sidx)
        return;
    end

    uptakeSign = uptakeFluxSign_pub(m,subRxn); % -1 if uptake is negative
    m.c(:) = 0;
    m.c(sidx) = 1;

    if uptakeSign < 0
        solMin = optimizeCbModel(m,'max');
        solMax = optimizeCbModel(m,'min');
    else
        solMin = optimizeCbModel(m,'min');
        solMax = optimizeCbModel(m,'max');
    end

    if isfield(solMin,'stat') && solMin.stat == 1 && ~isempty(solMin.x)
        minReq = uptakeMagnitudeFromSolution_pub(m,subRxn,solMin.x);
    end
    if isfield(solMax,'stat') && solMax.stat == 1 && ~isempty(solMax.x)
        maxPos = uptakeMagnitudeFromSolution_pub(m,subRxn,solMax.x);
    end
end

function m = selectOnlyBOF_pub(m,bof)
    nr = string(cellfun(@norm_pub,m.rxns,'UniformOutput',false));
    raw = lower(string(m.rxns));

    cand = find(contains(nr,'TOTALBIOMASS') | ...
                contains(nr,'BMTX') | endsWith(raw,"_bm_tx"));

    keep = string({bof.assemblyRxn,bof.sinkRxn});

    for j = cand(:)'
        if ~any(string(m.rxns{j}) == keep)
            m.lb(j) = 0;
            m.ub(j) = 0;
            if isfield(m,'c') && numel(m.c)>=j
                m.c(j) = 0;
            end
        end
    end

    aidx = find(strcmp(m.rxns,bof.assemblyRxn),1);
    sidx = find(strcmp(m.rxns,bof.sinkRxn),1);

    if isempty(aidx) || isempty(sidx)
        error('Selected BOF reactions are missing for %s.',bof.mode);
    end

    m.lb(aidx) = 0;
    m.ub(aidx) = max(1000,m.ub(aidx));
    m.lb(sidx) = 0;
    m.ub(sidx) = max(1000,m.ub(sidx));
    m = setObjective_pub(m,bof.sinkRxn);
end

function m = selectOnlyWorkbookPigmentProducer_pub(m,keepRxn)
    nr = string(cellfun(@norm_pub,m.rxns,'UniformOutput',false));

    oldProducers = find(contains(nr,'PIGMENTTOTALPOOLBIO'));
    oldSinks = find(contains(nr,'PIGMENTTOTALPOOLSINK'));
    analysisProducers = find(contains(nr,'S2PIGMENTTOTALPOOL') & contains(nr,'PRODUCER'));

    closeIdx = unique([oldProducers(:); oldSinks(:); analysisProducers(:)]);

    for j = closeIdx(:)'
        m.lb(j) = 0;
        m.ub(j) = 0;
        if isfield(m,'c') && numel(m.c) >= j
            m.c(j) = 0;
        end
    end

    idx = find(strcmp(m.rxns,keepRxn),1);
    if isempty(idx)
        error('New workbook-derived pigment producer not found: %s',keepRxn);
    end

    m.lb(idx) = 0;
    m.ub(idx) = 1000;
end

function ensureCobraReady_pub()
    coreFunctions = { ...
        'optimizeCbModel', ...
        'addReaction', ...
        'addMetabolite', ...
        'changeObjective', ...
        'removeRxns'};

    present = false(size(coreFunctions));
    for k = 1:numel(coreFunctions)
        present(k) = exist(coreFunctions{k},'file') == 2;
    end

    if all(present)
        return;
    end

    missing = coreFunctions(~present);

    if exist('initCobraToolbox','file') ~= 2
        error(['COBRA Toolbox is not initialized and initCobraToolbox is not on the MATLAB path. ' ...
               'Add the COBRA Toolbox folder to the MATLAB path first.']);
    end

    initFile = which('initCobraToolbox');
    cobraRoot = fileparts(initFile);
    tmpDir = fullfile(cobraRoot,'.tmp');
    if ~isfolder(tmpDir)
        try
            mkdir(tmpDir);
        catch MEtmp
            warning('Could not create COBRA .tmp folder: %s',MEtmp.message);
        end
    end

    try
        evalc('initCobraToolbox(false);');
    catch ME
        error(['COBRA initialization failed before the sole-substrate analysis.\n' ...
               'The model and BOF code have not run yet.\n' ...
               'Original COBRA initialization error:\n%s'],ME.message);
    end

    stillMissing = {};
    for k = 1:numel(coreFunctions)
        if exist(coreFunctions{k},'file') ~= 2
            stillMissing{end+1} = coreFunctions{k}; %#ok<AGROW>
        end
    end
    if ~isempty(stillMissing)
        error('COBRA initialized, but required functions are still unavailable: %s', ...
              strjoin(stillMissing,', '));
    end
end

function ensureLPSolverSelected_pub()
    global CBT_LP_SOLVER

    if ~isempty(CBT_LP_SOLVER)
        currentSolver = strtrim(char(string(CBT_LP_SOLVER)));
        if ~isempty(currentSolver)
            return;
        end
    end

    if exist('changeCobraSolver','file') ~= 2
        error(['COBRA Toolbox functions are available, but no LP solver is ' ...
               'selected and changeCobraSolver is not on the MATLAB path. ' ...
               'Run initCobraToolbox once, then rerun this analysis.']);
    end

    candidates = {'gurobi','glpk','linprog'};
    messages = strings(numel(candidates),1);

    for k = 1:numel(candidates)
        solverName = candidates{k};
        try
            evalc('ok = changeCobraSolver(solverName,''LP'',0);');
            selected = false;
            if ~isempty(ok)
                try
                    selected = logical(ok);
                catch
                    selected = false;
                end
            end
            if ~isempty(CBT_LP_SOLVER)
                selected = selected || strcmpi( ...
                    strtrim(char(string(CBT_LP_SOLVER))),solverName);
            end
            if selected
                return;
            end
            messages(k) = "changeCobraSolver returned without selecting solver";
        catch MEsolver
            messages(k) = string(MEsolver.message);
        end
    end

    if exist('initCobraToolbox','file') == 2
        try
            evalc('initCobraToolbox(false);');
        catch MEinit
            warning('Fallback initCobraToolbox failed: %s',MEinit.message);
        end

        if ~isempty(CBT_LP_SOLVER)
            currentSolver = strtrim(char(string(CBT_LP_SOLVER)));
            if ~isempty(currentSolver)
                return;
            end
        end

        for k = 1:numel(candidates)
            solverName = candidates{k};
            try
                evalc('ok = changeCobraSolver(solverName,''LP'',0);');
                if (~isempty(ok) && logical(ok)) || ...
                        (~isempty(CBT_LP_SOLVER) && strcmpi( ...
                        strtrim(char(string(CBT_LP_SOLVER))),solverName))
                    return;
                end
            catch
            end
        end
    end

    detail = "";
    for k = 1:numel(candidates)
        detail = detail + newline + "  " + candidates{k} + ": " + messages(k);
    end
    error(['COBRA Toolbox is available, but the analysis could not select an LP solver.' ...
           newline 'Tried Gurobi, GLPK and linprog.' ...
           newline 'Solver-selection messages:' char(detail) ...
           newline 'You can also run manually:' ...
           newline '  changeCobraSolver(''gurobi'',''LP'');']);
end

function m = loadCobraModelFromMat_pub(modelFile)
    X = load(modelFile);
    fn = fieldnames(X);
    hits = {};
    for k = 1:numel(fn)
        v = X.(fn{k});
        if isCobraStruct_pub(v)
            hits{end+1} = fn{k}; %#ok<AGROW>
        end
    end
    if isempty(hits)
        error('No COBRA model struct was found inside MAT file: %s',modelFile);
    elseif numel(hits) > 1
        error(['More than one COBRA-like model struct was found inside %s. ' ...
               'Load the desired one manually and pass it to this analysis. Variables: %s'], ...
               modelFile,strjoin(hits,', '));
    end
    m = X.(hits{1});
end

function tf = isCobraStruct_pub(x)
    tf = isstruct(x) && all(isfield(x,{'rxns','mets','S','lb','ub'}));
end

function [m,loaderUsed] = loadSBMLModel_pub(modelFile)
    if exist('TranslateSBML','file') == 2
        try
            evalc('m = readCbModel(modelFile,''fileType'',''SBML'');');
            loaderUsed = 'COBRA readCbModel/TranslateSBML';
            return;
        catch ME
        end
    else
    end

    m = readGaldieriaSBMLDirect_pub(modelFile);
    loaderUsed = 'Direct SBML Level-3/FBC parser';
end

function model = readGaldieriaSBMLDirect_pub(modelFile)
    if exist('xmlread','file') ~= 2 && exist('xmlread','builtin') ~= 5
        error(['MATLAB xmlread is unavailable, so the direct SBML parser cannot run. ' ...
               'A Java-enabled MATLAB installation is required.']);
    end

    doc = xmlread(modelFile);

    modelNodes = doc.getElementsByTagName('model');
    if modelNodes.getLength == 0
        error('No <model> element found in SBML file: %s',modelFile);
    end
    modelNode = modelNodes.item(0);

    model = struct();
    model.id = getAttrByLocalName_pub(modelNode,'id','Galdieria_SBML');
    model.description = sprintf('Direct SBML import: %s',modelFile);

    compNodes = doc.getElementsByTagName('compartment');
    nComp = double(compNodes.getLength);
    comps = cell(nComp,1);
    compNames = cell(nComp,1);
    compMap = containers.Map('KeyType','char','ValueType','double');
    for i = 1:nComp
        node = compNodes.item(i-1);
        cid = getAttrByLocalName_pub(node,'id',sprintf('comp%d',i));
        cname = getAttrByLocalName_pub(node,'name',cid);
        comps{i} = cid;
        compNames{i} = cname;
        compMap(cid) = i;
    end
    if nComp == 0
        comps = {'c'};
        compNames = {'cytosol'};
        compMap('c') = 1;
        nComp = 1;
    end
    model.comps = comps;
    model.compNames = compNames;

    speciesNodes = doc.getElementsByTagName('species');
    nMet = double(speciesNodes.getLength);
    if nMet == 0
        error('No species were found in the SBML file.');
    end

    rawMetIds = cell(nMet,1);
    mets = cell(nMet,1);
    metNames = cell(nMet,1);
    metComps = ones(nMet,1);
    metFormulas = cell(nMet,1);
    metCharges = nan(nMet,1);
    metBoundary = false(nMet,1);
    metMap = containers.Map('KeyType','char','ValueType','double');

    for i = 1:nMet
        node = speciesNodes.item(i-1);
        rawId = getAttrByLocalName_pub(node,'id','');
        if isempty(rawId)
            error('Encountered SBML species without an id at index %d.',i);
        end
        rawMetIds{i} = rawId;
        mets{i} = decodeSBMLId_pub(rawId);
        metNames{i} = getAttrByLocalName_pub(node,'name',mets{i});

        comp = getAttrByLocalName_pub(node,'compartment','c');
        if isKey(compMap,comp)
            metComps(i) = compMap(comp);
        else
            comps{end+1,1} = comp; %#ok<AGROW>
            compNames{end+1,1} = comp; %#ok<AGROW>
            compMap(comp) = numel(comps);
            metComps(i) = numel(comps);
        end

        metFormulas{i} = getAttrByLocalName_pub(node,'chemicalFormula','');
        chargeTxt = getAttrByLocalName_pub(node,'charge','');
        if ~isempty(chargeTxt)
            q = str2double(chargeTxt);
            if ~isnan(q), metCharges(i) = q; end
        end
        metBoundary(i) = parseBoolean_pub(getAttrByLocalName_pub(node,'boundaryCondition','false'));

        if isKey(metMap,rawId)
            error('Duplicate raw SBML species id: %s',rawId);
        end
        metMap(rawId) = i;
    end

    if numel(unique(string(mets))) ~= nMet
        error('Decoding SBML species IDs produced duplicate COBRA metabolite IDs.');
    end

    model.mets = mets;
    model.metNames = metNames;
    model.metComps = metComps;
    model.metFormulas = metFormulas;
    model.metCharges = metCharges;
    model.metBoundary = metBoundary;
    model.comps = comps;
    model.compNames = compNames;

    paramNodes = doc.getElementsByTagName('parameter');
    paramMap = containers.Map('KeyType','char','ValueType','double');
    for i = 1:double(paramNodes.getLength)
        node = paramNodes.item(i-1);
        pid = getAttrByLocalName_pub(node,'id','');
        valTxt = getAttrByLocalName_pub(node,'value','');
        if ~isempty(pid) && ~isempty(valTxt)
            val = str2double(valTxt);
            if ~isnan(val)
                paramMap(pid) = val;
            end
        end
    end

    reactionNodes = doc.getElementsByTagName('reaction');
    nRxn = double(reactionNodes.getLength);
    if nRxn == 0
        error('No reactions were found in the SBML file.');
    end

    rawRxnIds = cell(nRxn,1);
    rxns = cell(nRxn,1);
    rxnNames = cell(nRxn,1);
    lb = zeros(nRxn,1);
    ub = zeros(nRxn,1);
    reversible = false(nRxn,1);
    rxnMapRaw = containers.Map('KeyType','char','ValueType','double');

    I = zeros(0,1); J = zeros(0,1); V = zeros(0,1);

    for j = 1:nRxn
        rnode = reactionNodes.item(j-1);
        rawRid = getAttrByLocalName_pub(rnode,'id','');
        if isempty(rawRid)
            error('Encountered SBML reaction without an id at index %d.',j);
        end
        rawRxnIds{j} = rawRid;
        rxns{j} = decodeSBMLId_pub(rawRid);
        rxnNames{j} = getAttrByLocalName_pub(rnode,'name',rxns{j});
        reversible(j) = parseBoolean_pub(getAttrByLocalName_pub(rnode,'reversible','false'));

        lowerRef = getAttrByLocalName_pub(rnode,'lowerFluxBound','');
        upperRef = getAttrByLocalName_pub(rnode,'upperFluxBound','');
        lb(j) = resolveFBCBound_pub(lowerRef,paramMap, ternary_pub(reversible(j),-1000,0));
        ub(j) = resolveFBCBound_pub(upperRef,paramMap,1000);

        if isKey(rxnMapRaw,rawRid)
            error('Duplicate raw SBML reaction id: %s',rawRid);
        end
        rxnMapRaw(rawRid) = j;

        reactList = firstDescendantByLocalName_pub(rnode,'listOfReactants');
        if ~isempty(reactList)
            refs = reactList.getElementsByTagName('speciesReference');
            for k = 1:double(refs.getLength)
                ref = refs.item(k-1);
                rawSid = getAttrByLocalName_pub(ref,'species','');
                if ~isKey(metMap,rawSid)
                    error('Reaction %s references unknown reactant species %s.',rawRid,rawSid);
                end
                sto = getNumericAttr_pub(ref,'stoichiometry',1);
                I(end+1,1) = metMap(rawSid); %#ok<AGROW>
                J(end+1,1) = j; %#ok<AGROW>
                V(end+1,1) = -sto; %#ok<AGROW>
            end
        end

        prodList = firstDescendantByLocalName_pub(rnode,'listOfProducts');
        if ~isempty(prodList)
            refs = prodList.getElementsByTagName('speciesReference');
            for k = 1:double(refs.getLength)
                ref = refs.item(k-1);
                rawSid = getAttrByLocalName_pub(ref,'species','');
                if ~isKey(metMap,rawSid)
                    error('Reaction %s references unknown product species %s.',rawRid,rawSid);
                end
                sto = getNumericAttr_pub(ref,'stoichiometry',1);
                I(end+1,1) = metMap(rawSid); %#ok<AGROW>
                J(end+1,1) = j; %#ok<AGROW>
                V(end+1,1) = sto; %#ok<AGROW>
            end
        end
    end

    if numel(unique(string(rxns))) ~= nRxn
        error('Decoding SBML reaction IDs produced duplicate COBRA reaction IDs.');
    end

    model.rxns = rxns;
    model.rxnNames = rxnNames;
    model.S = sparse(I,J,V,nMet,nRxn);
    model.lb = lb;
    model.ub = ub;
    model.rev = double(lb < 0 & ub > 0);
    model.b = zeros(nMet,1);
    model.csense = repmat('E',nMet,1);

    c = zeros(nRxn,1);
    activeObjective = '';
    listObj = firstDescendantByLocalName_pub(modelNode,'listOfObjectives');
    if ~isempty(listObj)
        activeObjective = getAttrByLocalName_pub(listObj,'activeObjective','');
        objNodes = descendantsByLocalName_pub(listObj,'objective');
        selectedObj = [];
        for k = 1:numel(objNodes)
            oid = getAttrByLocalName_pub(objNodes{k},'id','');
            if isempty(selectedObj)
                selectedObj = objNodes{k};
            end
            if ~isempty(activeObjective) && strcmp(oid,activeObjective)
                selectedObj = objNodes{k};
                break;
            end
        end
        if ~isempty(selectedObj)
            fluxObjs = descendantsByLocalName_pub(selectedObj,'fluxObjective');
            for k = 1:numel(fluxObjs)
                rawRid = getAttrByLocalName_pub(fluxObjs{k},'reaction','');
                coeff = getNumericAttr_pub(fluxObjs{k},'coefficient',1);
                if isKey(rxnMapRaw,rawRid)
                    c(rxnMapRaw(rawRid)) = coeff;
                end
            end
        end
    end
    model.c = c;
    model.osenseStr = 'max';
    model.osense = -1;

    model.genes = cell(0,1);
    model.grRules = repmat({''},nRxn,1);
    model.rules = repmat({''},nRxn,1);
    model.rxnGeneMat = sparse(nRxn,0);

    model.sbmlRawMetIds = rawMetIds;
    model.sbmlRawRxnIds = rawRxnIds;

    if countElementsByLocalName_pub(modelNode,'userDefinedConstraint') > 0
        warning(['The SBML contains FBC userDefinedConstraint elements. Direct import ' ...
                 'does not transfer them. Use standard readCbModel if those constraints are required.']);
    end

end

function val = resolveFBCBound_pub(ref,paramMap,defaultVal)
    if isempty(ref)
        val = defaultVal;
        return;
    end
    if isKey(paramMap,ref)
        val = paramMap(ref);
        return;
    end
    x = str2double(ref);
    if ~isnan(x)
        val = x;
    else
        warning('Could not resolve FBC bound reference %s; using default %.12g.',ref,defaultVal);
        val = defaultVal;
    end
end

function node = firstDescendantByLocalName_pub(parent,target)
    node = [];
    allNodes = parent.getElementsByTagName('*');
    for i = 1:double(allNodes.getLength)
        x = allNodes.item(i-1);
        if strcmpi(nodeLocalName_pub(x),target)
            node = x;
            return;
        end
    end
end

function out = descendantsByLocalName_pub(parent,target)
    out = {};
    allNodes = parent.getElementsByTagName('*');
    for i = 1:double(allNodes.getLength)
        x = allNodes.item(i-1);
        if strcmpi(nodeLocalName_pub(x),target)
            out{end+1,1} = x; %#ok<AGROW>
        end
    end
end

function n = countElementsByLocalName_pub(parent,target)
    n = 0;
    allNodes = parent.getElementsByTagName('*');
    for i = 1:double(allNodes.getLength)
        if strcmpi(nodeLocalName_pub(allNodes.item(i-1)),target)
            n = n + 1;
        end
    end
end

function name = nodeLocalName_pub(node)
    try
        name = char(node.getLocalName());
    catch
        name = '';
    end
    if isempty(name)
        name = char(node.getNodeName());
        p = strfind(name,':');
        if ~isempty(p), name = name(p(end)+1:end); end
    end
end

function val = getAttrByLocalName_pub(node,target,defaultVal)
    val = defaultVal;
    if isempty(node) || ~node.hasAttributes()
        return;
    end
    attrs = node.getAttributes();
    for i = 1:double(attrs.getLength)
        a = attrs.item(i-1);
        nm = nodeLocalName_pub(a);
        if strcmpi(nm,target)
            val = char(a.getNodeValue());
            return;
        end
    end
end

function x = getNumericAttr_pub(node,target,defaultVal)
    txt = getAttrByLocalName_pub(node,target,'');
    if isempty(txt)
        x = defaultVal;
    else
        x = str2double(txt);
        if isnan(x), x = defaultVal; end
    end
end

function tf = parseBoolean_pub(txt)
    t = lower(strtrim(char(string(txt))));
    tf = any(strcmp(t,{'true','1','yes'}));
end

function out = ternary_pub(cond,a,b)
    if cond, out = a; else, out = b; end
end

function decoded = decodeSBMLId_pub(raw)
    decoded = char(raw);
    [st,en,tok] = regexp(decoded,'__(\d+)__','start','end','tokens');
    for k = numel(st):-1:1
        code = str2double(tok{k}{1});
        if ~isnan(code) && code >= 1 && code <= 255
            decoded = [decoded(1:st(k)-1), char(code), decoded(en(k)+1:end)]; %#ok<AGROW>
        end
    end
end

function [m,auditT] = closeOriginalBiomassOutputs_pub(m)
    ids = string(m.rxns);
    normIds = string(cellfun(@norm_pub,m.rxns,'UniformOutput',false));

    if isfield(m,'rxnNames')
        names = string(m.rxnNames);
        normNames = string(cellfun(@norm_pub,m.rxnNames,'UniformOutput',false));
    else
        names = repmat("",numel(m.rxns),1);
        normNames = repmat("",numel(m.rxns),1);
    end

    rawLower = lower(ids);
    rawNameLower = lower(names);

    isBmTx = endsWith(rawLower,"_bm_tx") | ...
             endsWith(rawNameLower,"_bm_tx") | ...
             contains(normIds,'BMTX') | contains(normNames,'BMTX');

    isTotalBiomass = contains(normIds,'TOTALBIOMASS') | ...
                     contains(normNames,'TOTALBIOMASS');

    idx = find(isBmTx | isTotalBiomass);

    oldLB = m.lb(idx);
    oldUB = m.ub(idx);
    oldC = zeros(numel(idx),1);

    if isfield(m,'c') && numel(m.c)==numel(m.rxns)
        oldC = m.c(idx);
        m.c(idx) = 0;
    end

    m.lb(idx) = 0;
    m.ub(idx) = 0;

    reason = strings(numel(idx),1);
    for k = 1:numel(idx)
        if isBmTx(idx(k)) && isTotalBiomass(idx(k))
            reason(k) = "bm_tx + TOTAL_BIOMASS";
        elseif isBmTx(idx(k))
            reason(k) = "pre-existing bm_tx";
        else
            reason(k) = "TOTAL_BIOMASS";
        end
    end

    auditT = table(ids(idx),names(idx),reason,oldLB,oldUB,oldC, ...
        zeros(numel(idx),1),zeros(numel(idx),1), ...
        'VariableNames',{'Reaction_ID','Reaction_name','Closure_reason', ...
        'Original_LB','Original_UB','Original_objective_coeff','New_LB','New_UB'});
end

function [m,cfg,auditT] = implementWorkbookMeanBOFs_pub(m,cfg,biomassFile)

    specs = { ...
        'AUTO','Reaction List Auto','TOTAL_BIOMASS_AUTO_MEAN', ...
            'PIGMENT_TOTAL_POOL_BIO_AUTOTROPH_NOFRAC'; ...
        'MIXO','Reaction List Mixo','TOTAL_BIOMASS_MIXO_MEAN', ...
            'PIGMENT_TOTAL_POOL_BIO_MIXOTROPH_NOFRAC'; ...
        'HETERO','Reaction List Heto','TOTAL_BIOMASS_HETER_MEAN', ...
            'PIGMENT_TOTAL_POOL_BIO_HETEROTROPH_GLUCOSE_NOFRAC'};

    % Shared energy and pigment metabolites are common. Macromolecular
    % pool metabolites are resolved per trophic state so the latest
    % condition-specific SBML pool layer can be used.
    shared.atp     = requireMetCore_pub(m,'ATP_CYTO');
    shared.water   = requireMetCore_pub(m,'WATER_CYTO');
    shared.adp     = requireMetCore_pub(m,'ADP_CYTO');
    shared.pi      = requireMetCore_pub(m,'PI_CYTO');
    shared.proton  = requireMetCore_pub(m,'PROTON_CYTO');

    pig.zea  = requireMetCore_pub(m,'CPD1F-130_THYL');
    pig.chla = requireMetAnyCore_pub(m,{'CHLOROPHYLL-A_PLAS','CHLOROPHYLL-A_THYL'});
    pig.beta = requireMetCore_pub(m,'CPD1F-129_THYL');
    pig.pcb  = requireMetCore_pub(m,'3Z-PHYCOCYANOBILIN_PLAS');

    cfg.resolvedPigmentMets = pig;
    cfg.resolvedBOFMets = resolveModeBOFMets_pub(m,'AUTO');
    rows = cell(size(specs,1),25);

    for i = 1:size(specs,1)
        mode = specs{i,1};
        sheetName = specs{i,2};
        workbookBOF = specs{i,3};
        workbookPigment = specs{i,4};
        metsMode = resolveModeBOFMets_pub(m,mode);

        sourceFormula = readWorkbookReactionFormula_pub( ...
            biomassFile,sheetName,workbookBOF);
        pigmentFormula = readWorkbookReactionFormula_pub( ...
            biomassFile,sheetName,workbookPigment);

        cProtein = coefficientForWorkbookMet_pub(sourceFormula,'m_PROTEIN_TOTAL_POOL[c]');
        cCarb    = coefficientForWorkbookMet_pub(sourceFormula,'m_CARBOHYDRATE_TOTAL_POOL[c]');
        cLipid   = coefficientForWorkbookMet_pub(sourceFormula,'m_LIPID_TOTAL_POOL[c]');
        cRNA     = coefficientForWorkbookMet_pub(sourceFormula,'m_RNA_TOTAL_POOL[c]');
        cDNA     = coefficientForWorkbookMet_pub(sourceFormula,'m_DNA_TOTAL_POOL[c]');
        cPigment = coefficientForWorkbookMet_pub(sourceFormula,'m_PIGMENT_TOTAL_POOL[c]');

        cATP     = coefficientForWorkbookMet_pub(sourceFormula,'m_ATP_Cyto[c]');
        cWater   = coefficientForWorkbookMet_pub(sourceFormula,'m_WATER_Cyto[c]');
        cADP     = coefficientForWorkbookMet_pub(sourceFormula,'m_ADP_Cyto[c]');
        cPi      = coefficientForWorkbookMet_pub(sourceFormula,'m_Pi_Cyto[c]');
        cProton  = coefficientForWorkbookMet_pub(sourceFormula,'m_PROTON_Cyto[c]');

        gamVec = [cATP,cWater,cADP,cPi,cProton];
        if any(abs(gamVec - 30) > 1e-9)
            error(['Unexpected GAM coefficients in %s / %s. ' ...
                   'Expected ATP/H2O/ADP/Pi/H = 30; obtained %s.'], ...
                   sheetName,workbookBOF,mat2str(gamVec,12));
        end

        pZea  = coefficientForWorkbookMet_pub(pigmentFormula,'m_CPD1F-130_Thyl[t]');
        pChlA = coefficientForWorkbookMet_pub(pigmentFormula,'m_CHLOROPHYLL-A_Plas[p]');
        pBeta = coefficientForWorkbookMet_pub(pigmentFormula,'m_CPD1F-129_Thyl[t]');
        pPCB  = coefficientForWorkbookMet_pub(pigmentFormula,'m_3Z-PHYCOCYANOBILIN_Plas[p]');

        pigSum = pZea + pChlA + pBeta + pPCB;
        if abs(pigSum - 1) > 0.01
            error(['NOFRAC pigment composition for %s does not sum to ~1. ' ...
                   'Obtained %.12g from TOTAL_BIOMASS.xlsx.'],mode,pigSum);
        end

        newPigMet = sprintf('m_S2_PIGMENT_TOTAL_POOL_%s_PUB[c]',mode);
        newPigRxn = sprintf('S2_PIGMENT_TOTAL_POOL_%s_PUB_PRODUCER_bm',mode);

        if ismember(newPigRxn,m.rxns)
            m = removeRxns(m,{newPigRxn});
        end
        if ~ismember(newPigMet,m.mets)
            m = addMetabolite(m,newPigMet, ...
                'metName',sprintf('Sole-substrate %s pigment total pool',mode), ...
                'metFormula','','Charge',0);
        end

        m = addReaction(m,newPigRxn, ...
            'reactionName',sprintf('%s pigment composition from TOTAL_BIOMASS.xlsx',mode), ...
            'metaboliteList',{pig.zea;pig.chla;pig.beta;pig.pcb;newPigMet}, ...
            'stoichCoeffList',[-pZea;-pChlA;-pBeta;-pPCB;1], ...
            'lowerBound',0,'upperBound',1000, ...
            'subSystem','Biomass pigment pool');
        m = padConstraintColumns_pub(m);

        newMet = sprintf('m_S2_TOTAL_BIOMASS_%s_MEAN_PUB[c]',mode);
        newAssembly = sprintf('S2_TOTAL_BIOMASS_%s_MEAN_PUB_bm',mode);
        newSink = sprintf('S2_TOTAL_BIOMASS_%s_MEAN_PUB_SINK_bm_tx',mode);

        if ismember(newAssembly,m.rxns)
            m = removeRxns(m,{newAssembly});
        end
        if ismember(newSink,m.rxns)
            m = removeRxns(m,{newSink});
        end
        if ~ismember(newMet,m.mets)
            m = addMetabolite(m,newMet, ...
                'metName',sprintf('Sole-substrate %s mean biomass',mode), ...
                'metFormula','','Charge',0);
        end

        m = addReaction(m,newAssembly, ...
            'reactionName',sprintf('Sole-substrate %s mean BOF from TOTAL_BIOMASS.xlsx',mode), ...
            'metaboliteList',{ ...
                metsMode.protein;metsMode.carb;metsMode.lipid;metsMode.rna;metsMode.dna;newPigMet; ...
                metsMode.atp;metsMode.water;metsMode.adp;metsMode.pi;metsMode.proton;newMet}, ...
            'stoichCoeffList',[ ...
                -cProtein;-cCarb;-cLipid;-cRNA;-cDNA;-cPigment; ...
                -cATP;-cWater;cADP;cPi;cProton;1], ...
            'lowerBound',0,'upperBound',1000, ...
            'subSystem','Biomass synthesis');
        m = padConstraintColumns_pub(m);

        m = addReaction(m,newSink, ...
            'reactionName',sprintf('Sole-substrate %s mean biomass sink',mode), ...
            'metaboliteList',{newMet}, ...
            'stoichCoeffList',-1, ...
            'lowerBound',0,'upperBound',1000, ...
            'subSystem','Biomass sink');
        m = padConstraintColumns_pub(m);

        bof.mode = string(mode);
        bof.assemblyRxn = newAssembly;
        bof.sinkRxn = newSink;
        bof.biomassMet = newMet;
        bof.pigmentRxn = newPigRxn;
        bof.pigmentMet = newPigMet;
        bof.workbookReaction = workbookBOF;
        bof.workbookSheet = sheetName;
        bof.workbookPigmentReaction = workbookPigment;
        bof.sourceFormula = sourceFormula;
        bof.pigmentSourceFormula = pigmentFormula;
        bof.proteinCoeff = cProtein;
        bof.carbohydrateCoeff = cCarb;
        bof.lipidCoeff = cLipid;
        bof.rnaCoeff = cRNA;
        bof.dnaCoeff = cDNA;
        bof.pigmentCoeff = cPigment;
        bof.GAM = cATP;
        bof.pigmentComposition = [pZea,pChlA,pBeta,pPCB];

        if strcmp(mode,'AUTO')
            cfg.AUTO = bof;
        elseif strcmp(mode,'MIXO')
            cfg.MIXO = bof;
        else
            cfg.HETERO = bof;
        end

        rows(i,:) = { ...
            string(mode),string(sheetName),string(workbookBOF),string(workbookPigment), ...
            cProtein,cCarb,cLipid,cRNA,cDNA,cPigment,cATP, ...
            pZea,pChlA,pBeta,pPCB, ...
            string(pig.zea),string(pig.chla),string(pig.beta),string(pig.pcb), ...
            string(newPigRxn),string(newPigMet), ...
            string(newAssembly),string(newSink), ...
            string(metsMode.protein),string(metsMode.carb)};
    end

    auditT = cell2table(rows,'VariableNames', { ...
        'Mode','Workbook_sheet','Workbook_mean_BOF','Workbook_pigment_NOFRAC', ...
        'Protein_coeff','Carbohydrate_coeff','Lipid_coeff', ...
        'RNA_coeff','DNA_coeff','Pigment_coeff','GAM_ATP_coeff', ...
        'Pigment_zeaxanthin_fraction','Pigment_ChlA_fraction', ...
        'Pigment_beta_carotene_fraction','Pigment_PCB_fraction', ...
        'Resolved_zeaxanthin_met','Resolved_ChlA_met', ...
        'Resolved_beta_carotene_met','Resolved_PCB_met', ...
        'New_pigment_producer','New_pigment_pool_met', ...
        'New_assembly_reaction','New_sink_reaction', ...
        'Resolved_protein_met','Resolved_carbohydrate_met'});
end

function mets = resolveModeBOFMets_pub(model,mode)
    % Detailed macromolecular composition is carried by condition-specific
    % pool metabolites generated from TOTAL_BIOMASS.xlsx in the final SBML.
    % Generic names are retained only as a compatibility fallback.
    mode = upper(string(mode));
    if ~ismember(mode,["AUTO","MIXO","HETERO"])
        error('Unsupported biomass mode: %s',mode);
    end
    tag = char(mode);

    mets.protein = requireMetAnyCore_pub(model,{ ...
        sprintf('PROTEIN_TOTAL_POOL_%s_CYTO',tag),'PROTEIN_TOTAL_POOL_CYTO'});
    mets.carb = requireMetAnyCore_pub(model,{ ...
        sprintf('CARBOHYDRATE_TOTAL_POOL_%s_CYTO',tag),'CARBOHYDRATE_TOTAL_POOL_CYTO'});
    mets.lipid = requireMetAnyCore_pub(model,{ ...
        sprintf('LIPID_TOTAL_POOL_%s_CYTO',tag),'LIPID_TOTAL_POOL_CYTO'});
    mets.rna = requireMetAnyCore_pub(model,{ ...
        sprintf('RNA_TOTAL_POOL_%s_CYTO',tag),'RNA_TOTAL_POOL_CYTO'});
    mets.dna = requireMetAnyCore_pub(model,{ ...
        sprintf('DNA_TOTAL_POOL_%s_CYTO',tag),'DNA_TOTAL_POOL_CYTO'});

    mets.atp = requireMetCore_pub(model,'ATP_CYTO');
    mets.water = requireMetCore_pub(model,'WATER_CYTO');
    mets.adp = requireMetCore_pub(model,'ADP_CYTO');
    mets.pi = requireMetCore_pub(model,'PI_CYTO');
    mets.proton = requireMetCore_pub(model,'PROTON_CYTO');
end

function met = requireMetAnyCore_pub(model,candidates)
    lastMessage = "";
    for i = 1:numel(candidates)
        try
            met = requireMetCore_pub(model,candidates{i});
            return;
        catch ME
            lastMessage = string(ME.message);
        end
    end
    error('Required GSM metabolite could not be resolved. Tried: %s. Last error: %s', ...
        strjoin(candidates,', '),lastMessage);
end

function formula = readWorkbookReactionFormula_pub(biomassFile,sheetName,reactionID)
    C = readcell(biomassFile,'Sheet',sheetName);

    hitRow = [];
    hitCol = [];

    for r = 1:size(C,1)
        for c = 1:size(C,2)
            v = C{r,c};
            if ischar(v) || isstring(v)
                if strcmpi(strtrim(string(v)),string(reactionID))
                    hitRow = r;
                    hitCol = c;
                    break;
                end
            end
        end
        if ~isempty(hitRow), break; end
    end

    if isempty(hitRow)
        error('Reaction %s was not found in workbook sheet %s.',reactionID,sheetName);
    end

    formula = '';
    for c = hitCol+1:size(C,2)
        v = C{hitRow,c};
        if ischar(v) || isstring(v)
            s = strtrim(string(v));
            if strlength(s) > 0 && contains(s,'->')
                formula = char(s);
                break;
            end
        end
    end

    if isempty(formula)
        error('Formula missing for %s in workbook sheet %s.',reactionID,sheetName);
    end
end

function c = coefficientForWorkbookMet_pub(formula,metToken)
    f = strrep(char(formula),',','.');
    token = regexptranslate('escape',char(metToken));
    pat = ['([0-9]+(?:\.[0-9]+)?(?:[eE][+\-]?[0-9]+)?)\s+' token];
    t = regexp(f,pat,'tokens','once');

    if isempty(t)
        error('Could not read coefficient for %s from workbook BOF: %s', ...
              metToken,formula);
    end

    c = str2double(t{1});
    if ~isfinite(c)
        error('Invalid coefficient for %s in workbook BOF.',metToken);
    end
end

function met = requireMetCore_pub(model,canonicalToken)
    target = normalizeMetCore_pub(canonicalToken);

    ids = string(model.mets);
    normIds = strings(numel(ids),1);
    for k = 1:numel(ids)
        normIds(k) = string(normalizeMetCore_pub(ids(k)));
    end

    idx = find(normIds == string(target));

    if isempty(idx) && isfield(model,'metNames')
        names = string(model.metNames);
        normNames = strings(numel(names),1);
        for k = 1:numel(names)
            normNames(k) = string(normalizeMetCore_pub(names(k)));
        end
        idx = find(normNames == string(target));
    end

    if isempty(idx)
        labels = upper(ids);
        if isfield(model,'metNames')
            labels = labels + " " + upper(string(model.metNames));
        end
        rawToken = upper(string(canonicalToken));
        idx = find(contains(labels,rawToken));
    end

    idx = unique(idx,'stable');

    if isempty(idx)
        error('Required GSM metabolite could not be resolved: %s',canonicalToken);
    end

    if numel(idx) > 1
        exactId = find(normIds(idx) == string(target));
        if numel(exactId) == 1
            idx = idx(exactId);
        else
            candidates = strjoin(cellstr(ids(idx)),', ');
            error('Ambiguous GSM metabolite mapping for %s: %s', ...
                  canonicalToken,candidates);
        end
    end

    met = model.mets{idx(1)};
end

function s = normalizeMetCore_pub(x)
    s = upper(strtrim(char(string(x))));
    s = regexprep(s,'^M_','');
    s = regexprep(s,'\[[A-Z]+\]$','');
    s = regexprep(s,'[^A-Z0-9]','');
end

function m = padConstraintColumns_pub(m)
    if ~isfield(m,'C') || isempty(m.C)
        return;
    end
    nRxn = numel(m.rxns);
    nCol = size(m.C,2);
    if nCol < nRxn
        m.C(:,nCol+1:nRxn) = 0;
    elseif nCol > nRxn
        error('model.C has more columns than reactions after BOF construction.');
    end
end

function m = addPhotonBiomassCoupling_pub(m,photonRxn,biomassRxn,k,name)
    pidx = find(strcmp(m.rxns,photonRxn),1);
    bidx = find(strcmp(m.rxns,biomassRxn),1);
    if isempty(pidx) || isempty(bidx)
        error('Photon or biomass reaction missing for HETERO coupling.');
    end

    signUptake = uptakeFluxSign_pub(m,photonRxn);
    row = sparse(1,numel(m.rxns));
    row(pidx) = 1;
    row(bidx) = -signUptake*k;

    m = ensureConstraintFields_pub(m);
    m.C(end+1,:) = row;
    m.d(end+1,1) = 0;
    m.dsense(end+1,1) = 'E';
    m.ctrs{end+1,1} = name;
end

function m = ensureConstraintFields_pub(m)
    nRxn = numel(m.rxns);
    if ~isfield(m,'C') || isempty(m.C)
        m.C = sparse(0,nRxn);
        m.d = zeros(0,1);
        m.dsense = char(zeros(0,1));
        m.ctrs = cell(0,1);
        return;
    end

    if size(m.C,2) ~= nRxn
        error('model.C column count does not match number of reactions.');
    end
    nCtr = size(m.C,1);
    if ~isfield(m,'d') || numel(m.d) ~= nCtr
        m.d = zeros(nCtr,1);
    end
    if ~isfield(m,'dsense') || numel(m.dsense) ~= nCtr
        m.dsense = repmat('E',nCtr,1);
    end
    if ~isfield(m,'ctrs') || numel(m.ctrs) ~= nCtr
        m.ctrs = arrayfun(@(x)sprintf('existing_constraint_%d',x), ...
            (1:nCtr)','UniformOutput',false);
    end
end

function m = setRubisco_pub(m,rxn,lb,ub)
    if isempty(rxn)
        warning('Rubisco reaction was not resolved; trophic Rubisco bound was not changed.');
        return;
    end
    idx = find(strcmp(m.rxns,rxn),1);
    m.lb(idx) = lb;
    m.ub(idx) = ub;
end

function m = setObjective_pub(m,rxn)
    idx = find(strcmp(m.rxns,rxn),1);
    if isempty(idx)
        error('Objective reaction not found: %s',rxn);
    end
    if ~isfield(m,'c') || numel(m.c) ~= numel(m.rxns)
        m.c = zeros(numel(m.rxns),1);
    end
    m.c(:) = 0;
    m.c(idx) = 1;
end

function m = closeUptake_pub(m,rxn)
    if isempty(rxn), return; end
    idx = find(strcmp(m.rxns,rxn),1);
    if isempty(idx), return; end
    sgn = uptakeFluxSign_pub(m,rxn);
    if sgn < 0
        m.lb(idx) = max(m.lb(idx),0);
    else
        m.ub(idx) = min(m.ub(idx),0);
    end
end

function m = openUptake_pub(m,rxn,q,preserveOtherDirection)
    if nargin < 4, preserveOtherDirection = true; end
    idx = find(strcmp(m.rxns,rxn),1);
    if isempty(idx)
        error('Reaction not found: %s',rxn);
    end
    sgn = uptakeFluxSign_pub(m,rxn);
    q = abs(q);

    if sgn < 0
        m.lb(idx) = -q;
        if ~preserveOtherDirection
            m.ub(idx) = 0;
        end
    else
        m.ub(idx) = q;
        if ~preserveOtherDirection
            m.lb(idx) = 0;
        end
    end
end

function m = setUptakeOnly_pub(m,rxn,q)
    idx = find(strcmp(m.rxns,rxn),1);
    if isempty(idx)
        error('Reaction not found: %s',rxn);
    end
    sgn = uptakeFluxSign_pub(m,rxn);
    q = abs(q);

    if sgn < 0
        m.lb(idx) = -q;
        m.ub(idx) = 0;
    else
        m.lb(idx) = 0;
        m.ub(idx) = q;
    end
end

function sgn = uptakeFluxSign_pub(model,rxn)
    idx = find(strcmp(model.rxns,rxn),1);
    if isempty(idx)
        error('Boundary reaction not found: %s',rxn);
    end

    s = full(model.S(:,idx));
    nz = find(abs(s) > 1e-12);
    if numel(nz) ~= 1
        error('Expected one-metabolite boundary reaction for uptake sign: %s',rxn);
    end

    if s(nz) < 0
        sgn = -1;
    else
        sgn = +1;
    end
end

function q = uptakeMagnitudeFromSolution_pub(model,rxn,x)
    idx = find(strcmp(model.rxns,rxn),1);
    if isempty(idx)
        q = NaN;
        return;
    end
    sgn = uptakeFluxSign_pub(model,rxn);
    if sgn < 0
        q = max(0,-x(idx));
    else
        q = max(0,x(idx));
    end
end

function r = requireRxn_pub(model,candidates)
    r = '';
    for k = 1:numel(candidates)
        r = resolveRxn_pub(model,candidates{k},false);
        if ~isempty(r), return; end
    end
    error('Required reaction not found: %s',strjoin(candidates,' / '));
end

function r = resolveRxn_pub(model,q,warnIfMissing)
    if nargin < 3, warnIfMissing = false; end
    r = '';
    q = char(string(q));

    idx = find(strcmp(model.rxns,q),1);
    if isempty(idx) && isfield(model,'rxnNames')
        idx = find(strcmpi(string(model.rxnNames),string(q)),1);
    end

    if isempty(idx)
        nq = norm_pub(q);
        nr = string(cellfun(@norm_pub,model.rxns,'UniformOutput',false));
        idx = find(nr == string(nq),1);
    end

    if ~isempty(idx)
        r = model.rxns{idx};
    elseif warnIfMissing
        warning('Reaction could not be resolved: %s',q);
    end
end

function s = norm_pub(x)
    raw = upper(strtrim(char(string(x))));
    if startsWith(raw,'R_')
        raw = raw(3:end);
    end
    s = upper(regexprep(raw,'[^A-Za-z0-9]',''));
end

function [nOpen,openRxns] = mappedSubstrateUptakeState_pub(m,cfg)
    openRxns = strings(0,1);
    for k = 1:numel(cfg.allMappedCarbonRxns)
        r = cfg.allMappedCarbonRxns{k};
        idx = find(strcmp(m.rxns,r),1);
        if isempty(idx), continue; end
        sgn = uptakeFluxSign_pub(m,r);
        if sgn < 0
            cap = max(0,-m.lb(idx));
        else
            cap = max(0,m.ub(idx));
        end
        if cap > 1e-12
            openRxns(end+1,1) = string(r); %#ok<AGROW>
        end
    end
    openRxns = unique(openRxns,'stable');
    nOpen = numel(openRxns);
end

function assertMappedSubstrateState_pub(m,selectedRxn,cfg,label)
    [nOpen,openRxns] = mappedSubstrateUptakeState_pub(m,cfg);
    selectedRxn = string(selectedRxn);

    if strlength(selectedRxn)==0
        if nOpen ~= 0
            error('%s: expected zero mapped organic substrates open, found %d: %s', ...
                label,nOpen,strjoin(cellstr(openRxns),', '));
        end
        return;
    end

    if nOpen ~= 1 || ~any(openRxns==selectedRxn)
        error(['%s: exactly one mapped substrate must be open. Selected=%s; ' ...
               'open count=%d; open=%s'], ...
               label,selectedRxn,nOpen,strjoin(cellstr(openRxns),', '));
    end
end

function tf = allOldBiomassOutputsClosed_pub(m,cfg)
    tf = true;
    if ~isfield(cfg,'preexistingClosedBiomassRxns'), return; end
    for k = 1:numel(cfg.preexistingClosedBiomassRxns)
        r = cfg.preexistingClosedBiomassRxns{k};
        idx = find(strcmp(m.rxns,r),1);
        if isempty(idx), continue; end
        if abs(m.lb(idx))>1e-12 || abs(m.ub(idx))>1e-12
            tf = false;
            return;
        end
    end
end

function tf = onlySelectedBiomassSinkOpen_pub(m,selectedSink)
    ids = string(m.rxns);
    raw = lower(ids);
    nr = string(cellfun(@norm_pub,m.rxns,'UniformOutput',false));
    idx = find(contains(nr,'BMTX') | endsWith(raw,"_bm_tx"));
    open = strings(0,1);

    for j = idx(:)'
        if abs(m.lb(j))>1e-12 || abs(m.ub(j))>1e-12
            open(end+1,1)=ids(j); %#ok<AGROW>
        end
    end

    open = unique(open,'stable');
    tf = numel(open)==1 && any(open==string(selectedSink));
end

function assertBiomassOutputState_pub(m,selectedSink,cfg,label)
    if ~allOldBiomassOutputsClosed_pub(m,cfg)
        error('%s: at least one pre-existing bm_tx/TOTAL_BIOMASS reaction is not 0;0.',label);
    end
    if ~onlySelectedBiomassSinkOpen_pub(m,selectedSink)
        error('%s: selected BOF sink is not the only open bm_tx reaction.',label);
    end
end

function [lb,ub] = boundsOf_pub(m,rxn)
    idx = find(strcmp(m.rxns,rxn),1);
    if isempty(idx)
        lb = NaN; ub = NaN;
    else
        lb = m.lb(idx); ub = m.ub(idx);
    end
end

function A = buildConditionAudit_pub(modelAUTO,modelMIXO,modelHETERO,cfg)
    modes = ["AUTO";"MIXO";"HETERO"];
    models = {modelAUTO;modelMIXO;modelHETERO};
    sinks = {cfg.AUTO.sinkRxn;cfg.MIXO.sinkRxn;cfg.HETERO.sinkRxn};

    n = 3;
    NGAM_LB=nan(n,1); NGAM_UB=nan(n,1);
    Photon_LB=nan(n,1); Photon_UB=nan(n,1);
    CO2_LB=nan(n,1); CO2_UB=nan(n,1);
    O2_LB=nan(n,1); O2_UB=nan(n,1);
    O2Dummy_LB=nan(n,1); O2Dummy_UB=nan(n,1);
    Rubisco_LB=nan(n,1); Rubisco_UB=nan(n,1);
    Open_mapped_organic_substrates=nan(n,1);
    Open_carbon_boundary_uptakes=nan(n,1);
    Open_carbon_boundary_list=strings(n,1);
    Old_bm_tx_all_closed=false(n,1);
    Only_selected_new_bm_tx_open=false(n,1);
    Selected_BOF_sink=strings(n,1);

    for i=1:n
        m=models{i};
        [NGAM_LB(i),NGAM_UB(i)] = boundsOf_pub(m,cfg.ngamRxn);
        [Photon_LB(i),Photon_UB(i)] = boundsOf_pub(m,cfg.photonRxn);
        [CO2_LB(i),CO2_UB(i)] = boundsOf_pub(m,cfg.co2Rxn);
        [O2_LB(i),O2_UB(i)] = boundsOf_pub(m,cfg.o2Rxn);
        [O2Dummy_LB(i),O2Dummy_UB(i)] = boundsOf_pub(m,cfg.o2DummyRxn);
        [Rubisco_LB(i),Rubisco_UB(i)] = boundsOf_pub(m,cfg.rubiscoRxn);
        [Open_mapped_organic_substrates(i),~] = mappedSubstrateUptakeState_pub(m,cfg);
        [Open_carbon_boundary_uptakes(i),openC] = carbonBoundaryUptakeState_pub(m,cfg);
        Open_carbon_boundary_list(i) = strjoin(cellstr(openC),'; ');
        Old_bm_tx_all_closed(i) = allOldBiomassOutputsClosed_pub(m,cfg);
        Only_selected_new_bm_tx_open(i) = onlySelectedBiomassSinkOpen_pub(m,sinks{i});
        Selected_BOF_sink(i) = string(sinks{i});
    end

    Condition_rule = [ ...
        "CO2 only; 6 C-mmol carbon; light non-binding then minimized"; ...
        "CO2 available + finite light + exactly one organic substrate during each run"; ...
        "no external inorganic carbon + exactly one organic substrate + calibrated small light"];

    A = table(modes,Condition_rule,Selected_BOF_sink, ...
        NGAM_LB,NGAM_UB,Photon_LB,Photon_UB,CO2_LB,CO2_UB,O2_LB,O2_UB, ...
        O2Dummy_LB,O2Dummy_UB,Rubisco_LB,Rubisco_UB,Open_mapped_organic_substrates, ...
        Open_carbon_boundary_uptakes,Open_carbon_boundary_list, ...
        Old_bm_tx_all_closed,Only_selected_new_bm_tx_open, ...
        'VariableNames',{'Mode','Condition_rule','Selected_BOF_sink', ...
        'NGAM_LB','NGAM_UB','Photon_LB','Photon_UB','CO2_LB','CO2_UB', ...
        'O2_LB','O2_UB','O2_Dummy_LB','O2_Dummy_UB', ...
        'Rubisco_LB','Rubisco_UB', ...
        'Open_mapped_organic_substrates_at_base', ...
        'Open_carbon_boundary_uptakes_at_base','Open_carbon_boundary_list', ...
        'Preexisting_bm_tx_all_0_0','Only_selected_new_bm_tx_open'});
end

function T = stratifiedSampleValidation_pub( ...
    modelAUTO,modelMIXO,modelHETERO,substrateName,resolvedExchange, ...
    carbonAtoms,maxSubstrateUptake,expectedGrowth, ...
    autoRun,mixoRuns,heteroRuns,cfg)

    rng(55,'twister');
    valid = find(strlength(resolvedExchange)>0);

    mandatory = [];
    for nm = ["D-Glucose","D-Xylose","Glycerol"]
        ii = find(strcmpi(strtrim(substrateName),nm),1);
        if ~isempty(ii) && any(valid==ii)
            mandatory(end+1) = ii; %#ok<AGROW>
        end
    end
    pool = setdiff(valid,mandatory,'stable');
    nr = min(2,numel(pool));
    if nr>0
        randomPick = pool(randperm(numel(pool),nr));
    else
        randomPick = [];
    end
    pick = unique([mandatory(:);randomPick(:)],'stable');

    nRows = 1 + 2*numel(pick);
    Sample=strings(nRows,1); Mode=strings(nRows,1); Substrate=strings(nRows,1);
    Selected_exchange=strings(nRows,1); Open_carbon_boundary_count=nan(nRows,1);
    First_pass_mu=nan(nRows,1); Repeat_mu=nan(nRows,1); Abs_mu_difference=nan(nRows,1);
    Preexisting_bm_tx_all_0_0=false(nRows,1);
    Only_selected_new_bm_tx_open=false(nRows,1);
    Carbon_boundary_rule_pass=false(nRows,1);
    Constraint_pass=false(nRows,1);
    Reproducibility_pass=false(nRows,1);
    Biological_growth=false(nRows,1);
    Experimental_agreement=strings(nRows,1);
    Overall_validation=strings(nRows,1);

    [repAuto,~] = optimizeAutoAdaptive_pub(modelAUTO,cfg.AUTO.sinkRxn,cfg);
    Sample(1)="AUTO_ADAPTIVE_REPEAT"; Mode(1)="AUTO"; Substrate(1)="CO2_ONLY";
    [Open_carbon_boundary_count(1),~]=carbonBoundaryUptakeState_pub(modelAUTO,cfg);
    First_pass_mu(1)=autoRun.Mu_h; Repeat_mu(1)=repAuto.Mu_h;
    Abs_mu_difference(1)=abs(autoRun.Mu_h-repAuto.Mu_h);
    Preexisting_bm_tx_all_0_0(1)=allOldBiomassOutputsClosed_pub(modelAUTO,cfg);
    Only_selected_new_bm_tx_open(1)=onlySelectedBiomassSinkOpen_pub(modelAUTO,cfg.AUTO.sinkRxn);
    Carbon_boundary_rule_pass(1)=Open_carbon_boundary_count(1)==1;
    Constraint_pass(1)=Carbon_boundary_rule_pass(1) && ...
        Preexisting_bm_tx_all_0_0(1) && Only_selected_new_bm_tx_open(1);
    Reproducibility_pass(1)=Abs_mu_difference(1)<1e-9;
    Biological_growth(1)=repAuto.PredictedGrowth;
    Experimental_agreement(1)="NOT_SCORED";
    if Constraint_pass(1) && Reproducibility_pass(1)
        if Biological_growth(1)
            Overall_validation(1)="MECHANICS_PASS_AUTO_GROWTH";
        else
            Overall_validation(1)="MECHANICS_PASS_AUTO_NO_GROWTH";
        end
    else
        Overall_validation(1)="MECHANICS_FAIL";
    end

    row=1;
    for p=1:numel(pick)
        i=pick(p); subRxn=char(resolvedExchange(i)); q=maxSubstrateUptake(i);
        for modeName=["MIXO","HETERO"]
            row=row+1;
            if modeName=="MIXO"
                m=modelMIXO; sink=cfg.MIXO.sinkRxn; first=mixoRuns(i);
                allowed={cfg.co2Rxn,subRxn};
            else
                m=modelHETERO; sink=cfg.HETERO.sinkRxn; first=heteroRuns(i);
                allowed={subRxn};
            end

            m=openUptake_pub(m,subRxn,q,false);
            assertCarbonBoundaryState_pub(m,allowed,cfg,char(modeName+"_SAMPLE"));
            assertBiomassOutputState_pub(m,sink,cfg,char(modeName+"_SAMPLE"));
            repeat=optimizeState_pub(m,sink,subRxn,carbonAtoms(i),cfg, ...
                char(modeName),substrateName(i));

            if any(i==mandatory)
                Sample(row)="CONTROL_"+substrateName(i);
            else
                Sample(row)="RANDOM_"+substrateName(i);
            end

            Mode(row)=modeName; Substrate(row)=substrateName(i);
            Selected_exchange(row)=string(subRxn);
            [Open_carbon_boundary_count(row),~]=carbonBoundaryUptakeState_pub(m,cfg);
            First_pass_mu(row)=first.Mu_h; Repeat_mu(row)=repeat.Mu_h;
            Abs_mu_difference(row)=abs(first.Mu_h-repeat.Mu_h);
            Preexisting_bm_tx_all_0_0(row)=allOldBiomassOutputsClosed_pub(m,cfg);
            Only_selected_new_bm_tx_open(row)=onlySelectedBiomassSinkOpen_pub(m,sink);
            Carbon_boundary_rule_pass(row)= ...
                Open_carbon_boundary_count(row)==numel(unique(string(allowed)));

            Constraint_pass(row)=Carbon_boundary_rule_pass(row) && ...
                Preexisting_bm_tx_all_0_0(row) && ...
                Only_selected_new_bm_tx_open(row);

            Reproducibility_pass(row)=first.Status==repeat.Status && ...
                Abs_mu_difference(row)<1e-9;

            Biological_growth(row)=repeat.PredictedGrowth;

            if isnan(expectedGrowth(i))
                Experimental_agreement(row)="NOT_SCORED";
            elseif logical(expectedGrowth(i))==Biological_growth(row)
                Experimental_agreement(row)="MATCH";
            else
                Experimental_agreement(row)="MISMATCH";
            end

            if ~Constraint_pass(row) || ~Reproducibility_pass(row)
                Overall_validation(row)="MECHANICS_FAIL";
            elseif Experimental_agreement(row)=="MISMATCH"
                Overall_validation(row)="MECHANICS_PASS_PHENOTYPE_MISMATCH";
            elseif Experimental_agreement(row)=="MATCH"
                Overall_validation(row)="MECHANICS_PASS_PHENOTYPE_MATCH";
            elseif Biological_growth(row)
                Overall_validation(row)="MECHANICS_PASS_NOT_SCORED_GROWTH";
            else
                Overall_validation(row)="MECHANICS_PASS_NOT_SCORED_NO_GROWTH";
            end
        end
    end

    T=table(Sample,Mode,Substrate,Selected_exchange,Open_carbon_boundary_count, ...
        First_pass_mu,Repeat_mu,Abs_mu_difference,Preexisting_bm_tx_all_0_0, ...
        Only_selected_new_bm_tx_open,Carbon_boundary_rule_pass,Constraint_pass, ...
        Reproducibility_pass,Biological_growth,Experimental_agreement, ...
        Overall_validation);

    if ~all(T.Constraint_pass & T.Reproducibility_pass)
        error(['Sample validation found a constraint or reproducibility ' ...
               'failure. Biological phenotype mismatches do NOT trigger this error.']);
    end
end

function [carbonRxns,auditT] = identifyCarbonBoundaryRxns_pub(model,cfg)
    nRxn=numel(model.rxns);
    rid=strings(nRxn,1); mid=strings(nRxn,1); mname=strings(nRxn,1);
    formula=strings(nRxn,1); nC=nan(nRxn,1); initiallyOpen=false(nRxn,1);
    keep=false(nRxn,1);

    for j=1:nRxn
        nz=find(abs(full(model.S(:,j)))>1e-12);
        if numel(nz)~=1, continue; end
        mi=nz(1); rid(j)=string(model.rxns{j}); mid(j)=string(model.mets{mi});
        if isfield(model,'metNames') && numel(model.metNames)>=mi
            mname(j)=string(model.metNames{mi});
        end
        if isfield(model,'metFormulas') && numel(model.metFormulas)>=mi
            formula(j)=string(model.metFormulas{mi});
        end
        nC(j)=formulaCarbonCount_pub(formula(j));
        if isnan(nC(j)) && (any(strcmp(model.rxns{j},cfg.allMappedCarbonRxns)) || ...
                strcmp(model.rxns{j},cfg.co2Rxn))
            nC(j)=1; % force inclusion when formula metadata is absent
        end
        if isfinite(nC(j)) && nC(j)>0
            keep(j)=true;
            sgn=uptakeFluxSign_pub(model,model.rxns{j});
            if sgn<0, initiallyOpen(j)=model.lb(j)<-1e-12;
            else, initiallyOpen(j)=model.ub(j)>1e-12; end
        end
    end

    idx=find(keep); carbonRxns=rid(idx);
    auditT=table(rid(idx),mid(idx),mname(idx),formula(idx),nC(idx), ...
        model.lb(idx),model.ub(idx),initiallyOpen(idx), ...
        'VariableNames',{'Reaction_ID','Metabolite_ID','Metabolite_name','Formula', ...
        'Carbon_atoms_from_formula','Original_LB','Original_UB','Uptake_initially_open'});
end

function nC = formulaCarbonCount_pub(formula)
    nC=NaN; f=char(strtrim(string(formula)));
    if isempty(f), return; end
    toks=regexp(f,'([A-Z][a-z]?)([0-9]*)','tokens');
    if isempty(toks), return; end
    nC=0;
    for k=1:numel(toks)
        if strcmp(toks{k}{1},'C')
            if isempty(toks{k}{2}), nC=nC+1;
            else, nC=nC+str2double(toks{k}{2}); end
        end
    end
end

function [nOpen,openRxns] = carbonBoundaryUptakeState_pub(m,cfg)
    openRxns=strings(0,1);
    for k=1:numel(cfg.carbonBoundaryRxns)
        r=cfg.carbonBoundaryRxns{k}; idx=find(strcmp(m.rxns,r),1);
        if isempty(idx), continue; end
        sgn=uptakeFluxSign_pub(m,r);
        if sgn<0, cap=max(0,-m.lb(idx)); else, cap=max(0,m.ub(idx)); end
        if cap>1e-12, openRxns(end+1,1)=string(r); end %#ok<AGROW>
    end
    openRxns=unique(openRxns,'stable'); nOpen=numel(openRxns);
end

function assertCarbonBoundaryState_pub(m,allowedRxns,cfg,label)
    [~,openRxns]=carbonBoundaryUptakeState_pub(m,cfg);
    allowed=unique(string(allowedRxns),'stable'); allowed=allowed(strlength(allowed)>0);
    if numel(openRxns)~=numel(allowed) || ~all(ismember(openRxns,allowed)) || ...
            ~all(ismember(allowed,openRxns))
        error(['%s carbon-source violation. Allowed={%s}; open={%s}.'], ...
            label,strjoin(cellstr(allowed),', '),strjoin(cellstr(openRxns),', '));
    end
end

function [alpha,k,T] = calibrateHeteroPhoton_pub(modelHET,glcRxn,qGlc,cfg)
    m=modelHET; m=openUptake_pub(m,glcRxn,qGlc,false);
    assertCarbonBoundaryState_pub(m,{glcRxn},cfg,'HETERO_PHOTON_CALIBRATION');
    bidx=find(strcmp(m.rxns,cfg.HETERO.sinkRxn),1);
    pidx=find(strcmp(m.rxns,cfg.photonRxn),1);
    gidx=find(strcmp(m.rxns,cfg.HETERO.pigmentRxn),1);
    if isempty(bidx)||isempty(pidx)||isempty(gidx), error('HETERO calibration mapping missing.'); end
    m.lb(bidx)=cfg.heteroPhotonCalibrationMu; m.ub(bidx)=cfg.heteroPhotonCalibrationMu;
    m.c(:)=0; m.c(pidx)=1; sgn=uptakeFluxSign_pub(m,cfg.photonRxn);
    if sgn<0, sol=optimizeCbModel(m,'max'); else, sol=optimizeCbModel(m,'min'); end
    if ~isfield(sol,'stat')||sol.stat~=1||isempty(sol.x)
        error('HETERO photon calibration infeasible at mu=%.4g.',cfg.heteroPhotonCalibrationMu);
    end
    minPhoton=uptakeMagnitudeFromSolution_pub(m,cfg.photonRxn,sol.x);
    pigmentFlux=sol.x(gidx);
    if pigmentFlux<=1e-12, error('HETERO pigment-pool flux is zero in calibration.'); end
    alpha=minPhoton/pigmentFlux; k=minPhoton/cfg.heteroPhotonCalibrationMu;
    T=table(cfg.heteroPhotonCalibrationMu,qGlc,minPhoton,pigmentFlux,alpha,k, ...
        cfg.expectedPhotonPerPigment,abs(alpha-cfg.expectedPhotonPerPigment), ...
        'VariableNames',{'Calibration_mu_h','Glucose_uptake_cap_mmol', ...
        'Minimum_photon_uptake','Pigment_pool_flux','Photon_per_pigment', ...
        'Photon_per_biomass','Expected_photon_per_pigment','Absolute_difference'});
end

function [run,T] = optimizeAutoAdaptive_pub(modelAUTO,biomassRxn,cfg)
    run=optimizeState_pub(modelAUTO,biomassRxn,'',NaN,cfg,'AUTO','CO2_ONLY');
    native=modelAUTO; native=setUptakeOnly_pub(native,cfg.photonRxn,cfg.autoNativePhotonDiagnostic);
    nativeRun=optimizeState_pub(native,biomassRxn,'',NaN,cfg,'AUTO_NATIVE_0P2','CO2_ONLY');
    minPhoton=NaN; maxPhoton=NaN;
    if run.PredictedGrowth
        target=cfg.autoPhotonCalibrationFraction*run.Mu_h;
        [minPhoton,maxPhoton]=substrateUseRangeAtBiomass_pub( ...
            modelAUTO,biomassRxn,cfg.photonRxn,target);
        run.PhotonUptake=minPhoton;
    end
    T=table(["Native photon 0.2";"AUTO photon cap 20"], ...
        [nativeRun.Mu_h;run.Mu_h],[nativeRun.PredictedGrowth;run.PredictedGrowth], ...
        [nativeRun.PhotonUptake;minPhoton], ...
        [cfg.autoNativePhotonDiagnostic;cfg.autoPhotonProbeMax], ...
        [cfg.autoCO2CarbonCap;cfg.autoCO2CarbonCap], ...
        [NaN;maxPhoton], ...
        'VariableNames',{'AUTO_test','Mu_h','Growth_feasible', ...
        'Photon_uptake_or_minimum_at_99pct_mu','Photon_cap','CO2_cap_mmol', ...
        'Maximum_photon_at_99pct_mu'});
end

function m = removeConstraintByName_pub(m,name)
    if ~isfield(m,'ctrs') || isempty(m.ctrs), return; end
    idx=find(strcmp(string(m.ctrs),string(name)));
    if isempty(idx), return; end
    m.C(idx,:)=[]; m.d(idx)=[]; m.dsense(idx)=[]; m.ctrs(idx)=[];
end

function m = removePigmentTermFromBOF_pub(m,bof)
    ridx=find(strcmp(m.rxns,bof.assemblyRxn),1);
    midx=find(strcmp(m.mets,bof.pigmentMet),1);
    if ~isempty(ridx)&&~isempty(midx), m.S(midx,ridx)=0; end
end

function T = buildFailureDiagnostics_pub( ...
    modelAUTO,modelMIXO,modelHETERO,substrateName,resolvedExchange, ...
    carbonAtoms,maxSubstrateUptake,autoRun,mixoRuns,heteroRuns,cfg)

    rows={};
    if ~autoRun.PredictedGrowth
        r0=optimizeState_pub(modelAUTO,cfg.AUTO.sinkRxn,'',NaN,cfg,'AUTO','CO2_ONLY');
        mn=modelAUTO; ni=find(strcmp(mn.rxns,cfg.ngamRxn),1); mn.lb(ni)=0; mn.ub(ni)=0;
        rn=optimizeState_pub(mn,cfg.AUTO.sinkRxn,'',NaN,cfg,'AUTO','CO2_ONLY');
        mp=removePigmentTermFromBOF_pub(modelAUTO,cfg.AUTO);
        rp=optimizeState_pub(mp,cfg.AUTO.sinkRxn,'',NaN,cfg,'AUTO','CO2_ONLY');
        rows(end+1,:)={'AUTO','CO2_ONLY','BASE_ADAPTIVE',r0.Mu_h,char(r0.Status)}; %#ok<AGROW>
        rows(end+1,:)={'AUTO','CO2_ONLY','NGAM_ZERO',rn.Mu_h,char(rn.Status)}; %#ok<AGROW>
        rows(end+1,:)={'AUTO','CO2_ONLY','NO_PIGMENT_TERM',rp.Mu_h,char(rp.Status)}; %#ok<AGROW>
    end

    for i=1:numel(substrateName)
        if strlength(resolvedExchange(i))==0, continue; end
        subRxn=char(resolvedExchange(i)); q=maxSubstrateUptake(i);
        for modeName=["MIXO","HETERO"]
            if modeName=="MIXO"
                first=mixoRuns(i); baseM=modelMIXO; sink=cfg.MIXO.sinkRxn; bof=cfg.MIXO;
            else
                first=heteroRuns(i); baseM=modelHETERO; sink=cfg.HETERO.sinkRxn; bof=cfg.HETERO;
            end
            if first.PredictedGrowth, continue; end
            m=baseM; m=openUptake_pub(m,subRxn,q,false);
            r0=optimizeState_pub(m,sink,subRxn,carbonAtoms(i),cfg,char(modeName),substrateName(i));
            m10=baseM; m10=openUptake_pub(m10,subRxn,q*cfg.failureProbeSubstrateMultiplier,false);
            r10=optimizeState_pub(m10,sink,subRxn,carbonAtoms(i),cfg,char(modeName),substrateName(i));
            mn=m; ni=find(strcmp(mn.rxns,cfg.ngamRxn),1); mn.lb(ni)=0; mn.ub(ni)=0;
            rn=optimizeState_pub(mn,sink,subRxn,carbonAtoms(i),cfg,char(modeName),substrateName(i));
            mp=removePigmentTermFromBOF_pub(m,bof);
            if modeName=="HETERO"
                mp=removeConstraintByName_pub(mp,'HETERO_PHOTON_BIOMASS_COUPLING_PUB');
            end
            rp=optimizeState_pub(mp,sink,subRxn,carbonAtoms(i),cfg,char(modeName),substrateName(i));
            rows(end+1,:)={char(modeName),char(substrateName(i)),'BASE',r0.Mu_h,char(r0.Status)}; %#ok<AGROW>
            rows(end+1,:)={char(modeName),char(substrateName(i)), ...
                sprintf('SUBSTRATE_%gx',cfg.failureProbeSubstrateMultiplier),r10.Mu_h,char(r10.Status)}; %#ok<AGROW>
            rows(end+1,:)={char(modeName),char(substrateName(i)),'NGAM_ZERO',rn.Mu_h,char(rn.Status)}; %#ok<AGROW>
            rows(end+1,:)={char(modeName),char(substrateName(i)),'NO_PIGMENT_TERM',rp.Mu_h,char(rp.Status)}; %#ok<AGROW>
        end
    end
    if isempty(rows)
        T=table(strings(0,1),strings(0,1),strings(0,1),zeros(0,1),strings(0,1), ...
            'VariableNames',{'Mode','Substrate','Probe','Mu_h','Status'});
    else
        T=cell2table(rows,'VariableNames',{'Mode','Substrate','Probe','Mu_h','Status'});
    end
end


function T = autoSingleBoundaryRescue_pub(modelAUTO,biomassRxn,carbonBoundaryAudit,cfg)

    rxns = string(cfg.carbonBoundaryRxns(:));
    rxns = rxns(rxns~=string(cfg.co2Rxn));

    n=numel(rxns);
    Reaction_ID=strings(n,1);
    Metabolite_ID=strings(n,1);
    Metabolite_name=strings(n,1);
    Formula=strings(n,1);
    Carbon_atoms=nan(n,1);
    Rescue_uptake_cap_mmol=nan(n,1);
    Mu_h=zeros(n,1);
    Rescues_growth=false(n,1);
    Actual_rescue_uptake_mmol=nan(n,1);
    Actual_CO2_uptake_mmol=nan(n,1);
    Photon_uptake=nan(n,1);
    Open_carbon_boundary_count=nan(n,1);
    Constraint_pass=false(n,1);

    for i=1:n
        r=char(rxns(i));
        Reaction_ID(i)=rxns(i);

        j=find(strcmp(string(carbonBoundaryAudit.Reaction_ID),rxns(i)),1);
        if ~isempty(j)
            if ismember('Metabolite_ID',carbonBoundaryAudit.Properties.VariableNames)
                Metabolite_ID(i)=string(carbonBoundaryAudit.Metabolite_ID(j));
            end
            if ismember('Metabolite_name',carbonBoundaryAudit.Properties.VariableNames)
                Metabolite_name(i)=string(carbonBoundaryAudit.Metabolite_name(j));
            end
            if ismember('Formula',carbonBoundaryAudit.Properties.VariableNames)
                Formula(i)=string(carbonBoundaryAudit.Formula(j));
            end
            if ismember('Carbon_atoms_from_formula',carbonBoundaryAudit.Properties.VariableNames)
                Carbon_atoms(i)=carbonBoundaryAudit.Carbon_atoms_from_formula(j);
            end
        end

        if ~isfinite(Carbon_atoms(i)) || Carbon_atoms(i)<=0
            Carbon_atoms(i)=1;
        end

        q=cfg.autoBoundaryRescueCarbon/Carbon_atoms(i);
        Rescue_uptake_cap_mmol(i)=q;

        m=modelAUTO;
        m=openUptake_pub(m,r,q,true);
        allowed={cfg.co2Rxn,r};

        try
            assertCarbonBoundaryState_pub(m,allowed,cfg,'AUTO_BOUNDARY_RESCUE');
            assertBiomassOutputState_pub(m,biomassRxn,cfg,'AUTO_BOUNDARY_RESCUE');
            Constraint_pass(i)=true;
        catch
            Constraint_pass(i)=false;
        end

        if ~Constraint_pass(i)
            continue;
        end

        rr=optimizeState_pub(m,biomassRxn,r,Carbon_atoms(i),cfg, ...
            'AUTO_RESCUE',Reaction_ID(i));

        Mu_h(i)=rr.Mu_h;
        Rescues_growth(i)=rr.PredictedGrowth;
        Actual_rescue_uptake_mmol(i)=rr.SubstrateUptake_mmol;
        Actual_CO2_uptake_mmol(i)=rr.CO2Uptake_mmol;
        Photon_uptake(i)=rr.PhotonUptake;
        [Open_carbon_boundary_count(i),~]=carbonBoundaryUptakeState_pub(m,cfg);
    end

    T=table(Reaction_ID,Metabolite_ID,Metabolite_name,Formula,Carbon_atoms, ...
        Rescue_uptake_cap_mmol,Mu_h,Rescues_growth, ...
        Actual_rescue_uptake_mmol,Actual_CO2_uptake_mmol,Photon_uptake, ...
        Open_carbon_boundary_count,Constraint_pass);

    if ~isempty(T)
        T=sortrows(T,{'Rescues_growth','Mu_h'},{'descend','descend'});
    end
end

function T = autoBOFComponentDiagnostic_pub(modelAUTO,bof,cfg)

    components = ["PROTEIN";"CARBOHYDRATE";"LIPID";"RNA";"DNA";"PIGMENT";"GAM"];
    n=numel(components);

    Mu_h=zeros(n,1);
    Rescues_growth=false(n,1);
    Diagnostic_metabolites=strings(n,1);

    for i=1:n
        m=modelAUTO;
        switch components(i)
            case "PROTEIN"
                mets={cfg.resolvedBOFMets.protein};
            case "CARBOHYDRATE"
                mets={cfg.resolvedBOFMets.carb};
            case "LIPID"
                mets={cfg.resolvedBOFMets.lipid};
            case "RNA"
                mets={cfg.resolvedBOFMets.rna};
            case "DNA"
                mets={cfg.resolvedBOFMets.dna};
            case "PIGMENT"
                mets={bof.pigmentMet};
            case "GAM"
                mets={cfg.resolvedBOFMets.atp,cfg.resolvedBOFMets.water, ...
                      cfg.resolvedBOFMets.adp,cfg.resolvedBOFMets.pi, ...
                      cfg.resolvedBOFMets.proton};
        end

        Diagnostic_metabolites(i)=strjoin(string(mets),"; ");
        m=zeroAssemblyStoichForMets_pub(m,bof.assemblyRxn,mets);
        r=optimizeState_pub(m,bof.sinkRxn,'',NaN,cfg, ...
            'AUTO_BOF_RELAX',components(i));

        Mu_h(i)=r.Mu_h;
        Rescues_growth(i)=r.PredictedGrowth;
    end

    T=table(components,Diagnostic_metabolites,Mu_h,Rescues_growth, ...
        'VariableNames',{'Removed_BOF_component','Affected_model_metabolites', ...
        'Mu_h_after_single_component_removal','Rescues_growth'});
end

function m = zeroAssemblyStoichForMets_pub(m,assemblyRxn,metList)
    ridx=find(strcmp(m.rxns,assemblyRxn),1);
    if isempty(ridx)
        error('Assembly reaction not found: %s',assemblyRxn);
    end
    for k=1:numel(metList)
        midx=find(strcmp(m.mets,metList{k}),1);
        if ~isempty(midx)
            m.S(midx,ridx)=0;
        end
    end
end

function T = autoPoolProductionDiagnostic_pub(modelAUTO,bof,cfg)

    component = ["PROTEIN";"CARBOHYDRATE";"LIPID";"RNA";"DNA";"PIGMENT"];
    met = string({cfg.resolvedBOFMets.protein;cfg.resolvedBOFMets.carb; ...
        cfg.resolvedBOFMets.lipid;cfg.resolvedBOFMets.rna; ...
        cfg.resolvedBOFMets.dna;bof.pigmentMet});

    scenarios=["NGAM_FIXED";"NGAM_ZERO"];
    n=numel(component)*numel(scenarios);

    Component=strings(n,1);
    Metabolite=strings(n,1);
    Scenario=strings(n,1);
    Max_pool_demand_flux=nan(n,1);
    Demand_feasible=false(n,1);
    Status=strings(n,1);

    row=0;
    for i=1:numel(component)
        for s=1:numel(scenarios)
            row=row+1;
            m=modelAUTO;

            ai=find(strcmp(m.rxns,bof.assemblyRxn),1);
            bi=find(strcmp(m.rxns,bof.sinkRxn),1);
            m.lb(ai)=0; m.ub(ai)=0;
            m.lb(bi)=0; m.ub(bi)=0;

            if scenarios(s)=="NGAM_ZERO"
                ni=find(strcmp(m.rxns,cfg.ngamRxn),1);
                m.lb(ni)=0; m.ub(ni)=0;
            end

            dm=sprintf('DIAG_DM_%s_%s',component(i),scenarios(s));
            if ismember(dm,m.rxns)
                m=removeRxns(m,{dm});
            end

            m=addReaction(m,dm, ...
                'metaboliteList',{char(met(i))}, ...
                'stoichCoeffList',-1, ...
                'lowerBound',0,'upperBound',cfg.autoDiagnosticDemandUB, ...
                'subSystem','Diagnostic demand');
            m=padConstraintColumns_pub(m);
            m=setObjective_pub(m,dm);

            sol=optimizeCbModel(m,'max');

            Component(row)=component(i);
            Metabolite(row)=met(i);
            Scenario(row)=scenarios(s);

            if isfield(sol,'stat') && sol.stat==1 && ~isempty(sol.f)
                Max_pool_demand_flux(row)=max(0,sol.f);
                Demand_feasible(row)=Max_pool_demand_flux(row)>cfg.autoDiagnosticFluxTol;
                Status(row)="OPTIMAL";
            else
                Max_pool_demand_flux(row)=0;
                Demand_feasible(row)=false;
                Status(row)="INFEASIBLE";
            end
        end
    end

    T=table(Component,Metabolite,Scenario,Max_pool_demand_flux, ...
        Demand_feasible,Status);
end

function T = autoPathwayDiagnostic_pub(modelAUTO,cfg,runFull)

    categories = { ...
        'LIGHT_INPUT',                 {'PHOTON'}; ...
        'PHOTOSYSTEM_II',              {'PHOTOSYSTEMII','PSII'}; ...
        'PHOTOSYSTEM_I',               {'PHOTOSYSTEMI','PSI'}; ...
        'PLASTOCYANIN_B6F',            {'PLASTOCYANIN','CYTOCHROMEB6'}; ...
        'FERREDOXIN_NADP_REDUCTASE',   {'FERREDOXINNADP','FERRIDOXINNADP'}; ...
        'ATP_SYNTHASE',                {'ATPSYNTH','COMPLEXV','PHOTOPHOS'}; ...
        'RUBISCO',                     {'RIBULOSEBISPHOSPHATECARBOXYLASE'}; ...
        'PHOSPHORIBULOKINASE',         {'PHOSPHORIBULOKINASE'}; ...
        'RIBULOSE_EPIMERASE',          {'RIBULOSEPHOSPHATE3EPIMERASE'}; ...
        'RIBOSE5P_ISOMERASE',          {'RIBOSE5PHOSPHATEISOMERASE'}; ...
        'FRUCTOSE_BISPHOSPHATASE',     {'FRUCTOSEBISPHOSPHATASE'}; ...
        'SEDOHEPTULOSE_BISPHOSPHATASE',{'SEDOHEPTULOSEBISPHOSPHATASE'}; ...
        'TRANSKETOLASE',               {'TRANSKETOLASE'}; ...
        'TRANSALDOLASE',               {'TRANSALDOLASE'}};

    rows={};

    if ~runFull
        rows(end+1,:)={'SKIPPED','','',false,NaN,NaN,NaN,NaN,false, ...
            'Single carbon-boundary rescue found; pathway scan not required.'}; %#ok<AGROW>
    else
        ids=string(modelAUTO.rxns);
        if isfield(modelAUTO,'rxnNames')
            names=string(modelAUTO.rxnNames);
        else
            names=repmat("",numel(ids),1);
        end

        normText=strings(numel(ids),1);
        for j=1:numel(ids)
            normText(j)=string(norm_pub(ids(j)))+" "+string(norm_pub(names(j)));
        end

        for c=1:size(categories,1)
            cat=categories{c,1};
            toks=categories{c,2};
            hit=false(numel(ids),1);

            for t=1:numel(toks)
                hit = hit | contains(normText,string(toks{t}));
            end

            if strcmp(cat,'LIGHT_INPUT')
                hit = hit | ids==string(cfg.photonRxn);
            elseif strcmp(cat,'RUBISCO')
                hit = hit | ids==string(cfg.rubiscoRxn);
            end

            idx=find(hit);

            if isempty(idx)
                rows(end+1,:)={cat,'','',false,NaN,NaN,NaN,NaN,true, ...
                    'No matching reaction ID/name found.'}; %#ok<AGROW>
                continue;
            end

            for j=idx(:)'
                [vmin,vmax]=reactionFluxRange_pub(modelAUTO,modelAUTO.rxns{j});
                blocked = (~isfinite(vmin) || abs(vmin)<=cfg.autoDiagnosticFluxTol) && ...
                          (~isfinite(vmax) || abs(vmax)<=cfg.autoDiagnosticFluxTol);

                rows(end+1,:)={cat,char(ids(j)),char(names(j)),true, ...
                    modelAUTO.lb(j),modelAUTO.ub(j),vmin,vmax,blocked,''}; %#ok<AGROW>
            end
        end
    end

    T=cell2table(rows,'VariableNames',{'Pathway_category','Reaction_ID', ...
        'Reaction_name','Reaction_found','LB','UB','Min_feasible_flux', ...
        'Max_feasible_flux','Blocked_under_strict_AUTO','Note'});
end

function [vmin,vmax] = reactionFluxRange_pub(m,rxn)
    vmin=NaN; vmax=NaN;
    idx=find(strcmp(m.rxns,rxn),1);
    if isempty(idx), return; end

    mm=m; mm.c(:)=0; mm.c(idx)=1;
    smin=optimizeCbModel(mm,'min');
    if isfield(smin,'stat') && smin.stat==1 && ~isempty(smin.f)
        vmin=smin.f;
    end

    mm=m; mm.c(:)=0; mm.c(idx)=1;
    smax=optimizeCbModel(mm,'max');
    if isfield(smax,'stat') && smax.stat==1 && ~isempty(smax.f)
        vmax=smax.f;
    end
end

function T = mixoCO2FractionScan_pub( ...
    modelMIXO,substrateName,resolvedExchange,carbonAtoms, ...
    maxSubstrateUptake,mixoRuns,cfg)

    fractions=cfg.mixoCO2ScanFractions(:);
    nValid=sum(strlength(resolvedExchange)>0 & [mixoRuns.PredictedGrowth]');
    nRows=nValid*numel(fractions);

    Substrate=strings(nRows,1);
    Exchange=strings(nRows,1);
    Max_mu_h=nan(nRows,1);
    Biomass_fraction=nan(nRows,1);
    Target_mu_h=nan(nRows,1);
    Min_CO2_uptake=nan(nRows,1);
    Max_CO2_uptake=nan(nRows,1);
    Min_substrate_uptake=nan(nRows,1);
    Max_substrate_uptake=nan(nRows,1);
    CO2_uptake_feasible=false(nRows,1);
    Interpretation=strings(nRows,1);

    row=0;
    for i=1:numel(substrateName)
        if strlength(resolvedExchange(i))==0 || ~mixoRuns(i).PredictedGrowth
            continue;
        end

        r=char(resolvedExchange(i));
        m=modelMIXO;
        m=openUptake_pub(m,r,maxSubstrateUptake(i),false);
        assertCarbonBoundaryState_pub(m,{cfg.co2Rxn,r},cfg,'MIXO_CO2_SCAN');

        for f=1:numel(fractions)
            row=row+1;
            target=fractions(f)*mixoRuns(i).Mu_h;

            [co2Min,co2Max]=substrateUseRangeAtBiomass_pub( ...
                m,cfg.MIXO.sinkRxn,cfg.co2Rxn,target);
            [subMin,subMax]=substrateUseRangeAtBiomass_pub( ...
                m,cfg.MIXO.sinkRxn,r,target);

            Substrate(row)=substrateName(i);
            Exchange(row)=string(r);
            Max_mu_h(row)=mixoRuns(i).Mu_h;
            Biomass_fraction(row)=fractions(f);
            Target_mu_h(row)=target;
            Min_CO2_uptake(row)=co2Min;
            Max_CO2_uptake(row)=co2Max;
            Min_substrate_uptake(row)=subMin;
            Max_substrate_uptake(row)=subMax;
            CO2_uptake_feasible(row)=isfinite(co2Max) && co2Max>cfg.uptakeTol;

            if CO2_uptake_feasible(row)
                if isfinite(co2Min) && co2Min>cfg.uptakeTol
                    Interpretation(row)="NET_CO2_UPTAKE_REQUIRED";
                else
                    Interpretation(row)="NET_CO2_UPTAKE_OPTIONAL";
                end
            else
                Interpretation(row)="NET_CO2_UPTAKE_NOT_FEASIBLE_AT_TARGET_MU";
            end
        end
    end

    if row<nRows
        Substrate=Substrate(1:row);
        Exchange=Exchange(1:row);
        Max_mu_h=Max_mu_h(1:row);
        Biomass_fraction=Biomass_fraction(1:row);
        Target_mu_h=Target_mu_h(1:row);
        Min_CO2_uptake=Min_CO2_uptake(1:row);
        Max_CO2_uptake=Max_CO2_uptake(1:row);
        Min_substrate_uptake=Min_substrate_uptake(1:row);
        Max_substrate_uptake=Max_substrate_uptake(1:row);
        CO2_uptake_feasible=CO2_uptake_feasible(1:row);
        Interpretation=Interpretation(1:row);
    end

    T=table(Substrate,Exchange,Max_mu_h,Biomass_fraction,Target_mu_h, ...
        Min_CO2_uptake,Max_CO2_uptake,Min_substrate_uptake, ...
        Max_substrate_uptake,CO2_uptake_feasible,Interpretation);
end


function T = buildMixoPhotonCO2Test_pub( ...
    modelMIXO,substrateName,resolvedExchange,carbonAtoms,maxSubstrateUptake,cfg)

    wanted = string(cfg.mixoPhotonSweepSubstrates);
    photonCaps = cfg.mixoPhotonSweep(:);
    forcedCO2 = cfg.mixoForcedCO2(:);

    rows = {};

    for w = 1:numel(wanted)
        i = find(strcmpi(strtrim(substrateName),wanted(w)),1);

        if isempty(i) || strlength(resolvedExchange(i))==0
            rows(end+1,:) = {char(wanted(w)),'',NaN,NaN,NaN,false,0,NaN,NaN,NaN,NaN,NaN, ...
                'SUBSTRATE_NOT_RESOLVED'}; %#ok<AGROW>
            continue;
        end

        subRxn = char(resolvedExchange(i));
        qSub = maxSubstrateUptake(i);

        for p = 1:numel(photonCaps)
            for c = 1:numel(forcedCO2)
                m = modelMIXO;

                m = openUptake_pub(m,subRxn,qSub,false);
                assertCarbonBoundaryState_pub(m,{cfg.co2Rxn,subRxn},cfg,'MIXO_PHOTON_CO2_TEST');
                assertBiomassOutputState_pub(m,cfg.MIXO.sinkRxn,cfg,'MIXO_PHOTON_CO2_TEST');

                m = setUptakeOnly_pub(m,cfg.photonRxn,photonCaps(p));

                m = forceMinimumUptake_pub(m,cfg.co2Rxn,forcedCO2(c));

                m = setObjective_pub(m,cfg.MIXO.sinkRxn);
                sol = optimizeCbModel(m,'max');

                feasible = isfield(sol,'stat') && sol.stat==1 && ...
                    isfield(sol,'x') && ~isempty(sol.x);

                mu = 0;
                qCO2 = NaN;
                qOrg = NaN;
                photon = NaN;
                rubisco = NaN;
                o2Dummy = NaN;

                if feasible
                    mu = max(0,sol.f);
                    qCO2 = uptakeMagnitudeFromSolution_pub(m,cfg.co2Rxn,sol.x);
                    qOrg = uptakeMagnitudeFromSolution_pub(m,subRxn,sol.x);
                    photon = uptakeMagnitudeFromSolution_pub(m,cfg.photonRxn,sol.x);

                    ridx = find(strcmp(m.rxns,cfg.rubiscoRxn),1);
                    if ~isempty(ridx), rubisco = sol.x(ridx); end

                    didx = find(strcmp(m.rxns,cfg.o2DummyRxn),1);
                    if ~isempty(didx), o2Dummy = sol.x(didx); end
                end

                if ~feasible
                    interpretation = 'INFEASIBLE';
                elseif forcedCO2(c)>0 && qCO2>=forcedCO2(c)-1e-8
                    interpretation = 'FORCED_CO2_ASSIMILATION_FEASIBLE';
                elseif qCO2>cfg.uptakeTol
                    interpretation = 'SPONTANEOUS_CO2_ASSIMILATION';
                else
                    interpretation = 'NO_NET_CO2_ASSIMILATION';
                end

                rows(end+1,:) = { ...
                    char(substrateName(i)),subRxn,carbonAtoms(i),qSub, ...
                    photonCaps(p),forcedCO2(c),feasible,mu,qCO2,qOrg, ...
                    photon,rubisco,o2Dummy,interpretation}; %#ok<AGROW>
            end
        end
    end

    T = cell2table(rows,'VariableNames',{ ...
        'Substrate','Exchange','Carbon_atoms','Substrate_cap_mmol', ...
        'Photon_cap','Forced_min_CO2_uptake','Feasible','Mu_h', ...
        'Actual_CO2_uptake','Actual_substrate_uptake','Photon_uptake', ...
        'Rubisco_flux','O2_Dummy_flux','Interpretation'});
end

function m = forceMinimumUptake_pub(m,rxn,q)
    if q<=0
        return;
    end

    idx = find(strcmp(m.rxns,rxn),1);
    if isempty(idx)
        error('Cannot force uptake; reaction not found: %s',rxn);
    end

    sgn = uptakeFluxSign_pub(m,rxn);

    if sgn < 0
        m.ub(idx) = min(m.ub(idx),-abs(q));
    else
        m.lb(idx) = max(m.lb(idx),abs(q));
    end

    if m.lb(idx) > m.ub(idx)
        error('Forced uptake creates inconsistent bounds for %s.',rxn);
    end
end


function T = buildMixoLightCompare_pub( ...
    substrateName,resolvedExchange,carbonAtoms,mixo25,mixo50,cfg)

    n = numel(substrateName);
    Substrate = substrateName;
    Exchange = resolvedExchange;
    Carbon_atoms = carbonAtoms;

    Mu_25 = [mixo25.Mu_h]';
    Mu_50 = [mixo50.Mu_h]';
    Growth_25 = [mixo25.PredictedGrowth]';
    Growth_50 = [mixo50.PredictedGrowth]';
    Substrate_uptake_25 = [mixo25.SubstrateUptake_mmol]';
    Substrate_uptake_50 = [mixo50.SubstrateUptake_mmol]';
    CO2_uptake_25 = [mixo25.CO2Uptake_mmol]';
    CO2_uptake_50 = [mixo50.CO2Uptake_mmol]';
    Photon_uptake_25 = [mixo25.PhotonUptake]';
    Photon_uptake_50 = [mixo50.PhotonUptake]';
    O2Dummy_flux_25 = [mixo25.O2DummySinkFlux]';
    O2Dummy_flux_50 = [mixo50.O2DummySinkFlux]';

    CO2_external_C_fraction_25 = nan(n,1);
    CO2_external_C_fraction_50 = nan(n,1);
    Genuine_mixotrophy_25 = false(n,1);
    Genuine_mixotrophy_50 = false(n,1);
    Recommendation = strings(n,1);

    for i=1:n
        org25 = max(0,Substrate_uptake_25(i))*Carbon_atoms(i);
        org50 = max(0,Substrate_uptake_50(i))*Carbon_atoms(i);
        co225 = max(0,CO2_uptake_25(i));
        co250 = max(0,CO2_uptake_50(i));

        if org25+co225 > 0
            CO2_external_C_fraction_25(i) = co225/(org25+co225);
        end
        if org50+co250 > 0
            CO2_external_C_fraction_50(i) = co250/(org50+co250);
        end

        Genuine_mixotrophy_25(i) = Growth_25(i) && ...
            Substrate_uptake_25(i)>cfg.uptakeTol && CO2_uptake_25(i)>cfg.uptakeTol;
        Genuine_mixotrophy_50(i) = Growth_50(i) && ...
            Substrate_uptake_50(i)>cfg.uptakeTol && CO2_uptake_50(i)>cfg.uptakeTol;

        if Genuine_mixotrophy_25(i)
            Recommendation(i) = "25_SUFFICIENT";
        elseif Genuine_mixotrophy_50(i)
            Recommendation(i) = "50_REQUIRED";
        elseif Growth_25(i) || Growth_50(i)
            Recommendation(i) = "GROWTH_WITHOUT_NET_CO2";
        else
            Recommendation(i) = "NO_GROWTH";
        end
    end

    Delta_mu_50_minus_25 = Mu_50-Mu_25;
    Delta_CO2_50_minus_25 = CO2_uptake_50-CO2_uptake_25;

    T = table(Substrate,Exchange,Carbon_atoms, ...
        Mu_25,Growth_25,Substrate_uptake_25,CO2_uptake_25,Photon_uptake_25, ...
        O2Dummy_flux_25,CO2_external_C_fraction_25,Genuine_mixotrophy_25, ...
        Mu_50,Growth_50,Substrate_uptake_50,CO2_uptake_50,Photon_uptake_50, ...
        O2Dummy_flux_50,CO2_external_C_fraction_50,Genuine_mixotrophy_50, ...
        Delta_mu_50_minus_25,Delta_CO2_50_minus_25,Recommendation);
end

function nC = inferCarbonAtoms_pub(substrateName,substrateClass)
    name = lower(strtrim(string(substrateName)));
    cls = lower(strtrim(string(substrateClass)));

    if contains(cls,'disaccharide')
        nC = 12;
    elseif contains(cls,'hexose') || contains(cls,'deoxyhexose') || contains(cls,'hexitol')
        nC = 6;
    elseif contains(cls,'pentose') || contains(cls,'pentitol')
        nC = 5;
    elseif contains(cls,'tetritol')
        nC = 4;
    elseif contains(cls,'triol')
        nC = 3;
    elseif contains(cls,'organic acid')
        if contains(name,'acetate')
            nC = 2;
        elseif contains(name,'pyruvate')
            nC = 3;
        else
            error('Unknown organic-acid carbon count for substrate: %s',substrateName);
        end
    elseif contains(cls,'polymer') || contains(name,'starch')
        nC = 6;
    else
        error('Cannot infer carbon atoms for substrate %s (class %s).', ...
            substrateName,substrateClass);
    end
end

function y = parseGrowthPhenotype_pub(txt)
    t = lower(strtrim(string(txt)));
    if contains(t,'yes')
        y = 1;
    elseif contains(t,'no')
        y = 0;
    else
        y = NaN;
    end
end

function txt = agreementText_pub(expected,predicted)
    if isnan(expected)
        txt = "NOT_SCORED";
    elseif logical(expected) == logical(predicted)
        txt = "MATCH";
    else
        txt = "MISMATCH";
    end
end

function r = emptyRun_pub()
    r = struct( ...
        'Mode',"", ...
        'Substrate',"", ...
        'Status',"NOT_RUN", ...
        'BiomassRxn',"", ...
        'SubstrateRxn',"", ...
        'CarbonAtoms',NaN, ...
        'Mu_h',NaN, ...
        'PredictedGrowth',false, ...
        'SubstrateUptake_mmol',NaN, ...
        'SubstrateUptake_Cmmol',NaN, ...
        'MinRequiredUptakeAtFraction_mmol',NaN, ...
        'MaxPossibleUptakeAtFraction_mmol',NaN, ...
        'CanConsumeAtFraction',false, ...
        'CO2Uptake_mmol',NaN, ...
        'PhotonUptake',NaN, ...
        'O2DummySinkFlux',NaN);
end

function r = failedRun_pub(mode,substrate,status)
    r = emptyRun_pub();
    r.Mode = string(mode);
    r.Substrate = string(substrate);
    r.Status = string(status);
    r.Mu_h = 0;
    r.PredictedGrowth = false;
end

function T = runStructsToTable_pub(runs)
    T = struct2table(runs);
end
```
