## Supplementary Data 3 for "Experimental trophic flexibility guides a compartmentalized genome-scale model of red microalgae *Galdieria sulphuraria*": S3_Galdieria_publication_FVA.m

```
function output = S3_Galdieria_publication_FVA(modelInput, substrateFile, biomassFile)
% Publication analysis: 3_Galdieria_publication_FVA
% Galdieria sulphuraria publication Script 3: targeted flux variability analysis.
%
% PURPOSE
%   Reaction-level robustness extension of Script 2 at the identical
%   experimentally anchored operating point, mu = 0.060 h^-1.
%
% INPUTS
%   Galdieria_sulphuraria_SBML_Final.xml
%   Model_Substrate_list.xlsx
%   TOTAL_BIOMASS.xlsx
%
% DESIGN
%   HETERO only; experimental HETERO BOF; NGAM 1.7; GAM 30;
%   external CO2 uptake closed; Rubisco disabled; established HETERO
%   photon/pigment coupling retained.
%   Exact substrates: D-glucose, D-xylose, meso-erythritol, glycerol,
%   acetate and pyruvate.
%   Biomass fixed at mu = 0.060 h^-1. Six C-mmol gDW^-1 h^-1 is only
%   the maximum organic-carbon availability. Minimum carbon requirement is
%   calculated first; targeted FVA then uses <=105% of that minimum uptake.
%   Approximately 5-8 informative reactions are tested per substrate.
%   A reaction is obligatory only when its FVA interval excludes zero.
%
% OUTPUT
%   3_Galdieria_publication_FVA.xlsx
%   Sheets: 03_FVA_SUMMARY, 03_FVA_REACTIONS, METHODS
%
%% ========================= PATHS / INPUTS ===============================
scriptDir = fileparts(mfilename('fullpath'));
if isempty(scriptDir), scriptDir = pwd; end

if nargin < 1 || isempty(modelInput)
    modelInput = 'Galdieria_sulphuraria_SBML_Final.xml';
end
if nargin < 2 || isempty(substrateFile)
    substrateFile = 'Model_Substrate_list.xlsx';
end
if nargin < 3 || isempty(biomassFile)
    biomassFile = 'TOTAL_BIOMASS.xlsx';
end

% Inputs may be stored beside this script OR in a shared Inputs folder
% several levels above it (the normalized Supplementary Material layout).
modelInput = resolvePublicationInput_s3(modelInput,scriptDir);
substrateFile = resolvePublicationInput_s3(substrateFile,scriptDir);
biomassFile = resolvePublicationInput_s3(biomassFile,scriptDir);

if ~isfile(modelInput)
    error(['Final SBML model not found: %s\n' ...
           'Expected beside the script or in an ancestor Inputs folder.'], ...
           char(string(modelInput)));
end
if ~isfile(substrateFile)
    error(['Substrate workbook not found: %s\n' ...
           'Expected beside the script or in an ancestor Inputs folder.'], ...
           char(string(substrateFile)));
end
if ~isfile(biomassFile)
    error(['Biomass workbook not found: %s\n' ...
           'Expected beside the script or in an ancestor Inputs folder.'], ...
           char(string(biomassFile)));
end

ensureCobraReady_s3();

fprintf('\n============================================================\n');
fprintf('Galdieria publication Step 3: targeted FVA from publication\n');
fprintf('============================================================\n');
fprintf('Model:      %s\n',modelInput);
fprintf('Substrates: %s\n',substrateFile);
fprintf('Biomass:    %s\n',biomassFile);

baseModel = loadModel_s3(modelInput);
validateBasicModel_s3(baseModel);

%% ========================= CONFIGURATION ================================
cfg.scriptDir = scriptDir;
cfg.modelFile = string(modelInput);
cfg.substrateFile = string(substrateFile);
cfg.biomassFile = string(biomassFile);
cfg.substrateSheet = 'Sole_substrate_ScrumPy';
cfg.resultsFile = fullfile(scriptDir,'3_Galdieria_publication_FVA.xlsx');

% publication workflow continuity.
cfg.referenceMu_d = 1.44;
cfg.referenceMu_h = cfg.referenceMu_d/24;     % 0.060 h^-1
cfg.referenceMuCondition = "KIN2_HET_MIX";
cfg.referenceMuMethod = "Log-linear fit, days 1-5";

cfg.NGAM = 1.7;
cfg.GAM = 30;
cfg.screenCarbon = 6;                         % C-mmol gDW^-1 h^-1
cfg.validationMaxCarbon = 120;                % non-binding min-demand opening
cfg.carbonFVATolerance = 0.05;                % <=105% min total carbon

cfg.o2MaxUptake = 1000;
cfg.autoPhotonProbeMax = 20;
cfg.heteroPhotonMaxUptake = 0.2;
cfg.mixoPhotonMaxUptake = 50;
cfg.autoCO2CarbonCap = cfg.screenCarbon;
cfg.mixoCO2MaxUptake = cfg.validationMaxCarbon;
cfg.rubiscoMinFluxAUTO = 0.01;
cfg.rubiscoMinFluxMIXO = 0.01;

cfg.expectedPhotonPerPigment = 0.58;
cfg.photonCalibrationTolerance = 0.02;

cfg.growthTol = 1e-8;
cfg.uptakeTol = 1e-9;
cfg.fluxTol = 1e-9;
cfg.strictCarbonBoundaryClosure = true;

cfg.targetFvaMode = "HETERO";
cfg.targetMu_h = 0.060;
cfg.targetUptakeTolerance = 0.05;              % <=105% minimum uptake
cfg.minimumTargetSubstrates = 6;
cfg.maxTargetSubstrates = 6;
cfg.maxTargetReactionsPerSubstrate = 8;
cfg.minTargetReactionsPerSubstrate = 5;
cfg.targetTopActiveReactions = 12;
cfg.targetLocalGraphDepth = 2;
cfg.preferredTargetSubstrates = [ ...
    "D-Glucose"
    "D-Xylose"
    "meso-Erythritol"
    "Glycerol"
    "Acetate"
    "Pyruvate"
];

% Explicit basal medium, retained from publication workflow.
cfg.basalMediumLB = -1000;
cfg.basalMediumUB = 1000;
cfg.o2DummyLB = 0;
cfg.o2DummyUB = 1000;

% Exact experimental BOF inputs used in the publication validation.
cfg.exp.MIXO.protein       = 0.4147;
cfg.exp.MIXO.carbohydrate  = 0.3377;
cfg.exp.MIXO.nucleic       = 0.0775;
cfg.exp.MIXO.lipid         = 0.0701;

cfg.exp.HETERO.protein      = 0.4016;
cfg.exp.HETERO.carbohydrate = 0.3609;
cfg.exp.HETERO.nucleic      = 0.0765;
cfg.exp.HETERO.lipid        = 0.0610;

cfg.reference.MIXO.rnaFraction = 0.06222707/(0.06222707+0.01091703);
cfg.reference.HETERO.rnaFraction = 0.085/(0.085+0.015);

cfg.reference.HETERO.pigmentMass = 0.00485;
cfg.reference.MIXO.PCB_mg  = 1.714658;
cfg.reference.MIXO.CHLA_mg = 4.450000;
cfg.reference.MIXO.BETA_mg = 0.240000;
cfg.reference.MIXO.ZEA_mg  = 0.350000;
cfg.reference.MIXO.pigmentMass = ...
    (cfg.reference.MIXO.PCB_mg + cfg.reference.MIXO.CHLA_mg + ...
     cfg.reference.MIXO.BETA_mg + cfg.reference.MIXO.ZEA_mg)/1000;

%% ========================= SUBSTRATE WORKBOOK ===========================
T = readtable(substrateFile,'Sheet',cfg.substrateSheet, ...
    'VariableNamingRule','preserve');

requiredCols = {'Substrate_ID','Class','Growth (1995)', ...
                'MetaCyc / model exchange ID'};
for k = 1:numel(requiredCols)
    if ~ismember(requiredCols{k},T.Properties.VariableNames)
        error('Required column missing from %s: %s',cfg.substrateSheet,requiredCols{k});
    end
end

substrateName = string(T.('Substrate_ID'));
substrateClass = string(T.('Class'));
expGrowthText = string(T.('Growth (1995)'));
requestedExchange = string(T.('MetaCyc / model exchange ID'));

if ismember('Modeling notes',T.Properties.VariableNames)
    modelingNotes = string(T.('Modeling notes'));
else
    modelingNotes = repmat("",height(T),1);
end

doiText = optionalColumn_s3(T,["DOI","doi"]);
referenceText = optionalColumn_s3(T, ...
    ["Reference","Literature reference","Literature","Source"]);

resolvedExchange = strings(height(T),1);
carbonAtoms = nan(height(T),1);
carbonAtomSource = strings(height(T),1);
expectedGrowth = nan(height(T),1);

for i = 1:height(T)
    resolvedExchange(i) = string(resolveRxn_s3(baseModel,requestedExchange(i),true));
    if strlength(resolvedExchange(i)) == 0
        [carbonAtoms(i),carbonAtomSource(i)] = ...
            inferCarbonAtoms_s3(baseModel,"",substrateName(i),substrateClass(i));
    else
        [carbonAtoms(i),carbonAtomSource(i)] = ...
            inferCarbonAtoms_s3(baseModel,resolvedExchange(i),substrateName(i),substrateClass(i));
    end
    expectedGrowth(i) = parseGrowthPhenotype_s3(expGrowthText(i));
end

nMissingExchange=sum(strlength(resolvedExchange)==0);
if nMissingExchange>0
    fprintf('Substrate mapping: %d/%d exchanges resolved; %d unresolved.\n', ...
        height(T)-nMissingExchange,height(T),nMissingExchange);
end

if any(~isfinite(carbonAtoms) | carbonAtoms<=0)
    bad = substrateName(~isfinite(carbonAtoms) | carbonAtoms<=0);
    error('Carbon count could not be determined for: %s',strjoin(cellstr(bad),', '));
end

duplicateExchange = false(height(T),1);
for i = 1:height(T)
    if strlength(resolvedExchange(i))>0
        duplicateExchange(i) = sum(resolvedExchange==resolvedExchange(i))>1;
    end
end

cfg.allMappedCarbonRxns = cellstr(unique( ...
    resolvedExchange(strlength(resolvedExchange)>0),'stable'));

%% ========================= EXACT MODEL MAPPING ==========================
cfg.ngamRxn = requireRxn_s3(baseModel, ...
    {'R_NGAM_ATP_Cyto','NGAM_ATP_Cyto','ATPM_NGAM_Cyto'});
cfg.photonRxn = requireRxn_s3(baseModel, ...
    {'R_Photon_Cyto_tx','Photon_Cyto_tx'});
cfg.co2Rxn = requireRxn_s3(baseModel, ...
    {'R_CO2_Cyto_tx','CO2_Cyto_tx'});
cfg.o2Rxn = requireRxn_s3(baseModel, ...
    {'R_O2_Cyto_tx','O2_Cyto_tx'});
cfg.o2DummyRxn = requireRxn_s3(baseModel, ...
    {'R_O2_Dummy_Cyto_tx','O2_Dummy_Cyto_tx'});
cfg.rubiscoRxn = requireRxn_s3(baseModel,{ ...
    'R_RIBULOSE-BISPHOSPHATE-CARBOXYLASE-RXN_Plas', ...
    'RIBULOSE-BISPHOSPHATE-CARBOXYLASE-RXN_Plas'});

cfg.basalRxns.Fe2 = requireRxn_s3(baseModel,{'R_FE+2_Cyto_tx','FE+2_Cyto_tx'});
cfg.basalRxns.Mg2 = requireRxn_s3(baseModel,{'R_MG+2_Cyto_tx','MG+2_Cyto_tx'});
cfg.basalRxns.NH4 = requireRxn_s3(baseModel,{'R_NH4_Cyto_tx','NH4_Cyto_tx'});
cfg.basalRxns.NO3 = requireRxn_s3(baseModel,{'R_NO3_Cyto_tx','NO3_Cyto_tx'});
cfg.basalRxns.H   = requireRxn_s3(baseModel,{'R_PROTON_Cyto_tx','PROTON_Cyto_tx'});
cfg.basalRxns.Pi  = requireRxn_s3(baseModel,{'R_Pi_Cyto_tx','Pi_Cyto_tx'});
cfg.basalRxns.SO4 = requireRxn_s3(baseModel,{'R_SO4_Cyto_tx','SO4_Cyto_tx'});
cfg.basalRxns.H2O = requireRxn_s3(baseModel,{'R_WATER_Cyto_tx','WATER_Cyto_tx'});

[cfg.carbonBoundaryRxns,carbonBoundaryAudit] = ...
    identifyCarbonBoundaryRxns_s3(baseModel,cfg);
cfg.carbonBoundaryRxns = cellstr(unique([ ...
    string(cfg.carbonBoundaryRxns(:)); ...
    string(cfg.allMappedCarbonRxns(:)); ...
    string(cfg.co2Rxn)],'stable'));

%% ========================= CLOSE OLD BIOMASS OUTPUTS ====================
[baseModel,oldBiomassAudit] = closeOriginalBiomassOutputs_s3(baseModel);

%% ========================= BUILD TROPHIC BOFs ===========================
% AUTO: dynamically rebuilt from TOTAL_BIOMASS_AUTO_MEAN.
[baseModel,cfg,autoBofAudit] = implementAutoWorkbookBOF_s3( ...
    baseModel,cfg,biomassFile);

% MIXO/HETERO: exact experimental BOF composition from cultivation
% validation, using the real GSM pool metabolites.
[baseModel,cfg,experimentalBofAudit] = ...
    implementExperimentalBOFs_s3(baseModel,cfg);

cfg.allAnalysisBOFRxns = { ...
    cfg.AUTO.assemblyRxn,cfg.AUTO.sinkRxn,cfg.AUTO.pigmentRxn, ...
    cfg.MIXO.assemblyRxn,cfg.MIXO.sinkRxn,cfg.MIXO.pigmentRxn, ...
    cfg.HETERO.assemblyRxn,cfg.HETERO.sinkRxn,cfg.HETERO.pigmentRxn};

%% ========================= PREPARE TROPHIC MODELS ======================
modelAUTO = prepareCondition_s3(baseModel,'AUTO',cfg);
modelMIXO = prepareCondition_s3(baseModel,'MIXO',cfg);
modelHETERO_uncoupled = prepareCondition_s3(baseModel,'HETERO',cfg);

% Calibrate the small HETERO pigment-linked photon requirement on glucose,
% exactly as a model-derived structural coupling, not as measured irradiance.
glcRow = find(strcmpi(strtrim(substrateName),'D-Glucose'),1);
if isempty(glcRow) || strlength(resolvedExchange(glcRow))==0
    error('D-Glucose must be present and mapped for HETERO photon calibration.');
end

[photonPerPigment,photonPerBiomass,photonCalibration] = ...
    calibrateHeteroPhoton_s3( ...
        modelHETERO_uncoupled,cfg.HETERO.sinkRxn,cfg.HETERO.pigmentRxn, ...
        char(resolvedExchange(glcRow)),carbonAtoms(glcRow),cfg);

cfg.HETERO.photonPerPigment = photonPerPigment;
cfg.HETERO.photonPerBiomass = photonPerBiomass;

if abs(photonPerPigment-cfg.expectedPhotonPerPigment) > ...
        cfg.photonCalibrationTolerance
    fprintf(2,['NOTICE: HETERO photon/pigment calibration = %.6g; publication workflow expectation ' ...
        'was %.6g +/- %.6g. publication calculated value is retained.\n'], ...
        photonPerPigment,cfg.expectedPhotonPerPigment,cfg.photonCalibrationTolerance);
end

modelHETERO = addPhotonBiomassCoupling_s3( ...
    modelHETERO_uncoupled,cfg.photonRxn,cfg.HETERO.sinkRxn, ...
    photonPerBiomass);

%% ========================= STEP 3A. HETERO QUALIFICATION ================
% A lightweight HETERO capacity pass is required only to confirm which
% mapped substrates are valid candidates for targeted FVA. These screening
% results are kept in memory and are NOT exported as a separate Step 2 sheet.
fprintf('\nStep 3A/3B: qualifying mapped HETERO substrates for targeted FVA...\n');

n = height(T);
heteroCapacity = repmat(emptyCapacityResult_s3(),n,1);

for i = 1:n
    if strlength(resolvedExchange(i))==0
        heteroCapacity(i) = failedCapacityResult_s3( ...
            'HETERO',substrateName(i),'MISSING_EXCHANGE');
        continue;
    end

    r = char(resolvedExchange(i));
    q = cfg.screenCarbon/carbonAtoms(i);

    h = modelHETERO;
    h = openUptake_s3(h,r,q,false);
    heteroCapacity(i) = capacityOne_s3( ...
        h,cfg.HETERO.sinkRxn,'HETERO', ...
        substrateName(i),r,carbonAtoms(i),cfg);
end

%% ========================= STEP 3B. TARGETED FVA ========================
fprintf('Step 3B/3B: selecting >=%d substrates and running fixed-mu FVA...\n', ...
    cfg.minimumTargetSubstrates);

targetRows = chooseTargetSubstrates_s3( ...
    substrateName,substrateClass,resolvedExchange,heteroCapacity,cfg);

[targetFvaSummary,targetFvaReactions] = runTargetedFVA_s3( ...
    modelHETERO,cfg.HETERO.sinkRxn, ...
    substrateName,substrateClass,resolvedExchange,carbonAtoms, ...
    expGrowthText,doiText,referenceText,targetRows,cfg);

% Publication-oriented summary order: put the literature information next
% to the phenotype and keep technical counting fields toward the right.
targetFvaSummary = reorderFvaSummary_s3(targetFvaSummary);

% Methods/settings are consolidated into ONE sheet rather than multiple
% audit/reference worksheets.
methodsTable = buildStep3MethodsTable_s3(cfg);

%% ========================= WRITE THREE-SHEET FINAL EXCEL ==============
if isfile(cfg.resultsFile)
    delete(cfg.resultsFile);
end

writetable(targetFvaSummary,cfg.resultsFile,'Sheet','03_FVA_SUMMARY');
writetable(targetFvaReactions,cfg.resultsFile,'Sheet','03_FVA_REACTIONS');
writetable(methodsTable,cfg.resultsFile,'Sheet','METHODS');

fprintf('\nDONE -- publication Step 3 targeted FVA.\n');
fprintf('Final Excel workbook: %s\n',cfg.resultsFile);
fprintf('Sheets: 03_FVA_SUMMARY | 03_FVA_REACTIONS | METHODS (simplified)\n');
fprintf('Selected substrates (%d): %s\n',numel(targetRows), ...
    strjoin(cellstr(substrateName(targetRows)),', '));

output = struct();
output.resultsFile = cfg.resultsFile;
output.targetedFVASummary = targetFvaSummary;
output.targetedFVAReactions = targetFvaReactions;
output.methods = methodsTable;
output.selectedSubstrates = substrateName(targetRows);
output.cfg = cfg;

end


%% ======================================================================
%% INPUT / COBRA
%% ======================================================================

function f = resolvePublicationInput_s3(f,scriptDir)
% Resolve a publication input without hard-coded absolute paths.
%
% Search order:
%   1. exact path supplied by the user
%   2. script folder
%   3. scriptFolder\Inputs
%   4. each ancestor folder
%   5. each ancestorFolder\Inputs
%
% This supports normalized archives where the analysis script can be nested
% several levels below Supplementary Material 8\Inputs.

requested = char(string(f));

% Exact path first.
if isfile(requested)
    f=requested;
    return;
end

% Work with the basename when the supplied/default path is not found.
[~,name,ext]=fileparts(requested);
if isempty(name)
    f=requested;
    return;
end
baseName=[name ext];

current=scriptDir;
maxLevels=20;

for level=0:maxLevels
    candidates={ ...
        fullfile(current,baseName), ...
        fullfile(current,'Inputs',baseName) ...
        };

    for k=1:numel(candidates)
        if isfile(candidates{k})
            f=candidates{k};
            return;
        end
    end

    parent=fileparts(current);
    if isempty(parent) || strcmp(parent,current)
        break;
    end
    current=parent;
end

% Not found: return the requested name so the caller can give a clear error.
f=requested;
end

function ensureCobraReady_s3()
required = {'readCbModel','optimizeCbModel','addReaction','addMetabolite'};
missing = {};
for i=1:numel(required)
    if exist(required{i},'file')~=2
        missing{end+1}=required{i}; %#ok<AGROW>
    end
end
if ~isempty(missing)
    error(['COBRA Toolbox functions are not on the MATLAB path: %s. ' ...
           'Start/configure COBRA Toolbox before running this script.'], ...
           strjoin(missing,', '));
end

% IMPORTANT: do not call initCobraToolbox or changeCobraSolver here.
% On some COBRA installations changeCobraSolver triggers toolbox setup,
% git/submodule checks and solver probing. This analysis must not modify or
% reinitialize the user's COBRA installation. optimizeCbModel will use the
% LP solver already configured in the current MATLAB/COBRA session.
solverName="configured COBRA LP solver";
try
    global CBT_LP_SOLVER
    if ~isempty(CBT_LP_SOLVER)
        solverName=string(CBT_LP_SOLVER);
    end
catch
end
fprintf('COBRA ready; using %s. No toolbox initialization will be run.\n',solverName);
end

function model = loadModel_s3(modelFile)
% readCbModel can print harmless annotation/subSystem warnings for this SBML.
% Capture those messages locally so the publication analysis log remains
% readable. Warning state is always restored, including after an error.
warnState=warning;
cleanupObj=onCleanup(@() warning(warnState)); %#ok<NASGU>
warning('off','all');

try
    evalc('model = readCbModel(modelFile);');
catch ME1
    try
        evalc('model = readCbModel(modelFile,''fileType'',''SBML'');');
    catch ME2
        error(['Unable to load SBML with readCbModel.\nFirst error: %s\n' ...
               'Second error: %s\nEnsure libSBML/TranslateSBML required by ' ...
               'your COBRA installation is available.'],ME1.message,ME2.message);
    end
end

fprintf('Model loaded quietly: %d reactions, %d metabolites.\n', ...
    numel(model.rxns),numel(model.mets));
end

function validateBasicModel_s3(model)
required={'rxns','mets','S','lb','ub'};
for i=1:numel(required)
    if ~isfield(model,required{i})
        error('Loaded model lacks required COBRA field: %s',required{i});
    end
end
if size(model.S,1)~=numel(model.mets) || size(model.S,2)~=numel(model.rxns)
    error('Model S dimensions do not match mets/rxns.');
end
if numel(model.lb)~=numel(model.rxns) || numel(model.ub)~=numel(model.rxns)
    error('Model bound vectors do not match rxns.');
end
end


%% ======================================================================
%% ID / WORKBOOK HELPERS
%% ======================================================================

function r = requireRxn_s3(model,candidates)
r = resolveRxn_s3(model,string(candidates),true);
if strlength(string(r))==0
    error('Required reaction not found. Tried: %s',strjoin(candidates,', '));
end
r = char(r);
end

function r = resolveRxn_s3(model,query,allowEmpty)
if nargin<3, allowEmpty=false; end
r = "";
qList = string(query(:));
rxns = string(model.rxns(:));

for q = qList'
    q = strtrim(q);
    if strlength(q)==0, continue; end

    idx=find(strcmp(rxns,q),1);
    if ~isempty(idx), r=rxns(idx); return; end

    if startsWith(q,"R_")
        alt=extractAfter(q,2);
    else
        alt="R_"+q;
    end
    idx=find(strcmp(rxns,alt),1);
    if ~isempty(idx), r=rxns(idx); return; end

    nq=normalizeId_s3(q);
    nr=strings(size(rxns));
    for kk=1:numel(rxns)
        nr(kk)=normalizeId_s3(rxns(kk));
    end
    idx=find(nr==nq,1);
    if ~isempty(idx), r=rxns(idx); return; end
end

if ~allowEmpty
    error('Reaction could not be resolved: %s',strjoin(cellstr(qList),', '));
end
end

function s = normalizeId_s3(s)
s=upper(strtrim(string(s)));
s=replace(s,["__45__","__43__","__91__","__93__"],["-","+","[","]"]);
s=replace(s,[" ","[","]"],"");
s=regexprep(s,'^R_','');
s=regexprep(s,'[^A-Z0-9+\-]','_');
s=regexprep(s,'_+','_');
end

function met = requireMetCandidates_s3(model,candidates)
mets=string(model.mets(:));
for q=string(candidates(:))'
    idx=find(strcmpi(mets,q),1);
    if ~isempty(idx), met=char(mets(idx)); return; end
end

normM=strings(size(mets));
for kk=1:numel(mets)
    normM(kk)=normalizeMetFull_s3(mets(kk));
end
for q=string(candidates(:))'
    nq=normalizeMetFull_s3(q);
    idx=find(normM==nq);
    if numel(idx)==1, met=char(mets(idx)); return; end
end

% Final safe fallback: compare compartment-free biochemical core only, but
% only accept a unique model metabolite.
coreM=strings(size(mets));
for kk=1:numel(mets)
    coreM(kk)=metCoreNoComp_s3(mets(kk));
end
for q=string(candidates(:))'
    nq=metCoreNoComp_s3(q);
    idx=find(coreM==nq);
    if numel(idx)==1, met=char(mets(idx)); return; end
end

error('Required metabolite not uniquely resolved. Tried: %s', ...
    strjoin(cellstr(string(candidates(:))),', '));
end

function s=normalizeMetFull_s3(s)
s=upper(strtrim(string(s)));
s=replace(s,["__45__","__43__","__91__","__93__"],["-","+","[","]"]);
s=regexprep(s,'^M_','');
s=replace(s," ","");
s=regexprep(s,'[^A-Z0-9+\-\[\]]','_');
s=regexprep(s,'_+','_');
end

function s=metCoreNoComp_s3(s)
s=normalizeMetFull_s3(s);
s=regexprep(s,'\[[A-Z]\]$','');
s=regexprep(s,'_(CYTO|MITO|PLAS|THYL|PERO)$','');
end

function out = optionalColumn_s3(T,candidates)
out=repmat("",height(T),1);
vars=string(T.Properties.VariableNames);
for q=string(candidates(:))'
    idx=find(strcmpi(vars,q),1);
    if ~isempty(idx)
        out=string(T.(T.Properties.VariableNames{idx}));
        return;
    end
end
end

function formula = readWorkbookReactionFormula_s3(file,sheet,abbr)
C=readcell(file,'Sheet',sheet);
formula="";
for i=1:size(C,1)
    for j=1:size(C,2)
        v=stringCell_s3(C{i,j});
        if strcmpi(strtrim(v),string(abbr))
            % Search the same row for a reaction-like expression.
            for k=1:size(C,2)
                s=stringCell_s3(C{i,k});
                if contains(s,"->") || contains(s,"<=>") || contains(s,"<->")
                    formula=s;
                    break;
                end
            end
            if strlength(formula)>0, return; end
        end
    end
end
error('Reaction formula not found in %s / %s for %s',file,sheet,abbr);
end

function s=stringCell_s3(v)
if isempty(v)
    s="";
elseif isstring(v)
    if all(ismissing(v(:))), s=""; else, s=string(v); end
elseif ischar(v)
    s=string(v);
elseif isnumeric(v) && isscalar(v)
    if isnan(v), s=""; else, s=string(v); end
elseif islogical(v) && isscalar(v)
    s=string(v);
else
    try
        tmp=string(v);
        if all(ismissing(tmp(:))), s=""; else, s=tmp; end
    catch
        s="";
    end
end
end

function c=coefficientForWorkbookMet_s3(formula,target)
% Extract an exact stoichiometric coefficient from a human-readable
% reaction formula. Excel may wrap the formula across lines; normalize all
% whitespace first. Delimiters are reaction '+' signs and the '->' arrow.
f=char(string(formula));
f=regexprep(f,'[\r\n\t]+',' ');
f=regexprep(f,'\s+',' ');
f=strtrim(f);
t=regexptranslate('escape',char(string(target)));

% Exact reaction term, e.g. "0.59 m_PROTEIN_TOTAL_POOL[c]".
% The separator itself is included in the match, which avoids fragile
% look-behind expressions and works with bracketed compartment IDs.
num='([0-9]*\.?[0-9]+(?:[eE][+\-]?[0-9]+)?)?';
pat=['(?:^|\+|->)\s*' num '\s*' t '\s*(?:\+|->|$)'];
tok=regexp(f,pat,'tokens','once');

if isempty(tok)
    error(['Metabolite %s was not found as an exact reaction term in ' ...
           'the workbook formula:\n%s'],target,f);
end

if isempty(tok{1})
    c=1;
else
    c=str2double(tok{1});
end
if ~isfinite(c)
    error('Invalid coefficient for %s in workbook formula.',target);
end
end


%% ======================================================================
%% CARBON COUNT / BOUNDARY
%% ======================================================================

function [nC,source] = inferCarbonAtoms_s3(model,exchange,substrateName,substrateClass)
nC=NaN; source="";

if strlength(string(exchange))>0
    ridx=find(strcmp(model.rxns,char(exchange)),1);
    if ~isempty(ridx)
        nz=find(abs(full(model.S(:,ridx)))>1e-12);
        if numel(nz)==1 && isfield(model,'metFormulas') && ...
                numel(model.metFormulas)>=nz
            formula=string(model.metFormulas{nz});
            [nC0,ok]=carbonCountFromFormula_s3(formula);
            if ok
                nC=nC0;
                source="Model metabolite formula: "+formula;
                return;
            end
        end
    end
end

name=lower(strtrim(string(substrateName)));
cls=lower(strtrim(string(substrateClass)));

if contains(cls,'disaccharide')
    nC=12;
elseif contains(cls,'hexose') || contains(cls,'deoxyhexose') || contains(cls,'hexitol')
    nC=6;
elseif contains(cls,'pentose') || contains(cls,'pentitol')
    nC=5;
elseif contains(cls,'tetritol')
    nC=4;
elseif contains(cls,'triol')
    nC=3;
elseif contains(cls,'organic acid')
    if contains(name,'acetate')
        nC=2;
    elseif contains(name,'pyruvate')
        nC=3;
    end
elseif contains(cls,'polymer') || contains(name,'starch')
    nC=6; % glucose-equivalent unit used in publication workflow
end

if isfinite(nC)
    source="publication workflow substrate-class fallback";
else
    source="UNRESOLVED";
end
end

function [nC,ok]=carbonCountFromFormula_s3(formula)
nC=NaN; ok=false;
formula=char(strtrim(string(formula)));
if isempty(formula), return; end
tok=regexp(formula,'(?:^|[^A-Za-z])C([0-9]*)','tokens','once');
if isempty(tok)
    % Standard formulas normally start with C; also handle directly.
    tok=regexp(formula,'^C([0-9]*)','tokens','once');
end
if isempty(tok), return; end
if isempty(tok{1})
    nC=1;
else
    nC=str2double(tok{1});
end
ok=isfinite(nC) && nC>0;
end

function [rxns,auditT]=identifyCarbonBoundaryRxns_s3(model,cfg)
nrxn=numel(model.rxns);
isBoundary=false(nrxn,1);
hasCarbon=false(nrxn,1);
metID=strings(nrxn,1);
formula=strings(nrxn,1);
nC=nan(nrxn,1);

for j=1:nrxn
    nz=find(abs(full(model.S(:,j)))>1e-12);
    if numel(nz)==1
        isBoundary(j)=true;
        metID(j)=string(model.mets{nz});
        if isfield(model,'metFormulas') && numel(model.metFormulas)>=nz
            formula(j)=string(model.metFormulas{nz});
            [nC(j),hasCarbon(j)]=carbonCountFromFormula_s3(formula(j));
        end
    end
end

idx=find(isBoundary & hasCarbon);
rxns=cellstr(string(model.rxns(idx)));

% Always include known mapped substrates and CO2 later in the caller.
auditT=table(string(model.rxns(idx)),metID(idx),formula(idx),nC(idx), ...
    'VariableNames',{'Reaction_ID','Boundary_metabolite','Formula','Carbon_atoms'});

if isempty(idx)
    fprintf(2,['NOTICE: no additional carbon boundary reactions were identified ' ...
        'from metFormulas; mapped substrate exchanges will still be closed.\n']);
end
end


%% ======================================================================
%% BIOMASS FUNCTIONS
%% ======================================================================

function [m,auditT]=closeOriginalBiomassOutputs_s3(m)
ids=string(m.rxns(:));
if isfield(m,'rxnNames')
    names=string(m.rxnNames(:));
else
    names=repmat("",numel(ids),1);
end

rawID=lower(ids); rawName=lower(names);
normID=strings(size(ids)); normName=strings(size(names));
for kk=1:numel(ids)
    normID(kk)=replace(normalizeId_s3(ids(kk)),"_","");
    normName(kk)=replace(normalizeId_s3(names(kk)),"_","");
end

isBmTx=endsWith(rawID,"_bm_tx") | endsWith(rawName,"_bm_tx") | ...
       contains(normID,"BMTX") | contains(normName,"BMTX");
isTotalBiomass=contains(normID,"TOTALBIOMASS") | contains(normName,"TOTALBIOMASS");
mask=isBmTx | isTotalBiomass;

idx=find(mask);
oldLB=m.lb(idx); oldUB=m.ub(idx);
if isfield(m,'c') && numel(m.c)==numel(m.rxns)
    oldC=m.c(idx);
else
    oldC=zeros(numel(idx),1);
    m.c=zeros(numel(m.rxns),1);
end

reason=strings(numel(idx),1);
for kk=1:numel(idx)
    j=idx(kk);
    if isBmTx(j) && isTotalBiomass(j)
        reason(kk)="bm_tx + TOTAL_BIOMASS";
    elseif isBmTx(j)
        reason(kk)="pre-existing bm_tx";
    else
        reason(kk)="TOTAL_BIOMASS";
    end
end

m.lb(idx)=0; m.ub(idx)=0; m.c(:)=0;

auditT=table(ids(idx),names(idx),reason,oldLB,oldUB,oldC, ...
    zeros(numel(idx),1),zeros(numel(idx),1), ...
    'VariableNames',{'Reaction_ID','Reaction_name','Closure_reason', ...
    'Original_LB','Original_UB','Original_objective','New_LB','New_UB'});
end

function [m,cfg,auditT]=implementAutoWorkbookBOF_s3(m,cfg,biomassFile)
% Resolve common real GSM metabolites.
mets.protein=requireMetCandidates_s3(m,{'m_PROTEIN_TOTAL_POOL_AUTO_Cyto[c]','PROTEIN_TOTAL_POOL_AUTO_CYTO','m_PROTEIN_TOTAL_POOL_Cyto[c]','PROTEIN_TOTAL_POOL_CYTO'});
mets.carb=requireMetCandidates_s3(m,{'m_CARBOHYDRATE_TOTAL_POOL_AUTO_Cyto[c]','CARBOHYDRATE_TOTAL_POOL_AUTO_CYTO','m_CARBOHYDRATE_TOTAL_POOL_Cyto[c]','CARBOHYDRATE_TOTAL_POOL_CYTO'});
mets.lipid=requireMetCandidates_s3(m,{'m_LIPID_TOTAL_POOL_AUTO_Cyto[c]','LIPID_TOTAL_POOL_AUTO_CYTO','m_LIPID_TOTAL_POOL_Cyto[c]','LIPID_TOTAL_POOL_CYTO'});
mets.rna=requireMetCandidates_s3(m,{'m_RNA_TOTAL_POOL_AUTO_Cyto[c]','RNA_TOTAL_POOL_AUTO_CYTO','m_RNA_TOTAL_POOL_Cyto[c]','RNA_TOTAL_POOL_CYTO'});
mets.dna=requireMetCandidates_s3(m,{'m_DNA_TOTAL_POOL_AUTO_Cyto[c]','DNA_TOTAL_POOL_AUTO_CYTO','m_DNA_TOTAL_POOL_Cyto[c]','DNA_TOTAL_POOL_CYTO'});
mets.atp=requireMetCandidates_s3(m,{'m_ATP_Cyto[c]','ATP_CYTO'});
mets.water=requireMetCandidates_s3(m,{'m_WATER_Cyto[c]','WATER_CYTO'});
mets.adp=requireMetCandidates_s3(m,{'m_ADP_Cyto[c]','ADP_CYTO'});
mets.pi=requireMetCandidates_s3(m,{'m_Pi_Cyto[c]','PI_CYTO'});
mets.proton=requireMetCandidates_s3(m,{'m_PROTON_Cyto[c]','PROTON_CYTO'});

pig.zea=requireMetCandidates_s3(m,{'m_CPD1F-130_Thyl[t]','CPD1F-130_THYL'});
pig.chla=requireMetCandidates_s3(m,{'m_CHLOROPHYLL-A_Plas[p]','m_CHLOROPHYLL-A_Thyl[t]', ...
                                    'CHLOROPHYLL-A_PLAS','CHLOROPHYLL-A_THYL'});
pig.beta=requireMetCandidates_s3(m,{'m_CPD1F-129_Thyl[t]','CPD1F-129_THYL'});
pig.pcb=requireMetCandidates_s3(m,{'m_3Z-PHYCOCYANOBILIN_Plas[p]','3Z-PHYCOCYANOBILIN_PLAS'});

cfg.resolvedBOFMets=mets;
cfg.resolvedPigmentMets=pig;

bofFormula=readWorkbookReactionFormula_s3( ...
    biomassFile,'Reaction List Auto','TOTAL_BIOMASS_AUTO_MEAN');
pigFormula=readWorkbookReactionFormula_s3( ...
    biomassFile,'Reaction List Auto','PIGMENT_TOTAL_POOL_BIO_AUTOTROPH_NOFRAC');

cProtein=coefficientForWorkbookMet_s3(bofFormula,'m_PROTEIN_TOTAL_POOL[c]');
cCarb=coefficientForWorkbookMet_s3(bofFormula,'m_CARBOHYDRATE_TOTAL_POOL[c]');
cLipid=coefficientForWorkbookMet_s3(bofFormula,'m_LIPID_TOTAL_POOL[c]');
cRNA=coefficientForWorkbookMet_s3(bofFormula,'m_RNA_TOTAL_POOL[c]');
cDNA=coefficientForWorkbookMet_s3(bofFormula,'m_DNA_TOTAL_POOL[c]');
cPig=coefficientForWorkbookMet_s3(bofFormula,'m_PIGMENT_TOTAL_POOL[c]');

% Read pigment composition dynamically and normalize it.
pZea=coefficientForWorkbookMet_s3(pigFormula,'m_CPD1F-130_Thyl[t]');
% Workbook versions use Plas for chlorophyll-a.
try
    pChl=coefficientForWorkbookMet_s3(pigFormula,'m_CHLOROPHYLL-A_Plas[p]');
catch
    pChl=coefficientForWorkbookMet_s3(pigFormula,'m_CHLOROPHYLL-A_Thyl[t]');
end
pBeta=coefficientForWorkbookMet_s3(pigFormula,'m_CPD1F-129_Thyl[t]');
pPCB=coefficientForWorkbookMet_s3(pigFormula,'m_3Z-PHYCOCYANOBILIN_Plas[p]');
pc=[pZea;pChl;pBeta;pPCB];
pc=pc/sum(pc);

pigMet='m_S3FVA_PIGMENT_TOTAL_AUTO[c]';
pigRxn='S3FVA_PIGMENT_TOTAL_AUTO_PRODUCER_bm';
bioMet='m_S3FVA_TOTAL_BIOMASS_AUTO[c]';
assembly='S3FVA_TOTAL_BIOMASS_AUTO_bm';
sink='S3FVA_TOTAL_BIOMASS_AUTO_SINK_bm_tx';

m=removeIfPresent_s3(m,{pigRxn,assembly,sink});
m=addMetIfMissing_s3(m,pigMet,'publication FVA AUTO pigment pool');
m=addMetIfMissing_s3(m,bioMet,'publication FVA AUTO total biomass');

m=addReaction(m,pigRxn, ...
    'reactionName','publication FVA AUTO pigment pool from TOTAL_BIOMASS.xlsx', ...
    'metaboliteList',{pig.zea;pig.chla;pig.beta;pig.pcb;pigMet}, ...
    'stoichCoeffList',[-pc;1], ...
    'lowerBound',0,'upperBound',1000,'subSystem','Biomass pigment pool');
m=padConstraintColumns_s3(m);

m=addReaction(m,assembly, ...
    'reactionName','publication FVA AUTO mean BOF from TOTAL_BIOMASS_AUTO_MEAN', ...
    'metaboliteList',{mets.protein;mets.carb;mets.lipid;mets.rna;mets.dna; ...
        pigMet;mets.atp;mets.water;mets.adp;mets.pi;mets.proton;bioMet}, ...
    'stoichCoeffList',[-cProtein;-cCarb;-cLipid;-cRNA;-cDNA;-cPig; ...
        -cfg.GAM;-cfg.GAM;cfg.GAM;cfg.GAM;cfg.GAM;1], ...
    'lowerBound',0,'upperBound',1000,'subSystem','Biomass synthesis');
m=padConstraintColumns_s3(m);

m=addReaction(m,sink, ...
    'reactionName','publication FVA AUTO biomass sink', ...
    'metaboliteList',{bioMet},'stoichCoeffList',-1, ...
    'lowerBound',0,'upperBound',1000,'subSystem','Biomass sink');
m=padConstraintColumns_s3(m);

b=struct();
b.mode="AUTO"; b.assemblyRxn=assembly; b.sinkRxn=sink;
b.biomassMet=bioMet; b.pigmentRxn=pigRxn; b.pigmentMet=pigMet;
b.proteinCoeff=cProtein; b.carbohydrateCoeff=cCarb; b.lipidCoeff=cLipid;
b.rnaCoeff=cRNA; b.dnaCoeff=cDNA; b.pigmentCoeff=cPig;
b.GAM=cfg.GAM; b.pigmentComposition=pc;
b.source="TOTAL_BIOMASS.xlsx / TOTAL_BIOMASS_AUTO_MEAN";
cfg.AUTO=b;

Component=["Protein";"Carbohydrate";"Lipid";"RNA";"DNA";"Pigment"];
Coefficient=[cProtein;cCarb;cLipid;cRNA;cDNA;cPig];
Raw_g_gDW=nan(6,1);
Mode=repmat("AUTO",6,1);
Source=repmat("TOTAL_BIOMASS.xlsx / TOTAL_BIOMASS_AUTO_MEAN",6,1);
auditT=table(Mode,Component,Raw_g_gDW,Coefficient,Source);
end

function [m,cfg,auditT]=implementExperimentalBOFs_s3(m,cfg)
[m,pigH]=addExperimentalPigmentPool_s3(m,'HETERO',cfg);
[m,pigM]=addExperimentalPigmentPool_s3(m,'MIXO',cfg);

[m,bH,aH]=createExperimentalBOF_s3(m,'HETERO',cfg.exp.HETERO, ...
    cfg.reference.HETERO.rnaFraction,cfg.reference.HETERO.pigmentMass, ...
    pigH.poolMet,pigH.rxn,cfg);

[m,bM,aM]=createExperimentalBOF_s3(m,'MIXO',cfg.exp.MIXO, ...
    cfg.reference.MIXO.rnaFraction,cfg.reference.MIXO.pigmentMass, ...
    pigM.poolMet,pigM.rxn,cfg);

cfg.HETERO=bH; cfg.MIXO=bM;
auditT=[aH;aM];
end

function [m,info]=addExperimentalPigmentPool_s3(m,mode,cfg)
mode=upper(string(mode));
pig=cfg.resolvedPigmentMets;

if mode=="HETERO"
    poolMet='m_S3FVA_PIGMENT_TOTAL_EXP_HETERO[c]';
    rxn='S3FVA_PIGMENT_TOTAL_EXP_HETERO_PRODUCER_bm';
    sheetName='Reaction List Heto';
    workbookRxn='PIGMENT_TOTAL_POOL_BIO_HETEROTROPH_GLUCOSE_NOFRAC';
elseif mode=="MIXO"
    poolMet='m_S3FVA_PIGMENT_TOTAL_EXP_MIXO[c]';
    rxn='S3FVA_PIGMENT_TOTAL_EXP_MIXO_PRODUCER_bm';
    sheetName='Reaction List Mixo';
    workbookRxn='PIGMENT_TOTAL_POOL_BIO_MIXOTROPH_NOFRAC';
else
    error('Unsupported pigment mode: %s',mode);
end

formula=readWorkbookReactionFormula_s3(cfg.biomassFile,sheetName,workbookRxn);
cZea=coefficientForWorkbookMet_s3(formula,'m_CPD1F-130_Thyl[t]');
try
    cChl=coefficientForWorkbookMet_s3(formula,'m_CHLOROPHYLL-A_Plas[p]');
catch
    cChl=coefficientForWorkbookMet_s3(formula,'m_CHLOROPHYLL-A_Thyl[t]');
end
cBeta=coefficientForWorkbookMet_s3(formula,'m_CPD1F-129_Thyl[t]');
cPCB=coefficientForWorkbookMet_s3(formula,'m_3Z-PHYCOCYANOBILIN_Plas[p]');

coeff=[cZea;cChl;cBeta;cPCB];
if any(~isfinite(coeff)) || sum(coeff)<=0
    error('Invalid %s pigment composition in TOTAL_BIOMASS.xlsx.',mode);
end
coeff=coeff/sum(coeff);

m=removeIfPresent_s3(m,{rxn});
m=addMetIfMissing_s3(m,poolMet,sprintf('publication FVA %s pigment pool',mode));
m=addReaction(m,rxn, ...
    'reactionName',sprintf('publication FVA %s pigment pool from TOTAL_BIOMASS.xlsx',mode), ...
    'metaboliteList',{pig.zea;pig.chla;pig.beta;pig.pcb;poolMet}, ...
    'stoichCoeffList',[-coeff;1], ...
    'lowerBound',0,'upperBound',1000,'subSystem','Biomass pigment pool');
m=padConstraintColumns_s3(m);

info.poolMet=poolMet; info.rxn=rxn; info.coeff=coeff;
info.workbookSheet=sheetName; info.workbookReaction=workbookRxn;
end

function [m,b,auditT]=createExperimentalBOF_s3( ...
    m,mode,expComp,rnaFraction,pigmentMass,pigmentPoolMet,pigmentRxn,cfg)

mode=upper(string(mode));
mets=resolveModeBOFMets_s3(m,mode);

raw=[expComp.protein;expComp.carbohydrate;expComp.lipid; ...
     expComp.nucleic*rnaFraction;expComp.nucleic*(1-rnaFraction); ...
     pigmentMass];
represented=sum(raw);
if represented<=0, error('Invalid %s BOF composition.',mode); end
% Measured macromolecules account for 0.9 g/gDW; do not rescale them.
% Pool-level molecular recipes retain workbook-derived mode specificity.
coeff=raw;
if abs(sum(raw(1:5))-0.9)>1e-8 || any(~isfinite(coeff)) || any(coeff<0) || represented>1+1e-8
    error('Invalid unnormalized experimental BOF for %s.',mode);
end

bioMet=sprintf('m_S3FVA_TOTAL_BIOMASS_EXP_%s[c]',mode);
assembly=sprintf('S3FVA_TOTAL_BIOMASS_EXP_%s_bm',mode);
sink=sprintf('S3FVA_TOTAL_BIOMASS_EXP_%s_SINK_bm_tx',mode);

m=removeIfPresent_s3(m,{assembly,sink});
m=addMetIfMissing_s3(m,bioMet,sprintf('publication FVA experimental %s biomass',mode));

m=addReaction(m,assembly, ...
    'reactionName',sprintf('publication FVA experimental %s BOF',mode), ...
    'metaboliteList',{mets.protein;mets.carb;mets.lipid;mets.rna;mets.dna; ...
        pigmentPoolMet;mets.atp;mets.water;mets.adp;mets.pi;mets.proton;bioMet}, ...
    'stoichCoeffList',[-coeff;-cfg.GAM;-cfg.GAM; ...
        cfg.GAM;cfg.GAM;cfg.GAM;1], ...
    'lowerBound',0,'upperBound',1000,'subSystem','Biomass synthesis');
m=padConstraintColumns_s3(m);

m=addReaction(m,sink, ...
    'reactionName',sprintf('publication FVA experimental %s biomass sink',mode), ...
    'metaboliteList',{bioMet},'stoichCoeffList',-1, ...
    'lowerBound',0,'upperBound',1000,'subSystem','Biomass sink');
m=padConstraintColumns_s3(m);

b=struct();
b.mode=mode; b.assemblyRxn=assembly; b.sinkRxn=sink; b.biomassMet=bioMet;
b.pigmentRxn=pigmentRxn; b.pigmentMet=pigmentPoolMet;
b.proteinCoeff=coeff(1); b.carbohydrateCoeff=coeff(2); b.lipidCoeff=coeff(3);
b.rnaCoeff=coeff(4); b.dnaCoeff=coeff(5); b.pigmentCoeff=coeff(6);
b.GAM=cfg.GAM; b.rawRepresentedMass=represented;
b.rawUnresolvedMass=max(0,1-represented);
b.source="Experimental cultivation-validation BOF";

Component=["Protein";"Carbohydrate";"Lipid";"RNA";"DNA";"Pigment"];
Mode=repmat(mode,6,1);
Coefficient=coeff;
Raw_g_gDW=raw;
Source=repmat("Experimental cultivation-validation BOF",6,1);
auditT=table(Mode,Component,Raw_g_gDW,Coefficient,Source);
end

function mets=resolveModeBOFMets_s3(model,mode)
mode=upper(string(mode));
if ~ismember(mode,["AUTO","MIXO","HETERO"])
    error('Unsupported biomass mode: %s',mode);
end
tag=char(mode);

mets.protein=requireMetCandidates_s3(model,{ ...
    sprintf('m_PROTEIN_TOTAL_POOL_%s_Cyto[c]',tag), ...
    sprintf('PROTEIN_TOTAL_POOL_%s_CYTO',tag), ...
    'm_PROTEIN_TOTAL_POOL_Cyto[c]','PROTEIN_TOTAL_POOL_CYTO'});
mets.carb=requireMetCandidates_s3(model,{ ...
    sprintf('m_CARBOHYDRATE_TOTAL_POOL_%s_Cyto[c]',tag), ...
    sprintf('CARBOHYDRATE_TOTAL_POOL_%s_CYTO',tag), ...
    'm_CARBOHYDRATE_TOTAL_POOL_Cyto[c]','CARBOHYDRATE_TOTAL_POOL_CYTO'});
mets.lipid=requireMetCandidates_s3(model,{ ...
    sprintf('m_LIPID_TOTAL_POOL_%s_Cyto[c]',tag), ...
    sprintf('LIPID_TOTAL_POOL_%s_CYTO',tag), ...
    'm_LIPID_TOTAL_POOL_Cyto[c]','LIPID_TOTAL_POOL_CYTO'});
mets.rna=requireMetCandidates_s3(model,{ ...
    sprintf('m_RNA_TOTAL_POOL_%s_Cyto[c]',tag), ...
    sprintf('RNA_TOTAL_POOL_%s_CYTO',tag), ...
    'm_RNA_TOTAL_POOL_Cyto[c]','RNA_TOTAL_POOL_CYTO'});
mets.dna=requireMetCandidates_s3(model,{ ...
    sprintf('m_DNA_TOTAL_POOL_%s_Cyto[c]',tag), ...
    sprintf('DNA_TOTAL_POOL_%s_CYTO',tag), ...
    'm_DNA_TOTAL_POOL_Cyto[c]','DNA_TOTAL_POOL_CYTO'});

mets.atp=requireMetCandidates_s3(model,{'m_ATP_Cyto[c]','ATP_CYTO'});
mets.water=requireMetCandidates_s3(model,{'m_WATER_Cyto[c]','WATER_CYTO'});
mets.adp=requireMetCandidates_s3(model,{'m_ADP_Cyto[c]','ADP_CYTO'});
mets.pi=requireMetCandidates_s3(model,{'m_Pi_Cyto[c]','PI_CYTO'});
mets.proton=requireMetCandidates_s3(model,{'m_PROTON_Cyto[c]','PROTON_CYTO'});
end

function m=removeIfPresent_s3(m,rxns)
for r=string(rxns(:))'
    if any(strcmp(m.rxns,char(r)))
        m=removeRxns(m,{char(r)});
    end
end
end

function m=addMetIfMissing_s3(m,metID,metName)
if ~any(strcmp(m.mets,metID))
    m=addMetabolite(m,metID,'metName',metName,'metFormula','','Charge',0);
end
end

function m=padConstraintColumns_s3(m)
n=numel(m.rxns);
if isfield(m,'C') && ~isempty(m.C) && size(m.C,2)<n
    m.C(:,end+1:n)=0;
end
end


%% ======================================================================
%% CONDITION SETUP / UPTAKE DIRECTION
%% ======================================================================

function m=prepareCondition_s3(m,mode,cfg)
mode=upper(string(mode));

% Close uptake through all known carbon-containing boundaries first.
for i=1:numel(cfg.carbonBoundaryRxns)
    r=cfg.carbonBoundaryRxns{i};
    if any(strcmp(m.rxns,r))
        m=closeUptake_s3(m,r);
    end
end
for i=1:numel(cfg.allMappedCarbonRxns)
    r=cfg.allMappedCarbonRxns{i};
    if any(strcmp(m.rxns,r))
        m=closeUptake_s3(m,r);
    end
end

% Reapply basal inorganic medium.
basal=struct2cell(cfg.basalRxns);
for i=1:numel(basal)
    idx=find(strcmp(m.rxns,basal{i}),1);
    m.lb(idx)=cfg.basalMediumLB;
    m.ub(idx)=cfg.basalMediumUB;
end

% Fixed NGAM.
idx=find(strcmp(m.rxns,cfg.ngamRxn),1);
m.lb(idx)=cfg.NGAM; m.ub(idx)=cfg.NGAM;

% Light.
if mode=="AUTO"
    m=setUptakeOnly_s3(m,cfg.photonRxn,cfg.autoPhotonProbeMax);
elseif mode=="MIXO"
    m=setUptakeOnly_s3(m,cfg.photonRxn,cfg.mixoPhotonMaxUptake);
elseif mode=="HETERO"
    m=setUptakeOnly_s3(m,cfg.photonRxn,cfg.heteroPhotonMaxUptake);
else
    error('Unsupported mode: %s',mode);
end

% O2 and light-specific dummy.
m=setUptakeOnly_s3(m,cfg.o2Rxn,cfg.o2MaxUptake);
if mode=="AUTO" || mode=="MIXO"
    m=setDirectBounds_s3(m,cfg.o2DummyRxn,cfg.o2DummyLB,cfg.o2DummyUB);
else
    m=setDirectBounds_s3(m,cfg.o2DummyRxn,0,0);
end

% Close all analysis-specific BOF reactions, then open only selected set.
for i=1:numel(cfg.allAnalysisBOFRxns)
    r=cfg.allAnalysisBOFRxns{i};
    if any(strcmp(m.rxns,r)), m=setDirectBounds_s3(m,r,0,0); end
end

if mode=="AUTO", b=cfg.AUTO;
elseif mode=="MIXO", b=cfg.MIXO;
else, b=cfg.HETERO;
end

m=setDirectBounds_s3(m,b.pigmentRxn,0,1000);
m=setDirectBounds_s3(m,b.assemblyRxn,0,1000);
m=setDirectBounds_s3(m,b.sinkRxn,0,1000);
m=setObjective_s3(m,b.sinkRxn);

% CO2 / Rubisco.
if mode=="AUTO"
    m=openUptake_s3(m,cfg.co2Rxn,cfg.autoCO2CarbonCap,true);
    m=setDirectBounds_s3(m,cfg.rubiscoRxn,cfg.rubiscoMinFluxAUTO,1000);
elseif mode=="MIXO"
    m=openUptake_s3(m,cfg.co2Rxn,cfg.mixoCO2MaxUptake,true);
    m=setDirectBounds_s3(m,cfg.rubiscoRxn,cfg.rubiscoMinFluxMIXO,1000);
else
    m=closeUptake_s3(m,cfg.co2Rxn);
    m=setDirectBounds_s3(m,cfg.rubiscoRxn,0,0);
end
end

function m=setDirectBounds_s3(m,rxn,lb,ub)
idx=find(strcmp(m.rxns,rxn),1);
if isempty(idx), error('Reaction not found: %s',rxn); end
m.lb(idx)=lb; m.ub(idx)=ub;
end

function m=setObjective_s3(m,rxn)
idx=find(strcmp(m.rxns,rxn),1);
if isempty(idx), error('Objective reaction not found: %s',rxn); end
m.c=zeros(numel(m.rxns),1);
m.c(idx)=1;
end

function m=clearObjective_s3(m)
m.c=zeros(numel(m.rxns),1);
end

function m=closeUptake_s3(m,rxn)
idx=find(strcmp(m.rxns,rxn),1);
if isempty(idx), return; end
sgn=uptakeFluxSign_s3(m,rxn);
if sgn<0
    m.lb(idx)=max(m.lb(idx),0);
else
    m.ub(idx)=min(m.ub(idx),0);
end
end

function m=openUptake_s3(m,rxn,q,preserveOtherDirection)
if nargin<4, preserveOtherDirection=true; end
idx=find(strcmp(m.rxns,rxn),1);
if isempty(idx), error('Reaction not found: %s',rxn); end
sgn=uptakeFluxSign_s3(m,rxn);
q=abs(q);
if sgn<0
    m.lb(idx)=-q;
    if ~preserveOtherDirection, m.ub(idx)=0; end
else
    m.ub(idx)=q;
    if ~preserveOtherDirection, m.lb(idx)=0; end
end
end

function m=setUptakeOnly_s3(m,rxn,q)
idx=find(strcmp(m.rxns,rxn),1);
if isempty(idx), error('Reaction not found: %s',rxn); end
sgn=uptakeFluxSign_s3(m,rxn);
q=abs(q);
if sgn<0
    m.lb(idx)=-q; m.ub(idx)=0;
else
    m.lb(idx)=0; m.ub(idx)=q;
end
end

function sgn=uptakeFluxSign_s3(model,rxn)
idx=find(strcmp(model.rxns,rxn),1);
if isempty(idx), error('Boundary reaction not found: %s',rxn); end
s=full(model.S(:,idx));
nz=find(abs(s)>1e-12);
if numel(nz)~=1
    error('Expected one-metabolite boundary reaction for uptake sign: %s',rxn);
end
if s(nz)<0, sgn=-1; else, sgn=1; end
end

function q=uptakeMagnitudeFromSolution_s3(model,rxn,x)
idx=find(strcmp(model.rxns,rxn),1);
if isempty(idx), q=NaN; return; end
sgn=uptakeFluxSign_s3(model,rxn);
q=max(0,sgn*x(idx));
end

function [qmin,qmax]=uptakeRangeFromRaw_s3(model,rxn,rawMin,rawMax)
sgn=uptakeFluxSign_s3(model,rxn);
vals=sgn*[rawMin rawMax];
qmin=max(0,min(vals));
qmax=max(0,max(vals));
end


%% ======================================================================
%% HETERO PHOTON CALIBRATION
%% ======================================================================

function [alpha,k,cal]=calibrateHeteroPhoton_s3( ...
    m,biomassRxn,pigmentRxn,glucoseRxn,nC,cfg)

m=openUptake_s3(m,glucoseRxn,cfg.validationMaxCarbon/nC,false);

bidx=find(strcmp(m.rxns,biomassRxn),1);
m.lb(bidx)=cfg.referenceMu_h;
m.ub(bidx)=cfg.referenceMu_h;

pidx=find(strcmp(m.rxns,cfg.photonRxn),1);
m=clearObjective_s3(m);
sgn=uptakeFluxSign_s3(m,cfg.photonRxn);
m.c(pidx)=sgn; % objective equals positive photon uptake magnitude
sol=optimizeCbModel(m,'min');

if ~isFeasibleSolution_s3(sol)
    error('HETERO photon calibration is infeasible at mu=%.6f h^-1.',cfg.referenceMu_h);
end

minPhoton=uptakeMagnitudeFromSolution_s3(m,cfg.photonRxn,sol.x);
pigIdx=find(strcmp(m.rxns,pigmentRxn),1);
pigFlux=sol.x(pigIdx);

if pigFlux<=cfg.fluxTol
    error('HETERO pigment flux is zero during photon calibration.');
end

alpha=minPhoton/pigFlux;
k=minPhoton/cfg.referenceMu_h;

cal=table(cfg.referenceMu_h,minPhoton,pigFlux,alpha,k, ...
    'VariableNames',{'Calibration_mu_h','Minimum_photon_uptake', ...
    'Pigment_pool_flux','Photon_per_pigment','Photon_per_biomass'});
end

function m=addPhotonBiomassCoupling_s3(m,photonRxn,biomassRxn,k)
% Exact Script-2 equality coupling: photon uptake is proportional to biomass.
pidx=find(strcmp(m.rxns,photonRxn),1);
bidx=find(strcmp(m.rxns,biomassRxn),1);
sgn=uptakeFluxSign_s3(m,photonRxn);

row=sparse(1,numel(m.rxns));
row(pidx)=1;
row(bidx)=-sgn*k;

if ~isfield(m,'C') || isempty(m.C)
    m.C=sparse(0,numel(m.rxns));
    m.d=zeros(0,1);
    m.dsense=char(zeros(0,1));
end
if size(m.C,2)<numel(m.rxns)
    m.C(:,end+1:numel(m.rxns))=0;
end

m.C(end+1,:)=row;
m.d(end+1,1)=0;
m.dsense(end+1,1)='E';
if ~isfield(m,'ctrs') || isempty(m.ctrs), m.ctrs=cell(0,1); end
m.ctrs{end+1,1}='S3FVA_HETERO_PHOTON_BIOMASS_COUPLING';
end


%% ======================================================================
%% CAPACITY SCREEN
%% ======================================================================

function out=capacityOne_s3(m,biomassRxn,mode,substrate,subRxn,nC,cfg)
out=emptyCapacityResult_s3();
out.Mode=string(mode);
out.Substrate=string(substrate);
out.Exchange=string(subRxn);
out.CarbonAtoms=nC;

m=setObjective_s3(m,biomassRxn);
sol=optimizeCbModel(m,'max');
out.Feasible=isFeasibleSolution_s3(sol);

if ~out.Feasible
    out.Status="INFEASIBLE";
    out.MaxMu_h=0;
    out.PredictedGrowth=false;
    return;
end

out.Status="OPTIMAL";
out.MaxMu_h=max(0,sol.f);
out.PredictedGrowth=out.MaxMu_h>cfg.growthTol;
out.PhotonUptake=uptakeMagnitudeFromSolution_s3(m,cfg.photonRxn,sol.x);
out.O2Uptake=uptakeMagnitudeFromSolution_s3(m,cfg.o2Rxn,sol.x);

if strcmpi(mode,'AUTO')
    out.SubstrateUptake=NaN;
    out.OrganicCarbon=0;
    out.CO2Uptake=uptakeMagnitudeFromSolution_s3(m,cfg.co2Rxn,sol.x);
else
    out.SubstrateUptake=uptakeMagnitudeFromSolution_s3(m,subRxn,sol.x);
    out.OrganicCarbon=nC*out.SubstrateUptake;
    if strcmpi(mode,'MIXO')
        out.CO2Uptake=uptakeMagnitudeFromSolution_s3(m,cfg.co2Rxn,sol.x);
    else
        out.CO2Uptake=0;
    end
end
out.TotalCarbon=out.OrganicCarbon+out.CO2Uptake;
end

function r=emptyCapacityResult_s3()
r=struct('Mode',"",'Substrate',"",'Exchange',"",'CarbonAtoms',NaN, ...
    'Status',"NOT_RUN",'Feasible',false,'PredictedGrowth',false, ...
    'MaxMu_h',NaN,'SubstrateUptake',NaN,'CO2Uptake',NaN, ...
    'OrganicCarbon',NaN,'TotalCarbon',NaN,'PhotonUptake',NaN,'O2Uptake',NaN);
end

function r=failedCapacityResult_s3(mode,substrate,status)
r=emptyCapacityResult_s3();
r.Mode=string(mode); r.Substrate=string(substrate); r.Status=string(status);
r.MaxMu_h=0; r.PredictedGrowth=false;
end

function tf=isFeasibleSolution_s3(sol)
tf=isstruct(sol) && isfield(sol,'stat') && sol.stat==1 && ...
   isfield(sol,'x') && ~isempty(sol.x);
end

function y=parseGrowthPhenotype_s3(txt)
t=lower(strtrim(string(txt)));
if ismissing(t) || strlength(t)==0
    y=NaN; return;
end
if any(strcmp(t,["no","no growth","negative","-","0","false"]))
    y=0;
elseif any(strcmp(t,["yes","growth","positive","+","++","+++","slow","slow growth","true"]))
    y=1;
elseif contains(t,'no growth')
    y=0;
elseif contains(t,'yes') || contains(t,'growth') || contains(t,'positive')
    y=1;
else
    y=NaN;
end
end

function txt=agreementText_s3(expected,predicted)
if isnan(expected)
    txt="NOT_SCORED";
elseif logical(expected)==logical(predicted)
    txt="MATCH";
else
    txt="MISMATCH";
end
end

function out=fixedMuCarbonAnalysis_s3( ...
    m,biomassRxn,mode,substrate,subRxn,nC,cfg)

out=emptyFixedResult_s3();
out.Mode=string(mode); out.Substrate=string(substrate);
out.Exchange=string(subRxn); out.CarbonAtoms=nC;
out.FixedMu_h=cfg.referenceMu_h;

mode=upper(string(mode));

% Enlarge relevant carbon opening for demand minimization.
if mode=="AUTO"
    % Demand objective represents positive external uptake. During this
    % minimization CO2 is therefore uptake-only; otherwise CO2 secretion
    % could subtract from a signed linear uptake objective.
    m=openUptake_s3(m,cfg.co2Rxn,cfg.validationMaxCarbon,false);
elseif mode=="MIXO"
    m=openUptake_s3(m,cfg.co2Rxn,cfg.validationMaxCarbon,false);
end

% Fixed biomass flux.
bidx=find(strcmp(m.rxns,biomassRxn),1);
m.lb(bidx)=cfg.referenceMu_h;
m.ub(bidx)=cfg.referenceMu_h;

% Objective = total external carbon uptake.
m=clearObjective_s3(m);
if mode=="AUTO"
    cidx=find(strcmp(m.rxns,cfg.co2Rxn),1);
    m.c(cidx)=uptakeFluxSign_s3(m,cfg.co2Rxn);
elseif mode=="HETERO"
    sidx=find(strcmp(m.rxns,subRxn),1);
    m.c(sidx)=nC*uptakeFluxSign_s3(m,subRxn);
else
    sidx=find(strcmp(m.rxns,subRxn),1);
    cidx=find(strcmp(m.rxns,cfg.co2Rxn),1);
    m.c(sidx)=nC*uptakeFluxSign_s3(m,subRxn);
    m.c(cidx)=uptakeFluxSign_s3(m,cfg.co2Rxn);
end

sol=optimizeCbModel(m,'min');
if ~isFeasibleSolution_s3(sol)
    out.Status="FIXED_MU_INFEASIBLE";
    return;
end

out.Status="OPTIMAL";
out.FixedMuFeasible=true;
out.MinimumTotalCarbon=max(0,sol.f);

if mode=="AUTO"
    out.SubstrateUptake=NaN;
    out.CO2Uptake=uptakeMagnitudeFromSolution_s3(m,cfg.co2Rxn,sol.x);
    out.MinimumOrganicCarbon=0;
elseif mode=="HETERO"
    out.SubstrateUptake=uptakeMagnitudeFromSolution_s3(m,subRxn,sol.x);
    out.CO2Uptake=0;
    out.MinimumOrganicCarbon=nC*out.SubstrateUptake;
else
    out.SubstrateUptake=uptakeMagnitudeFromSolution_s3(m,subRxn,sol.x);
    out.CO2Uptake=uptakeMagnitudeFromSolution_s3(m,cfg.co2Rxn,sol.x);

    % Separate MIXO minimum organic C at the same fixed biomass flux.
    mo=clearObjective_s3(m);
    sidx=find(strcmp(mo.rxns,subRxn),1);
    mo.c(sidx)=nC*uptakeFluxSign_s3(mo,subRxn);
    solOrg=optimizeCbModel(mo,'min');
    if isFeasibleSolution_s3(solOrg)
        out.MinimumOrganicCarbon=max(0,solOrg.f);
    end
end

tot=out.MinimumTotalCarbon;
if isfinite(tot) && tot>cfg.uptakeTol
    orgC=0;
    if mode~="AUTO" && isfinite(out.SubstrateUptake)
        orgC=nC*out.SubstrateUptake;
    end
    co2C=max(0,out.CO2Uptake);
    out.OrganicCarbonFraction=orgC/(orgC+co2C);
    out.CO2CarbonFraction=co2C/(orgC+co2C);
end
out.PhotonUptake=uptakeMagnitudeFromSolution_s3(m,cfg.photonRxn,sol.x);

% publication workflow-style carbon FVA: total external carbon <= 105% of minimum.
mFVA=addCarbonBudgetConstraint_s3(m,mode,subRxn,nC, ...
    out.MinimumTotalCarbon*(1+cfg.carbonFVATolerance),cfg);

if mode~="AUTO"
    [mn,mx]=optimizeRxnRawRange_s3(mFVA,subRxn);
    [out.SubstrateFVAmin,out.SubstrateFVAmax]= ...
        uptakeRangeFromRaw_s3(mFVA,subRxn,mn,mx);
end
if mode~="HETERO"
    [mn,mx]=optimizeRxnRawRange_s3(mFVA,cfg.co2Rxn);
    [out.CO2FVAmin,out.CO2FVAmax]= ...
        uptakeRangeFromRaw_s3(mFVA,cfg.co2Rxn,mn,mx);
end
end

function m=addCarbonBudgetConstraint_s3(m,mode,subRxn,nC,budget,cfg)
row=sparse(1,numel(m.rxns));
mode=upper(string(mode));

if mode=="AUTO"
    idx=find(strcmp(m.rxns,cfg.co2Rxn),1);
    row(idx)=uptakeFluxSign_s3(m,cfg.co2Rxn);
elseif mode=="HETERO"
    idx=find(strcmp(m.rxns,subRxn),1);
    row(idx)=nC*uptakeFluxSign_s3(m,subRxn);
else
    idx=find(strcmp(m.rxns,subRxn),1);
    row(idx)=nC*uptakeFluxSign_s3(m,subRxn);
    idx=find(strcmp(m.rxns,cfg.co2Rxn),1);
    row(idx)=uptakeFluxSign_s3(m,cfg.co2Rxn);
end
m=appendLinearConstraint_s3(m,row,budget,'L');
end

function m=appendLinearConstraint_s3(m,row,rhs,sense)
if ~isfield(m,'C') || isempty(m.C)
    m.C=sparse(0,numel(m.rxns));
    m.d=zeros(0,1);
    m.dsense=char(zeros(0,1));
end
if size(m.C,2)<numel(m.rxns)
    m.C(:,end+1:numel(m.rxns))=0;
elseif size(m.C,2)>numel(m.rxns)
    error('model.C has more columns than reactions.');
end
m.C(end+1,:)=row;
m.d(end+1,1)=rhs;
m.dsense(end+1,1)=sense;
if ~isfield(m,'ctrs') || isempty(m.ctrs), m.ctrs=cell(0,1); end
m.ctrs{end+1,1}=sprintf('S3FVA_LINEAR_%d',size(m.C,1));
end

function [mn,mx]=optimizeRxnRawRange_s3(m,rxn)
idx=find(strcmp(m.rxns,rxn),1);
if isempty(idx), mn=NaN; mx=NaN; return; end

m=clearObjective_s3(m); m.c(idx)=1;
s1=optimizeCbModel(m,'min');
s2=optimizeCbModel(m,'max');

if isFeasibleSolution_s3(s1), mn=s1.f; else, mn=NaN; end
if isFeasibleSolution_s3(s2), mx=s2.f; else, mx=NaN; end
end

function r=emptyFixedResult_s3()
r=struct('Mode',"",'Substrate',"",'Exchange',"",'CarbonAtoms',NaN, ...
    'Status',"NOT_RUN",'FixedMu_h',NaN,'FixedMuFeasible',false, ...
    'MinimumTotalCarbon',NaN,'MinimumOrganicCarbon',NaN, ...
    'SubstrateUptake',NaN,'CO2Uptake',NaN, ...
    'OrganicCarbonFraction',NaN,'CO2CarbonFraction',NaN, ...
    'SubstrateFVAmin',NaN,'SubstrateFVAmax',NaN, ...
    'CO2FVAmin',NaN,'CO2FVAmax',NaN,'PhotonUptake',NaN);
end

function r=failedFixedResult_s3(mode,substrate,status)
r=emptyFixedResult_s3();
r.Mode=string(mode); r.Substrate=string(substrate); r.Status=string(status);
end

function rows=chooseTargetSubstrates_s3( ...
    names,classes,exchanges,heteroCapacity,cfg) %#ok<INUSD>
% Require exactly the six publication substrates. No substitution.
rows=zeros(numel(cfg.preferredTargetSubstrates),1);
for k=1:numel(cfg.preferredTargetSubstrates)
    wanted=cfg.preferredTargetSubstrates(k);
    idx=find(strcmpi(strtrim(names),strtrim(wanted)),1);
    if isempty(idx)
        error('Required Script-3 substrate not found: %s',wanted);
    end
    if strlength(exchanges(idx))==0
        error('Required Script-3 substrate has no mapped exchange: %s',wanted);
    end
    if heteroCapacity(idx).MaxMu_h < (cfg.targetMu_h-cfg.growthTol)
        error(['Required substrate %s cannot sustain mu = %.3f h^-1 under the ' ...
               '%.1f C-mmol gDW^-1 h^-1 ceiling. Diagnostic mu_max = %.6f h^-1.'], ...
               wanted,cfg.targetMu_h,cfg.screenCarbon,heteroCapacity(idx).MaxMu_h);
    end
    rows(k)=idx;
end
end


function [summaryT,rxnT]=runTargetedFVA_s3( ...
    baseHetero,biomassRxn,names,classes,exchanges,nC, ...
    expText,doiText,referenceText,rows,cfg) %#ok<INUSD>
% Fixed-mu reaction-level robustness analysis at mu = 0.060 h^-1.
allRows=cell(0,13);
sumRows=cell(numel(rows),12);

for ii=1:numel(rows)
    i=rows(ii);
    sub=names(i);
    r=char(exchanges(i));
    maxUptakeAvailable=cfg.screenCarbon/nC(i);

    m=baseHetero;
    m=openUptake_s3(m,r,maxUptakeAvailable,false);
    bidx=find(strcmp(m.rxns,biomassRxn),1);
    m.lb(bidx)=cfg.targetMu_h;
    m.ub(bidx)=cfg.targetMu_h;

    % Minimum carbon requirement at fixed biomass.
    mMin=clearObjective_s3(m);
    sidx=find(strcmp(mMin.rxns,r),1);
    uptakeSign=uptakeFluxSign_s3(mMin,r);
    mMin.c(sidx)=nC(i)*uptakeSign;
    solMin=optimizeCbModel(mMin,'min');
    if ~isFeasibleSolution_s3(solMin)
        error('FVA infeasible for %s at mu = %.3f h^-1.',sub,cfg.targetMu_h);
    end
    minUptake=uptakeMagnitudeFromSolution_s3(mMin,r,solMin.x);
    minCarbonRequired=nC(i)*minUptake;
    if ~isfinite(minUptake) || minUptake<=cfg.uptakeTol
        error('Invalid minimum uptake for %s at mu = %.3f h^-1.',sub,cfg.targetMu_h);
    end

    uptakeLimit=min(maxUptakeAvailable,minUptake*(1+cfg.targetUptakeTolerance));
    mFVA=m;
    mFVA=openUptake_s3(mFVA,r,uptakeLimit,false);

    [rxnIdx,roles]=selectTargetReactions_s3(mFVA,solMin.x,r,biomassRxn,sub,cfg);
    exRawMin=NaN; exRawMax=NaN;
    pathwayObligatory=strings(0,1);
    pathwayFlexible=strings(0,1);
    respirationStatus=strings(0,1);

    for k=1:numel(rxnIdx)
        j=rxnIdx(k);
        rxn=char(mFVA.rxns{j});
        [mn,mx]=optimizeRxnRawRange_s3(mFVA,rxn);
        referenceFlux=solMin.x(j);
        width=mx-mn;
        zeroIncluded=isfinite(mn) && isfinite(mx) && ...
            (mn<=cfg.fluxTol) && (mx>=-cfg.fluxTol);
        obligatory=isfinite(mn) && isfinite(mx) && ~zeroIncluded;
        if strcmp(rxn,r), exRawMin=mn; exRawMax=mx; end

        rxnName=getRxnName_s3(mFVA,j);
        subsystem=getSubsystem_s3(mFVA,j);
        pubName=publicationReactionName_s3(mFVA,j,roles(k),sub,r,biomassRxn);
        if strcmp(rxn,biomassRxn), fluxUnit="h^-1"; else, fluxUnit="mmol gDW^-1 h^-1"; end
        if zeroIncluded, zeroText="YES"; else, zeroText="NO"; end
        if obligatory, obligatoryText="YES"; else, obligatoryText="NO"; end

        if roles(k)=="Entry" || roles(k)=="Central carbon"
            if obligatory
                pathwayObligatory(end+1,1)=pubName; %#ok<AGROW>
            else
                pathwayFlexible(end+1,1)=pubName; %#ok<AGROW>
            end
        elseif roles(k)=="Respiration" || roles(k)=="ATP synthesis" || roles(k)=="ATP/energy"
            if obligatory
                respirationStatus(end+1,1)=pubName+" - obligatory"; %#ok<AGROW>
            else
                respirationStatus(end+1,1)=pubName+" - zero included"; %#ok<AGROW>
            end
        end

        allRows(end+1,:)={sub,roles(k),pubName,string(rxn),rxnName,subsystem, ...
            fluxUnit,referenceFlux,mn,mx,width,zeroText,obligatoryText}; %#ok<AGROW>
    end

    if ~isfinite(exRawMin), [exRawMin,exRawMax]=optimizeRxnRawRange_s3(mFVA,r); end
    [fvaUptakeMin,fvaUptakeMax]=uptakeRangeFromRaw_s3(mFVA,r,exRawMin,exRawMax);

    if isempty(pathwayObligatory), obligatorySummary="None among selected pathway reactions";
    else, obligatorySummary=strjoin(unique(pathwayObligatory,'stable'),'; '); end
    if isempty(pathwayFlexible), flexibleSummary="None among selected pathway reactions";
    else, flexibleSummary=strjoin(unique(pathwayFlexible,'stable'),'; '); end
    if isempty(respirationStatus), respiratorySummary="No selected respiratory/ATP reaction available";
    else, respiratorySummary=strjoin(unique(respirationStatus,'stable'),'; '); end

    sumRows(ii,:)={sub,classes(i),expText(i),cfg.targetMu_h,minUptake, ...
        minCarbonRequired,uptakeLimit,fvaUptakeMin,fvaUptakeMax, ...
        obligatorySummary,flexibleSummary,respiratorySummary};
end

summaryT=cell2table(sumRows,'VariableNames',{ ...
    'Substrate','Class','Experimental_growth','Fixed_biomass_h_1', ...
    'Minimum_substrate_uptake_mmol_gDW_h','Minimum_C_required_Cmmol_gDW_h', ...
    'FVA_uptake_limit_105pct_mmol_gDW_h','FVA_uptake_min_mmol_gDW_h', ...
    'FVA_uptake_max_mmol_gDW_h','Obligatory_pathway_reactions', ...
    'Reactions_with_zero_in_FVA_interval','Respiration_ATP_requirement'});

rxnT=cell2table(allRows,'VariableNames',{ ...
    'Substrate','Role','Biochemical_reaction','Reaction_ID','Reaction_name', ...
    'Subsystem','Flux_unit','Flux_at_minimum_carbon','FVA_min','FVA_max', ...
    'FVA_range','Zero_in_FVA_interval','Obligatory'});
end


function T=reorderFvaSummary_s3(T)
wanted={ ...
    'Substrate','Class','Experimental_growth','Fixed_biomass_h_1', ...
    'Minimum_substrate_uptake_mmol_gDW_h','Minimum_C_required_Cmmol_gDW_h', ...
    'FVA_uptake_limit_105pct_mmol_gDW_h','FVA_uptake_min_mmol_gDW_h', ...
    'FVA_uptake_max_mmol_gDW_h','Obligatory_pathway_reactions', ...
    'Reactions_with_zero_in_FVA_interval','Respiration_ATP_requirement'};
T=T(:,wanted);
end


function T=buildStep3MethodsTable_s3(cfg)
Setting=[ ...
    "Condition"
    "Representative substrates"
    "Maximum organic carbon availability"
    "Fixed biomass"
    "Minimum-carbon calculation"
    "FVA uptake limit"
    "Reaction panel"
    "NGAM"
    "GAM"
    "Biomass objective ID"
    "External CO2"
    "Rubisco"
    ];
Value=[ ...
    "HETERO"
    "D-glucose; D-xylose; meso-erythritol; glycerol; acetate; pyruvate"
    sprintf('%.1f C-mmol gDW^-1 h^-1',cfg.screenCarbon)
    sprintf('mu = %.3f h^-1',cfg.targetMu_h)
    "Minimize substrate carbon uptake at fixed biomass"
    sprintf('<= %.0f%% of minimum substrate uptake',100*(1+cfg.targetUptakeTolerance))
    sprintf('%d-%d biologically informative reactions per substrate', ...
        cfg.minTargetReactionsPerSubstrate,cfg.maxTargetReactionsPerSubstrate)
    sprintf('%.3f mmol ATP gDW^-1 h^-1',cfg.NGAM)
    sprintf('%.1f mmol ATP gDW^-1 biomass',cfg.GAM)
    string(cfg.HETERO.sinkRxn)
    "Closed for uptake; secretion preserved"
    "Disabled"
    ];
T=table(Setting,Value);
end


function [idxOut,rolesOut]=selectTargetReactions_s3( ...
    m,x,exchangeRxn,biomassRxn,substrateName,cfg)
% Select 5-8 biologically interpretable reactions per substrate.
%
% Priority:
%   1. substrate exchange
%   2. first substrate-entry reaction(s)
%   3. one or two central-carbon reactions
%   4. respiration
%   5. ATP generation
%   6. biomass
%
% The exact six publication substrates use curated reaction panels.

maxN=cfg.maxTargetReactionsPerSubstrate;
minN=cfg.minTargetReactionsPerSubstrate;

ids=string(m.rxns(:));
nrxn=numel(ids);
chosen=false(nrxn,1);
role=strings(nrxn,1);

    function addRxn(rxnID,rxnRole)
        j=find(strcmp(ids,string(rxnID)),1);
        if ~isempty(j) && ~chosen(j)
            chosen(j)=true;
            role(j)=string(rxnRole);
        end
    end

% Always include exchange and biomass.
addRxn(exchangeRxn,"Substrate uptake");
addRxn(biomassRxn,"Biomass");

sub=lower(strtrim(string(substrateName)));

% Curated panels for the current six publication cases.
if strcmpi(sub,"d-glucose")
    addRxn('GLUCOKIN-RXN_Cyto',"Entry");
    addRxn('GLU6PDEHYDROG-RXN_Cyto',"Central carbon");
    addRxn('2PGADEHYDRAT-RXN_Cyto',"Central carbon");
    addRxn('1.10.2.2-RXN_Mito',"Respiration");
    addRxn('Complex-V_Mito',"ATP synthesis");

elseif strcmpi(sub,"d-xylose")
    addRxn('Beta-D-XYLOSE_TO_D-Xylopyranose_ARTIF_Cyto',"Entry");
    addRxn('RXN-8773_Cyto',"Entry");
    addRxn('RIBULP3EPIM-RXN_Cyto',"Central carbon");
    addRxn('RIB5PISOM-RXN_Cyto',"Central carbon");
    addRxn('1.10.2.2-RXN_Mito',"Respiration");
    addRxn('Complex-V_Mito',"ATP synthesis");

elseif contains(sub,"erythritol")
    addRxn('RXN-17763_Cyto',"Entry");
    addRxn('RXN-17761_Cyto',"Entry");
    addRxn('GAPOXNPHOSPHN-RXN_Cyto',"Central carbon");
    addRxn('GLU6PDEHYDROG-RXN_Cyto',"Central carbon");
    addRxn('1.10.2.2-RXN_Mito',"Respiration");
    addRxn('Complex-V_Mito',"ATP synthesis");

elseif strcmpi(sub,"glycerol")
    addRxn('GLYCEROL-KIN-RXN_Cyto',"Entry");
    addRxn('RXN0-5260_Cyto',"Entry");
    addRxn('GAPOXNPHOSPHN-RXN_Cyto',"Central carbon");
    addRxn('2PGADEHYDRAT-RXN_Cyto',"Central carbon");
    addRxn('1.10.2.2-RXN_Mito',"Respiration");
    addRxn('Complex-V_Mito',"ATP synthesis");

elseif strcmpi(sub,"acetate")
    addRxn('ACETATE--COA-LIGASE-ADP-FORMING-RXN_Cyto',"Entry");
    addRxn('MALSYN-RXN_Cyto',"Central carbon");
    addRxn('PEPCARBOXYKIN-RXN_Cyto',"Central carbon");
    addRxn('1.10.2.2-RXN_Mito',"Respiration");
    addRxn('Complex-V_Mito',"ATP synthesis");

elseif strcmpi(sub,"pyruvate")
    addRxn('PYRUVATE_H+_Mito_tx',"Entry");
    addRxn('PYRUVATEORTHOPHOSPHATE-DIKINASE-RXN_Cyto',"Central carbon");
    addRxn('CITSYN-RXN_Mito',"Central carbon");
    addRxn('1.10.2.2-RXN_Mito',"Respiration");
    addRxn('Complex-V_Mito',"ATP synthesis");
end

% ------------------------------------------------------------------
% Generic fallback / fill-in.
% This also protects the script if an exact curated reaction is absent.
% ------------------------------------------------------------------

% First active reactions directly connected to the exchanged metabolite.
exIdx=find(strcmp(ids,string(exchangeRxn)),1);
if ~isempty(exIdx)
    seedMets=find(abs(full(m.S(:,exIdx)))>cfg.fluxTol);
    localRxns=find(any(abs(m.S(seedMets,:))>cfg.fluxTol,1))';
    localRxns=localRxns(localRxns~=exIdx);
    activeLocal=localRxns(abs(x(localRxns))>cfg.fluxTol);

    [~,ord]=sort(abs(x(activeLocal)),'descend');
    activeLocal=activeLocal(ord);
    for j=activeLocal'
        if sum(chosen)>=maxN, break; end
        if nnz(abs(m.S(:,j))>cfg.fluxTol)>1 && ~chosen(j)
            chosen(j)=true;
            role(j)="Entry";
            if sum(role=="Entry")>=2, break; end
        end
    end
end

% Central carbon candidates, preferably active in the FBA optimum.
if sum(chosen)<maxN
    if isfield(m,'rxnNames'), names=upper(string(m.rxnNames(:)));
    else, names=repmat("",nrxn,1); end
    if isfield(m,'subSystems'), subs=upper(flattenSubsystems_s3(m.subSystems));
    else, subs=repmat("",nrxn,1); end
    txt=upper(ids)+" "+names+" "+subs;

    centralKeys=["GLYCOL","PENTOSE","TRANSKET","TRANSALD","PYRUV", ...
        "CITR","ISOCIT","SUCC","FUM","MALATE","OXALO","GLYCER", ...
        "GLYOX","PEPCARBOXY","2PGA","GAPOX"];
    isCentral=false(nrxn,1);
    for kw=centralKeys
        isCentral=isCentral | contains(txt,kw);
    end
    cand=find(isCentral & abs(x)>cfg.fluxTol & ~chosen);
    [~,ord]=sort(abs(x(cand)),'descend');
    cand=cand(ord);
    for j=cand'
        if sum(chosen)>=maxN, break; end
        if sum(role=="Central carbon")>=2, break; end
        chosen(j)=true;
        role(j)="Central carbon";
    end
end

% Ensure one respiration reaction.
if ~any(role=="Respiration")
    prefResp=["1.10.2.2-RXN_Mito","NADH-DEHYDROG-A-RXN_Mito"];
    for rr=prefResp
        j=find(strcmp(ids,rr),1);
        if ~isempty(j)
            chosen(j)=true; role(j)="Respiration"; break;
        end
    end
end

% Ensure one ATP-synthesis reaction.
if ~any(role=="ATP synthesis")
    j=find(strcmp(ids,"Complex-V_Mito"),1);
    if ~isempty(j)
        chosen(j)=true; role(j)="ATP synthesis";
    else
        j=find(strcmp(ids,"ATP_ADP_Mito_tx"),1);
        if ~isempty(j), chosen(j)=true; role(j)="ATP/energy"; end
    end
end

% If still below five, fill with most active internal reactions. These are
% labeled simply "Additional active" rather than exposing selection logic.
if sum(chosen)<minN
    isBoundary=false(nrxn,1);
    for j=1:nrxn
        isBoundary(j)=nnz(abs(m.S(:,j))>cfg.fluxTol)==1;
    end
    cand=find(abs(x)>cfg.fluxTol & ~isBoundary & ~chosen);
    [~,ord]=sort(abs(x(cand)),'descend');
    cand=cand(ord);
    for j=cand'
        if sum(chosen)>=minN, break; end
        chosen(j)=true;
        role(j)="Additional active";
    end
end

% Enforce maximum of eight while preserving biological priority.
idx=find(chosen);
priority=zeros(numel(idx),1);
for k=1:numel(idx)
    switch role(idx(k))
        case "Substrate uptake", priority(k)=1;
        case "Entry", priority(k)=2;
        case "Central carbon", priority(k)=3;
        case "Respiration", priority(k)=4;
        case {"ATP synthesis","ATP/energy"}, priority(k)=5;
        case "Biomass", priority(k)=6;
        otherwise, priority(k)=7;
    end
end

% Preserve role priority; within a role prefer higher absolute FBA flux.
[~,ord]=sortrows([priority,-abs(x(idx))],[1 2]);
idx=idx(ord);
if numel(idx)>maxN
    idx=idx(1:maxN);
end

% Make sure biomass survives truncation.
bidx=find(strcmp(ids,string(biomassRxn)),1);
if ~isempty(bidx) && ~ismember(bidx,idx)
    if numel(idx)>=maxN
        idx(end)=bidx;
    else
        idx(end+1)=bidx;
    end
end

idxOut=idx(:);
rolesOut=role(idxOut);
end

function s=appendReason_s3(s,newReason)
if strlength(s)==0
    s=string(newReason);
elseif ~contains(s,string(newReason))
    s=s+";"+string(newReason);
end
end

function name=getRxnName_s3(m,j)
if isfield(m,'rxnNames') && numel(m.rxnNames)>=j
    name=string(m.rxnNames{j});
else
    name="";
end
end

function name=publicationReactionName_s3(m,j,role,substrate,exchangeRxn,biomassRxn)
rxn=string(m.rxns{j});
if strcmp(rxn,string(exchangeRxn)), name=string(substrate)+" uptake"; return; end
if strcmp(rxn,string(biomassRxn)), name="Biomass formation"; return; end
switch rxn
    case "GLUCOKIN-RXN_Cyto", name="Glucokinase";
    case "GLU6PDEHYDROG-RXN_Cyto", name="Glucose-6-phosphate dehydrogenase";
    case "2PGADEHYDRAT-RXN_Cyto", name="Enolase";
    case "GLYCEROL-KIN-RXN_Cyto", name="Glycerol kinase";
    case "MALSYN-RXN_Cyto", name="Malate synthase";
    case "PEPCARBOXYKIN-RXN_Cyto", name="Phosphoenolpyruvate carboxykinase";
    case "CITSYN-RXN_Mito", name="Mitochondrial citrate synthase";
    case "1.10.2.2-RXN_Mito", name="Mitochondrial respiratory Complex III";
    case "Complex-V_Mito", name="Mitochondrial ATP synthase (Complex V)";
    case "PYRUVATE_H+_Mito_tx", name="Mitochondrial pyruvate transport";
    otherwise
        modelName=getRxnName_s3(m,j);
        if strlength(strtrim(modelName))>0, name=modelName; else, name=string(role); end
end
end

function s=getSubsystem_s3(m,j)
s="";
if isfield(m,'subSystems') && numel(m.subSystems)>=j
    v=m.subSystems{j};
    if iscell(v), s=strjoin(string(v),'; ');
    else, s=string(v); end
end
end

function s=flattenSubsystems_s3(v)
if isstring(v)
    s=v(:);
    return;
elseif ischar(v)
    s=string(cellstr(v));
    return;
end
s=strings(numel(v),1);
for i=1:numel(v)
    try
        x=v{i};
    catch
        x=v(i);
    end
    if iscell(x), s(i)=strjoin(string(x),'; ');
    else, s(i)=string(x); end
end
end

function [lb,ub]=bounds_s3(m,rxn)
idx=find(strcmp(m.rxns,rxn),1); lb=m.lb(idx); ub=m.ub(idx);
end
```
