## Supplementary Data 3 for "Experimental trophic flexibility guides a compartmentalized genome-scale model of red microalgae *Galdieria sulphuraria*": S4_Galdieria_publication_HVP.m

```
% 4_Galdieria_publication_HVP
%
% Publication high-value-product substrate optimization.
%
% The selected publication products are optimized separately with every
% mapped substrate in Model_Substrate_list.xlsx under HETERO and MIXO.
% Organic substrate supply is normalized to 6 C-mmol gDW^-1 h^-1.
%
% The script performs two deliberately separate analyses:
%
% A. PRIMARY THEORETICAL PRODUCTION-CAPACITY SCREEN
%   1) opens one organic-carbon substrate at a time;
%   2) calculates substrate-specific maximum growth;
%   3) maximizes each selected HVP at >=50% of that substrate-specific mu_max;
%   4) compares product capacity against D-glucose within the same trophic state.
%      These values are product-flux capacities and are NOT carbon yields.
%
% B. SECONDARY STANDARDIZED FIXED-GROWTH CARBON-EFFICIENCY ANALYSIS
%   5) fixes biomass exactly at mu = 0.060 h^-1;
%   6) re-maximizes each selected HVP under the same substrate supply;
%   7) reports HETERO yield per organic C and MIXO yields both per organic C
%      and per total assimilated C (organic C + CO2-C).
%
% The two analyses are exported and summarized separately so production
% capacity is never mixed with fixed-growth carbon efficiency.
%
% HETERO: organic substrate only; external CO2 uptake and Rubisco closed.
% MIXO:   the same organic substrate plus CO2 and light.
%
% Storage alpha-glucan is treated separately from extracellular substrates.
% Its 'substrate' case represents mobilization of an intracellular reserve
% assumed to have accumulated during a preceding AUTO phase. An analysis-only
% reserve source and lumped hydrolysis step (STARCH + H2O -> glucose) are used
% to make this phase-switch assumption explicit without changing the SBML file.
%
% Inputs are loaded automatically from the shared Inputs folder:
%   Galdieria_sulphuraria_SBML_Final.xml
%   Model_Substrate_list.xlsx
%   TOTAL_BIOMASS.xlsx
%
% Outputs:
%   4_Galdieria_publication_HVP.xlsx
%   4_Galdieria_publication_HVP_HETERO_vs_glucose_heatmap.png
%   4_Galdieria_publication_HVP_MIXO_vs_glucose_heatmap.png
%   4_Galdieria_publication_HVP_HETERO_fixed_growth_yields.png
%   4_Galdieria_publication_HVP_MIXO_fixed_growth_yields.png

%% PATHS / INPUTS
scriptDir = fileparts(mfilename('fullpath'));
if isempty(scriptDir), scriptDir = pwd; end
supplementDir = fileparts(fileparts(scriptDir));
inputDir = fullfile(supplementDir,'Inputs');

modelInput = fullfile(inputDir,'Galdieria_sulphuraria_SBML_Final.xml');
substrateFile = fullfile(inputDir,'Model_Substrate_list.xlsx');
biomassFile = fullfile(inputDir,'TOTAL_BIOMASS.xlsx');

modelInput = resolveLocalFile_hvp(modelInput,inputDir);
substrateFile = resolveLocalFile_hvp(substrateFile,inputDir);
biomassFile = resolveLocalFile_hvp(biomassFile,inputDir);

if ~isfile(modelInput), error('Model file not found: %s',modelInput); end
if ~isfile(substrateFile), error('Substrate workbook not found: %s',substrateFile); end
if ~isfile(biomassFile), error('Biomass workbook not found: %s',biomassFile); end

ensureCobraReady_hvp();

fprintf('\n============================================================\n');
fprintf('Galdieria publication HVP substrate optimization\n');
fprintf('============================================================\n');

baseModel = loadModel_hvp(modelInput);
validateBasicModel_hvp(baseModel);

%% CONFIGURATION
cfg.scriptDir = scriptDir;
cfg.modelFile = string(modelInput);
cfg.substrateFile = string(substrateFile);
cfg.biomassFile = string(biomassFile);
cfg.substrateSheet = 'Sole_substrate_ScrumPy';
cfg.resultsFile = fullfile(scriptDir,'4_Galdieria_publication_HVP.xlsx');
cfg.heteroHeatmapFile = fullfile(scriptDir,'4_Galdieria_publication_HVP_HETERO_vs_glucose_heatmap.png');
cfg.mixoHeatmapFile = fullfile(scriptDir,'4_Galdieria_publication_HVP_MIXO_vs_glucose_heatmap.png');
cfg.heteroFixedGrowthYieldFigure = fullfile(scriptDir,'4_Galdieria_publication_HVP_HETERO_fixed_growth_yields.png');
cfg.mixoFixedGrowthYieldFigure = fullfile(scriptDir,'4_Galdieria_publication_HVP_MIXO_fixed_growth_yields.png');
cfg.heatmapReferenceSubstrate = "D-Glucose";
cfg.heteroHeatmapAnnotationThresholdPct = 5;
cfg.mixoHeatmapAnnotationThresholdPct = 2;

cfg.referenceMu_d = 1.44;
cfg.referenceMu_h = cfg.referenceMu_d/24;
cfg.fixedMu_h = cfg.referenceMu_h;          % secondary standardized carbon-efficiency analysis (0.060 h^-1)
cfg.NGAM = 1.7;
cfg.GAM = 30;
cfg.screenCarbon = 6;
cfg.validationMaxCarbon = 120;

cfg.o2MaxUptake = 1000;
cfg.heteroPhotonMaxUptake = 0.2;
cfg.mixoPhotonMaxUptake = 50;
cfg.mixoCO2MaxUptake = cfg.validationMaxCarbon;
cfg.rubiscoMinFluxMIXO = 0.01;
cfg.autoPhotonMaxUptake = 20;
cfg.autoCO2MaxUptake = cfg.screenCarbon;
cfg.rubiscoMinFluxAUTO = 0.01;
cfg.noCarbonTol = 1e-8;

% Trophic-state product sets used for all state-specific HVP calculations,
% summaries, fixed-growth matrices, full-state analyses and heatmaps.
cfg.selectedProducts.AUTO = ["C18_2","BetaCarotene","PCB","Phytol","Glycogen"];
cfg.selectedProducts.MIXO = ["Glycogen","TAG","C18_2","PCB","Ergosterol"];
cfg.selectedProducts.HETERO = ["Glycogen","TAG","Floridoside","Ergosterol"];

cfg.expectedPhotonPerPigment = 0.58;
cfg.photonCalibrationTolerance = 0.02;
cfg.productGrowthFraction = 0.50;
cfg.fluxTol = 1e-9;
cfg.growthTol = 1e-8;
cfg.yieldTol = 1e-12;
cfg.strictCarbonBoundaryClosure = true;

cfg.basalMediumLB = -1000;
cfg.basalMediumUB = 1000;
cfg.o2DummyLB = 0;
cfg.o2DummyUB = 1000;

cfg.exp.MIXO.protein       = 0.4147;
cfg.exp.MIXO.carbohydrate  = 0.3377;
cfg.exp.MIXO.nucleic       = 0.0775;
cfg.exp.MIXO.lipid         = 0.0701;

cfg.exp.HETERO.protein      = 0.4016;
cfg.exp.HETERO.carbohydrate = 0.3609;
cfg.exp.HETERO.nucleic      = 0.0765;
cfg.exp.HETERO.lipid        = 0.0610;

cfg.reference.MIXO.rnaFraction = 0.06222707/(0.06222707+0.01091703);
cfg.reference.HETERO.rnaFraction = 0.085/(0.085+0.015);
cfg.reference.HETERO.pigmentMass = 0.00485;
cfg.reference.MIXO.PCB_mg  = 1.714658;
cfg.reference.MIXO.CHLA_mg = 4.450000;
cfg.reference.MIXO.BETA_mg = 0.240000;
cfg.reference.MIXO.ZEA_mg  = 0.350000;
cfg.reference.MIXO.pigmentMass = ...
    (cfg.reference.MIXO.PCB_mg + cfg.reference.MIXO.CHLA_mg + ...
     cfg.reference.MIXO.BETA_mg + cfg.reference.MIXO.ZEA_mg)/1000;

%% SUBSTRATES
Tsub = readtable(substrateFile,'Sheet',cfg.substrateSheet, ...
    'VariableNamingRule','preserve');

requiredCols = {'Substrate_ID','Class','MetaCyc / model exchange ID'};
for k = 1:numel(requiredCols)
    if ~ismember(requiredCols{k},Tsub.Properties.VariableNames)
        error('Required column missing from %s: %s',cfg.substrateSheet,requiredCols{k});
    end
end

substrateName = strtrim(string(Tsub.('Substrate_ID')));
substrateClass = strtrim(string(Tsub.('Class')));
requestedExchange = strtrim(string(Tsub.('MetaCyc / model exchange ID')));

% Replace database-style substrate labels with reader-facing chemical names.
substrateName(strcmpi(substrateName,'CPD-15829')) = "alpha-D-Ribopyranose";
substrateName(strcmpi(substrateName,'CPD-25028')) = "alpha-D-Xylopyranose";

resolvedExchange = strings(height(Tsub),1);
carbonAtoms = nan(height(Tsub),1);
carbonSource = strings(height(Tsub),1);

for i = 1:height(Tsub)
    if strlength(substrateName(i))==0 || strlength(requestedExchange(i))==0
        continue;
    end
    resolvedExchange(i) = string(resolveRxn_hvp(baseModel,requestedExchange(i),true));
    if strlength(resolvedExchange(i))==0
        warning('Substrate exchange could not be resolved: %s -> %s', ...
            substrateName(i),requestedExchange(i));
        continue;
    end
    [carbonAtoms(i),carbonSource(i)] = inferCarbonAtoms_hvp( ...
        baseModel,char(resolvedExchange(i)),substrateName(i),substrateClass(i));
    if ~isfinite(carbonAtoms(i)) || carbonAtoms(i)<=0
        error(['Carbon count could not be resolved for mapped substrate "%s" ' ...
            '(%s). Carbon-normalized HVP optimization cannot continue.'], ...
            substrateName(i),resolvedExchange(i));
    end
end

validSubstrate = strlength(resolvedExchange)>0 & isfinite(carbonAtoms) & carbonAtoms>0;
if ~any(validSubstrate)
    error('No mapped substrates with resolvable carbon counts were found.');
end

substrateName = substrateName(validSubstrate);
substrateClass = substrateClass(validSubstrate);
requestedExchange = requestedExchange(validSubstrate);
resolvedExchange = resolvedExchange(validSubstrate);
carbonAtoms = carbonAtoms(validSubstrate);
carbonSource = carbonSource(validSubstrate);
uptakeCap = cfg.screenCarbon ./ carbonAtoms;

% Starch is not interpreted as an extracellular substrate in this analysis.
% It represents intracellular storage alpha-glucan accumulated previously
% (e.g. during AUTO) and mobilized after switching to HETERO or MIXO.
isReserveSubstrate = contains(lower(substrateName),'starch') | ...
    contains(lower(resolvedExchange),'starch_cyto_tx');
carbonSourceRole = repmat("External organic substrate",numel(substrateName),1);
carbonSourceRole(isReserveSubstrate) = "Intracellular reserve";
reserveAssumption = repmat("",numel(substrateName),1);
reserveAssumption(isReserveSubstrate) = ...
    "Storage alpha-glucan assumed accumulated during preceding AUTO phase; mobilized during subsequent HETERO/MIXO phase.";

cfg.allMappedCarbonRxns = cellstr(unique(resolvedExchange,'stable'));

glcRow = find(strcmpi(substrateName,'D-Glucose'),1);
if isempty(glcRow)
    error('D-Glucose must be present for HETERO photon calibration.');
end
cfg.glucoseRxn = char(resolvedExchange(glcRow));
cfg.glucoseCarbonAtoms = carbonAtoms(glcRow);

fprintf('Mapped substrates to analyse: %d\n',numel(substrateName));

analysisSourceReaction = resolvedExchange;

substrateReference = table(substrateName,substrateClass,requestedExchange, ...
    resolvedExchange,analysisSourceReaction,carbonSourceRole,carbonAtoms, ...
    carbonSource,uptakeCap,reserveAssumption, ...
    'VariableNames',{'Substrate','Class','Requested_exchange', ...
    'Resolved_exchange','Analysis_source_reaction','Carbon_source_role', ...
    'Carbon_atoms','Carbon_count_source','Organic_uptake_cap_mmol', ...
    'Reserve_assumption'});

%% MODEL MAPPING
cfg.ngamRxn = requireRxn_hvp(baseModel, ...
    {'R_NGAM_ATP_Cyto','NGAM_ATP_Cyto','ATPM_NGAM_Cyto'});
cfg.photonRxn = requireRxn_hvp(baseModel, ...
    {'R_Photon_Cyto_tx','Photon_Cyto_tx'});
cfg.co2Rxn = requireRxn_hvp(baseModel, ...
    {'R_CO2_Cyto_tx','CO2_Cyto_tx'});
cfg.o2Rxn = requireRxn_hvp(baseModel, ...
    {'R_O2_Cyto_tx','O2_Cyto_tx'});
cfg.o2DummyRxn = requireRxn_hvp(baseModel, ...
    {'R_O2_Dummy_Cyto_tx','O2_Dummy_Cyto_tx'});
cfg.rubiscoRxn = requireRxn_hvp(baseModel,{ ...
    'R_RIBULOSE-BISPHOSPHATE-CARBOXYLASE-RXN_Plas', ...
    'RIBULOSE-BISPHOSPHATE-CARBOXYLASE-RXN_Plas'});

cfg.basalRxns.Fe2 = requireRxn_hvp(baseModel,{'R_FE+2_Cyto_tx','FE+2_Cyto_tx'});
cfg.basalRxns.Mg2 = requireRxn_hvp(baseModel,{'R_MG+2_Cyto_tx','MG+2_Cyto_tx'});
cfg.basalRxns.NH4 = requireRxn_hvp(baseModel,{'R_NH4_Cyto_tx','NH4_Cyto_tx'});
cfg.basalRxns.NO3 = requireRxn_hvp(baseModel,{'R_NO3_Cyto_tx','NO3_Cyto_tx'});
cfg.basalRxns.H   = requireRxn_hvp(baseModel,{'R_PROTON_Cyto_tx','PROTON_Cyto_tx'});
cfg.basalRxns.Pi  = requireRxn_hvp(baseModel,{'R_Pi_Cyto_tx','Pi_Cyto_tx'});
cfg.basalRxns.SO4 = requireRxn_hvp(baseModel,{'R_SO4_Cyto_tx','SO4_Cyto_tx'});
cfg.basalRxns.H2O = requireRxn_hvp(baseModel,{'R_WATER_Cyto_tx','WATER_Cyto_tx'});

% Analysis-only intracellular reserve module. This does not modify the SBML
% input on disk. It supplies stored alpha-glucan to the intracellular pool
% and represents its net hydrolytic mobilization as a glucose-equivalent unit.
[baseModel,cfg] = addStorageReserveModule_hvp( ...
    baseModel,cfg,resolvedExchange,isReserveSubstrate);
if any(isReserveSubstrate)
    analysisSourceReaction(isReserveSubstrate) = string(cfg.reserveSourceRxn);
    substrateReference.Analysis_source_reaction = analysisSourceReaction;
end

[cfg.carbonBoundaryRxns,~] = identifyCarbonBoundaryRxns_hvp(baseModel,cfg);
cfg.carbonBoundaryRxns = cellstr(unique([ ...
    string(cfg.carbonBoundaryRxns(:)); ...
    string(cfg.allMappedCarbonRxns(:)); ...
    string(cfg.co2Rxn)],'stable'));

%% BIOMASS FUNCTIONS
[baseModel,~] = closeOriginalBiomassOutputs_hvp(baseModel);
cfg = resolveBiomassMetabolites_hvp(baseModel,cfg);
[baseModel,cfg,~] = implementExperimentalBOFs_hvp(baseModel,cfg);
[baseModel,cfg,~] = implementAutoWorkbookBOF_hvp(baseModel,cfg,biomassFile);

% The latest trophic-state SBML contains condition-specific TAG biomass pools
% but retains the generic native TAG output reaction used by the publication
% HVP analysis. Recreate the former generic TAG producer in-memory when it is
% absent, using the HETERO TAG composition already encoded in the final SBML.
% This restores the published product definition without modifying the SBML.
baseModel = ensurePublicationTagProductAssembly_hvp(baseModel);

cfg.allAnalysisBOFRxns = { ...
    cfg.AUTO.assemblyRxn,cfg.AUTO.sinkRxn,cfg.AUTO.pigmentRxn, ...
    cfg.MIXO.assemblyRxn,cfg.MIXO.sinkRxn,cfg.MIXO.pigmentRxn, ...
    cfg.HETERO.assemblyRxn,cfg.HETERO.sinkRxn,cfg.HETERO.pigmentRxn};

%% PRODUCTS
products = defineProducts_hvp();
[products,productReference] = resolveProducts_hvp(baseModel,products);
baseModel = closeAllProductOutputs_hvp(baseModel,products);

% Enforce trophic-state HVP sets at the calculation level. Products not
% assigned to a trophic state are not optimized in that state's substrate
% screen or fixed-growth analysis.
productsHETERO = selectProducts_hvp(products,cfg.selectedProducts.HETERO);
productsMIXO = selectProducts_hvp(products,cfg.selectedProducts.MIXO);
productsByMode = {productsHETERO,productsMIXO};

%% AUTO / HETERO / MIXO BASE MODELS
modelAUTO = prepareCondition_hvp(baseModel,'AUTO',cfg);
modelMIXO = prepareCondition_hvp(baseModel,'MIXO',cfg);
modelHETERO_uncoupled = prepareCondition_hvp(baseModel,'HETERO',cfg);

assertModeBase_hvp(modelAUTO,'AUTO',cfg);
assertModeBase_hvp(modelMIXO,'MIXO',cfg);
assertModeBase_hvp(modelHETERO_uncoupled,'HETERO',cfg);

[photonPerPigment,photonPerBiomass,~] = calibrateHeteroPhoton_hvp( ...
    modelHETERO_uncoupled,cfg.HETERO.sinkRxn,cfg.HETERO.pigmentRxn, ...
    cfg.glucoseRxn,cfg.glucoseCarbonAtoms,cfg);

cfg.HETERO.photonPerPigment = photonPerPigment;
cfg.HETERO.photonPerBiomass = photonPerBiomass;

if abs(photonPerPigment-cfg.expectedPhotonPerPigment) > ...
        cfg.photonCalibrationTolerance
    fprintf(2,['NOTICE: HETERO photon/pigment calibration = %.6g; ' ...
        'reference expectation %.6g +/- %.6g. Calculated value retained.\n'], ...
        photonPerPigment,cfg.expectedPhotonPerPigment, ...
        cfg.photonCalibrationTolerance);
end

modelHETERO = addPhotonBiomassCoupling_hvp( ...
    modelHETERO_uncoupled,cfg.photonRxn,cfg.HETERO.sinkRxn, ...
    photonPerBiomass);

modeNames = ["HETERO","MIXO"];
modeModels = {modelHETERO,modelMIXO};
biomassRxns = {cfg.HETERO.sinkRxn,cfg.MIXO.sinkRxn};

%% SUBSTRATE-SPECIFIC GROWTH AND PRODUCT OPTIMIZATION
nS = numel(substrateName);
nP = numel(products);
nM = numel(modeNames);
nStateProducts = [numel(productsHETERO),numel(productsMIXO)];

muMax = nan(nS,nM);
detailRows = cell(nS*sum(nStateProducts),30);
rr = 0;

fprintf('\nOptimizing state-specific HVP sets across %d substrates: HETERO=%d, MIXO=%d...\n', ...
    nS,nStateProducts(1),nStateProducts(2));

for s = 1:nS
    subRxn = char(resolvedExchange(s));
    nC = carbonAtoms(s);
    qSub = uptakeCap(s);

    fprintf('  %2d/%2d  %s\n',s,nS,substrateName(s));

    for mIdx = 1:nM
        mode = modeNames(mIdx);
        bioRxn = biomassRxns{mIdx};

        isReserve = isReserveSubstrate(s);
        [mBase,sourceRxn] = openSingleSubstrate_hvp( ...
            modeModels{mIdx},subRxn,qSub,mode,cfg,isReserve);

        mGrowth = closeAllProductOutputs_hvp(mBase,products);
        mGrowth = setObjective_hvp(mGrowth,bioRxn);
        solGrowth = optimizeCbModel(mGrowth,'max');

        if isFeasibleSolution_hvp(solGrowth) && solGrowth.f > cfg.growthTol
            muMax(s,mIdx) = solGrowth.f;
        else
            muMax(s,mIdx) = 0;
        end

        modeProducts = productsByMode{mIdx};
        for p = 1:numel(modeProducts)
            rr = rr + 1;
            masterPidx = findProductKey_hvp(products,modeProducts(p).key);
            P = products(masterPidx);

            growthFloor = NaN;
            achievedGrowth = NaN;
            p50 = NaN;
            retained = NaN;
            subUptake = NaN;
            organicC = NaN;
            co2Uptake = NaN;
            totalCarbon = NaN;
            yieldPC = NaN;
            productCFlux = NaN;
            carbonYield = NaN;
            photonUptake = NaN;
            o2Uptake = NaN;
            pMax = NaN;
            analysisNote = "";

            % When storage alpha-glucan itself is the carbon reserve,
            % optimizing the same pool as the product is not meaningful.
            % All other HVPs are optimized using reserve mobilization.
            reserveSelfProduct = isReserve && strcmpi(string(P.key),'Glycogen');

            if reserveSelfProduct
                analysisNote = "Not evaluated: storage alpha-glucan is the intracellular carbon reserve itself in this phase-switch scenario.";
            else
                % Product capacity without a growth floor.
                m0 = openOnlyProductOutput_hvp(mBase,products,masterPidx);
                m0 = setBiomassFloor_hvp(m0,bioRxn,0);
                m0 = setObjective_hvp(m0,P.outputRxn);
                sol0 = optimizeCbModel(m0,'max');
                pMax = solutionObjectiveOrZero_hvp(sol0);

                if isReserve
                    analysisNote = "Intracellular storage alpha-glucan reserve mobilization after a preceding AUTO phase; not extracellular starch uptake.";
                end

                if muMax(s,mIdx) > cfg.growthTol
                    growthFloor = cfg.productGrowthFraction * muMax(s,mIdx);

                    m50 = openOnlyProductOutput_hvp(mBase,products,masterPidx);
                    m50 = setBiomassFloor_hvp(m50,bioRxn,growthFloor);
                    m50 = setObjective_hvp(m50,P.outputRxn);
                    sol50 = optimizeCbModel(m50,'max');

                    if isFeasibleSolution_hvp(sol50)
                        p50 = solutionObjectiveOrZero_hvp(sol50);
                        bidx = find(strcmp(m50.rxns,bioRxn),1);
                        achievedGrowth = sol50.x(bidx);

                        [subUptake,organicC,co2Uptake,totalCarbon, ...
                            photonUptake,o2Uptake] = ...
                            substrateUptakeMetrics_hvp( ...
                                m50,mode,sol50,sourceRxn,nC,cfg);

                        [yieldPC,carbonYield,productCFlux] = ...
                            productYieldsGeneral_hvp( ...
                                p50,P.carbonEquivalent, ...
                                totalCarbon,cfg.yieldTol);
                    end
                end

                if isfinite(p50) && isfinite(pMax) && pMax > cfg.fluxTol
                    retained = p50/pMax;
                end
            end

            detailRows(rr,:) = { ...
                substrateName(s),substrateClass(s),mode, ...
                string(subRxn),string(sourceRxn),carbonSourceRole(s), ...
                nC,qSub, ...
                muMax(s,mIdx),growthFloor,achievedGrowth, ...
                P.class,P.label,P.doi, ...
                P.outputRxn,P.carbonEquivalent, ...
                pMax,p50,retained, ...
                subUptake,organicC,co2Uptake,totalCarbon, ...
                yieldPC,productCFlux,carbonYield,photonUptake,o2Uptake, ...
                reserveAssumption(s),analysisNote};
        end
    end
end

results = cell2table(detailRows,'VariableNames',{ ...
    'Substrate','Substrate_class','Mode','Exchange', ...
    'Analysis_source_reaction','Carbon_source_role','Carbon_atoms', ...
    'Organic_uptake_cap_mmol','Mu_max_h','Growth_floor_h', ...
    'Achieved_growth_h','Product_class','Product','DOI', ...
    'Native_output_reaction','Product_C_equivalent', ...
    'Max_product_no_growth_floor','Product_at_50pct_mu_max', ...
    'Fraction_of_unconstrained_product_retained', ...
    'Organic_source_flux_mmol','Organic_source_C_mmolC', ...
    'CO2_uptake_mmolC','Total_C_input_mmolC', ...
    'Yield_product_mmol_per_C_mmol','Product_C_flux_mmolC', ...
    'Carbon_yield_Cmol_per_Cmol','Photon_uptake','O2_uptake', ...
    'Reserve_assumption','Analysis_note'});

% State-specific product selection is enforced during optimization, so the
% export table already contains only the HVPs assigned to each trophic state.
resultsExport = results;

heteroFlux = buildSubstrateMatrix_hvp( ...
    results,substrateReference,products,'HETERO','Product_at_50pct_mu_max');
mixoFlux = buildSubstrateMatrix_hvp( ...
    results,substrateReference,products,'MIXO','Product_at_50pct_mu_max');
heteroYield = buildSubstrateMatrix_hvp( ...
    results,substrateReference,products,'HETERO','Carbon_yield_Cmol_per_Cmol');
mixoYield = buildSubstrateMatrix_hvp( ...
    results,substrateReference,products,'MIXO','Carbon_yield_Cmol_per_Cmol');

% Additional publication-facing fixed-growth comparison. Each substrate is
% re-optimized only for the HVPs assigned to that trophic state at the same
% biomass flux (mu = 0.060 h^-1), so carbon yields can be compared without
% substrate-specific growth floors.
fixedMuHeteroYield = buildFixedMuYieldMatrix_hvp( ...
    modelHETERO,'HETERO',cfg.HETERO.sinkRxn, ...
    substrateName,substrateClass,resolvedExchange,carbonAtoms,uptakeCap, ...
    isReserveSubstrate,productsHETERO,products,cfg);
fixedMuMixoYield = buildFixedMuYieldMatrix_hvp( ...
    modelMIXO,'MIXO',cfg.MIXO.sinkRxn, ...
    substrateName,substrateClass,resolvedExchange,carbonAtoms,uptakeCap, ...
    isReserveSubstrate,productsMIXO,products,cfg);

% Reader-facing Supplementary fixed-growth tables are rounded to three
% decimals. Internal calculations retain full precision.
fixedMuHeteroYieldExport = roundNumericTable_hvp(fixedMuHeteroYield,3);
fixedMuMixoYieldExport = roundNumericTable_hvp(fixedMuMixoYield,3);

% Secondary fixed-growth summaries are calculated exclusively from the
% exact-mu matrices above. They are never derived from the >=50% mu_max
% capacity screen.
fixedMuHeteroSummary = buildFixedMuBestSummary_hvp( ...
    fixedMuHeteroYield,productsHETERO,'HETERO',cfg.heatmapReferenceSubstrate);
fixedMuMixoSummary = buildFixedMuBestSummary_hvp( ...
    fixedMuMixoYield,productsMIXO,'MIXO',cfg.heatmapReferenceSubstrate);

% Publication-facing post-analysis. Relative matrices use the master product
% schema, while only state-assigned HVPs contain calculated values.
heteroVsGlucose = buildReferenceRelativeMatrix_hvp( ...
    heteroFlux,productsHETERO,cfg.heatmapReferenceSubstrate);
mixoVsGlucose = buildReferenceRelativeMatrix_hvp( ...
    mixoFlux,productsMIXO,cfg.heatmapReferenceSubstrate);
heteroHeatmapLimitPct = relativeHeatmapLimit_hvp(heteroVsGlucose,productsHETERO);
mixoHeatmapLimitPct = relativeHeatmapLimit_hvp(mixoVsGlucose,productsMIXO);

hvpSummary = buildBestSolutionSummary_hvp( ...
    heteroFlux,mixoFlux,heteroYield,mixoYield, ...
    heteroVsGlucose,mixoVsGlucose,products,cfg.heatmapReferenceSubstrate);

% Publication-facing summaries follow the trophic-state product sets.
% Non-assigned HVP columns remain NaN in the master-schema matrices and are
% excluded from all state-specific statistics.
commonSelectedKeys = intersect(cfg.selectedProducts.HETERO, ...
    cfg.selectedProducts.MIXO,'stable');
productsCommon = selectProducts_hvp(products,commonSelectedKeys);

classSummary = buildSubstrateClassSummary_hvp( ...
    heteroFlux,mixoFlux,heteroYield,mixoYield, ...
    heteroVsGlucose,mixoVsGlucose,productsHETERO,productsMIXO, ...
    productsCommon,cfg.fluxTol,cfg.yieldTol);

overallSummary = buildOverallTrophicSummary_hvp( ...
    heteroFlux,mixoFlux,productsCommon,cfg.fluxTol);

% Compact reader-facing supplementary substrate summary: one row per
% substrate, summarizing the response across all selected HVPs. The full
% substrate x product x condition optimization table remains available in
% MATLAB as "results" but is not exported.
suppResults = buildCompactSubstrateResults_hvp( ...
    heteroFlux,mixoFlux,heteroYield,mixoYield, ...
    heteroVsGlucose,mixoVsGlucose,muMax,products, ...
    cfg.fluxTol,cfg.yieldTol);

% Remove technical implementation columns from the reader-facing Excel file.
% They remain available internally to the analysis code.
substrateReferenceExport = substrateReference(:,{ ...
    'Substrate','Class','Requested_exchange','Carbon_atoms', ...
    'Carbon_count_source','Organic_uptake_cap_mmol'});
productReferenceExport = productReference(:,{ ...
    'Class','Product','Target_metabolite','Product_C_equivalent', ...
    'Carbon_equivalent_note','DOI','Evidence_note','Model_unit_note'});
substrateReferenceExport = roundNumericTable_hvp(substrateReferenceExport,3);
productReferenceExport = roundNumericTable_hvp(productReferenceExport,3);

methodsTable = buildSubstrateHVPMethods_hvp( ...
    cfg,nS,nP,photonPerPigment,photonPerBiomass, ...
    heteroHeatmapLimitPct,mixoHeatmapLimitPct);

%% LITERATURE-GUIDED FULL TROPHIC-STATE FBA
% Use D-glucose for the full MIXO/HETERO FBA and CO2 only for AUTO.
qGlucose = cfg.screenCarbon / cfg.glucoseCarbonAtoms;
[modelHETERO_Full,~] = openSingleSubstrate_hvp( ...
    modelHETERO,cfg.glucoseRxn,qGlucose,'HETERO',cfg,false);
[modelMIXO_Full,~] = openSingleSubstrate_hvp( ...
    modelMIXO,cfg.glucoseRxn,qGlucose,'MIXO',cfg,false);
modelAUTO_Full = modelAUTO;

fullModeNames = ["AUTO","MIXO","HETERO"];
fullModeModels = {modelAUTO_Full,modelMIXO_Full,modelHETERO_Full};
fullBiomassRxns = {cfg.AUTO.sinkRxn,cfg.MIXO.sinkRxn,cfg.HETERO.sinkRxn};
fullMuMax = nan(1,3);
for c = 1:3
    mg = closeAllProductOutputs_hvp(fullModeModels{c},products);
    mg = setObjective_hvp(mg,fullBiomassRxns{c});
    sg = optimizeCbModel(mg,'max');
    if ~isFeasibleSolution_hvp(sg) || sg.f <= cfg.growthTol
        error('%s full HVP condition has no positive growth.',fullModeNames(c));
    end
    fullMuMax(c) = sg.f;
end

fullAUTO = runFullStateFBA_hvp(modelAUTO_Full,'AUTO',cfg.AUTO.sinkRxn, ...
    cfg.selectedProducts.AUTO,products,fullMuMax(1),cfg);
fullMIXO = runFullStateFBA_hvp(modelMIXO_Full,'MIXO',cfg.MIXO.sinkRxn, ...
    cfg.selectedProducts.MIXO,products,fullMuMax(2),cfg);
fullHETERO = runFullStateFBA_hvp(modelHETERO_Full,'HETERO',cfg.HETERO.sinkRxn, ...
    cfg.selectedProducts.HETERO,products,fullMuMax(3),cfg);

%% EXCEL OUTPUT -- SIX READER-FACING SHEETS
if isfile(cfg.resultsFile), delete(cfg.resultsFile); end

% Split the substrate-by-product results by trophic state for export. Each
% table already contains only the HVPs assigned to that trophic state.
resultsHETEROExport = resultsExport(strcmpi(string(resultsExport.Mode),'HETERO'),:);
resultsMIXOExport = resultsExport(strcmpi(string(resultsExport.Mode),'MIXO'),:);

% 1) Publication summary.
% Biological interpretation is trophic-state specific. HETERO and MIXO
% best-substrate summaries therefore contain only the literature-guided HVPs
% assigned to that condition. Products not selected for a state are not shown
% in that state's biological summary.
heteroLabels = string({productsHETERO.label});
mixoLabels = string({productsMIXO.label});
heteroKeep = ismember(string(hvpSummary.Product),heteroLabels);
mixoKeep = ismember(string(hvpSummary.Product),mixoLabels);

capacitySummaryHETERO = hvpSummary(heteroKeep,{ ...
    'Product','HETERO_best_flux_substrate', ...
    'HETERO_best_flux_change_vs_reference_pct'});
capacitySummaryMIXO = hvpSummary(mixoKeep,{ ...
    'Product','MIXO_best_flux_substrate', ...
    'MIXO_best_flux_change_vs_reference_pct'});
% Secondary fixed-growth summaries come only from the exact-mu analysis.
yieldSummaryHETERO = fixedMuHeteroSummary(:,{ ...
    'Product','Best_organicC_substrate','Best_organicC_yield', ...
    'Reference_organicC_yield','OrganicC_change_vs_reference_pct'});
yieldSummaryMIXO = fixedMuMixoSummary(:,{ ...
    'Product','Best_totalC_substrate','Best_totalC_yield', ...
    'Reference_totalC_yield','TotalC_change_vs_reference_pct', ...
    'Best_organicC_substrate','Best_organicC_yield'});

capacitySummaryHETERO = roundNumericTable_hvp(capacitySummaryHETERO,3);
capacitySummaryMIXO = roundNumericTable_hvp(capacitySummaryMIXO,3);
yieldSummaryHETERO = roundNumericTable_hvp(yieldSummaryHETERO,3);
yieldSummaryMIXO = roundNumericTable_hvp(yieldSummaryMIXO,3);

writecell({ ...
    'A. HETERO BEST PRODUCTION CAPACITY'; ...
    'Best substrate = highest product flux while growth is maintained at >=50% of that substrate''s maximum. Improvement is relative to D-glucose.' ...
    },cfg.resultsFile,'Sheet','HVP_SUMMARY','Range','A1');
writecell({'HVP','Best HETERO substrate','Improvement vs D-glucose (%)'}, ...
    cfg.resultsFile,'Sheet','HVP_SUMMARY','Range','A4');
writetable(capacitySummaryHETERO,cfg.resultsFile,'Sheet','HVP_SUMMARY', ...
    'Range','A5','WriteVariableNames',false);

rowMixoCap = 5 + height(capacitySummaryHETERO) + 3;
writecell({ ...
    'B. MIXO BEST PRODUCTION CAPACITY'; ...
    'Best substrate = highest product flux while growth is maintained at >=50% of that substrate''s maximum. Improvement is relative to D-glucose.' ...
    },cfg.resultsFile,'Sheet','HVP_SUMMARY','Range',sprintf('A%d',rowMixoCap));
writecell({'HVP','Best MIXO substrate','Improvement vs D-glucose (%)'}, ...
    cfg.resultsFile,'Sheet','HVP_SUMMARY','Range',sprintf('A%d',rowMixoCap+3));
writetable(capacitySummaryMIXO,cfg.resultsFile,'Sheet','HVP_SUMMARY', ...
    'Range',sprintf('A%d',rowMixoCap+4),'WriteVariableNames',false);

rowHeteroYield = rowMixoCap + 4 + height(capacitySummaryMIXO) + 3;
writecell({ ...
    'C. SECONDARY HETERO FIXED-GROWTH CARBON EFFICIENCY'; ...
    sprintf('Exact biomass flux: mu = %.3f h^-1. Yield = C-mol product-C per C-mol organic carbon actually taken up.',cfg.fixedMu_h) ...
    },cfg.resultsFile,'Sheet','HVP_SUMMARY','Range',sprintf('A%d',rowHeteroYield));
writecell({'HVP','Best HETERO substrate','Best organic-C yield', ...
    'D-glucose organic-C yield','Change vs D-glucose (%)'}, ...
    cfg.resultsFile,'Sheet','HVP_SUMMARY','Range',sprintf('A%d',rowHeteroYield+3));
writetable(yieldSummaryHETERO,cfg.resultsFile,'Sheet','HVP_SUMMARY', ...
    'Range',sprintf('A%d',rowHeteroYield+4),'WriteVariableNames',false);

rowMixoYield = rowHeteroYield + 4 + height(yieldSummaryHETERO) + 3;
writecell({ ...
    'D. SECONDARY MIXO FIXED-GROWTH CARBON EFFICIENCY'; ...
    sprintf('Exact biomass flux: mu = %.3f h^-1. Total-C yield uses organic C + assimilated CO2-C. Organic-C-normalized yield is also reported as an apparent process metric.',cfg.fixedMu_h) ...
    },cfg.resultsFile,'Sheet','HVP_SUMMARY','Range',sprintf('A%d',rowMixoYield));
writecell({'HVP','Best MIXO substrate by total-C yield','Best total-C yield', ...
    'D-glucose total-C yield','Change vs D-glucose (%)', ...
    'Best substrate by apparent organic-C yield','Best apparent organic-C yield'}, ...
    cfg.resultsFile,'Sheet','HVP_SUMMARY','Range',sprintf('A%d',rowMixoYield+3));
writetable(yieldSummaryMIXO,cfg.resultsFile,'Sheet','HVP_SUMMARY', ...
    'Range',sprintf('A%d',rowMixoYield+4),'WriteVariableNames',false);

overallKeep = ismember(string(overallSummary.Setting),[ ...
    "Unique model carbon-source/reserve reactions", ...
    "Matched substrate-product flux comparisons", ...
    "Median MIXO vs HETERO product-flux change"]);
overallSummaryExport = overallSummary(overallKeep,:);

rowOverall = rowMixoYield + 4 + height(yieldSummaryMIXO) + 3;
writecell({ ...
    'E. OVERALL TROPHIC-STATE SUMMARY'; ...
    'Cross-state product-flux comparisons use only HVPs selected in both HETERO and MIXO: glycogen, TAG and ergosterol.' ...
    },cfg.resultsFile,'Sheet','HVP_SUMMARY','Range',sprintf('A%d',rowOverall));
writecell({'Metric','Result'},cfg.resultsFile,'Sheet','HVP_SUMMARY', ...
    'Range',sprintf('A%d',rowOverall+3));
writetable(overallSummaryExport,cfg.resultsFile,'Sheet','HVP_SUMMARY', ...
    'Range',sprintf('A%d',rowOverall+4),'WriteVariableNames',false);

classSummaryExport = classSummary(:,{ ...
    'Substrate_class','Unique_model_sources', ...
    'HETERO_median_change_vs_glucose_pct', ...
    'MIXO_median_change_vs_glucose_pct', ...
    'Median_MIXO_vs_HETERO_flux_change_pct'});
classSummaryExport = roundNumericTable_hvp(classSummaryExport,3);
rowClass = rowOverall + 4 + height(overallSummaryExport) + 3;
writecell({ ...
    'F. SUBSTRATE-CLASS EFFECT'; ...
    'HETERO medians use the four HETERO-selected products; MIXO medians use the five MIXO-selected products. Cross-state medians use only glycogen, TAG and ergosterol, which are selected in both states.'; ...
    'Within-state columns compare each substrate class with D-glucose; 0.000% means no median difference after rounding.' ...
    },cfg.resultsFile,'Sheet','HVP_SUMMARY','Range',sprintf('A%d',rowClass));
writecell({ ...
    'Substrate class','Carbon sources (n)','HETERO HVP change vs glucose (%)', ...
    'MIXO HVP change vs glucose (%)','MIXO vs HETERO HVP change (%)' ...
    },cfg.resultsFile,'Sheet','HVP_SUMMARY','Range',sprintf('A%d',rowClass+3));
writetable(classSummaryExport,cfg.resultsFile,'Sheet','HVP_SUMMARY', ...
    'Range',sprintf('A%d',rowClass+4),'WriteVariableNames',false);

% 2) AUTO-specific HVP analysis. AUTO has no organic-substrate screen, so
% only the literature-guided full-state HVP FBA is required here.
writeAutoStateFBASheet_hvp(cfg.resultsFile,'AUTO_HVP',fullAUTO, ...
    'AUTO HVP FBA: C18:2, beta-carotene, PCB, phytol, glycogen');

% 3) HETERO-specific sheet: substrate-by-product capacity screen, fixed-mu
% carbon-yield matrix, and the compact D-glucose full-state HVP FBA block.
writeConsolidatedStateSheet_hvp(cfg.resultsFile,'HETERO_HVP','HETERO', ...
    resultsHETEROExport,fixedMuHeteroYieldExport,fullHETERO,cfg);

% 4) MIXO-specific sheet: substrate-by-product capacity screen, fixed-mu
% carbon-yield matrix with organic-C/CO2-C uptake, and the compact D-glucose
% full-state HVP FBA block.
writeConsolidatedStateSheet_hvp(cfg.resultsFile,'MIXO_HVP','MIXO', ...
    resultsMIXOExport,fixedMuMixoYieldExport,fullMIXO,cfg);

% 5) Carbon-source / intracellular-reserve mapping.
writecell({ ...
    'Substrate','Substrate class','Model substrate reaction','Carbon atoms', ...
    'Carbon-count basis','Flux limit (mmol gDW^-1 h^-1)' ...
    },cfg.resultsFile,'Sheet','SUBSTRATE_REFERENCE','Range','A1');
writetable(substrateReferenceExport,cfg.resultsFile,'Sheet','SUBSTRATE_REFERENCE', ...
    'Range','A2','WriteVariableNames',false);

% 6) Methods and product-reference information combined in one sheet.
writecell({'METHODS'},cfg.resultsFile,'Sheet','METHODS_PRODUCT_REFERENCE','Range','A1');
writecell({'Analysis setting','Value / description'},cfg.resultsFile, ...
    'Sheet','METHODS_PRODUCT_REFERENCE','Range','A2');
writetable(methodsTable,cfg.resultsFile,'Sheet','METHODS_PRODUCT_REFERENCE', ...
    'Range','A3','WriteVariableNames',false);
rowProductRef = height(methodsTable) + 6;
writecell({'PRODUCT REFERENCE'},cfg.resultsFile,'Sheet','METHODS_PRODUCT_REFERENCE', ...
    'Range',sprintf('A%d',rowProductRef));
writecell({ ...
    'Product class','HVP','Model target metabolite','Product carbon equivalent', ...
    'Carbon-equivalent basis','DOI','Evidence / interpretation','Model flux unit' ...
    },cfg.resultsFile,'Sheet','METHODS_PRODUCT_REFERENCE', ...
    'Range',sprintf('A%d',rowProductRef+1));
writetable(productReferenceExport,cfg.resultsFile,'Sheet','METHODS_PRODUCT_REFERENCE', ...
    'Range',sprintf('A%d',rowProductRef+2),'WriteVariableNames',false);

%% D-GLUCOSE-RELATIVE STATE-SPECIFIC SUBSTRATE-EFFECT HEATMAPS
% The plotting function enforces the trophic-state product sets internally:
% HETERO = glycogen, TAG, floridoside, ergosterol (4 products)
% MIXO   = glycogen, TAG, C18:2, PCB, ergosterol (5 products)
createReferenceRelativeHeatmap_hvp(heteroVsGlucose,products,'HETERO', ...
    cfg.heteroHeatmapFile,cfg.productGrowthFraction,heteroHeatmapLimitPct, ...
    cfg.heteroHeatmapAnnotationThresholdPct,cfg.heatmapReferenceSubstrate);
createReferenceRelativeHeatmap_hvp(mixoVsGlucose,products,'MIXO', ...
    cfg.mixoHeatmapFile,cfg.productGrowthFraction,mixoHeatmapLimitPct, ...
    cfg.mixoHeatmapAnnotationThresholdPct,cfg.heatmapReferenceSubstrate);

%% FIXED-GROWTH HVP CARBON-YIELD BAR CHARTS
% These figures use the same trophic-state-specific fixed-mu matrices exported
% in section B of HETERO_HVP and MIXO_HVP. Substrate-label colors are assigned
% automatically from the median HVP yield change relative to D-glucose.
createFixedGrowthYieldBarChart_hvp( ...
    fixedMuHeteroYield,productsHETERO,'HETERO', ...
    cfg.heteroFixedGrowthYieldFigure,cfg.heatmapReferenceSubstrate,cfg.fixedMu_h);
createFixedGrowthYieldBarChart_hvp( ...
    fixedMuMixoYield,productsMIXO,'MIXO', ...
    cfg.mixoFixedGrowthYieldFigure,cfg.heatmapReferenceSubstrate,cfg.fixedMu_h);

fprintf('\nDONE -- HVP substrate optimization.\n');
fprintf('Substrates analysed: %d\n',nS);
fprintf('Products analysed:   %d\n',nP);
fprintf('Conditions:          HETERO, MIXO\n');
fprintf('Excel workbook:      %s\n',cfg.resultsFile);
fprintf(['Excel sheets:         HVP_SUMMARY | AUTO_HVP | HETERO_HVP | MIXO_HVP | ' ...
    'SUBSTRATE_REFERENCE | METHODS_PRODUCT_REFERENCE\n']);
fprintf('Heatmap reference:    %s\n',cfg.heatmapReferenceSubstrate);
fprintf('HETERO heatmap scale: +/- %.0f%%\n',heteroHeatmapLimitPct);
fprintf('MIXO heatmap scale:   +/- %.0f%%\n',mixoHeatmapLimitPct);
fprintf('HETERO heatmap:      %s\n',cfg.heteroHeatmapFile);
fprintf('MIXO heatmap:        %s\n',cfg.mixoHeatmapFile);
fprintf('HETERO fixed-growth yield chart: %s\n',cfg.heteroFixedGrowthYieldFigure);
fprintf('MIXO fixed-growth yield chart:   %s\n',cfg.mixoFixedGrowthYieldFigure);


%% LITERATURE-GUIDED EXPORT / FULL-STATE FBA HELPERS

function T=applyLiteratureProductSelection_hvp(T,products,selected)
labels=string({products.label});
keys=string({products.key});
zeroVars={ ...
    'Achieved_growth_h', ...
    'Max_product_no_growth_floor', ...
    'Product_at_50pct_mu_max', ...
    'Fraction_of_unconstrained_product_retained', ...
    'Organic_source_flux_mmol', ...
    'Organic_source_C_mmolC', ...
    'CO2_uptake_mmolC', ...
    'Total_C_input_mmolC', ...
    'Yield_product_mmol_per_C_mmol', ...
    'Product_C_flux_mmolC', ...
    'Carbon_yield_Cmol_per_Cmol', ...
    'Photon_uptake','O2_uptake'};

for r=1:height(T)
    pidx=find(strcmp(labels,string(T.Product(r))),1);
    if isempty(pidx)
        error('Could not map substrate-result product: %s',string(T.Product(r)));
    end
    key=keys(pidx);
    mode=upper(string(T.Mode(r)));
    if mode=="HETERO"
        keep=any(strcmpi(key,selected.HETERO));
    elseif mode=="MIXO"
        keep=any(strcmpi(key,selected.MIXO));
    else
        keep=true;
    end
    if ~keep
        for j=1:numel(zeroVars)
            vn=zeroVars{j};
            if ismember(vn,T.Properties.VariableNames)
                T.(vn)(r)=0;
            end
        end
        if ismember('Analysis_note',T.Properties.VariableNames)
            T.Analysis_note(r)= ...
                "Not selected for this trophic state by literature-guided HVP classification.";
        end
    end
end
end

function T=runFullStateFBA_hvp(model,mode,biomassRxn,selectedKeys,products,muMax,cfg)
n=numel(selectedKeys);
rows=cell(n,30);
for i=1:n
    pidx=findProductKey_hvp(products,selectedKeys(i));
    P=products(pidx);

    m0=openOnlyProductOutput_hvp(model,products,pidx);
    m0=setBiomassFloor_hvp(m0,biomassRxn,0);
    m0=setObjective_hvp(m0,P.outputRxn);
    sol0=optimizeCbModel(m0,'max');
    pMax=solutionObjectiveOrZero_hvp(sol0);
    [glc0,co20,totalC0,ph0,o20]=fullStateUptakeMetrics_hvp(m0,mode,sol0,cfg); %#ok<ASGLU>
    [ypc0,pcc0,pC0,ypg0]=fullStateYields_hvp( ...
        pMax,P.carbonEquivalent,totalC0,glc0,cfg.yieldTol); %#ok<ASGLU>

    growthFloor=cfg.productGrowthFraction*muMax;
    m50=openOnlyProductOutput_hvp(model,products,pidx);
    m50=setBiomassFloor_hvp(m50,biomassRxn,growthFloor);
    m50=setObjective_hvp(m50,P.outputRxn);
    sol50=optimizeCbModel(m50,'max');
    p50=solutionObjectiveOrZero_hvp(sol50);

    achieved=NaN;
    if isFeasibleSolution_hvp(sol50)
        bidx=find(strcmp(m50.rxns,biomassRxn),1);
        achieved=sol50.x(bidx);
    end
    [glc,co2,totalC,photon,o2]=fullStateUptakeMetrics_hvp(m50,mode,sol50,cfg);
    [ypc50,pcc50,pC50,ypg50]=fullStateYields_hvp( ...
        p50,P.carbonEquivalent,totalC,glc,cfg.yieldTol);

    mNC=openOnlyProductOutput_hvp(model,products,pidx);
    mNC=closeAllCarbonUptakeFull_hvp(mNC,cfg);
    mNC=setBiomassFloor_hvp(mNC,biomassRxn,0);
    mNC=setObjective_hvp(mNC,P.outputRxn);
    solNC=optimizeCbModel(mNC,'max');
    pNC=solutionObjectiveOrZero_hvp(solNC);

    retained=NaN;
    if pMax>cfg.fluxTol
        retained=p50/pMax;
    end

    if ~isFeasibleSolution_hvp(sol0)
        status="INFEASIBLE_NO_GROWTH_FLOOR";
    elseif ~isFeasibleSolution_hvp(sol50)
        status="INFEASIBLE_AT_50pct_mu";
    elseif pNC>cfg.noCarbonTol
        status="SUSPICIOUS_NO_CARBON";
    elseif p50<=cfg.fluxTol
        status="NO_PRODUCT_AT_50pct_mu";
    else
        status="OK";
    end

    rows(i,:)={ ...
        P.class,P.label,upper(string(mode)),P.doi,P.metID,P.outputRxn, ...
        P.carbonEquivalent,P.carbonEquivalentNote,P.unitNote, ...
        muMax,growthFloor,achieved, ...
        pMax,totalC0,ypc0,pC0,pcc0, ...
        p50,retained,pNC,status, ...
        glc,co2,totalC,ypc50,pC50,pcc50,ypg50,photon,o2};
end

T=cell2table(rows,'VariableNames',{ ...
    'Class','Product','Mode','DOI','Target_metabolite','Native_output_reaction', ...
    'Product_C_equivalent','Carbon_equivalent_note','Model_unit_note', ...
    'Mu_max_h','Growth_floor_h','Achieved_growth_h', ...
    'Max_product_no_growth_floor','P0_Total_external_C_mmol', ...
    'P0_Yield_product_mmol_per_C_mmol','P0_Product_C_flux_mmolC', ...
    'P0_Carbon_yield_Cmol_per_Cmol', ...
    'Product_at_50pct_mu_max','Fraction_of_unconstrained_product_retained', ...
    'NoCarbon_product_flux','Status','Glucose_uptake','CO2_uptake', ...
    'Total_external_C_mmol','Yield_product_mmol_per_C_mmol', ...
    'Product_C_flux_mmolC','Carbon_yield_Cmol_per_Cmol', ...
    'Yield_product_mmol_per_mmol_glucose','Photon_uptake','O2_uptake'});
end

function writeConsolidatedStateSheet_hvp(file,sheetName,mode,Tsubstrate,Tfixed,Tfull,cfg)
% Consolidate all reader-facing results for one organic-carbon trophic state
% into a single worksheet. Underlying calculations remain unchanged.
mode=upper(string(mode));
if mode~="HETERO" && mode~="MIXO"
    error('Consolidated state sheet supports HETERO or MIXO only.');
end

% Reader-facing Analysis A rule: rows with exactly zero product capacity
% are omitted. NaN rows (e.g. biologically not applicable reserve self-product
% cases) are retained as N/A rather than silently discarded.
subCapacity=Tsubstrate.Product_at_50pct_mu_max;
keepSubstrateRows=~isfinite(subCapacity) | abs(subCapacity)>cfg.fluxTol;
Tsubstrate=Tsubstrate(keepSubstrateRows,:);

fullCapacity=Tfull.Product_at_50pct_mu_max;
keepFullRows=~isfinite(fullCapacity) | abs(fullCapacity)>cfg.fluxTol;
Tfull=Tfull(keepFullRows,:);

writecell({ ...
    sprintf('%s HVP ANALYSIS',char(mode)); ...
    'This sheet combines the substrate-capacity screen, fixed-growth carbon-yield matrix, and D-glucose full-state HVP FBA for this trophic condition.' ...
    },file,'Sheet',sheetName,'Range','A1');

% A. Compact substrate-by-product capacity screen.
writecell({'A. SUBSTRATE-BY-PRODUCT PRODUCTION CAPACITY'}, ...
    file,'Sheet',sheetName,'Range','A4');
writecell({'Each HVP is maximized while growth is maintained at >=50% of the substrate-specific maximum growth rate.'}, ...
    file,'Sheet',sheetName,'Range','A5');

if mode=="HETERO"
    Tout=Tsubstrate(:,{ ...
        'Substrate','Substrate_class','Carbon_atoms','Mu_max_h','Growth_floor_h', ...
        'Product_class','Product','DOI','Product_at_50pct_mu_max', ...
        'Organic_source_C_mmolC'});
    headers={ ...
        'Substrate','Substrate class','Carbon atoms','Maximum growth (h^-1)', ...
        '50% growth floor (h^-1)','Product class','HVP','DOI', ...
        'Product flux at >=50% max growth','Organic-C input (C-mmol gDW^-1 h^-1)'};
else
    Tout=Tsubstrate(:,{ ...
        'Substrate','Substrate_class','Carbon_atoms','Mu_max_h','Growth_floor_h', ...
        'Product_class','Product','DOI','Product_at_50pct_mu_max', ...
        'Organic_source_C_mmolC','CO2_uptake_mmolC','Total_C_input_mmolC'});
    headers={ ...
        'Substrate','Substrate class','Carbon atoms','Maximum growth (h^-1)', ...
        '50% growth floor (h^-1)','Product class','HVP','DOI', ...
        'Product flux at >=50% max growth','Organic-C input (C-mmol gDW^-1 h^-1)', ...
        'CO2-C input (C-mmol gDW^-1 h^-1)','Total C input (C-mmol gDW^-1 h^-1)'};
end
Tout=roundNumericTable_hvp(Tout,3);
writecell(headers,file,'Sheet',sheetName,'Range','A7');
writetable(Tout,file,'Sheet',sheetName,'Range','A8','WriteVariableNames',false);

% B. Fixed-growth carbon-yield matrix.
rowFixed=8+height(Tout)+3;
writecell({'B. FIXED-GROWTH HVP CARBON YIELDS'}, ...
    file,'Sheet',sheetName,'Range',sprintf('A%d',rowFixed));
if mode=="HETERO"
    writecell({ ...
        sprintf('Fixed growth: mu = %.3f h^-1',cfg.fixedMu_h); ...
        'Values are C-mol product-C per C-mol organic carbon supplied; values are rounded to three decimals.'; ...
        'State-specific matrix: glycogen, TAG, floridoside and ergosterol.' ...
        },file,'Sheet',sheetName,'Range',sprintf('A%d',rowFixed+1));
    fixedHeaders={'Substrate','Substrate class','Glycogen','TAG', ...
        'Floridoside','Ergosterol'};
    headerRow=rowFixed+5;
else
    writecell({ ...
        sprintf('Fixed growth: mu = %.3f h^-1',cfg.fixedMu_h); ...
        'Two MIXO yield definitions are exported: apparent yield vs organic C only, and mass-balance yield vs total assimilated C (organic C + CO2-C).'; ...
        'Carbon-uptake columns are medians across the five MIXO HVP optimizations; each HVP yield uses that HVP solution''s own carbon uptake.'; ...
        'State-specific matrix: glycogen, TAG, C18:2, PCB and ergosterol.' ...
        },file,'Sheet',sheetName,'Range',sprintf('A%d',rowFixed+1));
    fixedHeaders={'Substrate','Substrate class', ...
        'Median organic-C uptake (C-mmol gDW^-1 h^-1)', ...
        'Median CO2-C uptake (C-mmol gDW^-1 h^-1)', ...
        'Median total-C uptake (C-mmol gDW^-1 h^-1)', ...
        'Median CO2 contribution to total C (%)', ...
        'Glycogen yield vs organic-C','Glycogen yield vs total-C', ...
        'TAG yield vs organic-C','TAG yield vs total-C', ...
        'C18:2 yield vs organic-C','C18:2 yield vs total-C', ...
        'PCB yield vs organic-C','PCB yield vs total-C', ...
        'Ergosterol yield vs organic-C','Ergosterol yield vs total-C'};
    headerRow=rowFixed+6;
end
Tfixed=roundNumericTable_hvp(Tfixed,3);
writecell(fixedHeaders,file,'Sheet',sheetName,'Range',sprintf('A%d',headerRow));
writetable(Tfixed,file,'Sheet',sheetName,'Range',sprintf('A%d',headerRow+1), ...
    'WriteVariableNames',false);

% C. Compact D-glucose full-state FBA block retained from the previous
% state-specific FBA sheets.
rowFull=headerRow+1+height(Tfixed)+3;
writecell({'C. D-GLUCOSE FULL-STATE HVP FBA'}, ...
    file,'Sheet',sheetName,'Range',sprintf('A%d',rowFull));
writecell({'Selected HVPs under the validated trophic-state constraints.'}, ...
    file,'Sheet',sheetName,'Range',sprintf('A%d',rowFull+1));
TfullOut=compactFullStateForExport_hvp(Tfull);
TfullOut=roundNumericTable_hvp(TfullOut,3);
fullHeaders={'Product class','HVP','DOI','Maximum growth (h^-1)', ...
    '50% growth floor (h^-1)','Product flux at 50% max growth', ...
    'Glucose uptake','CO2 uptake','Total external C (C-mmol gDW^-1 h^-1)'};
writecell(fullHeaders,file,'Sheet',sheetName,'Range',sprintf('A%d',rowFull+3));
writetable(TfullOut,file,'Sheet',sheetName,'Range',sprintf('A%d',rowFull+4), ...
    'WriteVariableNames',false);
end

function Tout=compactFullStateForExport_hvp(T)
Tout=T(:,{ ...
    'Class','Product','DOI','Mu_max_h','Growth_floor_h', ...
    'Product_at_50pct_mu_max','Glucose_uptake','CO2_uptake', ...
    'Total_external_C_mmol'});
end

function writeAutoStateFBASheet_hvp(file,sheetName,T,titleText)
% AUTO has no glucose input. Keep only reader-facing AUTO variables and
% omit Status, Glucose uptake and Total external carbon from the worksheet.
writecell({ ...
    titleText; ...
    'Compact literature-guided autotrophic HVP FBA results.'; ...
    'Product flux is reported at >=50% of the condition-specific maximum growth rate.' ...
    },file,'Sheet',sheetName,'Range','A1');

Tout=T(:,{ ...
    'Class','Product','DOI','Mu_max_h','Growth_floor_h', ...
    'Product_at_50pct_mu_max','CO2_uptake'});
Tout=roundNumericTable_hvp(Tout,3);
headers={'Product class','HVP','DOI','Maximum growth (h^-1)', ...
    '50% growth floor (h^-1)','Product flux at 50% max growth','CO2 uptake'};
writecell(headers,file,'Sheet',sheetName,'Range','A5');
writetable(Tout,file,'Sheet',sheetName,'Range','A6','WriteVariableNames',false);
end

function idx=findProductKey_hvp(products,key)
idx=find(strcmpi(string({products.key}),string(key)),1);
if isempty(idx)
    error('Selected HVP key not found: %s',string(key));
end
end

function [glc,co2,totalC,photon,o2]=fullStateUptakeMetrics_hvp(m,mode,sol,cfg)
glc=NaN; co2=NaN; totalC=NaN; photon=NaN; o2=NaN;
if ~isFeasibleSolution_hvp(sol), return; end
mode=upper(string(mode));
if mode=="MIXO" || mode=="HETERO"
    glc=uptakeMagnitudeFromSolution_hvp(m,cfg.glucoseRxn,sol.x);
else
    glc=0;
end
if mode=="AUTO" || mode=="MIXO"
    co2=uptakeMagnitudeFromSolution_hvp(m,cfg.co2Rxn,sol.x);
else
    co2=0;
end
totalC=cfg.glucoseCarbonAtoms*glc+co2;
photon=uptakeMagnitudeFromSolution_hvp(m,cfg.photonRxn,sol.x);
o2=uptakeMagnitudeFromSolution_hvp(m,cfg.o2Rxn,sol.x);
end

function [yieldPC,yieldCC,productCFlux,yieldPerGlucose]= ...
    fullStateYields_hvp(productFlux,productCeq,totalCarbon,glucoseUptake,tol)
yieldPC=NaN; yieldCC=NaN; productCFlux=NaN; yieldPerGlucose=NaN;
if ~isfinite(productFlux) || ~isfinite(productCeq), return; end
productCFlux=productFlux*productCeq;
if isfinite(totalCarbon) && totalCarbon>tol
    yieldPC=productFlux/totalCarbon;
    yieldCC=productCFlux/totalCarbon;
end
if isfinite(glucoseUptake) && glucoseUptake>tol
    yieldPerGlucose=productFlux/glucoseUptake;
end
end

function m=closeAllCarbonUptakeFull_hvp(m,cfg)
for i=1:numel(cfg.carbonBoundaryRxns)
    r=cfg.carbonBoundaryRxns{i};
    if any(strcmp(m.rxns,r)), m=closeUptake_hvp(m,r); end
end
for i=1:numel(cfg.allMappedCarbonRxns)
    r=cfg.allMappedCarbonRxns{i};
    if any(strcmp(m.rxns,r)), m=closeUptake_hvp(m,r); end
end
if any(strcmp(m.rxns,cfg.co2Rxn)), m=closeUptake_hvp(m,cfg.co2Rxn); end
end

function [m,cfg,auditT]=implementAutoWorkbookBOF_hvp(m,cfg,biomassFile)
mets=cfg.resolvedBOFMets;
pig=cfg.resolvedPigmentMets;

bofFormula=readWorkbookReactionFormula_hvp( ...
    biomassFile,'Reaction List Auto','TOTAL_BIOMASS_AUTO_MEAN');
pigFormula=readWorkbookReactionFormula_hvp( ...
    biomassFile,'Reaction List Auto','PIGMENT_TOTAL_POOL_BIO_AUTOTROPH_NOFRAC');

cProtein=coefficientForWorkbookMet_hvp(bofFormula,'m_PROTEIN_TOTAL_POOL[c]');
cCarb=coefficientForWorkbookMet_hvp(bofFormula,'m_CARBOHYDRATE_TOTAL_POOL[c]');
cLipid=coefficientForWorkbookMet_hvp(bofFormula,'m_LIPID_TOTAL_POOL[c]');
cRNA=coefficientForWorkbookMet_hvp(bofFormula,'m_RNA_TOTAL_POOL[c]');
cDNA=coefficientForWorkbookMet_hvp(bofFormula,'m_DNA_TOTAL_POOL[c]');
cPig=coefficientForWorkbookMet_hvp(bofFormula,'m_PIGMENT_TOTAL_POOL[c]');

pZea=coefficientForWorkbookMet_hvp(pigFormula,'m_CPD1F-130_Thyl[t]');
try
    pChl=coefficientForWorkbookMet_hvp(pigFormula,'m_CHLOROPHYLL-A_Plas[p]');
catch
    pChl=coefficientForWorkbookMet_hvp(pigFormula,'m_CHLOROPHYLL-A_Thyl[t]');
end
pBeta=coefficientForWorkbookMet_hvp(pigFormula,'m_CPD1F-129_Thyl[t]');
pPCB=coefficientForWorkbookMet_hvp(pigFormula,'m_3Z-PHYCOCYANOBILIN_Plas[p]');
pc=[pZea;pChl;pBeta;pPCB];
pc=pc/sum(pc);

pigMet='m_PUBHVP_PIGMENT_TOTAL_AUTO[c]';
pigRxn='PUBHVP_PIGMENT_TOTAL_AUTO_PRODUCER_bm';
bioMet='m_PUBHVP_TOTAL_BIOMASS_AUTO[c]';
assembly='PUBHVP_TOTAL_BIOMASS_AUTO_bm';
sink='PUBHVP_TOTAL_BIOMASS_AUTO_SINK_bm_tx';

m=removeIfPresent_hvp(m,{pigRxn,assembly,sink});
m=addMetIfMissing_hvp(m,pigMet,'publication HVP AUTO pigment pool');
m=addMetIfMissing_hvp(m,bioMet,'publication HVP AUTO total biomass');

m=addReaction(m,pigRxn, ...
    'reactionName','publication HVP AUTO pigment pool', ...
    'metaboliteList',{pig.zea;pig.chla;pig.beta;pig.pcb;pigMet}, ...
    'stoichCoeffList',[-pc;1], ...
    'lowerBound',0,'upperBound',1000,'subSystem','Biomass pigment pool');
m=padConstraintColumns_hvp(m);

m=addReaction(m,assembly, ...
    'reactionName','publication HVP AUTO mean biomass objective', ...
    'metaboliteList',{mets.protein;mets.carb;mets.lipid;mets.rna;mets.dna; ...
        pigMet;mets.atp;mets.water;mets.adp;mets.pi;mets.proton;bioMet}, ...
    'stoichCoeffList',[-cProtein;-cCarb;-cLipid;-cRNA;-cDNA;-cPig; ...
        -cfg.GAM;-cfg.GAM;cfg.GAM;cfg.GAM;cfg.GAM;1], ...
    'lowerBound',0,'upperBound',1000,'subSystem','Biomass synthesis');
m=padConstraintColumns_hvp(m);

m=addReaction(m,sink, ...
    'reactionName','publication HVP AUTO biomass sink', ...
    'metaboliteList',{bioMet},'stoichCoeffList',-1, ...
    'lowerBound',0,'upperBound',1000,'subSystem','Biomass sink');
m=padConstraintColumns_hvp(m);

b=struct();
b.mode="AUTO"; b.assemblyRxn=assembly; b.sinkRxn=sink; b.biomassMet=bioMet;
b.pigmentRxn=pigRxn; b.pigmentMet=pigMet;
b.proteinCoeff=cProtein; b.carbohydrateCoeff=cCarb; b.lipidCoeff=cLipid;
b.rnaCoeff=cRNA; b.dnaCoeff=cDNA; b.pigmentCoeff=cPig;
b.GAM=cfg.GAM; b.pigmentComposition=pc;
b.source="TOTAL_BIOMASS.xlsx / TOTAL_BIOMASS_AUTO_MEAN";
cfg.AUTO=b;

Mode=repmat("AUTO",6,1);
Component=["Protein";"Carbohydrate";"Lipid";"RNA";"DNA";"Pigment"];
Raw_g_gDW=nan(6,1);
Coefficient=[cProtein;cCarb;cLipid;cRNA;cDNA;cPig];
Source=repmat("TOTAL_BIOMASS.xlsx / TOTAL_BIOMASS_AUTO_MEAN",6,1);
auditT=table(Mode,Component,Raw_g_gDW,Coefficient,Source);
end

function formula=readWorkbookReactionFormula_hvp(file,sheet,abbr)
C=readcell(file,'Sheet',sheet);
formula="";
for i=1:size(C,1)
    for j=1:size(C,2)
        v=stringCell_hvp(C{i,j});
        if strcmpi(strtrim(v),string(abbr))
            for k=1:size(C,2)
                s=stringCell_hvp(C{i,k});
                if contains(s,"->") || contains(s,"<=>") || contains(s,"<->")
                    formula=s;
                    break;
                end
            end
            if strlength(formula)>0, return; end
        end
    end
end
error('Reaction formula not found in %s / %s for %s',file,sheet,abbr);
end

function s=stringCell_hvp(v)
if isempty(v)
    s="";
elseif isstring(v)
    if all(ismissing(v(:))), s=""; else, s=string(v); end
elseif ischar(v)
    s=string(v);
elseif isnumeric(v) && isscalar(v)
    if isnan(v), s=""; else, s=string(v); end
elseif islogical(v) && isscalar(v)
    s=string(v);
else
    try
        tmp=string(v);
        if all(ismissing(tmp(:))), s=""; else, s=tmp; end
    catch
        s="";
    end
end
end

function c=coefficientForWorkbookMet_hvp(formula,target)
f=char(string(formula));
f=regexprep(f,'[\r\n\t]+',' ');
f=regexprep(f,'\s+',' ');
f=strtrim(f);
t=regexptranslate('escape',char(string(target)));
num='([0-9]*\.?[0-9]+(?:[eE][+\-]?[0-9]+)?)?';
pat=['(?:^|\+|->)\s*' num '\s*' t '\s*(?:\+|->|$)'];
tok=regexp(f,pat,'tokens','once');
if isempty(tok)
    error('Metabolite %s was not found in workbook reaction formula.',target);
end
if isempty(tok{1})
    c=1;
else
    c=str2double(tok{1});
end
if ~isfinite(c)
    error('Invalid coefficient for %s in workbook reaction formula.',target);
end
end

%% STORAGE-RESERVE / POST-ANALYSIS HELPERS

function [m,cfg] = addStorageReserveModule_hvp(m,cfg,resolvedExchange,isReserveSubstrate)
cfg.reserveSourceRxn="";
cfg.reserveMobilizationRxn="";
cfg.reserveOriginalExchange="";
if ~any(isReserveSubstrate), return; end

reserveRows=find(isReserveSubstrate);
reserveBoundary=char(resolvedExchange(reserveRows(1)));
for ii=2:numel(reserveRows)
    if ~strcmp(char(resolvedExchange(reserveRows(ii))),reserveBoundary)
        error('Multiple different storage alpha-glucan exchange reactions were mapped.');
    end
end

starchMet=singleBoundaryMet_hvp(m,reserveBoundary);
glucoseMet=singleBoundaryMet_hvp(m,cfg.glucoseRxn);
waterMet=singleBoundaryMet_hvp(m,cfg.basalRxns.H2O);

sourceRxn='HVP_STARCH_RESERVE_SOURCE_Cyto';
mobilRxn='HVP_STARCH_RESERVE_MOBILIZATION_Cyto';
if any(strcmp(m.rxns,sourceRxn)), m=removeRxns(m,{sourceRxn}); end
if any(strcmp(m.rxns,mobilRxn)), m=removeRxns(m,{mobilRxn}); end

% Source reaction represents withdrawal from a pre-existing intracellular
% storage pool. Positive flux supplies one glucosyl-equivalent STARCH unit.
m=addReaction(m,sourceRxn, ...
    'reactionName','analysis-only intracellular storage alpha-glucan reserve source', ...
    'metaboliteList',{starchMet},'stoichCoeffList',1, ...
    'lowerBound',0,'upperBound',0, ...
    'subSystem','Analysis-only intracellular reserve mobilization');
m=padConstraintColumns_hvp(m);

% Lumped hydrolysis proxy for reserve use: (C6H10O5)n unit + H2O -> glucose.
% This is used only in the analysis model and does not alter the SBML input.
m=addReaction(m,mobilRxn, ...
    'reactionName','analysis-only storage alpha-glucan hydrolysis to glucose equivalent', ...
    'metaboliteList',{starchMet,waterMet,glucoseMet}, ...
    'stoichCoeffList',[-1;-1;1], ...
    'lowerBound',0,'upperBound',0, ...
    'subSystem','Analysis-only intracellular reserve mobilization');
m=padConstraintColumns_hvp(m);

cfg.reserveSourceRxn=sourceRxn;
cfg.reserveMobilizationRxn=mobilRxn;
cfg.reserveOriginalExchange=reserveBoundary;
end

function met=singleBoundaryMet_hvp(m,rxn)
j=find(strcmp(m.rxns,rxn),1);
if isempty(j), error('Boundary reaction not found: %s',rxn); end
nz=find(abs(full(m.S(:,j)))>1e-12);
if numel(nz)~=1
    error('Expected one-metabolite boundary reaction for %s; found %d metabolites.',rxn,numel(nz));
end
met=char(m.mets{nz});
end

function Trel = buildReferenceRelativeMatrix_hvp(T,products,referenceSubstrate)
% Percentage change in product flux relative to one reference substrate.
% The reference is evaluated separately within each trophic condition.
Trel=T;
refRow=find(strcmpi(string(T.Substrate),string(referenceSubstrate)),1);
if isempty(refRow)
    error('Heatmap reference substrate was not found: %s',referenceSubstrate);
end
for p=1:numel(products)
    varName=matlab.lang.makeValidName(char(products(p).key));
    vals=T.(varName);
    refVal=vals(refRow);
    if ~isfinite(refVal) || abs(refVal)<1e-12
        error('Reference substrate %s has no valid flux for %s.', ...
            referenceSubstrate,products(p).label);
    end
    Trel.(varName)=100*(vals-refVal)./refVal;
end
end

function limPct = relativeHeatmapLimit_hvp(Trel,products)
% Use a separate symmetric scale for each trophic state. This preserves the
% true percentage values while making the smaller MIXO substrate effects
% visible instead of compressing them on the wider HETERO scale.
vals=[];
for p=1:numel(products)
    varName=matlab.lang.makeValidName(char(products(p).key));
    vals=[vals; Trel.(varName)(:)]; %#ok<AGROW>
end
vals=vals(isfinite(vals));
if isempty(vals)
    limPct=10;
    return;
end
limPct=max(10,5*ceil(max(abs(vals))/5));
end

function createReferenceRelativeHeatmap_hvp(Trel,products,mode,outFile, ...
    growthFraction,limPct,annotationThresholdPct,referenceSubstrate)
% Publication-style diverging heatmap of substrate effect relative to D-glucose.
% Heatmap columns are enforced here from the trophic-state-specific biological
% interpretation sets so a stale caller cannot accidentally plot all nine HVPs.
mode=upper(string(mode));
if mode=="HETERO"
    selectedKeys=["Glycogen","TAG","Floridoside","Ergosterol"];
    expectedCount=4;
elseif mode=="MIXO"
    selectedKeys=["Glycogen","TAG","C18_2","PCB","Ergosterol"];
    expectedCount=5;
else
    error('Heatmap mode must be HETERO or MIXO.');
end
products=selectProducts_hvp(products,selectedKeys);
if numel(products)~=expectedCount
    error('%s heatmap product selection failed: expected %d products, found %d.', ...
        char(mode),expectedCount,numel(products));
end

productVars=cell(numel(products),1);
for p=1:numel(products)
    productVars{p}=matlab.lang.makeValidName(char(products(p).key));
    if ~ismember(productVars{p},Trel.Properties.VariableNames)
        error('%s heatmap is missing product column %s.',char(mode),productVars{p});
    end
end
% Explicitly keep only substrate metadata plus the selected product columns.
% This makes it impossible for non-selected HVPs to reach the plotting matrix.
keepVars=[{'Substrate','Class'}, productVars'];
Ts=Trel(:,keepVars);
Ts=sortrows(Ts,{'Class','Substrate'});

data=nan(height(Ts),numel(products));
for p=1:numel(products)
    data(:,p)=Ts.(productVars{p});
end

% Reader-facing substrate labels: classes are omitted from every row to
% reduce crowding; rows remain grouped by substrate class through sorting.
rowLabels=string(Ts.Substrate);
rowLabels(contains(lower(rowLabels),'starch'))="Storage alpha-glucan reserve";
rowLabels=replace(rowLabels,"alpha-D-Ribopyranose",string(char(945))+"-D-Ribopyranose");
rowLabels=replace(rowLabels,"alpha-D-Xylopyranose",string(char(945))+"-D-Xylopyranose");

prettyProducts=strings(1,numel(products));
for p=1:numel(products)
    key=string(products(p).key);
    switch key
        case "Glycogen",     prettyProducts(p)="Glycogen";
        case "TAG",          prettyProducts(p)="TAG";
        case "C18_2",        prettyProducts(p)="C18:2";
        case "BetaCarotene", prettyProducts(p)=string(char(946))+"-Carotene";
        case "PCB",          prettyProducts(p)="PCB";
        case "Floridoside",  prettyProducts(p)="Floridoside";
        case "Squalene",     prettyProducts(p)="Squalene";
        case "Ergosterol",   prettyProducts(p)="Ergosterol";
        case "Phytol",       prettyProducts(p)="Phytol";
        otherwise,            prettyProducts(p)=string(products(p).label);
    end
end

% Approximately full journal-column width when exported; height is scaled
% to the number of substrates so y-axis labels remain readable.
figHeight=max(15.5,0.47*height(Ts)+3.5);
f=figure('Visible','off','Color','w','Units','centimeters', ...
    'Position',[2 2 18.3 figHeight]);
ax=axes(f);
h=imagesc(ax,data,[-limPct limPct]);
set(h,'AlphaData',~isnan(data));
set(ax,'Color',[0.92 0.92 0.92]);
colormap(ax,orangeWhiteBlue_hvp(256));
cb=colorbar(ax);
cb.Label.String='Change vs D-glucose (%)';
cb.FontName='Arial';
cb.FontSize=8;
cb.Label.FontName='Arial';
cb.Label.FontSize=8;

ax.XTick=1:numel(products);
ax.XTickLabel=cellstr(prettyProducts);
ax.YTick=1:height(Ts);
ax.YTickLabel=cellstr(rowLabels);
ax.TickLabelInterpreter='none';
ax.FontName='Arial';
ax.FontSize=8;
ax.LineWidth=0.75;
ax.TickDir='out';
ax.Box='off';
xtickangle(ax,45);
xlabel(ax,'High-value products','FontName','Arial','FontSize',9);
ylabel(ax,'Carbon source','FontName','Arial','FontSize',9);
title(ax,char(mode),'FontName','Arial','FontSize',10,'FontWeight','bold', ...
    'Interpreter','none');

% Thin separators between substrate classes retain visual grouping without
% repeating class names in every y-axis label.
classNames=string(Ts.Class);
for r=1:height(Ts)-1
    if classNames(r)~=classNames(r+1)
        yline(ax,r+0.5,'-','Color',[0.72 0.72 0.72],'LineWidth',0.45);
    end
end

% N/A cells are explicit, e.g. glycogen when the storage alpha-glucan pool
% itself is serving as the intracellular reserve source.
[nr,nc]=find(isnan(data));
for k=1:numel(nr)
    text(ax,nc(k),nr(k),'N/A','HorizontalAlignment','center', ...
        'VerticalAlignment','middle','FontName','Arial','FontSize',7, ...
        'Color',[0.25 0.25 0.25]);
end

% Label effects >= threshold. Also label the strongest positive substrate
% for each product even when MIXO substrate sensitivity is small.
for p=1:size(data,2)
    col=data(:,p);
    finiteRows=find(isfinite(col));
    bestRow=[];
    if ~isempty(finiteRows)
        [~,kk]=max(col(finiteRows));
        bestRow=finiteRows(kk);
    end
    for r=1:size(data,1)
        if ~isfinite(data(r,p)), continue; end
        showLabel=abs(data(r,p))>=annotationThresholdPct || ...
            (~isempty(bestRow) && r==bestRow && data(r,p)>0);
        if showLabel
            fw='normal';
            if ~isempty(bestRow) && r==bestRow, fw='bold'; end
            text(ax,p,r,sprintf('%+.0f',data(r,p)), ...
                'HorizontalAlignment','center','VerticalAlignment','middle', ...
                'FontName','Arial','FontSize',7,'FontWeight',fw, ...
                'Color',[0.10 0.10 0.10]);
        end
    end
end

% Remove any previous image first so the output cannot be confused with a
% stale nine-product heatmap from an earlier script run.
if isfile(outFile)
    delete(outFile);
end
try
    exportgraphics(f,outFile,'Resolution',600,'BackgroundColor','white');
catch
    print(f,outFile,'-dpng','-r600');
end
close(f);
end

function createFixedGrowthYieldBarChart_hvp(Tfixed,products,mode,outFile,referenceSubstrate,fixedMu_h)
% Grouped fixed-growth carbon-yield chart using exactly the trophic-state
% products present in the fixed-mu matrix. Product bars use a stable MATLAB
% palette; only substrate x-axis labels are colored by median response class.
% Classification rules are intentionally not printed in the figure itself.

mode=upper(string(mode));
if mode~="HETERO" && mode~="MIXO"
    error('Fixed-growth yield chart mode must be HETERO or MIXO.');
end

nS=height(Tfixed);
nP=numel(products);
if nS<1 || nP<1
    error('%s fixed-growth yield chart has no data.',char(mode));
end

Y=nan(nS,nP);
productVars=cell(nP,1);
prettyProducts=strings(1,nP);
for p=1:nP
    baseVar=matlab.lang.makeValidName(char(products(p).key));
    if mode=="MIXO"
        productVars{p}=matlab.lang.makeValidName([baseVar '_TotalC']);
    else
        productVars{p}=baseVar;
    end
    if ~ismember(productVars{p},Tfixed.Properties.VariableNames)
        error('%s fixed-growth table is missing product column %s.', ...
            char(mode),productVars{p});
    end
    Y(:,p)=Tfixed.(productVars{p});
    switch string(products(p).key)
        case "Glycogen",     prettyProducts(p)="Glycogen";
        case "TAG",          prettyProducts(p)="TAG";
        case "C18_2",        prettyProducts(p)="C18:2";
        case "BetaCarotene", prettyProducts(p)=string(char(946))+"-Carotene";
        case "PCB",          prettyProducts(p)="PCB";
        case "Floridoside",  prettyProducts(p)="Floridoside";
        case "Squalene",     prettyProducts(p)="Squalene";
        case "Ergosterol",   prettyProducts(p)="Ergosterol";
        case "Phytol",       prettyProducts(p)="Phytol";
        otherwise,            prettyProducts(p)=string(products(p).label);
    end
end

refRow=find(strcmpi(string(Tfixed.Substrate),string(referenceSubstrate)),1);
if isempty(refRow)
    error('Fixed-growth chart reference substrate was not found: %s',referenceSubstrate);
end
refY=Y(refRow,:);
medianChange=nan(nS,1);
for s=1:nS
    pct=nan(1,nP);
    for p=1:nP
        if isfinite(Y(s,p)) && isfinite(refY(p)) && abs(refY(p))>1e-12
            pct(p)=100*(Y(s,p)-refY(p))/refY(p);
        end
    end
    medianChange(s)=medianFinite_hvp(pct);
end

substrateLabels=string(Tfixed.Substrate);
substrateLabels=replace(substrateLabels,"Dulcitol (Galactitol)","Dulcitol");
substrateLabels=replace(substrateLabels,"Ribitol (Adonitol)","Ribitol");
substrateLabels=replace(substrateLabels,"alpha-D-Ribopyranose",string(char(945))+"-D-Ribopyranose");
substrateLabels=replace(substrateLabels,"alpha-D-Xylopyranose",string(char(945))+"-D-Xylopyranose");
reserveMask=contains(lower(substrateLabels),"starch") | ...
    contains(lower(substrateLabels),"alpha-glucan");
substrateLabels(reserveMask)="Storage "+string(char(945))+"-glucan reserve";

% Response class affects only substrate-label color. The class legend/rules
% are intentionally left to the manuscript caption/text below the figure.
cHigh=[0.00 0.45 0.20];
cRef=[0.10 0.35 0.75];
cLow=[0.90 0.45 0.05];
cStrong=[0.75 0.10 0.10];
cReserve=[0.35 0.35 0.35];
labelColors=repmat(cRef,nS,1);
for s=1:nS
    if reserveMask(s)
        labelColors(s,:)=cReserve;
        continue;
    end
    d=medianChange(s);
    if ~isfinite(d)
        labelColors(s,:)=cReserve;
        continue;
    end
    if mode=="HETERO"
        if d>=5
            labelColors(s,:)=cHigh;
        elseif d>=-5
            labelColors(s,:)=cRef;
        elseif d>=-20
            labelColors(s,:)=cLow;
        else
            labelColors(s,:)=cStrong;
        end
    else
        if d>0.5
            labelColors(s,:)=cHigh;
        elseif d>=-0.5
            labelColors(s,:)=cRef;
        elseif d>=-1.0
            labelColors(s,:)=cLow;
        else
            labelColors(s,:)=cStrong;
        end
    end
end

% Scientific Reports/Nature-style publication layout. The figure is built at
% approximately double-column final width (18.3 cm) so font sizes are already
% close to their final printed size. Arial is used consistently, the background
% is white, axis strokes are >=1 pt, and PNG export remains at 300 dpi. Rotated
% substrate names occupy a compact lower band; the centered product legend is
% placed immediately below the substrate names without overlap.
f=figure('Visible','off','Color','w','Units','centimeters', ...
    'Position',[2 2 18.3 11.5]);
ax=axes(f,'Position',[0.095 0.325 0.885 0.61]);
hold(ax,'on');
b=bar(ax,1:nS,Y,'grouped','BarWidth',0.86);

% Fixed publication palette. HETERO therefore appears as blue / orange /
% grey / purple; TAG (orange) and ergosterol (purple) are deliberately very
% different. The same HVP keeps the same color in the MIXO figure.
canonicalKeys=["Glycogen","TAG","C18_2","BetaCarotene","PCB", ...
    "Floridoside","Squalene","Ergosterol","Phytol"];
productColors=[ ...
    0.20 0.45 0.70; ... % Glycogen - blue
    0.90 0.35 0.05; ... % TAG - orange
    0.00 0.60 0.50; ... % C18:2 - teal/green
    0.95 0.75 0.15; ... % beta-carotene - yellow
    0.80 0.47 0.65; ... % PCB - magenta
    0.55 0.55 0.55; ... % Floridoside - grey
    0.35 0.70 0.90; ... % Squalene - sky blue
    0.45 0.20 0.65; ... % Ergosterol - purple
    0.60 0.45 0.25];    % Phytol - brown
for p=1:nP
    ci=find(canonicalKeys==string(products(p).key),1);
    if isempty(ci), ci=min(p,size(productColors,1)); end
    b(p).FaceColor=productColors(ci,:);
    b(p).EdgeColor='none';
end

xlim(ax,[0.35 nS+0.65]);
ax.XTick=1:nS;
ax.XTickLabel=repmat({''},1,nS);
ax.FontName='Arial';
ax.FontSize=8.5;
ax.LineWidth=1.0;
ax.TickDir='out';
ax.Box='off';
ax.YGrid='on';
ax.XGrid='off';
ax.GridAlpha=0.14;
if mode=="HETERO"
    ylabel(ax,'Carbon yield (C-mol product-C / C-mol organic-C)', ...
        'FontName','Arial','FontSize',9.5);
else
    ylabel(ax,'Carbon yield (C-mol product-C / C-mol total-C)', ...
        'FontName','Arial','FontSize',9.5);
end
title(ax,sprintf('%s fixed-growth HVP carbon yields (\\mu = %.3f h^{-1})', ...
    char(mode),fixedMu_h),'FontName','Arial','FontSize',10.5, ...
    'FontWeight','bold','Interpreter','tex');

% Product legend only. The substrate-response classification remains encoded
% by the x-axis label colors and is explained in the manuscript caption. The
% legend is centered immediately below the rotated substrate labels. A separate
% centered carbon-source label is used instead of the MATLAB legend title so its
% alignment is stable for both the four-product HETERO and five-product MIXO
% figures.
lgd=legend(ax,b,cellstr(prettyProducts), ...
    'Orientation','horizontal','Box','off','FontName','Arial','FontSize',8.5);
try
    lgd.NumColumns=nP;
catch
end
try
    lgd.Units='normalized';
    legendWidth=min(0.68,max(0.42,0.115*nP));
    lgd.Position=[0.5-legendWidth/2 0.018 legendWidth 0.050];
    lgd.Title.String='';
catch
end
annotation(f,'textbox',[0.35 0.082 0.30 0.028], ...
    'String','Carbon source','HorizontalAlignment','center', ...
    'VerticalAlignment','middle','EdgeColor','none', ...
    'FontName','Arial','FontSize',9.0,'FontWeight','normal');

% MATLAB applies one color to all native tick labels, so individual colored
% substrate names are drawn as text objects below the axis.
yl=ylim(ax);
yr=yl(2)-yl(1);
if yr<=0, yr=1; end
yText=yl(1)-0.022*yr;
for s=1:nS
    text(ax,s,yText,char(substrateLabels(s)), ...
        'Rotation',65,'HorizontalAlignment','right', ...
        'VerticalAlignment','top','FontName','Arial','FontSize',8.0, ...
        'Color',labelColors(s,:),'Interpreter','none','Clipping','off');
end

% No separate x-axis title is used: the colored tick labels are substrate
% names, and the product legend/title is deliberately positioned below them.
xlabel(ax,'');

% Keep a zero baseline for direct yield comparison.
yl=ylim(ax);
if yl(1)>0
    ylim(ax,[0 yl(2)]);
end

if isfile(outFile)
    delete(outFile);
end
try
    exportgraphics(f,outFile,'Resolution',300,'BackgroundColor','white');
catch
    print(f,outFile,'-dpng','-r300');
end
close(f);
end

function cmap=orangeWhiteBlue_hvp(n)
% Diverging, non-red/green palette: negative = orange, zero = white,
% positive = blue. This is suitable for a zero-centered relative-effect map.
if nargin<1, n=256; end
neg=[0.85 0.33 0.10];
mid=[1.00 1.00 1.00];
pos=[0.15 0.42 0.68];
n1=floor(n/2);
n2=n-n1;
a=linspace(0,1,n1)';
b=linspace(0,1,n2)';
cmap1=neg.*(1-a)+mid.*a;
cmap2=mid.*(1-b)+pos.*b;
cmap=[cmap1;cmap2];
end

function T = buildBestSolutionSummary_hvp(HF,MF,HY,MY,HD,MD,products,referenceSubstrate)
% One compact row per HVP for the PRIMARY >=50% substrate-specific mu_max
% production-capacity screen. Carbon-yield fields remain internal legacy
% diagnostics and are not used for fixed-growth carbon-efficiency summaries.
n=numel(products);
Product_class=strings(n,1); Product=strings(n,1);
HETERO_reference_flux=nan(n,1); HETERO_best_flux_substrate=strings(n,1);
HETERO_best_flux=nan(n,1); HETERO_best_flux_change_vs_reference_pct=nan(n,1);
HETERO_best_yield_substrate=strings(n,1); HETERO_best_carbon_yield=nan(n,1);
MIXO_reference_flux=nan(n,1); MIXO_best_flux_substrate=strings(n,1);
MIXO_best_flux=nan(n,1); MIXO_best_flux_change_vs_reference_pct=nan(n,1);
MIXO_best_yield_substrate=strings(n,1); MIXO_best_carbon_yield=nan(n,1);
MIXO_vs_HETERO_best_flux_change_pct=nan(n,1);

refH=find(strcmpi(string(HF.Substrate),string(referenceSubstrate)),1);
refM=find(strcmpi(string(MF.Substrate),string(referenceSubstrate)),1);
if isempty(refH) || isempty(refM)
    error('Reference substrate %s is missing from summary matrices.',referenceSubstrate);
end

for p=1:n
    Product_class(p)=products(p).class;
    Product(p)=products(p).label;
    varName=matlab.lang.makeValidName(char(products(p).key));

    hv=HF.(varName); mv=MF.(varName);
    hy=HY.(varName); my=MY.(varName);
    hd=HD.(varName); md=MD.(varName);

    HETERO_reference_flux(p)=hv(refH);
    MIXO_reference_flux(p)=mv(refM);

    [hi,HETERO_best_flux(p)]=maxFinite_hvp(hv);
    [hyi,HETERO_best_carbon_yield(p)]=maxFinite_hvp(hy);
    [mi,MIXO_best_flux(p)]=maxFinite_hvp(mv);
    [myi,MIXO_best_carbon_yield(p)]=maxFinite_hvp(my);

    if ~isempty(hi)
        HETERO_best_flux_substrate(p)=string(HF.Substrate(hi));
        HETERO_best_flux_change_vs_reference_pct(p)=hd(hi);
    end
    if ~isempty(hyi), HETERO_best_yield_substrate(p)=string(HY.Substrate(hyi)); end
    if ~isempty(mi)
        MIXO_best_flux_substrate(p)=string(MF.Substrate(mi));
        MIXO_best_flux_change_vs_reference_pct(p)=md(mi);
    end
    if ~isempty(myi), MIXO_best_yield_substrate(p)=string(MY.Substrate(myi)); end

    if isfinite(HETERO_best_flux(p)) && HETERO_best_flux(p)>0 && isfinite(MIXO_best_flux(p))
        MIXO_vs_HETERO_best_flux_change_pct(p)= ...
            100*(MIXO_best_flux(p)/HETERO_best_flux(p)-1);
    end
end

T=table(Product_class,Product, ...
    HETERO_reference_flux,HETERO_best_flux_substrate,HETERO_best_flux, ...
    HETERO_best_flux_change_vs_reference_pct,HETERO_best_yield_substrate, ...
    HETERO_best_carbon_yield, ...
    MIXO_reference_flux,MIXO_best_flux_substrate,MIXO_best_flux, ...
    MIXO_best_flux_change_vs_reference_pct,MIXO_best_yield_substrate, ...
    MIXO_best_carbon_yield,MIXO_vs_HETERO_best_flux_change_pct);
end

function [idx,val]=maxFinite_hvp(v)
idx=[]; val=NaN;
valid=find(isfinite(v));
if isempty(valid), return; end
[valLocal,k]=max(v(valid));
idx=valid(k);
val=valLocal;
end

function T = buildSubstrateClassSummary_hvp(HF,MF,HY,MY,HD,MD, ...
    productsHETERO,productsMIXO,productsCommon,fluxTol,yieldTol)
% Collapse exact duplicate model exchanges (e.g. D-/L-glucose mapping to
% the same exchange) before summarizing substrate classes. Within-state
% summaries use the literature-guided product set for that trophic state;
% cross-state comparisons use only products selected in both states.
[~,ia]=unique(string(HF.Resolved_exchange),'stable');
HF=HF(ia,:); MF=MF(ia,:); HY=HY(ia,:); MY=MY(ia,:); HD=HD(ia,:); MD=MD(ia,:);
classes=unique(string(HF.Class),'stable');
n=numel(classes);
Substrate_class=classes;
Unique_model_sources=zeros(n,1);
HETERO_median_change_vs_glucose_pct=nan(n,1);
MIXO_median_change_vs_glucose_pct=nan(n,1);
Median_MIXO_vs_HETERO_flux_change_pct=nan(n,1);
HETERO_median_carbon_yield=nan(n,1);
MIXO_median_carbon_yield=nan(n,1);
Median_MIXO_vs_HETERO_carbon_yield_change_pct=nan(n,1);

for c=1:n
    rows=strcmp(string(HF.Class),classes(c));
    Unique_model_sources(c)=sum(rows);

    hYield=productMatrix_hvp(HY(rows,:),productsHETERO);
    mYield=productMatrix_hvp(MY(rows,:),productsMIXO);
    hDelta=productMatrix_hvp(HD(rows,:),productsHETERO);
    mDelta=productMatrix_hvp(MD(rows,:),productsMIXO);

    HETERO_median_change_vs_glucose_pct(c)=medianFinite_hvp(hDelta(:));
    MIXO_median_change_vs_glucose_pct(c)=medianFinite_hvp(mDelta(:));
    HETERO_median_carbon_yield(c)=medianFinite_hvp(hYield(:));
    MIXO_median_carbon_yield(c)=medianFinite_hvp(mYield(:));

    hFluxCommon=productMatrix_hvp(HF(rows,:),productsCommon);
    mFluxCommon=productMatrix_hvp(MF(rows,:),productsCommon);
    validFlux=isfinite(hFluxCommon) & isfinite(mFluxCommon) & hFluxCommon>fluxTol;
    if any(validFlux(:))
        x=100*(mFluxCommon(validFlux)./hFluxCommon(validFlux)-1);
        Median_MIXO_vs_HETERO_flux_change_pct(c)=medianFinite_hvp(x);
    end

    hYieldCommon=productMatrix_hvp(HY(rows,:),productsCommon);
    mYieldCommon=productMatrix_hvp(MY(rows,:),productsCommon);
    validYield=isfinite(hYieldCommon) & isfinite(mYieldCommon) & hYieldCommon>yieldTol;
    if any(validYield(:))
        x=100*(mYieldCommon(validYield)./hYieldCommon(validYield)-1);
        Median_MIXO_vs_HETERO_carbon_yield_change_pct(c)=medianFinite_hvp(x);
    end
end

T=table(Substrate_class,Unique_model_sources, ...
    HETERO_median_change_vs_glucose_pct,MIXO_median_change_vs_glucose_pct, ...
    Median_MIXO_vs_HETERO_flux_change_pct,HETERO_median_carbon_yield, ...
    MIXO_median_carbon_yield,Median_MIXO_vs_HETERO_carbon_yield_change_pct);
end

function T = buildOverallTrophicSummary_hvp(HF,MF,products,fluxTol)
[~,ia]=unique(string(HF.Resolved_exchange),'stable');
HFu=HF(ia,:); MFu=MF(ia,:);
hFlux=productMatrix_hvp(HFu,products);
mFlux=productMatrix_hvp(MFu,products);

validFlux=isfinite(hFlux) & isfinite(mFlux) & hFlux>fluxTol;
fluxChange=100*(mFlux(validFlux)./hFlux(validFlux)-1);

Setting=[ ...
    "Unique model carbon-source/reserve reactions"
    "Matched substrate-product flux comparisons"
    "Median MIXO vs HETERO product-flux change"
    ];
Value=[ ...
    string(height(HFu))
    string(sum(validFlux(:)))
    sprintf('%+.3f%%',medianFinite_hvp(fluxChange))
];
T=table(Setting,Value);
end

function P=selectProducts_hvp(products,selectedKeys)
% Return product structs matching a literature-guided trophic-state key set,
% preserving the master product order.
allKeys=string({products.key});
keep=false(size(allKeys));
for i=1:numel(selectedKeys)
    keep=keep | strcmpi(allKeys,string(selectedKeys(i)));
end
P=products(keep);
end

function M=productMatrix_hvp(T,products)
M=nan(height(T),numel(products));
for p=1:numel(products)
    varName=matlab.lang.makeValidName(char(products(p).key));
    M(:,p)=T.(varName);
end
end

function v=medianFinite_hvp(x)
x=x(isfinite(x));
if isempty(x), v=NaN; else, v=median(x); end
end


%% SUBSTRATE-SCREEN HELPERS

function cfg = resolveBiomassMetabolites_hvp(m,cfg)
mets.protein=requireMetCandidates_hvp(m,{'m_PROTEIN_TOTAL_POOL_AUTO_Cyto[c]','PROTEIN_TOTAL_POOL_AUTO_CYTO','m_PROTEIN_TOTAL_POOL_Cyto[c]','PROTEIN_TOTAL_POOL_CYTO'});
mets.carb=requireMetCandidates_hvp(m,{'m_CARBOHYDRATE_TOTAL_POOL_AUTO_Cyto[c]','CARBOHYDRATE_TOTAL_POOL_AUTO_CYTO','m_CARBOHYDRATE_TOTAL_POOL_Cyto[c]','CARBOHYDRATE_TOTAL_POOL_CYTO'});
mets.lipid=requireMetCandidates_hvp(m,{'m_LIPID_TOTAL_POOL_AUTO_Cyto[c]','LIPID_TOTAL_POOL_AUTO_CYTO','m_LIPID_TOTAL_POOL_Cyto[c]','LIPID_TOTAL_POOL_CYTO'});
mets.rna=requireMetCandidates_hvp(m,{'m_RNA_TOTAL_POOL_AUTO_Cyto[c]','RNA_TOTAL_POOL_AUTO_CYTO','m_RNA_TOTAL_POOL_Cyto[c]','RNA_TOTAL_POOL_CYTO'});
mets.dna=requireMetCandidates_hvp(m,{'m_DNA_TOTAL_POOL_AUTO_Cyto[c]','DNA_TOTAL_POOL_AUTO_CYTO','m_DNA_TOTAL_POOL_Cyto[c]','DNA_TOTAL_POOL_CYTO'});
mets.atp=requireMetCandidates_hvp(m,{'m_ATP_Cyto[c]','ATP_CYTO'});
mets.water=requireMetCandidates_hvp(m,{'m_WATER_Cyto[c]','WATER_CYTO'});
mets.adp=requireMetCandidates_hvp(m,{'m_ADP_Cyto[c]','ADP_CYTO'});
mets.pi=requireMetCandidates_hvp(m,{'m_Pi_Cyto[c]','PI_CYTO'});
mets.proton=requireMetCandidates_hvp(m,{'m_PROTON_Cyto[c]','PROTON_CYTO'});

pig.zea=requireMetCandidates_hvp(m,{'m_CPD1F-130_Thyl[t]','CPD1F-130_THYL'});
pig.chla=requireMetCandidates_hvp(m,{'m_CHLOROPHYLL-A_Plas[p]','m_CHLOROPHYLL-A_Thyl[t]', ...
                                    'CHLOROPHYLL-A_PLAS','CHLOROPHYLL-A_THYL'});
pig.beta=requireMetCandidates_hvp(m,{'m_CPD1F-129_Thyl[t]','CPD1F-129_THYL'});
pig.pcb=requireMetCandidates_hvp(m,{'m_3Z-PHYCOCYANOBILIN_Plas[p]','3Z-PHYCOCYANOBILIN_PLAS'});

cfg.resolvedBOFMets=mets;
cfg.resolvedPigmentMets=pig;
end

function [m,sourceRxn] = openSingleSubstrate_hvp(m,subRxn,qSub,mode,cfg,isReserve)
for i=1:numel(cfg.allMappedCarbonRxns)
    r=cfg.allMappedCarbonRxns{i};
    if any(strcmp(m.rxns,r))
        m=closeUptake_hvp(m,r);
    end
end
if isfield(cfg,'reserveSourceRxn') && strlength(string(cfg.reserveSourceRxn))>0 && ...
        any(strcmp(m.rxns,cfg.reserveSourceRxn))
    m=closeUptake_hvp(m,cfg.reserveSourceRxn);
end

if isfield(cfg,'reserveMobilizationRxn') && strlength(string(cfg.reserveMobilizationRxn))>0 && ...
        any(strcmp(m.rxns,cfg.reserveMobilizationRxn))
    m=setDirectBounds_hvp(m,cfg.reserveMobilizationRxn,0,0);
end

if isReserve
    if ~isfield(cfg,'reserveSourceRxn') || strlength(string(cfg.reserveSourceRxn))==0
        error('Intracellular storage alpha-glucan reserve source was not configured.');
    end
    sourceRxn=char(cfg.reserveSourceRxn);
    m=setDirectBounds_hvp(m,cfg.reserveMobilizationRxn,0,1000);
else
    sourceRxn=subRxn;
end

m=openUptake_hvp(m,sourceRxn,qSub,false);
assertSingleSubstrate_hvp(m,subRxn,sourceRxn,qSub,mode,cfg,isReserve);
end

function assertModeBase_hvp(m,mode,cfg)
mode=upper(string(mode));
tol=1e-10;

for i=1:numel(cfg.allMappedCarbonRxns)
    r=cfg.allMappedCarbonRxns{i};
    if any(strcmp(m.rxns,r)) && isUptakeOpen_hvp(m,r,tol)
        error('%s base model has unintended organic-carbon uptake through %s.',mode,r);
    end
end
if isfield(cfg,'reserveSourceRxn') && strlength(string(cfg.reserveSourceRxn))>0 && ...
        any(strcmp(m.rxns,cfg.reserveSourceRxn)) && ...
        isUptakeOpen_hvp(m,cfg.reserveSourceRxn,tol)
    error('%s base model has unintended intracellular storage-reserve mobilization.',mode);
end

if mode=="HETERO"
    if isUptakeOpen_hvp(m,cfg.co2Rxn,tol)
        error('HETERO external CO2 uptake must be closed.');
    end
    [rLB,rUB]=bounds_hvp(m,cfg.rubiscoRxn);
    if abs(rLB)>tol || abs(rUB)>tol
        error('HETERO Rubisco must be closed.');
    end
elseif mode=="MIXO"
    qCO2=uptakeCapacity_hvp(m,cfg.co2Rxn);
    if abs(qCO2-cfg.mixoCO2MaxUptake)>tol
        error('MIXO CO2 uptake cap differs from publication validation.');
    end
    [rLB,rUB]=bounds_hvp(m,cfg.rubiscoRxn);
    if abs(rLB-cfg.rubiscoMinFluxMIXO)>tol || abs(rUB-1000)>tol
        error('MIXO Rubisco bounds differ from publication validation.');
    end
elseif mode=="AUTO"
    qCO2=uptakeCapacity_hvp(m,cfg.co2Rxn);
    if abs(qCO2-cfg.autoCO2MaxUptake)>tol
        error('AUTO CO2 uptake cap differs from publication validation.');
    end
    [rLB,rUB]=bounds_hvp(m,cfg.rubiscoRxn);
    if abs(rLB-cfg.rubiscoMinFluxAUTO)>tol || abs(rUB-1000)>tol
        error('AUTO Rubisco bounds differ from publication validation.');
    end
else
    error('Unsupported mode: %s',mode);
end
end

function assertSingleSubstrate_hvp(m,subRxn,sourceRxn,qSub,mode,cfg,isReserve)
tol=1e-10;
for i=1:numel(cfg.allMappedCarbonRxns)
    r=cfg.allMappedCarbonRxns{i};
    if ~any(strcmp(m.rxns,r)), continue; end
    isOpen=isUptakeOpen_hvp(m,r,tol);
    if ~isReserve && strcmp(r,subRxn)
        if ~isOpen
            error('%s substrate uptake is not open: %s',mode,subRxn);
        end
        q=uptakeCapacity_hvp(m,r);
        if abs(q-qSub)>tol
            error('%s substrate uptake cap mismatch for %s.',mode,subRxn);
        end
    elseif isOpen
        error('%s has unintended organic-carbon uptake through %s.',mode,r);
    end
end

if isfield(cfg,'reserveSourceRxn') && strlength(string(cfg.reserveSourceRxn))>0 && ...
        any(strcmp(m.rxns,cfg.reserveSourceRxn))
    reserveOpen=isUptakeOpen_hvp(m,cfg.reserveSourceRxn,tol);
    if isReserve
        if ~reserveOpen || ~strcmp(sourceRxn,cfg.reserveSourceRxn)
            error('%s intracellular storage alpha-glucan reserve source is not open.',mode);
        end
        q=uptakeCapacity_hvp(m,cfg.reserveSourceRxn);
        if abs(q-qSub)>tol
            error('%s intracellular reserve mobilization cap mismatch.',mode);
        end
    elseif reserveOpen
        error('%s has unintended intracellular storage-reserve mobilization.',mode);
    end
end
end

function [subUptake,organicC,co2Uptake,totalCarbon,photonUptake,o2Uptake] = ...
    substrateUptakeMetrics_hvp(m,mode,sol,subRxn,nC,cfg)

subUptake=NaN; organicC=NaN; co2Uptake=NaN; totalCarbon=NaN;
photonUptake=NaN; o2Uptake=NaN;

if ~isFeasibleSolution_hvp(sol) || ~isfield(sol,'x') || isempty(sol.x)
    return;
end

subUptake=uptakeMagnitudeFromSolution_hvp(m,subRxn,sol.x);
organicC=nC*subUptake;

mode=upper(string(mode));
if mode=="MIXO"
    co2Uptake=uptakeMagnitudeFromSolution_hvp(m,cfg.co2Rxn,sol.x);
else
    co2Uptake=0;
end

totalCarbon=organicC+co2Uptake;
photonUptake=uptakeMagnitudeFromSolution_hvp(m,cfg.photonRxn,sol.x);
o2Uptake=uptakeMagnitudeFromSolution_hvp(m,cfg.o2Rxn,sol.x);
end

function [yieldPC,yieldCC,productCFlux] = ...
    productYieldsGeneral_hvp(productFlux,productCeq,totalCarbon,tol)

yieldPC=NaN; yieldCC=NaN; productCFlux=NaN;
if ~isfinite(productFlux) || ~isfinite(productCeq), return; end

productCFlux=productFlux*productCeq;
if isfinite(totalCarbon) && totalCarbon>tol
    yieldPC=productFlux/totalCarbon;
    yieldCC=productCFlux/totalCarbon;
end
end

function T = buildFixedMuYieldMatrix_hvp( ...
    modeModel,mode,biomassRxn,substrateName,substrateClass, ...
    resolvedExchange,carbonAtoms,uptakeCap,isReserveSubstrate, ...
    selectedProducts,allProducts,cfg)
% Secondary fixed-growth analysis at one exact biomass flux. Rows are
% substrates and columns are state-assigned HVPs only.
%
% HETERO:
%   yield = product-C / organic-C actually taken up.
%
% MIXO:
%   apparent organic-C yield = product-C / organic-C actually taken up;
%   total-C yield = product-C / (organic-C + assimilated CO2-C).
% The total-C yield is the mass-balance carbon-efficiency metric. Both are
% exported because the organic-C-normalized value is useful for process
% accounting but can include product carbon originating from CO2.

nS=numel(substrateName);
nP=numel(selectedProducts);
Yorganic=nan(nS,nP);
Ytotal=nan(nS,nP);
organicCByProduct=nan(nS,nP);
co2CByProduct=nan(nS,nP);
totalCByProduct=nan(nS,nP);

for s=1:nS
    subRxn=char(resolvedExchange(s));
    nC=carbonAtoms(s);
    qSub=uptakeCap(s);
    isReserve=isReserveSubstrate(s);

    [mBase,sourceRxn]=openSingleSubstrate_hvp( ...
        modeModel,subRxn,qSub,mode,cfg,isReserve);

    % Check once that this substrate can sustain the common biomass flux.
    mFeas=closeAllProductOutputs_hvp(mBase,allProducts);
    mFeas=setBiomassExact_hvp(mFeas,biomassRxn,cfg.fixedMu_h);
    mFeas=setObjective_hvp(mFeas,biomassRxn);
    solFeas=optimizeCbModel(mFeas,'max');
    if ~isFeasibleSolution_hvp(solFeas)
        continue;
    end

    for p=1:nP
        if isReserve && strcmpi(string(selectedProducts(p).key),'Glycogen')
            continue;
        end

        masterPidx=findProductKey_hvp(allProducts,selectedProducts(p).key);
        m=openOnlyProductOutput_hvp(mBase,allProducts,masterPidx);
        m=setBiomassExact_hvp(m,biomassRxn,cfg.fixedMu_h);
        m=setObjective_hvp(m,selectedProducts(p).outputRxn);
        sol=optimizeCbModel(m,'max');

        if ~isFeasibleSolution_hvp(sol)
            continue;
        end

        productFlux=solutionObjectiveOrZero_hvp(sol);
        productCFlux=productFlux*selectedProducts(p).carbonEquivalent;
        [~,organicC,co2Uptake,totalCarbon,~,~]=substrateUptakeMetrics_hvp( ...
            m,mode,sol,sourceRxn,nC,cfg);

        organicCByProduct(s,p)=organicC;
        co2CByProduct(s,p)=co2Uptake;
        totalCByProduct(s,p)=totalCarbon;

        if isfinite(organicC) && organicC>cfg.yieldTol
            Yorganic(s,p)=productCFlux/organicC;
        end
        if isfinite(totalCarbon) && totalCarbon>cfg.yieldTol
            Ytotal(s,p)=productCFlux/totalCarbon;
        end
    end
end

T=table(string(substrateName(:)),string(substrateClass(:)), ...
    'VariableNames',{'Substrate','Substrate_class'});

mode=upper(string(mode));
if mode=="MIXO"
    medianOrganicC=nan(nS,1);
    medianCO2C=nan(nS,1);
    medianTotalC=nan(nS,1);
    for s=1:nS
        medianOrganicC(s)=medianFinite_hvp(organicCByProduct(s,:));
        medianCO2C(s)=medianFinite_hvp(co2CByProduct(s,:));
        medianTotalC(s)=medianFinite_hvp(totalCByProduct(s,:));
    end
    T.Organic_C_uptake_mmolC=medianOrganicC;
    T.CO2_C_uptake_mmolC=medianCO2C;
    T.Total_C_uptake_mmolC=medianTotalC;
    co2ContributionPct=nan(nS,1);
    validTotal=isfinite(medianTotalC) & medianTotalC>cfg.yieldTol;
    co2ContributionPct(validTotal)=100*medianCO2C(validTotal)./medianTotalC(validTotal);
    T.CO2_contribution_pct=co2ContributionPct;

    for p=1:nP
        base=matlab.lang.makeValidName(char(selectedProducts(p).key));
        T.(matlab.lang.makeValidName([base '_OrganicC']))=Yorganic(:,p);
        T.(matlab.lang.makeValidName([base '_TotalC']))=Ytotal(:,p);
    end
else
    for p=1:nP
        base=matlab.lang.makeValidName(char(selectedProducts(p).key));
        T.(base)=Yorganic(:,p);
    end
end
end

function T = buildFixedMuBestSummary_hvp(Tfixed,products,mode,referenceSubstrate)
% Best-substrate summary for the secondary exact-mu analysis only.
mode=upper(string(mode));
nP=numel(products);
Product=strings(nP,1);
Best_organicC_substrate=strings(nP,1);
Best_organicC_yield=nan(nP,1);
Reference_organicC_yield=nan(nP,1);
OrganicC_change_vs_reference_pct=nan(nP,1);
Best_totalC_substrate=strings(nP,1);
Best_totalC_yield=nan(nP,1);
Reference_totalC_yield=nan(nP,1);
TotalC_change_vs_reference_pct=nan(nP,1);

refRow=find(strcmpi(string(Tfixed.Substrate),string(referenceSubstrate)),1);
if isempty(refRow)
    error('Fixed-growth reference substrate was not found: %s',referenceSubstrate);
end

for p=1:nP
    Product(p)=string(products(p).label);
    base=matlab.lang.makeValidName(char(products(p).key));

    if mode=="MIXO"
        orgVar=matlab.lang.makeValidName([base '_OrganicC']);
        totalVar=matlab.lang.makeValidName([base '_TotalC']);
    else
        orgVar=base;
        totalVar=base;
    end

    yOrg=Tfixed.(orgVar);
    yTotal=Tfixed.(totalVar);

    Reference_organicC_yield(p)=yOrg(refRow);
    Reference_totalC_yield(p)=yTotal(refRow);

    [iOrg,Best_organicC_yield(p)]=maxFinite_hvp(yOrg);
    if ~isempty(iOrg), Best_organicC_substrate(p)=string(Tfixed.Substrate(iOrg)); end
    [iTot,Best_totalC_yield(p)]=maxFinite_hvp(yTotal);
    if ~isempty(iTot), Best_totalC_substrate(p)=string(Tfixed.Substrate(iTot)); end

    if isfinite(Reference_organicC_yield(p)) && abs(Reference_organicC_yield(p))>1e-12 ...
            && isfinite(Best_organicC_yield(p))
        OrganicC_change_vs_reference_pct(p)= ...
            100*(Best_organicC_yield(p)/Reference_organicC_yield(p)-1);
    end
    if isfinite(Reference_totalC_yield(p)) && abs(Reference_totalC_yield(p))>1e-12 ...
            && isfinite(Best_totalC_yield(p))
        TotalC_change_vs_reference_pct(p)= ...
            100*(Best_totalC_yield(p)/Reference_totalC_yield(p)-1);
    end
end

T=table(Product,Best_organicC_substrate,Best_organicC_yield, ...
    Reference_organicC_yield,OrganicC_change_vs_reference_pct, ...
    Best_totalC_substrate,Best_totalC_yield,Reference_totalC_yield, ...
    TotalC_change_vs_reference_pct);
end

function T = roundNumericTable_hvp(T,nDigits)
% Round numeric variables for reader-facing Excel export only.
for j=1:width(T)
    varName=T.Properties.VariableNames{j};
    if isnumeric(T.(varName))
        T.(varName)=round(T.(varName),nDigits);
    end
end
end

function m = setBiomassExact_hvp(m,biomassRxn,muValue)
idx=find(strcmp(m.rxns,biomassRxn),1);
if isempty(idx), error('Biomass sink not found: %s',biomassRxn); end
m.lb(idx)=muValue;
m.ub(idx)=muValue;
end

function T = buildCompactSubstrateResults_hvp( ...
    HF,MF,HY,MY,HD,MD,muMax,products,fluxTol,yieldTol)
% One publication-facing row per substrate. Only metrics needed to interpret
% the glucose-relative heatmaps and the trophic-state comparison are kept.
% Product-specific results remain internal and are summarized separately in
% HVP_SUMMARY.

n=height(HF);
Substrate=string(HF.Substrate);
Substrate_class=string(HF.Class);
HETERO_mu_max_h=muMax(:,1);
MIXO_mu_max_h=muMax(:,2);
HETERO_median_HVP_change_vs_glucose_pct=nan(n,1);
MIXO_median_HVP_change_vs_glucose_pct=nan(n,1);
Median_MIXO_vs_HETERO_HVP_flux_change_pct=nan(n,1);

for i=1:n
    hFlux=productMatrix_hvp(HF(i,:),products);
    mFlux=productMatrix_hvp(MF(i,:),products);
    hDelta=productMatrix_hvp(HD(i,:),products);
    mDelta=productMatrix_hvp(MD(i,:),products);

    HETERO_median_HVP_change_vs_glucose_pct(i)=medianFinite_hvp(hDelta(:));
    MIXO_median_HVP_change_vs_glucose_pct(i)=medianFinite_hvp(mDelta(:));

    validFlux=isfinite(hFlux) & isfinite(mFlux) & hFlux>fluxTol;
    if any(validFlux(:))
        x=100*(mFlux(validFlux)./hFlux(validFlux)-1);
        Median_MIXO_vs_HETERO_HVP_flux_change_pct(i)=medianFinite_hvp(x(:));
    end
end

T=table(Substrate,Substrate_class,HETERO_mu_max_h,MIXO_mu_max_h, ...
    HETERO_median_HVP_change_vs_glucose_pct, ...
    MIXO_median_HVP_change_vs_glucose_pct, ...
    Median_MIXO_vs_HETERO_HVP_flux_change_pct);
end

function x=minFiniteScalar_hvp(v)
v=v(isfinite(v));
if isempty(v), x=NaN; else, x=min(v); end
end

function x=maxFiniteScalar_hvp(v)
v=v(isfinite(v));
if isempty(v), x=NaN; else, x=max(v); end
end

function T = buildSubstrateMatrix_hvp(results,substrateReference,products,mode,valueVar)
nS=height(substrateReference);
T=substrateReference(:,{'Substrate','Class','Resolved_exchange','Carbon_atoms'});
for p=1:numel(products)
    vals=nan(nS,1);
    for s=1:nS
        idx = strcmpi(string(results.Mode),string(mode)) & ...
              strcmp(string(results.Substrate),string(substrateReference.Substrate(s))) & ...
              strcmp(string(results.Exchange),string(substrateReference.Resolved_exchange(s))) & ...
              strcmp(string(results.Product),string(products(p).label));
        if any(idx)
            v=results.(valueVar)(find(idx,1));
            vals(s)=v;
        end
    end
    varName=matlab.lang.makeValidName(char(products(p).key));
    T.(varName)=vals;
end
end

function T = buildGrowthTable_hvp(substrateReference,muH,muM,growthFraction)
T=substrateReference(:,{'Substrate','Class','Resolved_exchange', ...
    'Analysis_source_reaction','Carbon_source_role','Carbon_atoms', ...
    'Organic_uptake_cap_mmol'});
T.HETERO_mu_max_h=muH;
T.MIXO_mu_max_h=muM;
T.HETERO_product_growth_floor_h=growthFraction*muH;
T.MIXO_product_growth_floor_h=growthFraction*muM;
T.HETERO_product_growth_floor_h(muH<=0)=NaN;
T.MIXO_product_growth_floor_h(muM<=0)=NaN;
end

function T = buildSubstrateHVPMethods_hvp(cfg,nS,nP,alpha,k,heteroHeatmapLimitPct,mixoHeatmapLimitPct)
Setting = [ ...
    "Substrates analysed"
    "Products analysed"
    "Conditions"
    "Organic carbon normalization"
    "Substrate opening rule"
    "Heatmap reference"
    "Heatmap metric"
    "HETERO heatmap scale"
    "MIXO heatmap scale"
    "Heatmap cell labels"
    "Product objective"
    "Primary production-capacity constraint"
    "Primary heatmap interpretation"
    "Secondary fixed-growth analysis"
    "Secondary HETERO yield denominator"
    "Secondary MIXO yield outputs"
    "HETERO external CO2"
    "HETERO Rubisco"
    "HETERO photon uptake cap"
    "MIXO external CO2 uptake cap"
    "MIXO Rubisco minimum flux"
    "MIXO photon uptake cap"
    "NGAM"
    "GAM"
    "HETERO photon per pigment"
    "HETERO photon per biomass"
    "Excel output"
    ];

Value = [ ...
    string(nS)
    string(nP)
    "HETERO; MIXO"
    sprintf('%.6g C-mmol gDW^-1 h^-1',cfg.screenCarbon)
    "One carbon source active at a time"
    string(cfg.heatmapReferenceSubstrate)
    "Percent change in product flux relative to D-glucose, calculated separately within HETERO and MIXO"
    sprintf('+/- %.0f%%',heteroHeatmapLimitPct)
    sprintf('+/- %.0f%%',mixoHeatmapLimitPct)
    sprintf('HETERO labels >= %.0f%%; MIXO labels >= %.0f%%; each product maximum is labelled', ...
        cfg.heteroHeatmapAnnotationThresholdPct,cfg.mixoHeatmapAnnotationThresholdPct)
    "Each native HVP output reaction maximized separately"
    sprintf('>= %.0f%% of substrate- and condition-specific maximum growth',100*cfg.productGrowthFraction)
    "Theoretical product-flux capacity relative to D-glucose within the same trophic state; not a carbon yield"
    sprintf('Exact biomass flux mu = %.3f h^-1; re-optimize only the state-assigned HVPs',cfg.fixedMu_h)
    "C-mol product-C per C-mol organic carbon actually taken up"
    "Apparent yield vs organic C and mass-balance yield vs total C (organic C + assimilated CO2-C); median CO2-C contribution is exported for each substrate"
    "Closed"
    "Closed"
    sprintf('%.6g',cfg.heteroPhotonMaxUptake)
    sprintf('%.6g',cfg.mixoCO2MaxUptake)
    sprintf('%.6g',cfg.rubiscoMinFluxMIXO)
    sprintf('%.6g',cfg.mixoPhotonMaxUptake)
    sprintf('%.6g mmol ATP gDW^-1 h^-1',cfg.NGAM)
    sprintf('%.6g mmol ATP gDW^-1 biomass',cfg.GAM)
    sprintf('%.9g',alpha)
    sprintf('%.9g',k)
    "Six reader-facing sheets: summary, AUTO, HETERO, MIXO, substrate reference, and methods/product reference"
    ];
T=table(Setting,Value);
end


%% ======================================================================
%% PRODUCT DEFINITIONS / MAPPING
%% ======================================================================

function P = defineProducts_hvp()
% Publication-facing products. Every product MUST resolve to its curated
% native cytosolic output reaction inherited from the original model
% definition (x_PRODUCT <- PRODUCT_Cyto). No substitute demand is allowed.

P = repmat(struct( ...
    'key',"",'label',"",'class',"",'outputRxnCandidates',{{}}, ...
    'outputRxn',"",'metID',"",'doi',"",'evidenceNote',"", ...
    'unitNote',"",'carbonEquivalent',NaN,'carbonEquivalentNote',""),9,1);

P(1).key="Glycogen";
P(1).label="Glycogen / storage alpha-glucan";
P(1).class="Storage carbohydrate";
P(1).outputRxnCandidates={'STARCH_Cyto_tx','R_STARCH_Cyto_tx'};
P(1).doi="10.1016/j.biortech.2015.11.014; 10.1016/j.ijbiomac.2016.04.051";
P(1).evidenceNote="Direct G. sulphuraria glycogen accumulation and structural characterization; GSM uses STARCH_Cyto for the storage alpha-glucan.";
P(1).unitNote="mmol glucosyl-equivalent gDW^-1 h^-1 (model STARCH repeat unit)";

P(2).key="TAG";
P(2).label="Triacylglycerol (TAG)";
P(2).class="Neutral lipid";
P(2).outputRxnCandidates={'Triacylglycerols_Cyto_TOTAL_Cyto_tx','R_Triacylglycerols_Cyto_TOTAL_Cyto_tx','TAG_Cyto_tx','R_TAG_Cyto_tx'};
P(2).doi="10.1016/j.biortech.2015.11.014";
P(2).evidenceNote="Native total-TAG output Triacylglycerols_Cyto_TOTAL_Cyto_tx (TAG_Cyto_tx in the source definition) is used directly; no artificial TAG demand is introduced.";
P(2).unitNote="model TAG output flux gDW^-1 h^-1";

P(3).key="C18_2";
P(3).label="Linoleic acid (C18:2)";
P(3).class="PUFA";
P(3).outputRxnCandidates={'LINOLEIC_ACID_Cyto_tx','R_LINOLEIC_ACID_Cyto_tx'};
P(3).doi="10.1039/C2FO30198A";
P(3).evidenceNote="Direct G. sulphuraria fatty-acid data; C18:2 is a major PUFA and is substantially more abundant than C18:3 in the cited AUTO profile.";
P(3).unitNote="mmol gDW^-1 h^-1";

P(4).key="BetaCarotene";
P(4).label="beta-Carotene";
P(4).class="Pigment";
P(4).outputRxnCandidates={'CPD1F-129_Cyto_tx','R_CPD1F-129_Cyto_tx'};
P(4).doi="10.3389/fpls.2022.978246";
P(4).evidenceNote="beta-Carotene was directly quantified in G. sulphuraria and shows strong trophic-state dependence.";
P(4).unitNote="mmol gDW^-1 h^-1";

P(5).key="PCB";
P(5).label="Phycocyanobilin (PCB)";
P(5).class="Pigment";
P(5).outputRxnCandidates={'3Z-PHYCOCYANOBILIN_Cyto_tx','R_3Z-PHYCOCYANOBILIN_Cyto_tx'};
P(5).doi="10.1111/febs.70391";
P(5).evidenceNote="Direct biochemical validation of the G. sulphuraria PCB biosynthetic route.";
P(5).unitNote="mmol gDW^-1 h^-1";

P(6).key="Floridoside";
P(6).label="Floridoside";
P(6).class="Compatible solute / glycoside";
P(6).outputRxnCandidates={'FLORIDOSIDE_Cyto_tx','R_FLORIDOSIDE_Cyto_tx'};
P(6).doi="10.1111/nph.13108; 10.1186/s13568-016-0244-6";
P(6).evidenceNote="Native FLORIDOSIDE_Cyto_tx output is used directly for free floridoside.";
P(6).unitNote="mmol gDW^-1 h^-1";

P(7).key="Squalene";
P(7).label="Squalene";
P(7).class="Isoprenoid";
P(7).outputRxnCandidates={'SQUALENE_Cyto_tx','R_SQUALENE_Cyto_tx'};
P(7).doi="";
P(7).evidenceNote="Native SQUALENE_Cyto_tx output is used; squalene is treated as a pathway target without assigning a direct G. sulphuraria accumulation DOI.";
P(7).unitNote="mmol gDW^-1 h^-1";

P(8).key="Ergosterol";
P(8).label="Ergosterol";
P(8).class="Isoprenoid";
P(8).outputRxnCandidates={'ERGOSTEROL_Cyto_tx','R_ERGOSTEROL_Cyto_tx'};
P(8).doi="10.3390/md18030169; 10.3390/ijms22031247";
P(8).evidenceNote="Ergosterol was directly detected in G. sulphuraria and is modulated by trophic/light conditions.";
P(8).unitNote="mmol gDW^-1 h^-1";

P(9).key="Phytol";
P(9).label="Phytol";
P(9).class="Isoprenoid";
P(9).outputRxnCandidates={'PHYTOL_Cyto_tx','R_PHYTOL_Cyto_tx'};
P(9).doi="10.3390/md18030169; 10.3390/ijms22031247";
P(9).evidenceNote="Phytol was directly detected in G. sulphuraria and shows trophic/light dependence.";
P(9).unitNote="mmol gDW^-1 h^-1";
end

function m = ensurePublicationTagProductAssembly_hvp(m)
% Recreate the legacy generic TAG product-pool producer only for analysis.
% The final SBML now uses condition-specific AUTO/MIXO/HETERO TAG biomass
% pools, while the publication HVP target remains the generic native output
% R_Triacylglycerols_Cyto_TOTAL_Cyto_tx.  The former generic producer used
% exactly the HETERO TAG composition, so copy that stoichiometry when needed.

genericRxn = resolveRxn_hvp(m,{ ...
    'R_Triacylglycerols_Cyto_BIO_NO_LIPID_TOTAL_bm', ...
    'Triacylglycerols_Cyto_BIO_NO_LIPID_TOTAL_bm'},true);
if strlength(string(genericRxn))>0
    return;
end

sourceRxn = requireRxn_hvp(m,{ ...
    'R_Triacylglycerols_Cyto_BIO_NO_LIPID_TOTAL_HETERO_bm', ...
    'Triacylglycerols_Cyto_BIO_NO_LIPID_TOTAL_HETERO_bm'});
genericMet = requireMetCandidates_hvp(m,{ ...
    'm_Triacylglycerols_Cyto_TOTAL_Cyto[c]', ...
    'Triacylglycerols_Cyto_TOTAL_Cyto'});

j = find(strcmp(m.rxns,sourceRxn),1);
if isempty(j)
    error('HETERO TAG assembly reaction missing after resolution.');
end
s = full(m.S(:,j));
react = find(s < -1e-12);
prod = find(s > 1e-12);
if isempty(react) || numel(prod)~=1
    error('Unexpected HETERO TAG assembly stoichiometry in %s.',sourceRxn);
end

metList = [m.mets(react); {genericMet}];
coeffList = [s(react); s(prod)];

m = addReaction(m,'R_Triacylglycerols_Cyto_BIO_NO_LIPID_TOTAL_bm', ...
    'reactionName','publication generic TAG product-pool assembly', ...
    'metaboliteList',metList, ...
    'stoichCoeffList',coeffList, ...
    'lowerBound',0,'upperBound',1000, ...
    'subSystem','HVP product definition');
m = padConstraintColumns_hvp(m);
end

function [P,T] = resolveProducts_hvp(model,P)
% Exact native outputs are mandatory. Resolve each and infer its intracellular
% product metabolite from the one-metabolite SBML boundary reaction.
for i=1:numel(P)
    r=resolveRxn_hvp(model,string(P(i).outputRxnCandidates),true);
    if strlength(string(r))==0
        error(['Required native HVP output reaction is missing for %s. ' ...
            'Expected %s. No artificial demand will be created.'], ...
            P(i).label,strjoin(P(i).outputRxnCandidates,', '));
    end
    P(i).outputRxn=string(r);
    P(i).metID=string(productMetFromNativeOutput_hvp(model,char(r),P(i).label));
    [P(i).carbonEquivalent,P(i).carbonEquivalentNote] = ...
        productCarbonEquivalent_hvp(model,char(P(i).metID),char(P(i).key));
end
T=refreshProductReference_hvp(P);

fprintf('\nNative HVP output reactions resolved:\n');
for i=1:numel(P)
    fprintf('  %-20s -> %-35s [%s], C-eq = %.6g\n', ...
        P(i).key,P(i).outputRxn,P(i).metID,P(i).carbonEquivalent);
end
end

function met = productMetFromNativeOutput_hvp(model,rxn,label)
j=find(strcmp(model.rxns,rxn),1);
if isempty(j), error('Native product output reaction not found: %s',rxn); end
nz=find(abs(full(model.S(:,j)))>1e-12);
if numel(nz)~=1
    error(['Native product output %s for %s must contain exactly one model ' ...
        'metabolite after x_* external species removal; found %d.'], ...
        rxn,label,numel(nz));
end
coef=full(model.S(nz,j));
if coef>=0
    error(['Native product output %s for %s must consume the intracellular ' ...
        'product in the forward direction; stoichiometry is %.6g.'],rxn,label,coef);
end
met=char(model.mets{nz});
end

function T = refreshProductReference_hvp(P)
n=numel(P);
Class=strings(n,1); Product=strings(n,1); Target_metabolite=strings(n,1);
Native_output_reaction=strings(n,1); DOI=strings(n,1); Evidence_note=strings(n,1);
Model_unit_note=strings(n,1); Product_C_equivalent=nan(n,1);
Carbon_equivalent_note=strings(n,1);
for i=1:n
    Class(i)=P(i).class; Product(i)=P(i).label;
    Target_metabolite(i)=P(i).metID; Native_output_reaction(i)=P(i).outputRxn;
    DOI(i)=P(i).doi; Evidence_note(i)=P(i).evidenceNote;
    Model_unit_note(i)=P(i).unitNote;
    Product_C_equivalent(i)=P(i).carbonEquivalent;
    Carbon_equivalent_note(i)=P(i).carbonEquivalentNote;
end
T=table(Class,Product,Target_metabolite,Native_output_reaction, ...
    Product_C_equivalent,Carbon_equivalent_note,DOI,Evidence_note,Model_unit_note);
end

function [cEq,note] = productCarbonEquivalent_hvp(model,met,key)
% Carbon equivalent used for carbon-recovery yield. For ordinary metabolites
% this is the number of C atoms in the FBC formula. The composite TAG pool
% has symbolic formula R, so its effective carbon content is derived from
% the native weighted TAG-pool assembly reaction.
mi=find(strcmp(model.mets,met),1);
if isempty(mi), error('Product metabolite not found for carbon accounting: %s',met); end
formula="";
if isfield(model,'metFormulas') && numel(model.metFormulas)>=mi && ...
        ~isempty(model.metFormulas{mi})
    formula=string(model.metFormulas{mi});
end
c=parseCarbonAtoms_hvp(formula);
if isfinite(c) && c>0
    cEq=c;
    note="From target-metabolite formula " + formula;
    return;
end

if strcmpi(key,'TAG')
    assembly=requireRxn_hvp(model,{ ...
        'R_Triacylglycerols_Cyto_BIO_NO_LIPID_TOTAL_bm', ...
        'Triacylglycerols_Cyto_BIO_NO_LIPID_TOTAL_bm'});
    j=find(strcmp(model.rxns,assembly),1);
    outCoef=full(model.S(mi,j));
    if outCoef<=0
        error('TAG total pool is not produced by %s as expected.',assembly);
    end
    react=find(full(model.S(:,j)) < -1e-12);
    carbonFlux=0;
    for k=1:numel(react)
        rk=react(k);
        rf="";
        if isfield(model,'metFormulas') && numel(model.metFormulas)>=rk && ...
                ~isempty(model.metFormulas{rk})
            rf=string(model.metFormulas{rk});
        end
        rc=parseCarbonAtoms_hvp(rf);
        if ~isfinite(rc) || rc<=0
            error('Cannot derive TAG carbon equivalent: formula missing for %s.',model.mets{rk});
        end
        carbonFlux = carbonFlux + (-full(model.S(rk,j)))*rc;
    end
    cEq=carbonFlux/outCoef;
    note=sprintf(['Derived from %s weighted reactants; effective TAG-pool ' ...
        'carbon equivalent = %.6g C-mmol per mmol model-pool flux'],assembly,cEq);
    return;
end

error('No numeric carbon formula/equivalent available for product %s (%s).',key,met);
end

function c = parseCarbonAtoms_hvp(formula)
c=NaN;
formula=char(string(formula));
if isempty(formula), return; end
tok=regexp(formula,'(?:^|[^A-Za-z])C(\d*)','tokens','once');
if isempty(tok)
    if startsWith(formula,'C')
        tok=regexp(formula,'^C(\d*)','tokens','once');
    else
        return;
    end
end
if isempty(tok), return; end
if isempty(tok{1}), c=1; else, c=str2double(tok{1}); end
end

%% ======================================================================
%% PRODUCT CONDITION / OPTIMIZATION HELPERS
%% ======================================================================

function [lb,ub]=bounds_hvp(m,rxn)
idx=find(strcmp(m.rxns,rxn),1);
if isempty(idx), error('Reaction not found: %s',rxn); end
lb=m.lb(idx); ub=m.ub(idx);
end

function tf=isUptakeOpen_hvp(m,rxn,tol)
if nargin<3, tol=1e-10; end
idx=find(strcmp(m.rxns,rxn),1);
if isempty(idx), tf=false; return; end
sgn=uptakeFluxSign_hvp(m,rxn);
if sgn<0, tf=m.lb(idx)<-tol; else, tf=m.ub(idx)>tol; end
end

function q=uptakeCapacity_hvp(m,rxn)
idx=find(strcmp(m.rxns,rxn),1);
if isempty(idx), error('Reaction not found: %s',rxn); end
sgn=uptakeFluxSign_hvp(m,rxn);
if sgn<0, q=max(0,-m.lb(idx)); else, q=max(0,m.ub(idx)); end
end

function m = closeAllProductOutputs_hvp(m,P)
% Close all curated native product outputs during growth optimization.
for i=1:numel(P)
    r=char(P(i).outputRxn);
    if ~any(strcmp(m.rxns,r))
        error('Required native product output disappeared from model: %s',r);
    end
    m=setDirectBounds_hvp(m,r,0,0);
end
end

function m = openOnlyProductOutput_hvp(m,P,selectedIndex)
% Open exactly one native product output in the forward secretion direction.
m=closeAllProductOutputs_hvp(m,P);
r=char(P(selectedIndex).outputRxn);
m=setDirectBounds_hvp(m,r,0,1000);
end

function m = setBiomassFloor_hvp(m,biomassRxn,floorValue)
idx=find(strcmp(m.rxns,biomassRxn),1);
if isempty(idx), error('Biomass sink not found: %s',biomassRxn); end
m.lb(idx)=max(0,floorValue);
m.ub(idx)=max(m.ub(idx),m.lb(idx));
end

function v = solutionObjectiveOrZero_hvp(sol)
if isFeasibleSolution_hvp(sol) && isfield(sol,'f') && isfinite(sol.f)
    v=max(0,sol.f);
else
    v=0;
end
end

function f = resolveLocalFile_hvp(f,scriptDir)
f = char(string(f));
if isfile(f), return; end
candidate = fullfile(scriptDir,f);
if isfile(candidate), f = candidate; end
end

function ensureCobraReady_hvp()
required = {'readCbModel','optimizeCbModel','addReaction','addMetabolite'};
missing = {};
for i=1:numel(required)
    if exist(required{i},'file')~=2
        missing{end+1}=required{i}; %#ok<AGROW>
    end
end
if ~isempty(missing)
    error(['COBRA Toolbox functions are not on the MATLAB path: %s. ' ...
           'Start/configure COBRA Toolbox before running this script.'], ...
           strjoin(missing,', '));
end

% IMPORTANT: do not call initCobraToolbox or changeCobraSolver here.
% On some COBRA installations changeCobraSolver triggers toolbox setup,
% git/submodule checks and solver probing. This analysis must not modify or
% reinitialize the user's COBRA installation. optimizeCbModel will use the
% LP solver already configured in the current MATLAB/COBRA session.
solverName="configured COBRA LP solver";
try
    global CBT_LP_SOLVER
    if ~isempty(CBT_LP_SOLVER)
        solverName=string(CBT_LP_SOLVER);
    end
catch
end
fprintf('COBRA ready; using %s. No toolbox initialization will be run.\n',solverName);
end

function model = loadModel_hvp(modelFile)
% readCbModel can print harmless annotation/subSystem warnings for this SBML.
% Capture those messages locally so the publication analysis log remains
% readable. Warning state is always restored, including after an error.
warnState=warning;
cleanupObj=onCleanup(@() warning(warnState)); %#ok<NASGU>
warning('off','all');

try
    evalc('model = readCbModel(modelFile);');
catch ME1
    try
        evalc('model = readCbModel(modelFile,''fileType'',''SBML'');');
    catch ME2
        error(['Unable to load SBML with readCbModel.\nFirst error: %s\n' ...
               'Second error: %s\nEnsure libSBML/TranslateSBML required by ' ...
               'your COBRA installation is available.'],ME1.message,ME2.message);
    end
end

fprintf('Model loaded quietly: %d reactions, %d metabolites.\n', ...
    numel(model.rxns),numel(model.mets));
end

function validateBasicModel_hvp(model)
required={'rxns','mets','S','lb','ub'};
for i=1:numel(required)
    if ~isfield(model,required{i})
        error('Loaded model lacks required COBRA field: %s',required{i});
    end
end
if size(model.S,1)~=numel(model.mets) || size(model.S,2)~=numel(model.rxns)
    error('Model S dimensions do not match mets/rxns.');
end
if numel(model.lb)~=numel(model.rxns) || numel(model.ub)~=numel(model.rxns)
    error('Model bound vectors do not match rxns.');
end
end


%% ======================================================================
%% ID / WORKBOOK HELPERS
%% ======================================================================

function r = requireRxn_hvp(model,candidates)
r = resolveRxn_hvp(model,string(candidates),true);
if strlength(string(r))==0
    error('Required reaction not found. Tried: %s',strjoin(candidates,', '));
end
r = char(r);
end

function r = resolveRxn_hvp(model,query,allowEmpty)
if nargin<3, allowEmpty=false; end
r = "";
qList = string(query(:));
rxns = string(model.rxns(:));

for q = qList'
    q = strtrim(q);
    if strlength(q)==0, continue; end

    idx=find(strcmp(rxns,q),1);
    if ~isempty(idx), r=rxns(idx); return; end

    if startsWith(q,"R_")
        alt=extractAfter(q,2);
    else
        alt="R_"+q;
    end
    idx=find(strcmp(rxns,alt),1);
    if ~isempty(idx), r=rxns(idx); return; end

    nq=normalizeId_hvp(q);
    nr=strings(size(rxns));
    for kk=1:numel(rxns)
        nr(kk)=normalizeId_hvp(rxns(kk));
    end
    idx=find(nr==nq,1);
    if ~isempty(idx), r=rxns(idx); return; end
end

if ~allowEmpty
    error('Reaction could not be resolved: %s',strjoin(cellstr(qList),', '));
end
end

function s = normalizeId_hvp(s)
s=upper(strtrim(string(s)));
s=replace(s,["__45__","__43__","__91__","__93__"],["-","+","[","]"]);
s=replace(s,[" ","[","]"],"");
s=regexprep(s,'^R_','');
s=regexprep(s,'[^A-Z0-9+\-]','_');
s=regexprep(s,'_+','_');
end

function met = requireMetCandidates_hvp(model,candidates)
mets=string(model.mets(:));
for q=string(candidates(:))'
    idx=find(strcmpi(mets,q),1);
    if ~isempty(idx), met=char(mets(idx)); return; end
end

normM=strings(size(mets));
for kk=1:numel(mets)
    normM(kk)=normalizeMetFull_hvp(mets(kk));
end
for q=string(candidates(:))'
    nq=normalizeMetFull_hvp(q);
    idx=find(normM==nq);
    if numel(idx)==1, met=char(mets(idx)); return; end
end

% Final safe fallback: compare compartment-free biochemical core only, but
% only accept a unique model metabolite.
coreM=strings(size(mets));
for kk=1:numel(mets)
    coreM(kk)=metCoreNoComp_hvp(mets(kk));
end
for q=string(candidates(:))'
    nq=metCoreNoComp_hvp(q);
    idx=find(coreM==nq);
    if numel(idx)==1, met=char(mets(idx)); return; end
end

error('Required metabolite not uniquely resolved. Tried: %s', ...
    strjoin(cellstr(string(candidates(:))),', '));
end

function s=normalizeMetFull_hvp(s)
s=upper(strtrim(string(s)));
s=replace(s,["__45__","__43__","__91__","__93__"],["-","+","[","]"]);
s=regexprep(s,'^M_','');
s=replace(s," ","");
s=regexprep(s,'[^A-Z0-9+\-\[\]]','_');
s=regexprep(s,'_+','_');
end

function s=metCoreNoComp_hvp(s)
s=normalizeMetFull_hvp(s);
s=regexprep(s,'\[[A-Z]\]$','');
s=regexprep(s,'_(CYTO|MITO|PLAS|THYL|PERO)$','');
end

function [nC,source] = inferCarbonAtoms_hvp(model,exchange,substrateName,substrateClass)
nC=NaN; source="";

if strlength(string(exchange))>0
    ridx=find(strcmp(model.rxns,char(exchange)),1);
    if ~isempty(ridx)
        nz=find(abs(full(model.S(:,ridx)))>1e-12);
        if numel(nz)==1 && isfield(model,'metFormulas') && ...
                numel(model.metFormulas)>=nz
            formula=string(model.metFormulas{nz});
            [nC0,ok]=carbonCountFromFormula_hvp(formula);
            if ok
                nC=nC0;
                source="Model metabolite formula: "+formula;
                return;
            end
        end
    end
end

name=lower(strtrim(string(substrateName)));
cls=lower(strtrim(string(substrateClass)));

if contains(cls,'disaccharide')
    nC=12;
elseif contains(cls,'hexose') || contains(cls,'deoxyhexose') || contains(cls,'hexitol')
    nC=6;
elseif contains(cls,'pentose') || contains(cls,'pentitol')
    nC=5;
elseif contains(cls,'tetritol')
    nC=4;
elseif contains(cls,'triol')
    nC=3;
elseif contains(cls,'organic acid')
    if contains(name,'acetate')
        nC=2;
    elseif contains(name,'pyruvate')
        nC=3;
    end
elseif contains(cls,'polymer') || contains(name,'starch')
    nC=6; % glucose-equivalent unit used in the sole-substrate screen
end

if isfinite(nC)
    source="substrate-class fallback";
else
    source="UNRESOLVED";
end
end

function [nC,ok]=carbonCountFromFormula_hvp(formula)
nC=NaN; ok=false;
formula=char(strtrim(string(formula)));
if isempty(formula), return; end
tok=regexp(formula,'(?:^|[^A-Za-z])C([0-9]*)','tokens','once');
if isempty(tok)
    % Standard formulas normally start with C; also handle directly.
    tok=regexp(formula,'^C([0-9]*)','tokens','once');
end
if isempty(tok), return; end
if isempty(tok{1})
    nC=1;
else
    nC=str2double(tok{1});
end
ok=isfinite(nC) && nC>0;
end

function [rxns,auditT]=identifyCarbonBoundaryRxns_hvp(model,cfg)
nrxn=numel(model.rxns);
isBoundary=false(nrxn,1);
hasCarbon=false(nrxn,1);
metID=strings(nrxn,1);
formula=strings(nrxn,1);
nC=nan(nrxn,1);

for j=1:nrxn
    nz=find(abs(full(model.S(:,j)))>1e-12);
    if numel(nz)==1
        isBoundary(j)=true;
        metID(j)=string(model.mets{nz});
        if isfield(model,'metFormulas') && numel(model.metFormulas)>=nz
            formula(j)=string(model.metFormulas{nz});
            [nC(j),hasCarbon(j)]=carbonCountFromFormula_hvp(formula(j));
        end
    end
end

idx=find(isBoundary & hasCarbon);
rxns=cellstr(string(model.rxns(idx)));

% Always include known mapped substrates and CO2 later in the caller.
auditT=table(string(model.rxns(idx)),metID(idx),formula(idx),nC(idx), ...
    'VariableNames',{'Reaction_ID','Boundary_metabolite','Formula','Carbon_atoms'});

if isempty(idx)
    fprintf(2,['NOTICE: no additional carbon boundary reactions were identified ' ...
        'from metFormulas; mapped substrate exchanges will still be closed.\n']);
end
end


%% ======================================================================
%% BIOMASS FUNCTIONS
%% ======================================================================

function [m,auditT]=closeOriginalBiomassOutputs_hvp(m)
ids=string(m.rxns(:));
if isfield(m,'rxnNames')
    names=string(m.rxnNames(:));
else
    names=repmat("",numel(ids),1);
end

rawID=lower(ids); rawName=lower(names);
normID=strings(size(ids)); normName=strings(size(names));
for kk=1:numel(ids)
    normID(kk)=replace(normalizeId_hvp(ids(kk)),"_","");
    normName(kk)=replace(normalizeId_hvp(names(kk)),"_","");
end

isBmTx=endsWith(rawID,"_bm_tx") | endsWith(rawName,"_bm_tx") | ...
       contains(normID,"BMTX") | contains(normName,"BMTX");
isTotalBiomass=contains(normID,"TOTALBIOMASS") | contains(normName,"TOTALBIOMASS");
mask=isBmTx | isTotalBiomass;

idx=find(mask);
oldLB=m.lb(idx); oldUB=m.ub(idx);
if isfield(m,'c') && numel(m.c)==numel(m.rxns)
    oldC=m.c(idx);
else
    oldC=zeros(numel(idx),1);
    m.c=zeros(numel(m.rxns),1);
end

reason=strings(numel(idx),1);
for kk=1:numel(idx)
j=idx(kk);
    if isBmTx(j) && isTotalBiomass(j)
        reason(kk)="bm_tx + TOTAL_BIOMASS";
    elseif isBmTx(j)
        reason(kk)="pre-existing bm_tx";
    else
        reason(kk)="TOTAL_BIOMASS";
    end
end

m.lb(idx)=0; m.ub(idx)=0; m.c(:)=0;

auditT=table(ids(idx),names(idx),reason,oldLB,oldUB,oldC, ...
    zeros(numel(idx),1),zeros(numel(idx),1), ...
    'VariableNames',{'Reaction_ID','Reaction_name','Closure_reason', ...
    'Original_LB','Original_UB','Original_objective','New_LB','New_UB'});
end

function [m,cfg,auditT]=implementExperimentalBOFs_hvp(m,cfg)
[m,pigH]=addExperimentalPigmentPool_hvp(m,'HETERO',cfg);
[m,pigM]=addExperimentalPigmentPool_hvp(m,'MIXO',cfg);

[m,bH,aH]=createExperimentalBOF_hvp(m,'HETERO',cfg.exp.HETERO, ...
    cfg.reference.HETERO.rnaFraction,cfg.reference.HETERO.pigmentMass, ...
    pigH.poolMet,pigH.rxn,cfg);

[m,bM,aM]=createExperimentalBOF_hvp(m,'MIXO',cfg.exp.MIXO, ...
    cfg.reference.MIXO.rnaFraction,cfg.reference.MIXO.pigmentMass, ...
    pigM.poolMet,pigM.rxn,cfg);

cfg.HETERO=bH; cfg.MIXO=bM;
auditT=[aH;aM];
end

function [m,info]=addExperimentalPigmentPool_hvp(m,mode,cfg)
mode=upper(string(mode));
pig=cfg.resolvedPigmentMets;

if mode=="HETERO"
    poolMet='m_PUBHVP_PIGMENT_TOTAL_EXP_HETERO[c]';
    rxn='PUBHVP_PIGMENT_TOTAL_EXP_HETERO_PRODUCER_bm';
    sheetName='Reaction List Heto';
    workbookRxn='PIGMENT_TOTAL_POOL_BIO_HETEROTROPH_GLUCOSE_NOFRAC';
elseif mode=="MIXO"
    poolMet='m_PUBHVP_PIGMENT_TOTAL_EXP_MIXO[c]';
    rxn='PUBHVP_PIGMENT_TOTAL_EXP_MIXO_PRODUCER_bm';
    sheetName='Reaction List Mixo';
    workbookRxn='PIGMENT_TOTAL_POOL_BIO_MIXOTROPH_NOFRAC';
else
    error('Unsupported pigment mode: %s',mode);
end

formula=readWorkbookReactionFormula_hvp(cfg.biomassFile,sheetName,workbookRxn);
cZea=coefficientForWorkbookMet_hvp(formula,'m_CPD1F-130_Thyl[t]');
try
    cChl=coefficientForWorkbookMet_hvp(formula,'m_CHLOROPHYLL-A_Plas[p]');
catch
    cChl=coefficientForWorkbookMet_hvp(formula,'m_CHLOROPHYLL-A_Thyl[t]');
end
cBeta=coefficientForWorkbookMet_hvp(formula,'m_CPD1F-129_Thyl[t]');
cPCB=coefficientForWorkbookMet_hvp(formula,'m_3Z-PHYCOCYANOBILIN_Plas[p]');

coeff=[cZea;cChl;cBeta;cPCB];
if any(~isfinite(coeff)) || sum(coeff)<=0
    error('Invalid %s pigment composition in TOTAL_BIOMASS.xlsx.',mode);
end
coeff=coeff/sum(coeff);

m=removeIfPresent_hvp(m,{rxn});
m=addMetIfMissing_hvp(m,poolMet,sprintf('publication HVP %s pigment pool',mode));
m=addReaction(m,rxn, ...
    'reactionName',sprintf('publication HVP %s pigment pool from TOTAL_BIOMASS.xlsx',mode), ...
    'metaboliteList',{pig.zea;pig.chla;pig.beta;pig.pcb;poolMet}, ...
    'stoichCoeffList',[-coeff;1], ...
    'lowerBound',0,'upperBound',1000,'subSystem','Biomass pigment pool');
m=padConstraintColumns_hvp(m);

info.poolMet=poolMet; info.rxn=rxn; info.coeff=coeff;
info.workbookSheet=sheetName; info.workbookReaction=workbookRxn;
end

function [m,b,auditT]=createExperimentalBOF_hvp( ...
    m,mode,expComp,rnaFraction,pigmentMass,pigmentPoolMet,pigmentRxn,cfg)

mode=upper(string(mode));
mets=resolveModeBOFMets_hvp(m,mode);

raw=[expComp.protein;expComp.carbohydrate;expComp.lipid; ...
     expComp.nucleic*rnaFraction;expComp.nucleic*(1-rnaFraction); ...
     pigmentMass];
represented=sum(raw);
if represented<=0, error('Invalid %s BOF composition.',mode); end
% Measured macromolecules account for 0.9 g/gDW; do not rescale them.
% Pool-level molecular recipes retain workbook-derived mode specificity.
coeff=raw;
if abs(sum(raw(1:5))-0.9)>1e-8 || any(~isfinite(coeff)) || any(coeff<0) || represented>1+1e-8
    error('Invalid unnormalized experimental BOF for %s.',mode);
end

bioMet=sprintf('m_PUBHVP_TOTAL_BIOMASS_EXP_%s[c]',mode);
assembly=sprintf('PUBHVP_TOTAL_BIOMASS_EXP_%s_bm',mode);
sink=sprintf('PUBHVP_TOTAL_BIOMASS_EXP_%s_SINK_bm_tx',mode);

m=removeIfPresent_hvp(m,{assembly,sink});
m=addMetIfMissing_hvp(m,bioMet,sprintf('publication HVP experimental %s biomass',mode));

m=addReaction(m,assembly, ...
    'reactionName',sprintf('publication HVP experimental %s BOF',mode), ...
    'metaboliteList',{mets.protein;mets.carb;mets.lipid;mets.rna;mets.dna; ...
        pigmentPoolMet;mets.atp;mets.water;mets.adp;mets.pi;mets.proton;bioMet}, ...
    'stoichCoeffList',[-coeff;-cfg.GAM;-cfg.GAM; ...
        cfg.GAM;cfg.GAM;cfg.GAM;1], ...
    'lowerBound',0,'upperBound',1000,'subSystem','Biomass synthesis');
m=padConstraintColumns_hvp(m);

m=addReaction(m,sink, ...
    'reactionName',sprintf('publication HVP experimental %s biomass sink',mode), ...
    'metaboliteList',{bioMet},'stoichCoeffList',-1, ...
    'lowerBound',0,'upperBound',1000,'subSystem','Biomass sink');
m=padConstraintColumns_hvp(m);

b=struct();
b.mode=mode; b.assemblyRxn=assembly; b.sinkRxn=sink; b.biomassMet=bioMet;
b.pigmentRxn=pigmentRxn; b.pigmentMet=pigmentPoolMet;
b.proteinCoeff=coeff(1); b.carbohydrateCoeff=coeff(2); b.lipidCoeff=coeff(3);
b.rnaCoeff=coeff(4); b.dnaCoeff=coeff(5); b.pigmentCoeff=coeff(6);
b.GAM=cfg.GAM; b.rawRepresentedMass=represented;
b.rawUnresolvedMass=max(0,1-represented);
b.source="Experimental cultivation-validation BOF";

Component=["Protein";"Carbohydrate";"Lipid";"RNA";"DNA";"Pigment"];
Mode=repmat(mode,6,1);
Coefficient=coeff;
Raw_g_gDW=raw;
Source=repmat("Experimental cultivation-validation BOF",6,1);
auditT=table(Mode,Component,Raw_g_gDW,Coefficient,Source);
end

function mets=resolveModeBOFMets_hvp(model,mode)
mode=upper(string(mode));
if ~ismember(mode,["AUTO","MIXO","HETERO"])
    error('Unsupported biomass mode: %s',mode);
end
tag=char(mode);

mets.protein=requireMetCandidates_hvp(model,{ ...
    sprintf('m_PROTEIN_TOTAL_POOL_%s_Cyto[c]',tag), ...
    sprintf('PROTEIN_TOTAL_POOL_%s_CYTO',tag), ...
    'm_PROTEIN_TOTAL_POOL_Cyto[c]','PROTEIN_TOTAL_POOL_CYTO'});
mets.carb=requireMetCandidates_hvp(model,{ ...
    sprintf('m_CARBOHYDRATE_TOTAL_POOL_%s_Cyto[c]',tag), ...
    sprintf('CARBOHYDRATE_TOTAL_POOL_%s_CYTO',tag), ...
    'm_CARBOHYDRATE_TOTAL_POOL_Cyto[c]','CARBOHYDRATE_TOTAL_POOL_CYTO'});
mets.lipid=requireMetCandidates_hvp(model,{ ...
    sprintf('m_LIPID_TOTAL_POOL_%s_Cyto[c]',tag), ...
    sprintf('LIPID_TOTAL_POOL_%s_CYTO',tag), ...
    'm_LIPID_TOTAL_POOL_Cyto[c]','LIPID_TOTAL_POOL_CYTO'});
mets.rna=requireMetCandidates_hvp(model,{ ...
    sprintf('m_RNA_TOTAL_POOL_%s_Cyto[c]',tag), ...
    sprintf('RNA_TOTAL_POOL_%s_CYTO',tag), ...
    'm_RNA_TOTAL_POOL_Cyto[c]','RNA_TOTAL_POOL_CYTO'});
mets.dna=requireMetCandidates_hvp(model,{ ...
    sprintf('m_DNA_TOTAL_POOL_%s_Cyto[c]',tag), ...
    sprintf('DNA_TOTAL_POOL_%s_CYTO',tag), ...
    'm_DNA_TOTAL_POOL_Cyto[c]','DNA_TOTAL_POOL_CYTO'});

mets.atp=requireMetCandidates_hvp(model,{'m_ATP_Cyto[c]','ATP_CYTO'});
mets.water=requireMetCandidates_hvp(model,{'m_WATER_Cyto[c]','WATER_CYTO'});
mets.adp=requireMetCandidates_hvp(model,{'m_ADP_Cyto[c]','ADP_CYTO'});
mets.pi=requireMetCandidates_hvp(model,{'m_Pi_Cyto[c]','PI_CYTO'});
mets.proton=requireMetCandidates_hvp(model,{'m_PROTON_Cyto[c]','PROTON_CYTO'});
end

function m=removeIfPresent_hvp(m,rxns)
for r=string(rxns(:))'
    if any(strcmp(m.rxns,char(r)))
        m=removeRxns(m,{char(r)});
    end
end
end

function m=addMetIfMissing_hvp(m,metID,metName)
if ~any(strcmp(m.mets,metID))
    m=addMetabolite(m,metID,'metName',metName,'metFormula','','Charge',0);
end
end

function m=padConstraintColumns_hvp(m)
n=numel(m.rxns);
if isfield(m,'C') && ~isempty(m.C) && size(m.C,2)<n
    m.C(:,end+1:n)=0;
end
end


%% ======================================================================
%% CONDITION SETUP / UPTAKE DIRECTION
%% ======================================================================

function m=prepareCondition_hvp(m,mode,cfg)
mode=upper(string(mode));
if mode~="AUTO" && mode~="MIXO" && mode~="HETERO"
    error('Unsupported HVP trophic condition: %s',mode);
end

for i=1:numel(cfg.carbonBoundaryRxns)
    r=cfg.carbonBoundaryRxns{i};
    if any(strcmp(m.rxns,r)), m=closeUptake_hvp(m,r); end
end
for i=1:numel(cfg.allMappedCarbonRxns)
    r=cfg.allMappedCarbonRxns{i};
    if any(strcmp(m.rxns,r)), m=closeUptake_hvp(m,r); end
end
if isfield(cfg,'reserveSourceRxn') && strlength(string(cfg.reserveSourceRxn))>0 && ...
        any(strcmp(m.rxns,cfg.reserveSourceRxn))
    m=closeUptake_hvp(m,cfg.reserveSourceRxn);
end
if isfield(cfg,'reserveMobilizationRxn') && strlength(string(cfg.reserveMobilizationRxn))>0 && ...
        any(strcmp(m.rxns,cfg.reserveMobilizationRxn))
    m=setDirectBounds_hvp(m,cfg.reserveMobilizationRxn,0,0);
end

basal=struct2cell(cfg.basalRxns);
for i=1:numel(basal)
    idx=find(strcmp(m.rxns,basal{i}),1);
    m.lb(idx)=cfg.basalMediumLB;
    m.ub(idx)=cfg.basalMediumUB;
end

idx=find(strcmp(m.rxns,cfg.ngamRxn),1);
m.lb(idx)=cfg.NGAM;
m.ub(idx)=cfg.NGAM;

if mode=="AUTO"
    m=setUptakeOnly_hvp(m,cfg.photonRxn,cfg.autoPhotonMaxUptake);
elseif mode=="MIXO"
    m=setUptakeOnly_hvp(m,cfg.photonRxn,cfg.mixoPhotonMaxUptake);
else
    m=setUptakeOnly_hvp(m,cfg.photonRxn,cfg.heteroPhotonMaxUptake);
end

m=setUptakeOnly_hvp(m,cfg.o2Rxn,cfg.o2MaxUptake);
if mode=="AUTO" || mode=="MIXO"
    m=setDirectBounds_hvp(m,cfg.o2DummyRxn,cfg.o2DummyLB,cfg.o2DummyUB);
else
    m=setDirectBounds_hvp(m,cfg.o2DummyRxn,0,0);
end

for i=1:numel(cfg.allAnalysisBOFRxns)
    r=cfg.allAnalysisBOFRxns{i};
    if any(strcmp(m.rxns,r)), m=setDirectBounds_hvp(m,r,0,0); end
end

if mode=="AUTO"
    b=cfg.AUTO;
elseif mode=="MIXO"
    b=cfg.MIXO;
else
    b=cfg.HETERO;
end

m=setDirectBounds_hvp(m,b.pigmentRxn,0,1000);
m=setDirectBounds_hvp(m,b.assemblyRxn,0,1000);
m=setDirectBounds_hvp(m,b.sinkRxn,0,1000);
m=setObjective_hvp(m,b.sinkRxn);

if mode=="AUTO"
    m=openUptake_hvp(m,cfg.co2Rxn,cfg.autoCO2MaxUptake,true);
    m=setDirectBounds_hvp(m,cfg.rubiscoRxn,cfg.rubiscoMinFluxAUTO,1000);
elseif mode=="MIXO"
    m=openUptake_hvp(m,cfg.co2Rxn,cfg.mixoCO2MaxUptake,true);
    m=setDirectBounds_hvp(m,cfg.rubiscoRxn,cfg.rubiscoMinFluxMIXO,1000);
else
    m=closeUptake_hvp(m,cfg.co2Rxn);
    m=setDirectBounds_hvp(m,cfg.rubiscoRxn,0,0);
end
end

function m=setDirectBounds_hvp(m,rxn,lb,ub)
idx=find(strcmp(m.rxns,rxn),1);
if isempty(idx), error('Reaction not found: %s',rxn); end
m.lb(idx)=lb; m.ub(idx)=ub;
end

function m=setObjective_hvp(m,rxn)
idx=find(strcmp(m.rxns,rxn),1);
if isempty(idx), error('Objective reaction not found: %s',rxn); end
m.c=zeros(numel(m.rxns),1);
m.c(idx)=1;
end

function m=clearObjective_hvp(m)
m.c=zeros(numel(m.rxns),1);
end

function m=closeUptake_hvp(m,rxn)
idx=find(strcmp(m.rxns,rxn),1);
if isempty(idx), return; end
sgn=uptakeFluxSign_hvp(m,rxn);
if sgn<0
    m.lb(idx)=max(m.lb(idx),0);
else
    m.ub(idx)=min(m.ub(idx),0);
end
end

function m=openUptake_hvp(m,rxn,q,preserveOtherDirection)
if nargin<4, preserveOtherDirection=true; end
idx=find(strcmp(m.rxns,rxn),1);
if isempty(idx), error('Reaction not found: %s',rxn); end
sgn=uptakeFluxSign_hvp(m,rxn);
q=abs(q);
if sgn<0
    m.lb(idx)=-q;
    if ~preserveOtherDirection, m.ub(idx)=0; end
else
    m.ub(idx)=q;
    if ~preserveOtherDirection, m.lb(idx)=0; end
end
end

function m=setUptakeOnly_hvp(m,rxn,q)
idx=find(strcmp(m.rxns,rxn),1);
if isempty(idx), error('Reaction not found: %s',rxn); end
sgn=uptakeFluxSign_hvp(m,rxn);
q=abs(q);
if sgn<0
    m.lb(idx)=-q; m.ub(idx)=0;
else
    m.lb(idx)=0; m.ub(idx)=q;
end
end

function sgn=uptakeFluxSign_hvp(model,rxn)
idx=find(strcmp(model.rxns,rxn),1);
if isempty(idx), error('Boundary reaction not found: %s',rxn); end
s=full(model.S(:,idx));
nz=find(abs(s)>1e-12);
if numel(nz)~=1
    error('Expected one-metabolite boundary reaction for uptake sign: %s',rxn);
end
if s(nz)<0, sgn=-1; else, sgn=1; end
end

function q=uptakeMagnitudeFromSolution_hvp(model,rxn,x)
idx=find(strcmp(model.rxns,rxn),1);
if isempty(idx), q=NaN; return; end
sgn=uptakeFluxSign_hvp(model,rxn);
q=max(0,sgn*x(idx));
end

function [alpha,k,cal]=calibrateHeteroPhoton_hvp( ...
    m,biomassRxn,pigmentRxn,glucoseRxn,nC,cfg)

m=openUptake_hvp(m,glucoseRxn,cfg.validationMaxCarbon/nC,false);

bidx=find(strcmp(m.rxns,biomassRxn),1);
m.lb(bidx)=cfg.referenceMu_h;
m.ub(bidx)=cfg.referenceMu_h;

pidx=find(strcmp(m.rxns,cfg.photonRxn),1);
m=clearObjective_hvp(m);
sgn=uptakeFluxSign_hvp(m,cfg.photonRxn);
m.c(pidx)=sgn; % objective equals positive photon uptake magnitude
sol=optimizeCbModel(m,'min');

if ~isFeasibleSolution_hvp(sol)
    error('HETERO photon calibration is infeasible at mu=%.6f h^-1.',cfg.referenceMu_h);
end

minPhoton=uptakeMagnitudeFromSolution_hvp(m,cfg.photonRxn,sol.x);
pigIdx=find(strcmp(m.rxns,pigmentRxn),1);
pigFlux=sol.x(pigIdx);

if pigFlux<=cfg.fluxTol
    error('HETERO pigment flux is zero during photon calibration.');
end

alpha=minPhoton/pigFlux;
k=minPhoton/cfg.referenceMu_h;

cal=table(cfg.referenceMu_h,minPhoton,pigFlux,alpha,k, ...
    'VariableNames',{'Calibration_mu_h','Minimum_photon_uptake', ...
    'Pigment_pool_flux','Photon_per_pigment','Photon_per_biomass'});
end

function m=addPhotonBiomassCoupling_hvp(m,photonRxn,biomassRxn,k)
pidx=find(strcmp(m.rxns,photonRxn),1);
bidx=find(strcmp(m.rxns,biomassRxn),1);
sgn=uptakeFluxSign_hvp(m,photonRxn);

row=sparse(1,numel(m.rxns));
row(pidx)=sgn;
row(bidx)=-k;

if ~isfield(m,'C') || isempty(m.C)
    m.C=sparse(0,numel(m.rxns));
    m.d=zeros(0,1);
    m.dsense=char(zeros(0,1));
end
if size(m.C,2)<numel(m.rxns)
    m.C(:,end+1:numel(m.rxns))=0;
end

m.C(end+1,:)=row;
m.d(end+1,1)=0;
m.dsense(end+1,1)='G';
if ~isfield(m,'ctrs') || isempty(m.ctrs), m.ctrs=cell(0,1); end
m.ctrs{end+1,1}='PUBHVP_HETERO_PHOTON_BIOMASS_COUPLING';
end


%% ======================================================================
%% CAPACITY SCREEN
%% ======================================================================


function tf=isFeasibleSolution_hvp(sol)
tf=isstruct(sol) && isfield(sol,'stat') && sol.stat==1 && ...
   isfield(sol,'x') && ~isempty(sol.x);
end
```
