## Supplementary Data 3 for "Experimental trophic flexibility guides a compartmentalized genome-scale model of red microalgae *Galdieria sulphuraria*": RUN_ALL_PUBLICATION_ANALYSES.m

```
% RUN_ALL_PUBLICATION_ANALYSES
% Executes publication analyses 1-4 sequentially and stops on the first error.

scriptsDir = fileparts(mfilename('fullpath'));
if isempty(scriptsDir), scriptsDir = pwd; end
supplementDir = fileparts(scriptsDir);
inputDir = fullfile(supplementDir,'Inputs');

requiredInputs = { ...
    'Galdieria_sulphuraria_SBML_Final.xml', ...
    'Model_Substrate_list.xlsx', ...
    'TOTAL_BIOMASS.xlsx'};

for i = 1:numel(requiredInputs)
    f = fullfile(inputDir,requiredInputs{i});
    if ~isfile(f)
        error('Required publication input is missing: %s',f);
    end
end

jobDirs = { ...
    '1_Galdieria_publication_growth_validation', ...
    '2_Galdieria_publication_substrate_optimisation', ...
    '3_Galdieria_publication_FVA', ...
    '4_Galdieria_publication_HVP'};

fprintf('\nGaldieria publication analyses\n');
for i = 1:4
    jobDir = fullfile(scriptsDir,jobDirs{i});
    if ~isfolder(jobDir)
        error('Required analysis folder is missing: %s',jobDir);
    end
    addpath(jobDir);
    fprintf('[%d/4] %s ... ',i,jobDirs{i});
    switch i
        case 1
            S1_Galdieria_publication_growth_validation();
        case 2
            S2_Galdieria_publication_substrate_optimisation();
        case 3
            S3_Galdieria_publication_FVA();
        case 4
            run(fullfile(jobDir,'S4_Galdieria_publication_HVP.m'));
    end
    fprintf('DONE\n');
end
fprintf('All four publication analyses completed successfully.\n\n');
```
