## Supplementary figures and images for "Experimental trophic flexibility guides a compartmentalized genome-scale model of red microalgae *Galdieria sulphuraria*"

### 4_Galdieria_publication_HVP_HETERO_fixed_growth_yields.png

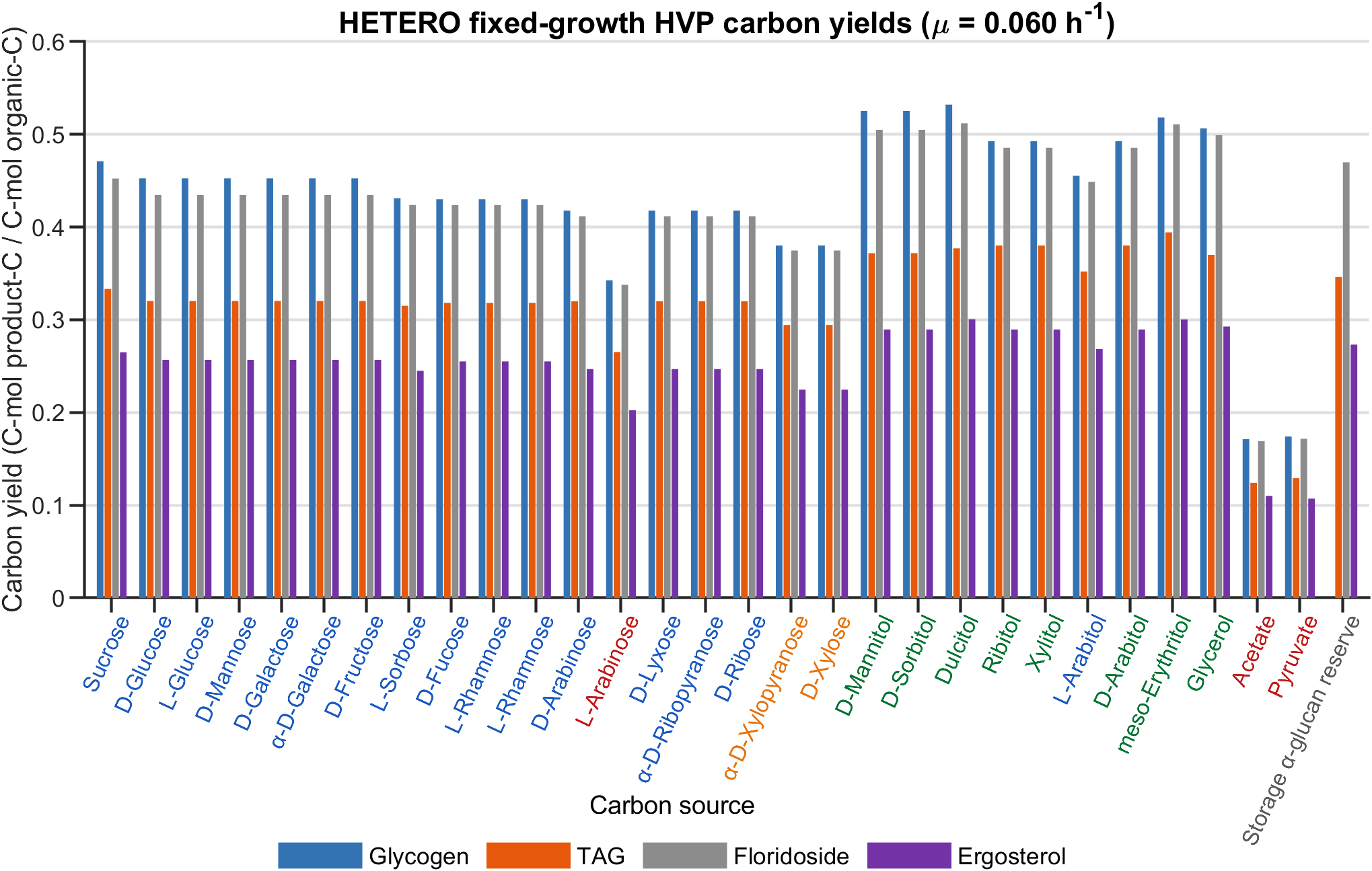

### 4_Galdieria_publication_HVP_HETERO_vs_glucose_heatmap.png

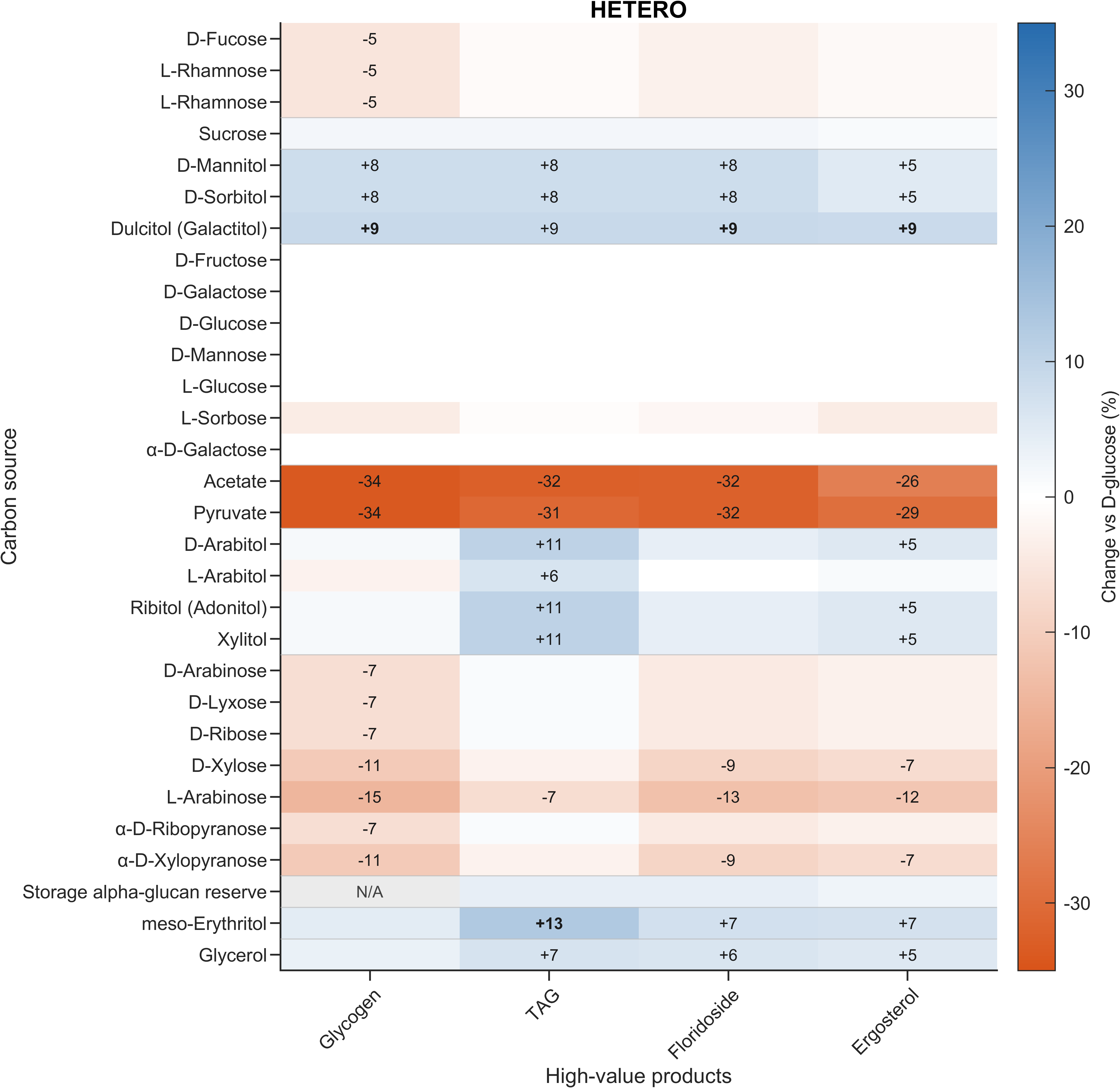

### 4_Galdieria_publication_HVP_MIXO_fixed_growth_yields.png

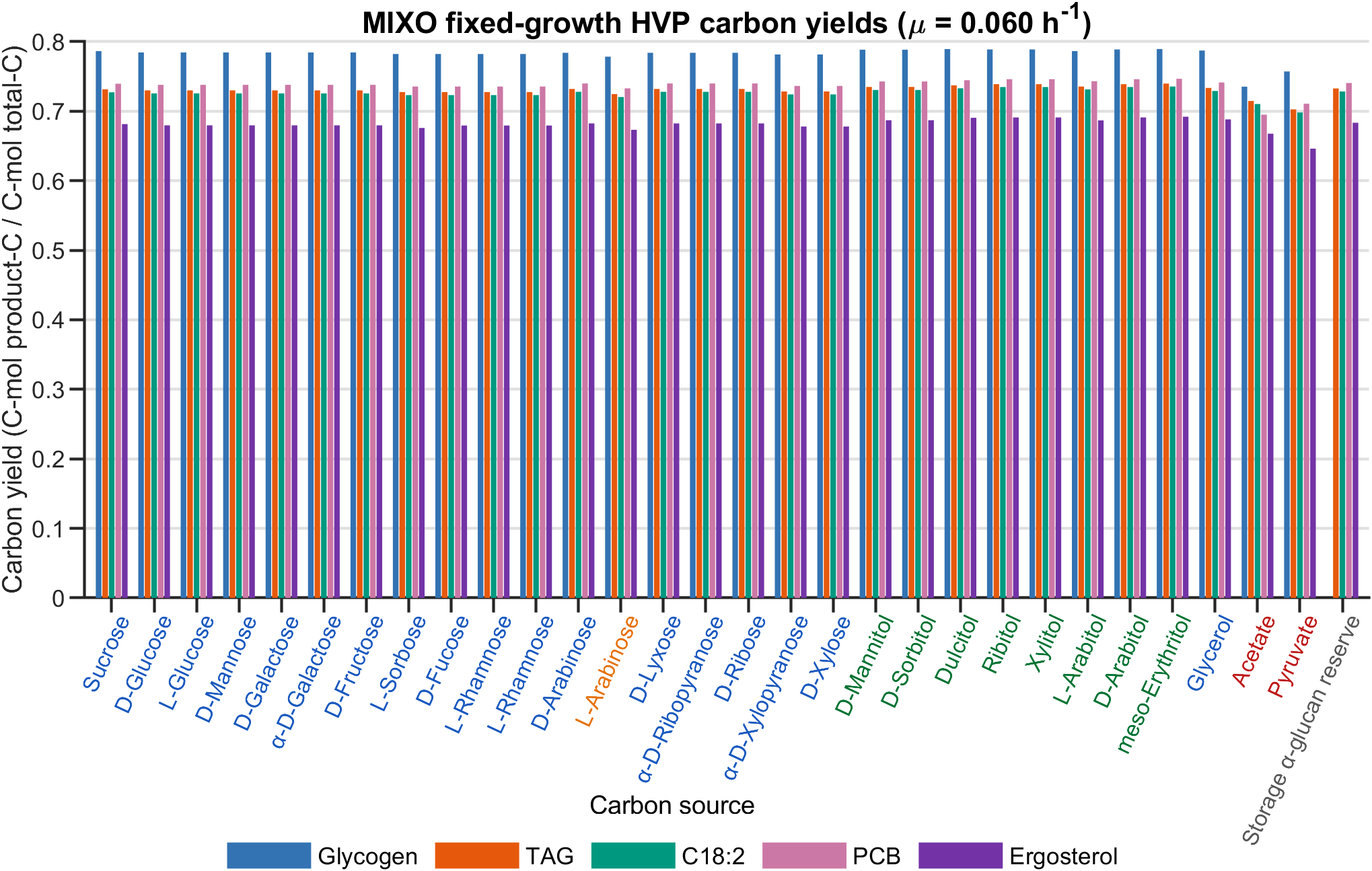

### 4_Galdieria_publication_HVP_MIXO_vs_glucose_heatmap.png

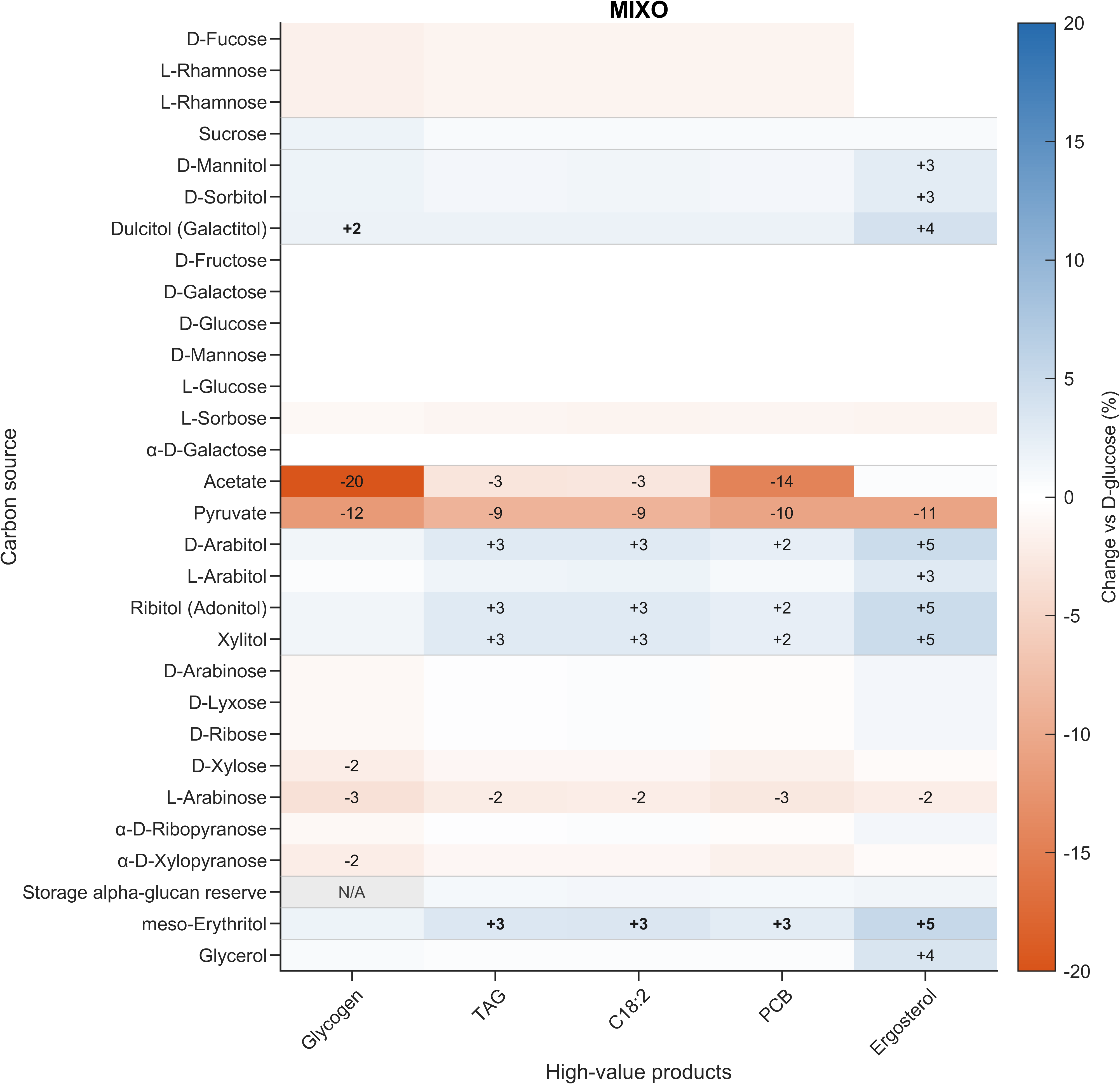
