## Supplementary Data 4 for "Experimental trophic flexibility guides a compartmentalized genome-scale model of red microalgae *Galdieria sulphuraria*": Supplementary_specific_growth_rate_methods.docx

Supplementary Methods and Data: Specific Growth Rates (μ) of *Galdieria sulphuraria*

Condition-specific literature dataset for autotrophic, mixotrophic and heterotrophic cultivation.

### 1. Dataset scope and organization

This supplementary dataset describes autotrophic, mixotrophic and heterotrophic growth-rate records for Galdieria sulphuraria. Each Excel row represents one reported or transparently derived condition-specific observation. The dataset excludes biomass concentration, productivity, unresolved records, model constraints and congeneric taxa. The source workbook is Mju_Biomass_data_final.xlsx.

Table S1. Dataset coverage and evidence composition.

| **Trophic state** | **Records** | **Minimum μ** | **Maximum μ** | **With uncertainty** | **Primary** | **Derived** | **Secondary** | **Operational** |
| --- | --- | --- | --- | --- | --- | --- | --- | --- |
| Autotrophic | 45 | 0,128 | 0,500 | 44 | 2 | 43 | 0 | 0 |
| Mixotrophic | 10 | 0,228 | 1,490 | 4 | 6 | 0 | 2 | 2 |
| Heterotrophic | 80 | 0,020 | 1,530 | 15 | 38 | 20 | 22 | 0 |

The numerical ranges summarize coverage only and must not be interpreted as pooled biological estimates, because strains, substrates, temperatures, pH values, cultivation modes and evidence types differ among records.

### 2. Growth-rate calculation and uncertainty

Directly reported μ values were retained in day⁻¹. For records reported as doubling time, td, in hours, the specific growth rate was calculated as:

**μ = ln(2) × 24 / td**

When the doubling-time standard deviation was available, uncertainty was propagated by first-order error propagation: SDμ = μ × SDtd / td. For steady-state chemostat or continuous-flow records, μ was equated to the reported dilution rate only when the evidence and validation note explicitly identify that operational interpretation. 'NA' indicates information that was not reported or could not be recovered with sufficient confidence.

The Excel field 'SD / uncertainty' is retained without re-labelling all entries as standard deviations: it may contain a reported SD, a propagated SD, or another source-defined uncertainty. The Evidence and Validation note columns identify the provenance and interpretation of each value.

### 3. Evidence classes and interpretation

**• Primary measurement:** A condition-specific μ value reported in the primary publication.

**• Operational steady-state measurement:** A dilution-rate value used as operational μ under a documented steady state; it is not automatically μmax.

**• Derived from reported doubling time:** μ calculated from a published doubling time using the equation above.

**• Secondary extraction:** A value recovered from a comparison table or secondary summary and retained with that limitation.

**• Grouped or replicate-specific result:** A publication-reported grouped average or individual-culture value that must not be treated as an independent population mean.

### 4. Supplementary growth-rate tables

The Excel file reproduces the current workbook records. μ and uncertainty values are displayed to three decimal places with decimal commas; underlying Excel values retain full numeric precision. DOI identifiers are given without the https://doi.org/ prefix.

### 5. DOI index

The following DOI identifiers occur in the retained autotrophic, mixotrophic and heterotrophic workbook records. The condition-level Excel rows remain the authoritative mapping between each measurement and its source.

10.1002/bit.20417 (4 records)

10.1002/elsc.202000075 (3 records)

10.1007/s00253-007-1150-2 (1 record)

10.1007/s10811-019-01957-9 (8 records)

10.1007/s10811-023-03012-0 (10 records)

10.1007/s12649-023-02036-4 (1 record)

10.1016/j.algal.2021.102513 (1 record)

10.1016/j.algal.2022.102752 (4 records)

10.1016/j.biortech.2017.04.043 (2 records)

10.1016/j.biortech.2021.125477 (7 records)

10.1016/j.biortech.2023.129281 (9 records)

10.1016/j.biortech.2024.130628 (2 records)

10.1016/j.biteb.2019.100269 (2 records)

10.1016/j.enzmictec.2005.05.010 (5 records)

10.1016/j.scowo.2025.100116 (2 records)

10.1039/C2FO30198A (43 records)

10.1080/09670269910001736322 (18 records)

10.3389/fmicb.2019.00927 (2 records)

10.3389/fpls.2022.978246 (2 records)

10.3390/su151914313 (1 record)

10.3934/environsci.2021028 (8 records)
