## Supplementary Data 5 for "Experimental trophic flexibility guides a compartmentalized genome-scale model of red microalgae *Galdieria sulphuraria*": GSM_COMP_KEGG_LINK_reading_guide.docx

**How to read GSM_COMP_KEGG_LINK_FINAL.xlsx**

*Guide to compartment-specific reactions, model provenance and database cross-references*

**Purpose.** This workbook is the publication-facing reference table for the final *Galdieria sulphuraria* genome-scale metabolic model. It contains 2,487 reactions and 2,270 compartment-specific metabolites parsed directly from Galdieria_sulphuraria_SBML_Final.xml; the final SBML also contains 1,050 gene products. Selected provenance fields were retained from the previous workbook only when they could be matched safely.

**1. Workbook structure**

| **Worksheet** | **What it contains** | **How it should be used** |
| --- | --- | --- |
| Reaction List | Final compartment-specific reactions, equations, bounds, GPRs, pathways, database identifiers and selected provenance. | Primary sheet for reviewing the implemented reaction network and its biological annotation. |
| Metabolite List | Compartment-specific metabolite identifiers, names, charged formulae and chemical cross-references. | Use to interpret metabolites appearing in reaction equations. |
| Data Dictionary | Dataset scope, final SBML source, model counts, default objective, column definitions, missing-value rule and compartment key. | Consult first when a field, abbreviation or objective-handling rule is unclear. |

**2. Reading the Reaction List**

**Read each row from left to right.** Reaction ID, stoichiometric equation, compartment and flux bounds describe the implemented final model; annotation and provenance columns provide biological context and origin.

| **Column group** | **Fields** | **Interpretation** |
| --- | --- | --- |
| Identity and chemistry | Reaction ID; Reaction name; Stoichiometric equation; Compartment | The final model implementation. Suffixes and metabolite tags identify localization; arrows show directionality. |
| Flux constraints | Lower bound; Upper bound | Allowed flux range. A negative lower bound normally permits reverse flux; 0-1000 is forward-only. |
| Biological annotation | GPR; Subsystem; EC number; KEGG reaction ID | Gene association, pathway assignment and external enzyme/reaction cross-references. |
| Provenance and curation | Source-model reaction ID; Implementation source; Notes | Documents safely retained source-model mapping, gene/DOI/source evidence, gap-filled status and curation notes. |

**Authoritative fields.** For the current model, use Reaction ID, Stoichiometric equation, Compartment, Lower bound and Upper bound from this workbook. Source-model reaction ID, Implementation source and Notes are provenance/context fields and do not override the final SBML implementation.

**3. Compartment and equation notation**

| **Code or notation** | **Meaning** | **Typical identifier tag** |
| --- | --- | --- |
| CYTO | Cytosol | _Cyto and [c] |
| MITO | Mitochondrion | _Mito and [m] |
| PLAS | Plastid | _Plas and [p] |
| THYL | Thylakoid | _Thyl and [t] |
| PERO | Peroxisome | _Pero |
| EXTRACELLULAR | Extracellular space | extracellular identifier |
| A <=> B | Reversible or transport reaction connecting listed participants/compartments | participant identifiers carry compartment tags |
| A -> (sink) | Sink, demand, exchange or one-compartment transport-style reaction | often one-sided equation |

**Equation arrows.** "->" indicates the implemented forward direction and "<=>" indicates reversibility. **Bounds remain decisive:** always interpret the equation together with the numerical lower and upper bounds.

**4. Reading the Metabolite List**

| **Field** | **Meaning** |
| --- | --- |
| Metabolite ID | Unique model identifier, including its compartment-specific suffix/tag. |
| Metabolite name | Human-readable chemical name; alternative names may be separated by “//”. |
| Charged formula | Formula used for the represented charged metabolite state. |
| Compartment | Full-name localization corresponding to the model identifier. |
| ChEBI ID | External Chemical Entities of Biological Interest cross-reference. |
| InChI | International Chemical Identifier, where available. |

**5. Interpretation rules and quality notes**

- Blank cells mean that information was not available or could not be transferred unambiguously; blank does not mean zero.
- GPR expressions use AND/OR logic: AND indicates jointly required gene products, whereas OR indicates alternatives or isoenzymes.
- Implementation source may contain a gene ID, DOI, source label (for example GAP FILLED), or another provenance note.
- Source-model reaction ID, Implementation source and Notes document provenance where a safe match was available; blank provenance fields should not be interpreted as absence of biological support.
- Reaction compartments are derived from the final reaction and metabolite identifiers. Transport reactions list all participating compartments.
- The workbook is a scientific supplementary reference, not the executable model. The final SBML is authoritative for reaction structure, bounds, gene products and the default objective. The workbook intentionally omits a static Objective coefficient column because Scripts 1-4 assign biomass or product objectives dynamically for each analysis.

**6. Recommended reading sequence**

**1)** Locate the Reaction ID; **2)** verify equation, compartment and bounds; **3)** review GPR, subsystem and KEGG/EC links; **4)** check provenance fields when the reconstruction source is relevant; **5)** use the Data Dictionary for dataset scope, missing-value rules and the default SBML objective.
